# Programming Biomolecular Interactions with All-Atom Generative Model

**DOI:** 10.64898/2026.03.12.711044

**Authors:** Xiangzhe Kong, Junwei Chen, Ziting Zhang, Gaodeng Li, Qingyuan Zhu, Lei Wei, Mingyu Li, Yan Shi, Weiyang Dai, Zishen Zhang, Wenjuan Tan, Rui Jiao, Xiaolun Wang, Jiqing Zheng, Ziyang Yu, Qilong Wu, Zhiye Guo, Li Zhang, Wentao Li, Qiaojing Huang, Tian Zhu, Xiaowo Wang, Wenbing Huang, Yuli She, Jian Zhang, Yang Liu, Kai Liu, Jianzhu Ma

## Abstract

Biomolecular interactions lie at the core of cellular life, spanning diverse molecular modalities from small molecules to nucleic acids and proteins. Nevertheless, design strategies remain separated despite shared physicochemical principles of molecular recognition. Here we present AnewOmni, a unified generative framework trained on more than 5 million biomolecular complexes, that enables transferable molecular design across molecular scales by assembling chemically meaningful building blocks at atomic resolution. We further introduce programmable graph prompts to support user-defined chemical, topological, and geometric steering during generation, exploring hybrid and unconventional chemistries beyond canonical structures. We demonstrate that transferable learning of interaction patterns and physical constraints across molecular modalities is possible, via an atom-to-block latent space capturing both atomic details and structural priors. The framework successfully designed small molecules, peptides, and nanobodies targeting the challenging KRAS G12D switch II pocket, as well as orthosteric peptides and allosteric small-molecule inhibitors for PCSK9 in the absence of known binding site, achieving 23%-75% success with only low-throughput validation, bypassing modality-specific high-throughput screening. AnewOmni is the first to succeed in functional molecular design across all scales, from small chemical entities to large biologics, and represents a stepstone towards general molecular reasoning engines, advocating a generative foundation model for biomolecular interactions to enter regimes where data and human intuition remain limited.

## Introduction

Biomolecular interactions underlie cellular function and constitute a fundamental basis of life. Throughout human history, diverse classes of molecules, including small organic molecules, proteins, and nucleic acids, etc., have been developed to modulate these interactions and thereby control biological processes^1–4^. However, such target-binding molecules are typically designed using distinct representations, rules, and protocols^5–7^, reflecting historical divergences in chemical and biological technologies rather than fundamental mechanisms of molecular recognition. Indeed, it is the atomic-scale interaction geometry that constitutes the primary determinants of biological recognition, thus advocating the potential of a unified, modality-agnostic design framework for regulating biomolecular interactions. Ideally, such a unified methodology should encourage transferability of the shared physical design principles to 1) enhance generation of each molecular modality, and to 2) enable exploration of hybrid or unconventional chemistries beyond training data.

While divergences in wet-lab experimental toolkits may be difficult to reconcile, recent advances in computational molecular design offer an alternative path forward. Structure-based generative models have demonstrated considerable promise in designing target-binding molecules across diverse fields, including small molecules^8^, peptides^9^, mini-proteins^10,11^, and antibodies^12^. However, most existing approaches continue to rely on modality-specific representations and model architectures, which limit their extensibility to other modalities or even unconventional chemistries such as non-canonical amino acids, cyclic scaffolds, or post-translationally modifications. As a result, current methods still struggle to provide a modality-agnostic and customizable framework for molecular binder design, particularly when extrapolating beyond the distributions observed during training.

Although models like AlphaFold3^13^ and Boltz2^14^ exhibit remarkable accuracy in joint modeling diverse biomolecular complexes, their success is largely confined in structure prediction rather than generative molecular design. Recent efforts on generative approaches are exploring this direction, yet achieving generalizable cross-modality design remains challenging. One line of work (e.g., BoltzGen^15^ and ODesign^16^) represents molecules at separate granularity, with large molecules like proteins in tokens of amino acids, and small molecules in tokens of atoms. During generation, they first generate coordinates of representative atoms in tokens, such as C_↵_, and subsequently infer token identities via inverse folding. Such methodology limits the transferability across modalities since the atomic geometries, where the shared physical principle lies, are not explicitly considered during generation. Another line of work like PocketXMol^17^ treats all modalities as purely atomic graphs. While this formulation captures fine-grained interactions, it neglects higher-order structural priors shaped by biological evolution, such as conserved repeating patterns in amino acids and nucleobases. Consequently, it is largely restricted to small molecules and short peptides, with limited generalizability to larger assemblies such as antibodies. Together, these limitations highlight a central open question: *Is there a generative framework unifying molecular modalities in a manner that preserves transferable atomic-level interaction principles while retaining modality-specific structural priors required for molecular design across vastly different scales?*

In pursuit of this goal, we present AnewOmni, an atom-scale generative foundation model trained on over five million biomolecular complexes curated from the literature. Given a target binding site, it is capable of designing binders across diverse molecular modalities by assembling atomic building blocks mined from data. Atomic details are explicitly encoded into the latent points of building blocks through a trained all-atom variational autoencoder, enabling efficient latent diffusion that simultaneously preserves transferable atomic interaction geometries and heuristic structural priors during generation. Such modeling principle is also evidenced by successful expert experience, where reusable chemical building blocks were transfered and adapted across molecular modalities^18,19^. The model is further equipped with programmable graph-based prompts that enable explicit control over chemical composition, topology, and structural constraints, such as cyclization, incorporation of non-canonical amino acids (ncAAs), covalent interaction, and molecular growing on existing scaffolds, supporting flexible customizations even beyond the training data.

Extensive *in silico* benchmarks across small molecules, peptides, and antibodies demonstrate that unified atomic modeling improves physical plausibility within each modality, achieving superior performance over state-of-the-art modality-specific generative models. Further analysis indicates that the model learns to transfer generalizable interaction patterns, maintaining critical interactions across different molecular modalities. Consistent with this property, AnewOmni generalizes to previously unseen scenarios, successfully generating physically favorable binders for targets such as ribonucleic acid (RNA), deoxyribonucleic acid (DNA), and glycans. On recently published RNA–small-molecule datasets with experimental validation^20^, our model exhibits zero-shot discriminative signal between binding and non-binding pairs through generative likelihood estimation, indicating strong generalization considering the absence of RNA-related data during training.

As experimental proof of concept, we applied AnewOmni to design binders across distinct modalities yet targeting the same functional site on kirsten rat sarcoma viral oncogene homolog (KRAS) G12D, a long-standing and challenging therapeutic target^21^. Inspiringly, the model successfully identified de novo small molecule, peptide, and nanobody binders, achieving success rates of 23%-75%, without reliance on high-throughput display technologies. Notably, the new small molecule inhibitors adopt novo scaffolds, with the similarity to MRTX-1133^22^, the well known inhibitor, as low as 0.12-0.15. Next, AnewOmni identified peptides targeting the orthosteric binding site on proprotein convertase subtilisin/kexin type 9 (PCSK9) for disrupting its interaction with the low-density lipoprotein receptor (LDLR), which is a key regulator of cholesterol homeostasis^23^. Furthermore, the model discovered small molecules that bind a cryptic allosteric pocket of PCSK9 and modulate its activity potentially by inhibiting the secretion, leading to increased LDLR function, as evidenced by cellular assays. Remarkably, high-resolution crystal structures confirmed that the experimentally observed binding pose of the small molecule closely matches the pose generated by AnewOmni, with a root mean square deviation of 0.92 Å. Together, these results highlight the potential of all-atom generative foundation models to unify design of molecular binder specific to target binding sites across traditionally separate paradigms. To the best of our knowledge, this work provides the first evidence that, with an appropriate representation granularity, joint generative training across molecular modalities can yield mutual performance gains and improved generalization beyond the training distribution. We are also the first to demonstrate experimental success of binders in modalities spanning diverse scales, from small entities (e.g., small molecules) to large biologics (e.g., nanobodies) within a single unified framework.

## Results

### Unified Generation across Molecular Modalities

We aim to enable molecular generation across diverse modalities within a single unified framework. Building upon atom-scale resolution which supports generalizable interaction learning, we further organize atoms into atomic building blocks to maintain well-structured chemistry (Fig. 1a), such as amino acids, nucleotides, or molecular fragments mined from data (Supplementary Note 1.2.1). Generation is performed by operating on these building blocks within a shared geometric latent space conditioned on the atomic environment of the target binding site (Fig. 1b). The model first learns a reversible mapping between all-atom geometries and the coarse-level latent space, where atoms are compressed into their building blocks by context-aware embeddings that summarize their local atomic structures and environments. Molecular generation is then formulated as a diffusion process operating on latent point clouds to support scalable and efficient generation (Methods). The model is trained on a large-scale dataset comprising over five million pairs of binding sites and binders curated from the literature, spanning small molecules, peptides, and antibodies (Fig. 1c). This scale provides high diversity, which encourages the model to learn a wide spectrum of atomic interaction patterns across modalities. We further introduce programmable graph prompts to steer the generative process towards user-defined conditions. This mechanism associates latent points or their edges with specified chemical entities and spatial arrangements, driving them towards the desired chemical topology and three-dimensional geometry (Supplementary Note 1.3.3). It allows flexible customization of generation for a wide range of design objectives, including enforcing specific chemical motifs such as disulfide bonds or non-canonical amino acids, constraining overall geometry via cyclization, directing interaction growth from existing molecular scaffolds, etc. (Fig. 1d).

**Figure 1.**
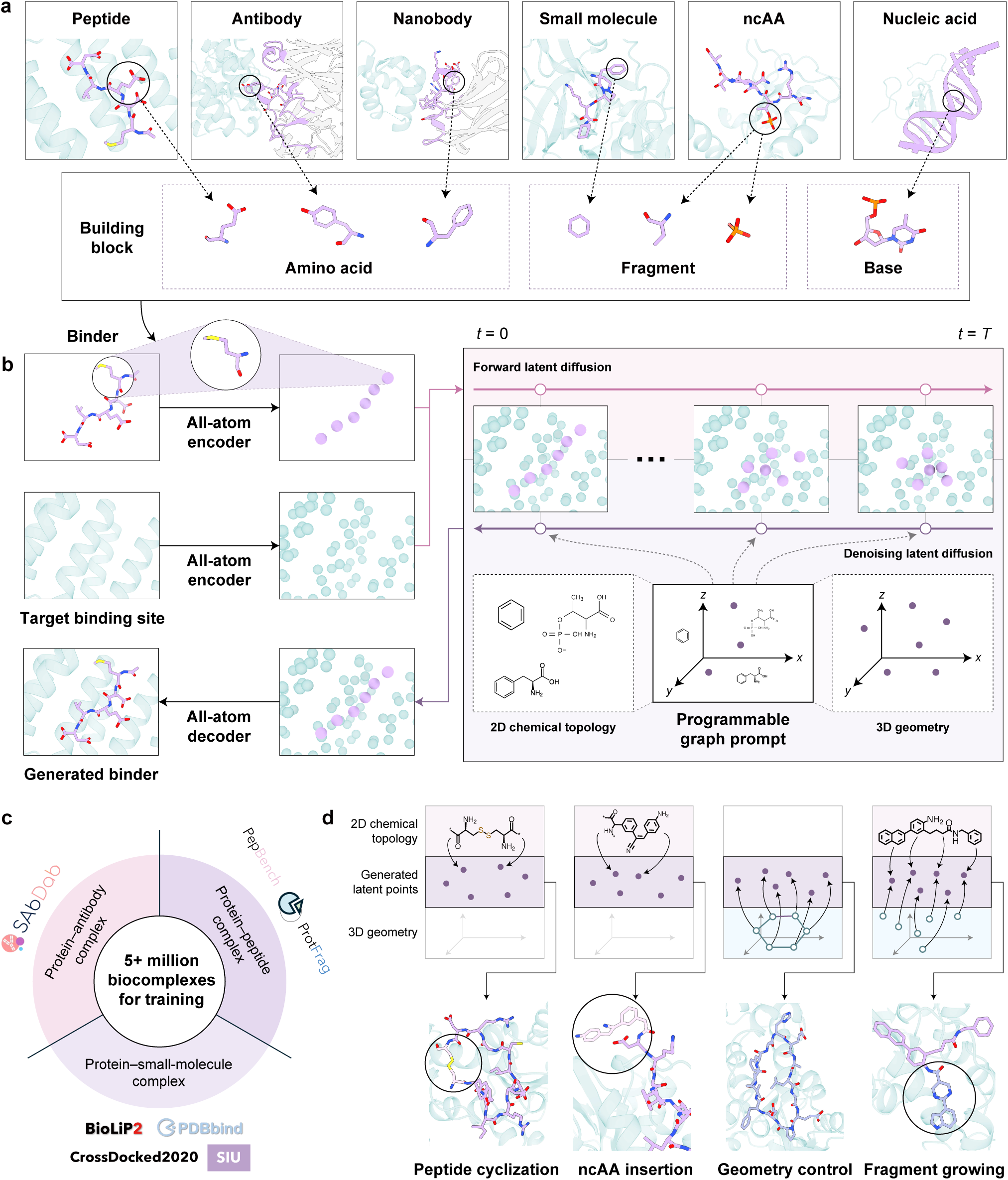
Overview of AnewOmni. The model unifies representations of diverse molecular modalities into building blocks with atomic resolution and supports flexible, programmable control during generation. **a**, Atoms from different molecular modalities are organized into chemically coherent building blocks, including canonical units such as amino acids and nucleobases. Other chemical entities, such as small molecules and non-canonical amino acids (ncAAs), are decomposed into frequent fragments mined from ChEMBL^24^. **b**, The architecture of AnewOmni combines an all-atom autoencoder with a latent diffusion model. The autoencoder establishes a reversible mapping between all-atom geometries and low-dimensional block-level latent representations, while the diffusion model is trained to generate plausible latent point clouds. During inference, generated latent points are decoded back into all-atom geometries via the autoencoder. **c**, AnewOmni is trained on large-scale, multi-source datasets spanning protein complexes with small molecules, peptides, and antibodies, comprising over five million biocomplexes in total. **d**, AnewOmni enables customized generation through programmable graph prompts. Chemical, topological, and geometric constraints are injected as node- or edge-level prompts via chemical graphs or spatial coordinates, allowing fine-grained control over generation. This mechanism supports diverse downstream applications, such as peptide cyclization via the prompt of a disulfide bond, ncAA insertion through the prompt of its chemical structure, macrocycle conformation control using geometric constraints, and molecular growth by prompting part of the latent point cloud with the scaffold while allowing the remainder to be generated freely.

We evaluated the unified architecture on established public benchmarks to enable fair comparison with state-of-the-art, modality-specific generative models. Complementary metrics reflecting binding energy from statistical force fields, physiochemical validity, and distribution fidelity were normalized and aggregated into a composite score between 0 and 1 (Supplementary Note 2.4). Across small molecules, peptides, and antibodies, AnewOmni consistently surpassed specialized baselines as well as its counterparts trained on single-modality data, demonstrating that joint training across molecular modalities inspiringly enhances rather than compromises generative performance (Fig. 2a). Visualization of the block-level latent space further illustrates interpretable patterns, with hydrophobic and polar amino acids separated by clear boundaries, while molecular fragments blend in with amino acids (Fig. 2b). Notably, molecular fragments that resemble amino acid sidechains cluster closely in the latent space: the thiomethyl group aligns with cysteine, hydroxymethyl with serine, acetamido with asparagine, ethanoic acid with aspartic acid, and hydroxyethyl with threonine (Fig. 2b). This organization reflects a meaningful latent representation that captures the functional similarity of diverse building blocks in molecular recognition. To investigate how cross-modality training shaped learned interaction patterns, we extracted eight recently released targets (Supplementary Note 3.1) for AnewOmni to generate binders from multiple molecular modalities targeting the same binding sites. We detected inter-molecular interactions with ProLIF^28^ for generated and native complexes, and found 50%-70% native interactions could be recovered from generated complexes, indicating that the model reliably reproduced key contact patterns regardless of molecular modalities (Fig. 2c, left). Peptides exhibited the greatest interaction flexibility, achieving the highest recovery rates while also introducing the most novel contacts. With cross-modality training, all modalities showed enhanced recovery and novel interactions, with small molecules and antibodies benefiting most substantially—recovering about 10% more native interactions than their single-modality counterparts. Interestingly, this improvement may arise from mutual knowledge transfer between antibody and small-molecule data, with peptides acting as an intermediate bridge. Consistent with this interpretation, cross-modality training increased the reliance of small molecules on hydrophobic interactions, which should be more characteristic of antibody recognition, whereas antibodies displayed the opposite shift, and peptides showed minimal changes in interaction preference (Fig. 2c, right). Explicit visualization of small molecules, peptides, and antibodies bound to the same target (PDB ID: 9DMV), together with interaction statistics, showed that shared interactions largely recovered native ones (e.g., Arg23 and His72). In contrast, modality-specific novel interactions displayed distinct preferences, such as Arg30 for small molecules and Arg64 for antibodies, whereas peptides exhibited the broadest interaction coverage, spanning nearly all observed contacts (Fig. 2d, Supplementary Fig. 2).

**Figure 2.**
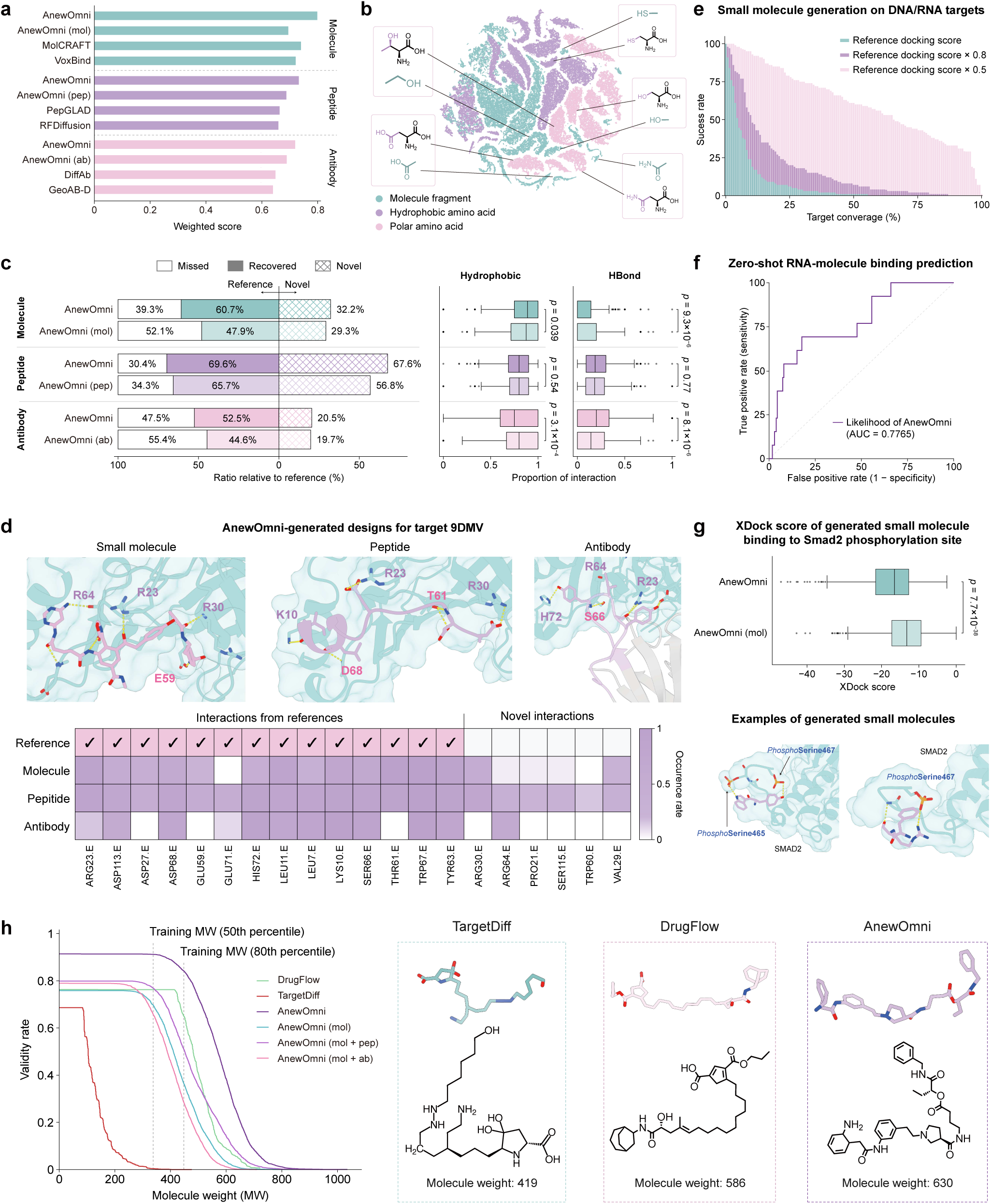
In silico analysis of AnewOmni. A comprehensive evaluation is conducted to assess physical validity, computational binding energy, interaction fidelity, and cross-modality generalization of designs given by AnewOmni, with focus on both in-distribution and out-of-distribution settings. **a**, Overall weighted score aggregating multiple *in silico* metrics from public benchmarks for *de novo* design of small molecules, peptides, and antibodies (Supplementary Note 2.4). AnewOmni (mol), AnewOmni (pep), AnewOmni (ab) are trained solely with molecules, peptides, and antibodies, respectively. **b**, Latent space of building blocks visualized using t-SNE and colored by block type, including molecular fragments, hydrophobic and polar amino acids. Adjacent molecular fragments and amino acids are annotated with their chemical structures, with backbones and sidechains of amino acids marked in different colors. **c**, Left: distribution of interaction types generated by small molecules, peptides, and antibodies compared with native binders, showing the proportions of missed, recovered, and novel interactions. Results from AnewOmni trained on single-modality data are included to assess the impact of cross-modality training. Right: differences in hydrophobic interactions and hydrogen bonds (HBonds) generated by AnewOmni under joint training versus single-modality training. **d**, Visualization of small molecule, peptide, and antibody binders generated for the same target binding site (PDB ID: 9DMV). Interaction statistics across modalities are compared with the native binder, with deeper colors indicating higher generation frequency. **e**, Success rates of small-molecule generation on 139 DNA/RNA targets, with binding energies evaluated using XDock^25^. Multiple success thresholds are reported, defined as achieving 100%, 80%, or 50% of the native binder’s binding energy. **f**, Zero-shot prediction of RNA-binding small molecules via generative likelihood on a newly reported experimental assay with binary binding labels^20^. **g**, Distribution of XDock scores for small molecules generated against a phosphorylation site, along with a representative structural visualization. **h**, PoseBusters validity of generated small molecules versus molecular weight. Models trained on small molecules only (mol), small molecules with peptides (mol+pep), and small molecules with antibodies (mol+ab) are compared to assess the contribution of each modality. The 50th (339.2) and 80th (444.3) percentiles of molecular weight in the training set are indicated on the axis. Representative molecules with high molecular weights from TargetDiff^26^, DrugFlow^27^, and AnewOmni are visualized for comparison.

We next assessed the ability of our model to generalize on distinct scenarios completely unseen during training. We collected 139 RNA/DNA targets from the literature and generated small-molecule binders, which were evaluated using XDock^25^, a physics-based energy function supporting nucleic acid-ligand and non-canonical protein target docking (Supplementary Note 3.2). For nearly all targets, AnewOmni generated at least one candidate with binding energy exceeding 50% of the corresponding native binder within 100 generations, with success rates of 60% or higher observed in 50%–60% of cases. Under a more strict threshold of 80% of native binding energy, 70%-80% of targets still yielded satisfactory candidates, and for a substantial fraction (50%-60%) of targets, the model could produce strong binders even with energies comparable to native ligands (Fig. 2e). Similar trends were also observed when designing peptides targeting these RNAs/DNAs (Supplementary Fig. 3a). To further challenge the model, we evaluated zero-shot generalization on a recently published RNA–small-molecule dataset with binary binding labels validated experimentally^20^ by controlling AnewOmni to generate exactly the same molecules in the dataset for likelihood estimation (Supplementary Note 3.3). Without having seen any RNA–small-molecule data during training, the generative likelihood produced by AnewOmni achieved an area under the receiver operating characteristic curve (AUC–ROC) of 0.77 in distinguishing binding from non-binding ligands (Fig. 2f). Beyond nucleic acids, we assessed generalization to biologically relevant yet less regular targets, including glycans and phosphorylation sites (Supplementary Note 3.4). In both cases, AnewOmni generated physically favorable peptide and small-molecule binders in a zero-shot setting as evaluated by XDock, exhibiting significant advantages over the single-modality counterparts (Fig. 2g, Supplementary Fig. 3c-e). Together, these results indicate that unified all-atom generative modeling enables robust generalization to diverse and previously unseen molecular interaction landscapes. Beyond interactions, AnewOmni maintained high physical validity as molecular size increases, even at elevated molecular weights far exceeding conventional small molecules (Fig. 2h), outperforming literature baselines^26,27^ as well as ablated variants trained solely on small-molecule data. Further addition of either peptide or antibody data to the small-molecule variant of AnewOmni led to the conclusion that such advantages came from both peptides and antibodies (Fig. 2h). This capability of generating medium- to large-sized molecules are particularly desirable for targeting extended and binding sites, which are common for regulators of protein–protein interactions.

### Customized Generation through Programmability

Beyond unified generation across molecular modalities, a central requirement for practical molecular design is the flexibility to exert precise and customizable control over the generation process, such as molecular topology, chemical composition, interaction patterns, and extensions from existing scaffolds. Our model supports such programmability through graph-based prompts that encode different formats of chemical and spatial constraints as node-level or edge-level conditions on the latent point clouds, directly steering the generative process (Fig. 1d, Supplementary Note 1.3.3). Specifically, users can assign arbitrary chemical structures to designated latent points as node labels or define their connectivity as edge labels, while spatial coordinates could also be specified to guide approximate positioning. These elements can be composed into more complicated graphs for prompted generation. This abstraction is highly expressive, allowing diverse design requirements to be combined and modified flexibly in a single representation system without retraining.

We started by topological control, a common requirement which is receiving growing attention in drug discovery, especially in peptide design^4^. Turning linear topology into cyclic ones in peptides encourages rigidity by restricting the accessible conformational space, and thus potentially enhances properties such as stability and binding affinity^29^. The simplest form of cyclization connects the free amino and carboxyl groups at two ends of the peptide via an amide bond. By explicitly specifying this connection as an edge-level prompt between the terminal building blocks, AnewOmni managed to generate chemically valid head-to-tail cyclic peptides with favorable target interactions (Fig. 3a). Beyond backbone cyclization, disulfide bonds provide an alternative and widely used mechanism for stabilizing peptide topology when at least two cysteines exist^29^. This is also achievable for our model by constraining the presence of cysteine residues through node-level prompts and enforcing a disulfide linkage via an edge-level prompt. Programmed by these prompts, AnewOmni successfully generated valid candidates with cyclic structures built by disulfide bonds (Fig. 3b).

**Figure 3.**
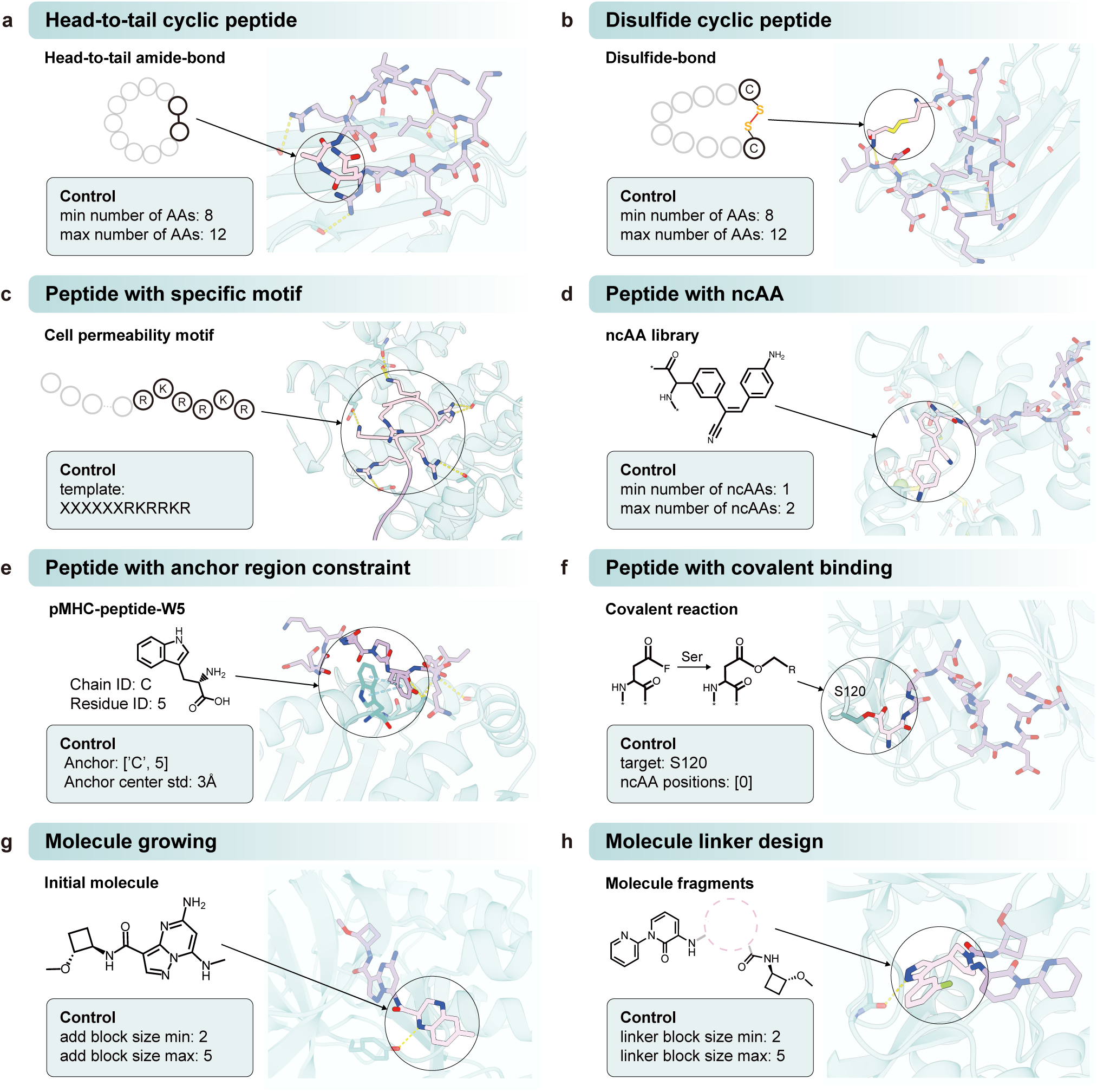
Programmable design with customized controls. AnewOmni supports flexible incorporation of chemical, topological, and geometric control during molecular generation. **a**, Head-to-tail cyclic peptide generation with a specified number of amino acids (AAs) by enforcing an amide bond between the N- and C-terminus. **b**, Disulfide cyclic peptide generation with a specified number of AAs by enforcing two cysteine residues connected by a disulfide bond. **c**, Peptide generation incorporating a predefined cell-permeability motif by fixing part of the sequence template to the specified motif. **d**, Peptide generation with a specified number of non-canonical amino acid (ncAA) insertions sampled from a user-defined library. **e**, Peptide generation guided to form interactions with a specified anchor residue on the binding site. **f**, Peptide generation forming a covalent bond with a specified target residue through incorporation of a reactive ncAA. **g**, Scaffold-based molecular growth to extend an existing molecule into deeper regions of the binding site with desired size. **h**, Linker design connecting two predefined fragments within the binding site.

Another critical requirement is control over chemical composition during generation. In many design scenarios, specific sequence motifs are known to correlate with desirable properties, such as cell permeability^30^. By encoding such motifs as node-level prompts, AnewOmni preserved the specified chemical pattern while simultaneously optimizing its spatial placement relative to the target binding site, enabling favorable interactions to be formed around the constrained motif (Fig. 3c). A more challenging setting involves incorporation of user-defined libraries of non-canonical amino acids (ncAAs), a widely adopted strategy for exploring unconventional chemical space and addressing otherwise intractable design problems^31,32^. As an illustrative example, we included a fluorescent ncAA whose emission is activated only under conformational restriction^31^ and applied the model to calmodulin, which undergoes a well-characterized structural transition upon Ca^2+^ binding^33^. By generating peptides that preferentially target the Ca^2+^-bound conformation with the ncAA, the methodology could, in principle, support the discovery of Ca^2+^ sensors (Fig. 3d).

We next demonstrated control over desired interaction patterns during molecular generation. In many biological systems, functional binding requires engaging specific anchor regions on the target surface, such as in recognition of peptide–major histocompatibility complex (pMHC) proteins^34^. To fulfill such requirements, we imposed node-level spatial prompts on the latent points, biasing their placement toward user-defined anchor regions with adjustable spatial tolerance. Under these constraints, AnewOmni could yield binders that form interactions at the specified locations while maintaining overall structural compatibility with the binding site (Fig. 3e). Beyond non-covalent interactions, some applications require covalent bonding to achieve specificity or durability. As an illustrative design scenario, we considered TNF-like ligand 1A (TL1A), where selective inhibition of its interaction with death receptor 3 (DR3), but not the closely related decoy receptor 3 (DcR3), is desirable^35^. Prior studies identified residue Ser120 as a key differentiating site^35^. By specifying a reactive non-canonical amino acid and enforcing a covalent bond between the designed peptide and Ser120 through node-level and edge-level prompts, respectively, AnewOmni generated peptide binders that satisfied the prescribed covalent interaction geometry (Fig. 3f).

Finally, we showed that the programmability of AnewOmni naturally extended to extension of existing molecules, a common task in structure-based functional molecule design. By constraining a subset of latent points to preserve an existing scaffold with node-level and edge-level prompts, the model was able to generate and extend additional fragments either freely or with directed growth toward specified regions of the binding site. In the given example (Fig. 3g), the newly grown fragment extends deeper into the pocket and forms a hydrogen bond with a nearby tyrosine residue. This algorithm also supports linker design just with minor changes that the "scaffold" is now two separate functional fragments already placed in the binding site. AnewOmni designed a linker featuring rigid aromatic rings for connection and further grew one fragment to form a hydrogen bond with a deeper asparagine residue in the binding site (Fig. 3h).

Although the above-mentioned scenarios represent only a preliminary of the diverse requirements encountered in real-world applications, together they demonstrate how the programmability of AnewOmni enables flexible and precise customization for structure-based molecular discovery.

### Targeting the Switch II Site on KRAS with Different Modalities

Cancer remains one of the leading causes of death worldwide^36^. Among oncogenic drivers, Kirsten rat sarcoma viral oncogene homolog (KRAS) mutations occur in over 30% of human cancers, with the G12D mutation representing the most prevalent KRAS variant^22,37^. Nevertheless, owing to its high affinity for guanosine diphosphate (GDP) and guanosine triphosphate (GTP), together with the absence of deep or persistent ligandable pockets, KRAS has been regarded as a prototypical “undruggable” target until recent years^22^. To this end, we challenged our model to design small molecules, peptides, and antibodies targeting the same switch II pocket of GDP-bound KRAS G12D, enabling a direct comparison of cross-modality design within a single binding site.

We started by designing small-molecule binders, which remains exceptionally challenging due to the shallow binding pocket and the requirement for large scaffolds beyond common sizes to reach the scattered key residues. Most prior efforts have therefore focused on optimizing derivatives of the breakthrough inhibitor MRTX1133^22^, rather than exploring fundamentally new chemotypes^38,39^. However, the clinical translation of this pyrido[4,3-*d*] pyrimidine scaffold has been hindered by its low bioavailability^40^. Here, we instead leveraged AnewOmni to perform *de novo* small-molecule design targeting the Switch II pocket with scaffolds distinct from existing inhibitors. Molecules were generated and filtered them based on physicochemical validity, engagement of key residues^22,41^ (Asp12, Asp69), redocking consistency by Glide^42^, and molecular dynamics simulations (Methods, Supplementary Fig. 4). Three synthesizable candidates were advanced to experimental evaluation using a homogeneous time-resolved fluorescence (HTRF) competition assay against a reference ligand known to occupy the same pocket^22^ (Methods). Beyond quantifying inhibitory potency, this assay directly verifies whether binding occurs at the intended site. Two compounds exhibited IC_50_ values of 24 µM and 36 µM (Fig. 4b–c, Supplementary Fig. 5b-c), confirming successful discovery of inhibitors with moderate affinity. Notably, the most potent candidate, KRAS G12D-compound-3, was synthesized as a mixture of four stereoisomers owing to synthetic complexity, which may under-estimate its true potency. Importantly, KRAS G12D-compound-1 and -3 show minimal Tanimoto similarity^43^ to the famous inhibitor MRTX1133 (0.12 and 0.15, respectively) and low similarity to each other (0.25), demonstrating identification of previously unobserved chemical scaffolds targeting this challenging pocket. KRAS G12D–compound-3 constitutes a structurally differentiated chemotype with a conserved yet distinct interaction network relative to MRTX1133. Structural analyses suggest that it preserves several critical polar contacts observed in the MRTX1133 co-crystal structure (Supplementary Fig. 6). The protonated N-methyl group of the tetrahydroisoquinoline (THIQ) core functionally emulates the C4-bicyclic amine of MRTX1133, maintaining the essential salt bridge with Asp12. Similarly, the phenolic hydroxyl group forms a hydrogen bond with Asp69, recapitulating the interaction pattern of the naphthol moiety in MRTX1133. Despite adopting a distinct spatial orientation, the ethynyl substituent, together with the chloro group and the tetrahydrobenzo[b]thiophene fragment, efficiently occupies the adjacent hydrophobic cavity. The thiophene ring further contributes through an aromatic *π*–cation interaction with Arg68, reinforcing binding stabilization. Importantly, our design extends beyond mimicry of the reference scaffold. An amide-linked aniline projects into the solvent-exposed region, establishing a complementary triad of interactions with Glu62, His95, and Asp92. The amide NH hydrogen bonds with the Glu62 carboxylate, the aniline ring engages in face-to-face *π*–*π* stacking with His95, and the meta-amino group forms a previously unexploited hydrogen bond with Asp92, an interaction absent in MRTX1133. Collectively, this expanded interaction network illustrates AnewOmni’s capacity to systematically interrogate and exploit underexplored binding subpockets, thereby capturing additional binding energy and spatial opportunity within the KRAS G12D pocket.

**Figure 4.**
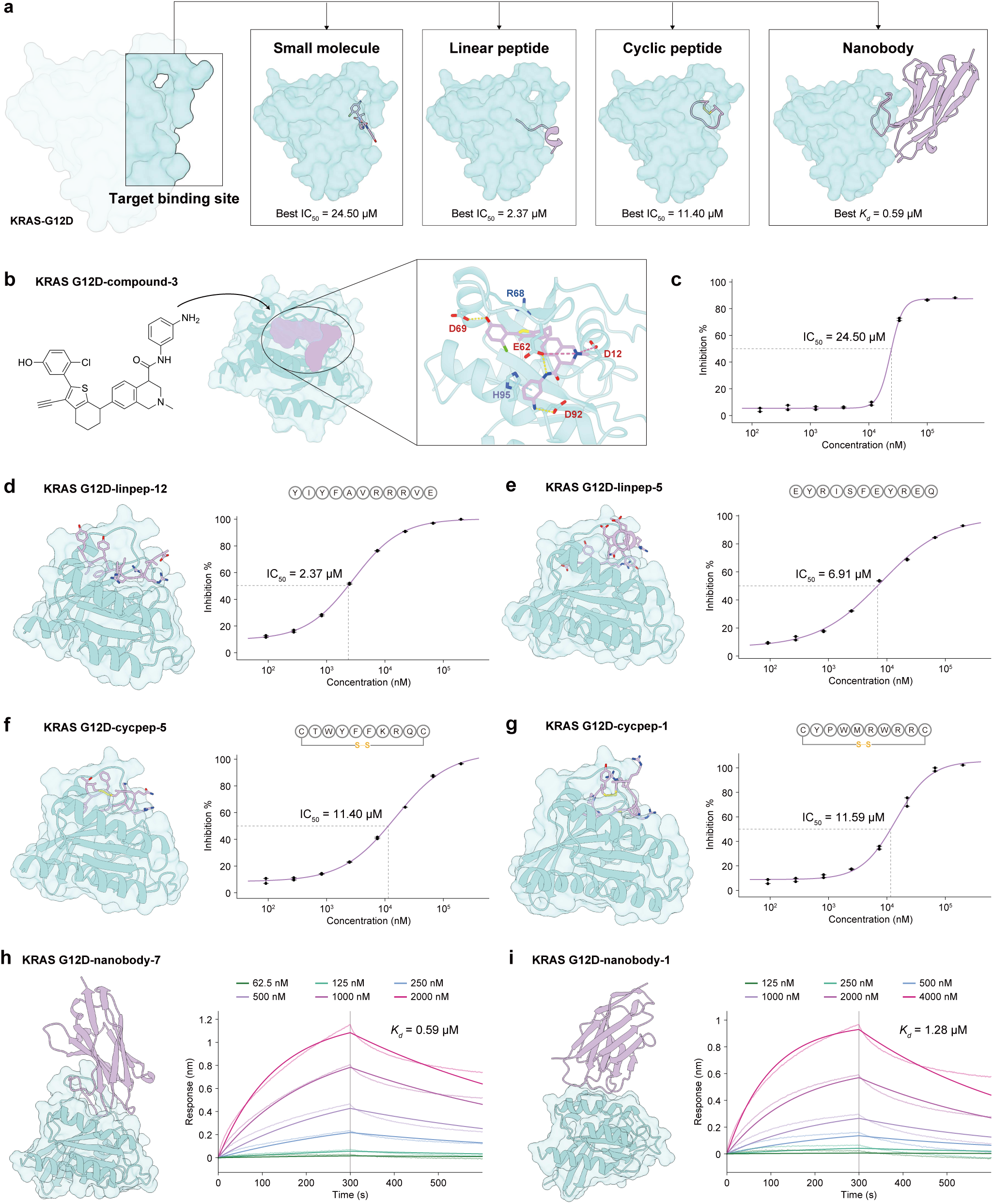
Designed binders of different modalities targeting KRAS G12D. Small molecules, peptides, and nanobodies were designed to bind the same switch II binding site of KRAS G12D. **a**, Representative visualizations of the designed small molecules, peptides, and nanobodies, together with the best half-maximal inhibitory concentration (IC_50_) relative to a reference ligand^22^ for this site or the dissociation constant (K_d_) measured across tested generations. **b**, Chemical structure and three-dimensional binding geometry of KRAS G12D–compound-3 generated by our AnewOmni. **c**, IC_50_ of KRAS G12D–compound-3 measured by the homogeneous time-resolved fluorescence (HTRF) assay. **d-g**, Structures of KRAS G12D-linpep-12, -5 and KRAS G12D-cycpep-5, -1, as well as their IC_50_ to a reference ligand^22^ determined by HTRF. **h-i**, Structure of KRAS G12D-nanobody-7 and -1, as well as their binding affinities to KRAS G12D determined by BLI.

We next applied AnewOmni to the design of linear and cyclic peptides targeting the same Switch II pocket of KRAS G12D. The peptides were first generated and subjected to simple physicochemical filtering (Methods, Supplementary Fig. 7a–b). Valid candidates further underwent molecular dynamics (MD) simulations for 100 ns, and those exhibiting stable trajectories were retained. Binding affinities were subsequently estimated using MM/PBSA for ranking (Methods, Supplementary Fig. 7c). In total, 50 peptides (top 30 linear and top 20 cyclic) were selected for synthesis and experimentally tested by the HTRF assay^22,44^ to determine IC_50_ to the reference compound^22^ at the designed binding site (Methods). Among linear peptides, 7 of 30 produced > 50% inhibition at 100 µM (Supplementary Table 1, Supplementary Fig. 5e,h-k), corresponding to a 23% success rate for IC_50_ < 100 µM, with the most potent reaching IC_50_ of 2.4 and 6.9 µM (Fig. 4d-e). For cyclic peptides, 7 of 20 achieved IC_50_ < 100 µM (Supplementary Table 1, Supplementary Fig. 5f,l-q), likewise yielding a 35% success rate, with the strongest binders reaching IC_50_ of 11 µM and 12 µM (Fig. 4f-g). Notably, as no cyclic peptides existed in the training data, the success of cyclic peptide designs arose purely from the programmability of AnewOmni and the transferable cross-modality design principles it learned. The effectiveness of MD-based filtering further indicates that the generated structures provide reasonable initial binding conformations, as MD stability depends strongly on the starting pose.

Finally, we focused on finding antibody or single-domain antibody (VHH, or nanobody) hits for the same binding site. We built a pipeline involving our AnewOmni and Protenix^45^ to automatically select suitable framework and redesign Complementarity Determining Regions (CDRs) binding to the given epitope on the target. We first curated a set of antibody and nanobody frameworks from SAbDab^46^ in which CDRs contribute the majority of binding contacts, thereby minimizing influence of frameworks on binding (Supplementary Table 2). Starting from these frameworks, we alternated between CDR generation with AnewOmni and complex structure prediction with Protenix. Candidates were ranked mostly based on epitope coverage and ratio of interactions formed by CDR to provide initializations with plausible binding poses for subsequent iterations (Methods). Finally, we observed nanobodies evolved from three frameworks (PDB ID: 8G70, 3JBE, 4W6X) enriched during iterations (Supplementary Fig. 8a), none of which has a target similar to KRAS G12D in their native complexes. For experimental validation, we explored two complementary selection strategies. In the first, we selected designs exhibiting low self-consistent root mean square deviation (scRMSD) between the generated structures and Protenix-predicted complexes, with a threshold of 5.0 Å. It is worth noting that Protenix is biased towards other epitopes instead of the one we desired (Supplementary Fig. 8b), thus few designs could pass the scRMSD filter (Supplementary Fig. 8c), making this problem even tougher. Of four such designs synthesized, three nanobodies bound KRAS G12D with single-digit micromolar affinity as measured by bio-layer interferometry (BLI), achieving a 75% success rate (Supplementary Fig. 9a-d). In the second, more aggressive strategy, we selected designs solely based on high generative likelihood without enforcing structural self-consistency. Among the three candidates tested, one nanobody exhibited a sub-micromolar binding affinity of 587 nM (Fig. 3h, Supplementary Fig. 9e-g), achieving a stronger affinity than the first strategy (Fig. 3i, Supplementary Fig. 9i-j) yet with a lower success rate.

### Orthosteric and Allosteric Inhibition of PCSK9

Cardiovascular disease causes over four million deaths annually in Europe, accounting for nearly half (47%) of all deaths, with elevated low-density lipoprotein (LDL) cholesterol as a major contributing factor^47^. As a key regulator of LDL, proprotein convertase subtilisin-like/kexin type 9 (PCSK9) thus emerges as a highly attractive target for both scientific and pharmaceutical research^48^. Under physiological conditions, LDL is cleared from circulation through binding to the low-density lipoprotein receptor (LDLR), followed by endocytosis. LDLR subsequently dissociates from LDL and is recycled back to the cell surface, whereas LDL is directed toward lysosomal degradation^48^ (Fig. 5a). PCSK9 disrupts this process by binding to LDLR and stabilizing its conformation, thereby blocking recycling and instead routing the PCSK9–LDLR complex to lysosomal degradation (Fig. 5b). As a result, inhibition of the PCSK9–LDLR protein–protein interaction (PPI) represents a compelling strategy to enhance LDLR recycling and reduce plasma LDL cholesterol levels. However, the PPI interface on PCSK9 is large and flat^49^, presenting a substantial challenge for hit identification. To this end, we pursued two complementary paradigms using AnewOmni: 1) Designing orthosteric peptides on the PPI interface (Fig. 5c); 2) Identifying an alternative binding site and designing allosteric small-molecule modulators (Fig. 5c).

**Figure 5.**
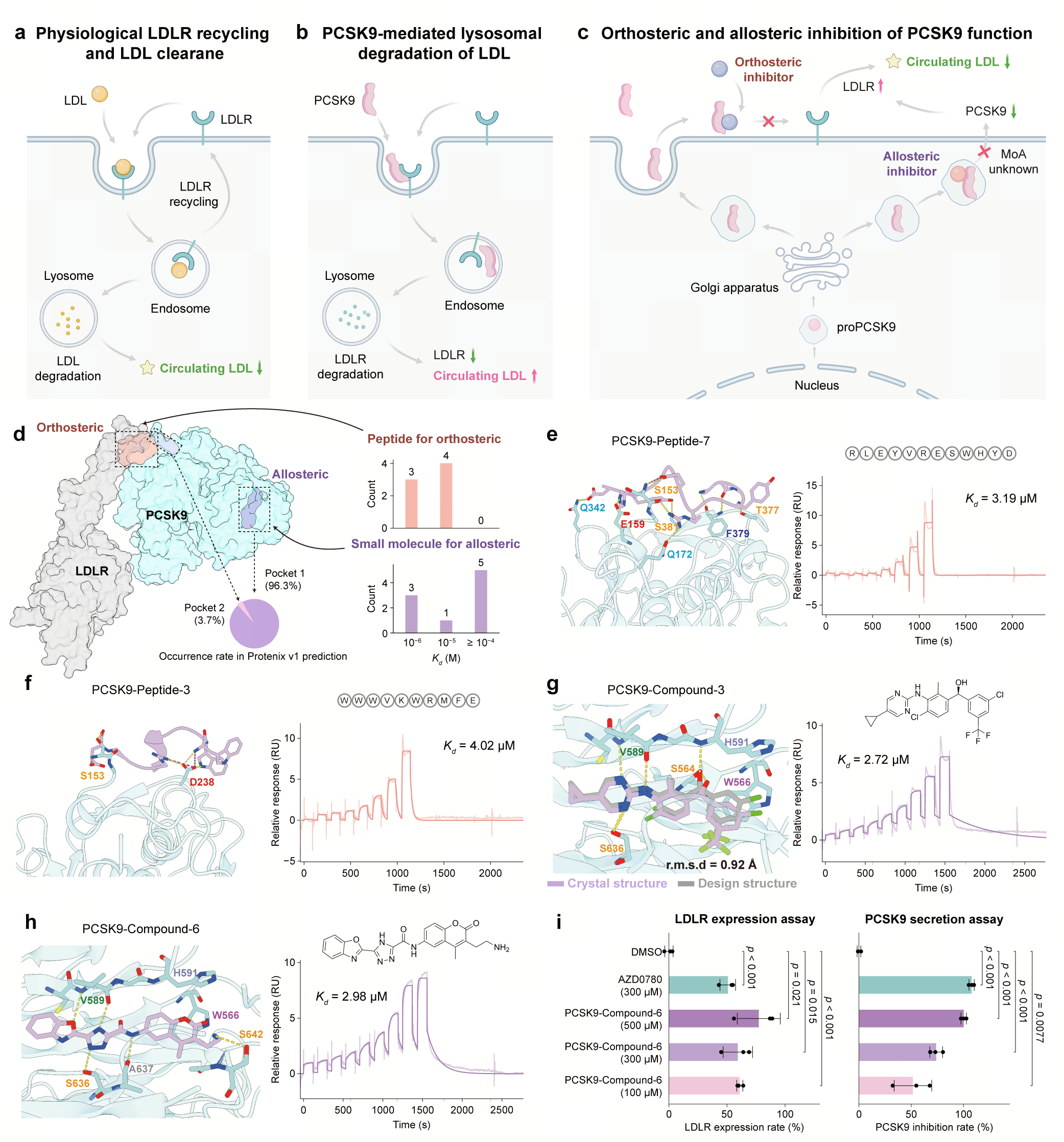
Orthosteric and allosteric inhibitor designs for PCSK9. First, orthosteric peptides were designed against PCSK9. Furthermore, identification of an allosteric binding site enabled the design of small-molecule inhibitors with a potentially novel mechanism of action (MoA). **a**, Schematic representation of physiological LDLR recycling and LDL clearance. LDL binds LDLR at the cell surface, is internalized via endocytosis, and subsequently degraded in lysosomes, while LDLR is recycled back to the cell surface. **b**, Schematic representation of PCSK9-mediated lysosomal degradation of LDLR. PCSK9 binding promotes internalization and degradation of the PCSK9–LDLR complex, reducing LDLR abundance and increasing circulating LDL. **c**, Schematic representation orthosteric and allosteric inhibition of PCSK9 function. Orthosteric inhibitors directly block the PCSK9–LDLR interaction, whereas the identified allosteric inhibitors are likely to hinder PCSK9 secretion, thereby indirectly preventing PCSK9-mediated LDLR degradation. **d**, Left: Orthosteric and allosteric binding sites on PCSK9, with the allosteric site revealed by Protenix prediction of the complex between PCSK9 and a clinical-stage compound lacking structural or mechanistic characterization^50^. Right: Distribution of measured K_d_ levels for designed orthosteric peptides and allosteric small molecules. **e-f**, Designed structures of PCSK9-peptide-7 and PCSK9-peptide-3, as well as their binding affinity to PCSK9 determined by SPR. **g-h**, Designed structures of PCSK9-compound-3 and PCSK9-compound-6, as well as their binding affinity to PCSK9 determined by SPR. Crystal structure is solved for PCSK9-compound-3, and exhibits an RMSD of 0.92 Å to the designed structure. **i**, Cell-based assays showing concentration-dependent upregulation of LDLR and inhibition of PCSK9 secretion, with AZD0780 (from TOPSCIENCE, TSID: T64362, CAS: 2455427-91-3) as the positive control. Both assays were measured with triplicates (N=3), and the error bars represent the standard deviations. Panel **a-c** were created using BioRender (https://biorender.com).

For the orthosteric strategy, we designed linear peptides of 10 to 12 residues targeting the PPI interface on PCSK9. Encouraged by the performance of likelihood-based ranking in nanobody design on the KRAS target mentioned above, we explored whether this lightweight and efficient strategy could generalize to peptide selection. Specifically, we generated 5,000 peptide candidates and selected designs for synthesis and experimental validation based solely on generative likelihood. The mean (µ) and standard deviation (a-) of the likelihood distribution were computed (Supplementary Fig. 10a), and a selection threshold of µ + 3a- was applied to identify candidates with exceptionally high likelihoods. Seven peptides exceeded this threshold and were advanced for experimental testing. Surface plasmon resonance (SPR) measurements revealed that four of the seven synthesized peptides bound PCSK9 with dissociation constants (K_d_) below 10 µM, corresponding to a 57% hit rate (Fig. 5e-f, Supplementary Fig. 10c-g, Methods). The strongest ones achieved K_d_ values of 3.19 µM and 4.02 µM (Fig. 5e-f). These results provide further evidence that unsupervised generative likelihood might serve as an effective ranking criterion, offering a computationally efficient alternative to physics-based scoring while maintaining accuracy and generalizability across targets and molecular modalities.

AZD0780 is a recently reported small-molecule modulator of PCSK9 with favorable clinical profiles, despite not directly inhibiting the PCSK9–LDLR interaction^50^. Its precise binding site and mechanism of action remain unclear^50^. Nevertheless, this observation suggests an alternative opportunity for allosteric modulation of PCSK9 function. Motivated by this possibility, we sought to identify a novel allosteric binding site, design small-molecule binders targeting this site, and explore their functional mechanism in regulating LDLR degradation. We first employed the state-of-the-art cofolding model Protenix^45^ to predict 1,000 complex structures between PCSK9 and AZD0780. Strikingly, 94.5% of the predictions converged on a cryptic binding site located on the C-terminal domain (CTD) of PCSK9 (Fig. 5f). Using this predicted allosteric site as the design target, we applied AnewOmni to generate candidate small molecules and synthesized nine designs passing the filters (Methods, Supplementary Fig. 11) for experimental evaluation (Supplementary Note 7). SPR measurements showed that three of the ten compounds bound PCSK9 with K_d_ below 10 µM, with the most potent candidates (PCSK9-compound-3 and 6) achieving a K_d_ of 2.72 and 2.98 µM (Fig. 5g-h, Supplementary Fig. 10i-o). We next evaluated the functional impact of PCSK9-compound-6 in cellular assays measuring LDLR expression. Notably, treatment with PCSK9-compound-6 at 100 µM resulted in LDLR expression levels comparable to those achieved by AZD0780 at 300 µM (Fig. 5i, Methods). Given that the identified allosteric site is located on CTD of PCSK9, which is related to PCSK9 secretion^51^, we further assessed the effects of these compounds on PCSK9 secretion. Cellular assays revealed that both AZD0780 and PCSK9-compound-6 effectively suppressed PCSK9 secretion (Fig. 5i, Methods), suggesting a potential mechanism in which allosteric binding reduces PCSK9-mediated LDLR degradation by impairing PCSK9 secretion (Fig. 5c). To confirm the binding site and binding mode of PCSK9-compound-3, we solved the crystal structure of the compound in complex with PCSK9 at a resolution of 2.70 Å (Fig. 5g, Extended Data Table 1). Structural alignment on PCSK9 revealed that PCSK9-compound-3 occupies exactly the predicted allosteric pocket, in agreement with the design model. Moreover, the binding pose generated by AnewOmni closely matches the experimentally determined structure, with a root mean square deviation (RMSD) of 0.92 Å. These results demonstrate the high accuracy of the model in designing molecular binders at atomic resolution.

## Discussion

The discovery of artificial modulators of specific biomolecular interactions has long been a cornerstone problem, spanning a variety of molecular modalities such as small organic molecules, proteins, and nucleic acids. Despite shared physical principles of molecular interactions, their design has traditionally been treated as fundamentally distinct problems due to historical divergence of experimental toolkits. In this work, we introduce AnewOmni, a unified generative framework that models diverse molecular modalities at atomic resolution within a shared latent space, enabling programmable and modality-agnostic molecular design, which is essential for the transfer of fundamental recognition principles across modalities, leading to enhanced performance and generalization beyond conventional chemistries.

We demonstrated that joint training on different modalities with our AnewOmni yielded consistent benefits across public benchmarks, in-distribution analysis, and out-of-distribution generalization, as reflected by enhanced *in silico* metrics (Fig. 2a), richer interaction patterns (Fig. 2c,d), and zero-shot generalization to unseen targets, including DNA, RNA, glycans, and phosphorylation sites (Fig. 2e-g), as well as the ability to generate large molecules beyond commonly explored molecular weight ranges (Fig. 2h). The programmable graph prompt further enables fine-grained control over chemistry, topology, and geometry, allowing a single representation system to support a spectrum of design downstream applications such as cyclic peptide design, motif control, non-canonical amino acid incorporation, anchor-conditioned design, covalent interactions, molecular growth, and linker design (Fig. 3). Together, these results suggest that despite differences in chemical composition and structure, diverse molecular modalities obey shared geometric and physical principles of molecular recognition that can be learned and transferred from large-scale atomic-resolution data using AnewOmni.

Experimental validation confirmed the practical effectiveness of our model in generating binders from different modalities targeting the same switch II binding site on KRAS G12D, a well-known and challenging cancer target, requiring only low-throughput experiments. Small-molecule design yielded two hits with IC_50_ of 24 µM and 36 µM from only three synthesized compounds, while peptide design achieved 23% and 35% success rates for linear and cyclic ones, with IC_50_ reaching as low as 2.37 µM. Nanobody design achieved a success rate of 75% under self-consistent RMSD ranking with Protenix, and 33.3% based on unsupervised likelihood, with the best binder exhibiting a K_d_ of 587 nM from seven synthesized candidates. The atomic-resolution design capability of AnewOmni further supports exploration of distinct binding sites for functional modulation, as illustrated by the PCSK9 case. Peptide design targeting the orthosteric site achieved a 57% success rate from seven synthesized candidates, with a best K_d_ of 3.19 µM. Leveraging Protenix, we identified a previously uncharacterized cryptic allosteric site on PCSK9 and successfully designed small-molecule binders targeting this site, achieving a 30% success rate from nine synthesized compounds and a best K_d_ of 2.72 µM. Cellular assays revealed functional modulation of LDLR expression comparable to a drug in clinical trial, and mechanistic analysis suggested that the allosteric small molecule potentially function by interfering with PCSK9 secretion. Finally, a crystal structure of a designed binder confirmed the structural accuracy of AnewOmni, with the designed structure closely matching the experimentally solved conformation at an RMSD of 0.92 Å.

Despite the encouraging success, several limitations of the current pipeline remain. First, post-filtering steps based on Protenix predictions or physics-based molecular simulations are computationally intensive. Although our preliminary results indicate that the unsupervised likelihood produced by the generative model itself may serve as a lightweight proxy to partially bypass these filters, its general applicability requires further validation across a broader range of targets and modalities. Second, the current framework does not yet provide explicit control over binding affinity. While the generative pipeline is effective at producing positive binders, none of the computational metrics have indication of the affinity levels of individual designs. Nevertheless, AnewOmni points toward a general foundational paradigm for molecular modulator discovery. The programmable graph prompt mechanism enables fine-grained control over chemistry, topology, and geometry, fulfilling the complicated customization demands of diverse downstream applications. Cross-modal transfer within a unified framework further suggests the possibility of mimicries between molecular modalities while preserving functions to acquire modality-specific properties, such as converting antibodies into peptides to enhance cell permeability while preserving interaction patterns. More broadly, it envisions future molecular design systems that move beyond modality-specific pipelines toward more general molecular reasoning engines, with the potential to extend molecular design into regimes where data and human intuition remain limited.

## Methods

### Dataset

To train AnewOmni as a unified generative foundation model across molecular modalities, we curated a comprehensive multi-modal dataset spanning diverse distributions, including small molecules, peptides, and antibodies. For small molecules, we assembled a high-quality structural corpus by integrating BioLip2^52^, PDBBind v2020^53^, and CrossDocked2020^54^. Following established practice^54^, the test split of CrossDocked2020 was held out exclusively for *in silico* evaluation. All small molecule data were subjected to a rigorous filtering pipeline to remove overlaps with test sets and exclude nonspecific ligands, such as solvents and inorganic ions (Supplementary Table 3). To further enhance the ability of the model to learn diverse fine-grained atomic interactions and three-dimensional binding poses, we augmented the training data with SIU^55^, a large-scale synthetic dataset comprising over 5.3 million protein–ligand complexes with associated bioactivity information. For peptides, we used PepBench and ProtFrag^9,56^ as training data, which provide a diverse collection of protein–peptide complexes enriched with synthetic samples derived from intra-domain protein contacts. The LNR benchmark^56^ was reserved exclusively for evaluation following previous convention^9,56^. For antibodies, training data were obtained from the Structural Antibody Database (SAbDab)^46^, with the RAbD dataset^57^ held out for testing. To prevent data leakage across modalities and benchmarks, a sequence identity cutoff of 40% was enforced between training and test sets, following previous literature^58,59^. Detailed dataset statistics and preprocessing procedures for all modalities are provided in Supplementary Note 1.2.2.

### Model Architecture

Our AnewOmni consists of four main components: atom-to-block decomposition, an all-atom variational autoencoder (VAE), a latent diffusion model, and an exact likelihood estimation module (Supplementary Notes 1).

In the atom-to-block decomposition stage, chemically coherent building blocks are defined to balance atomistic resolution and structured chemical priors. All standard biomolecular units are preserved explicitly, including canonical amino acids and nucleobases. Other chemical entities are decomposed into frequent molecular fragments mined from ChEMBL^24^, while rare substructures with low frequencies are further decomposed into individual atoms to ensure complete coverage of arbitrary chemistries (Supplementary Notes 1.2.1).

The all-atom VAE establishes a reversible mapping between atom-level molecular structures and compact block-level latent representations. Each building block is encoded into a latent embedding ***h*** ∈ ℝ^d^ together with a latent coordinate *x^⤑^* ∈ ℝ^3^, forming a continuous latent point cloud with compressed chemical priors and context-aware atomistic geometries (Supplementary Notes 1.3.1). Generation is then performed by a latent diffusion model operating in this smooth, compressed latent space, conditioned on the target binding site (Supplementary Notes 1.3.2).

To enable explicit and flexible control over the generative process, we introduce a programmable graph prompt mechanism that imposes constraints on chemical composition, topology, and three-dimensional geometry. These prompts are integrated directly into the denoising process of the latent diffusion model stepwise, which serve as node-level or edge-level controls on the latent points during generation. For training, the conditions are randomly sampled from ground-truth structures to promote robustness on arbitrary combination of controls (Supplementary Notes 1.3.3). During inference, arbitrary user-specified constraints can be enforced via classifier-free guidance^60^.

Both the VAE and the latent diffusion module are parameterized using an equivariant transformer that incorporates geometric symmetries while remaining compatible with the conventional transformer architecture to support large-scale training (Supplementary Notes 1.3.4). Finally, we implement exact likelihood estimation along the generative trajectory (Supplementary Notes 1.8.1), which serves as an unsupervised, modality-agnostic self-ranking metric. Although we also explored supervised confidence heads for pairwise distance error prediction (Supplementary Notes 1.8.2), we observed that such confidence estimates tend to overfit to seen molecular distributions and generalize poorly to unseen targets (Fig. 2f, Supplementary Fig. 3b). We therefore adopt unsupervised likelihood as the primary criterion for candidate ranking throughout this work.

### Training Regimen

Our AnewOmni was trained on 48 NVIDIA Tesla A800 GPUs using the AdamW optimizer with an initial learning rate of 1 × 10^-4^. The all-atom autoencoder, the latent diffusion model, and the supervised confidence model were trained for 500, 500, and 100 epochs, respectively, where SIU was subsampled to 5% per epoch. Learning rates were reduced by a factor of 0.8 when the validation loss plateaued for five consecutive evaluations. For the autoencoder and the latent diffusion model, validation was performed every 10 epochs, whereas the supervised confidence module was validated at every epoch. Gradient norms were clipped to 10.0 to stabilize training. Batch sizes were dynamically adjusted to accommodate the quadratic complexity of attention while minimizing padding, with implementation details provided in Supplementary Note 1.5.1 and Supplementary Table 1.5.1. A complete list of hyperparameters is reported in Supplementary Table 5.

### Pipeline for Small Molecule Design

Because the chemical synthesis of small molecules is costly, we designed a post-filtering pipeline to prioritize generated candidates with favorable chemical and physical properties for synthesis. The pipeline processes the raw generated candidates through the following sequential stages:

1. PAINS filter: We first apply the Pan-assay interference compounds^61^ (PAINS) rule as an "alert structure exclusion" filter, to remove molecules containing substructures known to be reactive or prone to assay interference.
2. 3D structure validity check: We assess the 3D pose generated by AnewOmni by PoseBusters^62^ toolkit, evaluating both intramolecular and intermolecular validity. Intramolecular checks verify geometric plausibility, including bond angles, bond lengths, planarity, and internal energy, as well as detecting internal clashes. While intramolecular checks identify poses exhibiting severe steric clashes against the protein pocket.
3. Key Interaction Constraints: A set of key interactions is pre-defined based on the target mechanism. We leverage ProLif^28^ package to detect interaction between pockets and small molecules, and strictly select molecules that satisfy all the interactions.
4. Physics-based Refinement and Stability Assessment: Selected candidates undergo pose refinements using GLIDE^42^. We assume that the generated poses are near-native starting points, requiring only local optimization to filter-out high-energy artifacts. Thus, "refine only" mod was selected, and we calculate the RMSD between the original pose and refined pose to drop candidates with high RMSD.
5. In the final stage, we use Molecular Dynamics (MD) simulations to evaluate the stability and dynamic behavior within the binding pocket, root mean square deviation (RMSD) and key interaction lifetime are used as key metrics.

Detailed implementations for MD simulations and the key interaction definitions are provided in Supplementary Note 4.2 and 5, respectively.

### Pipeline for Peptide Design

For generated peptides, we applied a set of basic physicochemical filters to remove unfavorable candidates. Specifically, the following criteria were used:

1. Geometry: Simple geometric checks were performed on atomic structures, and peptides containing covalent bonds longer than 2.0 Å were discarded.
2. Clash: Peptides with clash rates exceeding 10% were removed. A clashed atom on peptides was defined as having a Van der Waals radius overlap of ≥ 0.4 Å with a neighboring atom on the target^63^.
3. Cysteine exclusion: Cysteine residues were omitted to simplify synthesis and experimental validation, as disulfide formation or isolated cysteines would require additional synthetic steps.
4. Hydrophobicity: The GRAVY score^64^ was computed, and peptides with scores above zero were discarded due to excessive hydrophobicity that could hinder synthesis.
5. L-amino acids: Because the model can generate D-amino acids through all-atom reconstruction, only peptides composed entirely of L-amino acids were retained to facilitate synthesizability.

For disulfide cyclic peptides, cysteine residues are required, thus we skipped the normal cystenine exclusion filter. Instead, we applied specialized filters to exclude sequences containing cysteines beyond the intended disulfide pair or exhibiting distorted disulfide geometry. Acceptable disulfide bonds were constrained to lengths of 1.5–2.5 Å. The C_{3_ atom was required to remain bonded to the sulfur atom with a distance below 2.0 Å, and C_{3_ atoms from paired cysteines were required to remain non-clashing, with separations greater than 3.0 Å.

For the KRAS G12D target, peptide generation continued until 400 cyclic peptides and 600 linear peptides passed all filters. These candidates were subsequently evaluated using molecular dynamics simulations (Supplementary Note 4.1), retaining structures with trajectory RMSD below 3.5 Å, and were further ranked by binding energies computed using MM/PBSA^65^. For the PCSK9 target, generation proceeded until 5,000 peptides passed the filters, after which candidates were ranked by model likelihood to select those with high confidence beyond 3-sigma range.

### Pipeline for Antibody Design

For antibody design, we aim to design target-binding antibodies by designing CDRs on the frameworks in a given library to disentangle the binding characteristics, which are dominated by CDRs, and the developability, which are dominated by frameworks. While in practical applications, the framework library are typically provided by the users according to developability, in this paper we curated a library of 48 antibodies and single-domain antibodies (nanobodies) from SAbDab^46^, filtered to retain those in which more than 80% of binding contacts are contributed by CDRs (Supplementary Table 2). Contact ratios were computed based on heavy-atom distances below 5.0 Å. Such frameworks are well suited for epitope-directed CDR redesign because the specificity is dominated by CDRs instead of frameworks. While AnewOmni focuses on generative modeling of the local interfaces, we design an iterative pipeline to leverage state-of-the-art cofolding models ^45^ for global structure validation. The pipeline involves iterations of three stages: (1) Sequence-structure codesign of CDRs with AnewOmni targeting the given epitope; (2) Structure prediction of the complex of the target protein and the designed antibody; (3) Metric calculation and candidate ranking.

At the beginning of each iteration, we collect ten parent candidates: the top eight ranked by the composite score defined below, together with two candidates sampled randomly to encourage exploration. For each parent candidate, we leverage its docked complex for AnewOmni to generate 100 new CDRs in a sequence-structure codesign manner, which are then ranked together by likelihood (Supplementary Note 1.8.1) to get the top five generations, yielding a total of 50 candidates for structure prediction in the next stage. For structure prediction, multiple sequence alignments (MSA) and templates are provided for the target protein, while the generated antibody structures are supplied as templates for the antibody. Next, the following metrics are calculated to derive a composite score:

1. Binding site overlap ratio between the generated complex and the desired binding site (weight 10.0)
2. Ratio of contacts mediated by CDRs (weight 5.0)
3. ipTM between target and antibody (weight 0.3)
4. pLDDT of the antibody (weight 0.2)
5. normalized ipAE between the target and the antibody (weight 0.1)
6. average likelihood of subsequent CDR generations derived from this candidate (weight 0.2)
7. normalized self-consistent root mean square error (scRMSD) between the generated structure and the predicted structure (weight 0.2)

Detailed implementations for each metric are provided in Supplementary Notes 6.1. We assign the highest weight to binding-site overlap to prioritize epitope-specific generation. The fraction of CDR-mediated contacts is assigned the second-highest weight to discourage false positives arising from framework-dominated interactions, as these frameworks should be irrelevant to the target protein. Weights of other metrics sum up to 1.0 to adjust the ranking order when binding site overlap and contacts ratio of CDRs are at similar levels. The likelihood term is evaluated based on subsequent generations, reflecting whether a candidate provides a favorable docking orientation in the learned distribution of AnewOmni. After 60 iterations, corresponding to approximately 3,000 evaluated candidates, we first apply plausibility filters requiring a binding-site overlap ratio greater than 0.4 and a CDR contact fraction greater than 0.5. Final candidates are then selected using two strategies. The first is a conservative strategy that retains only candidates with scRMSD below 5.0 Å, leveraging Protenix as an orthogonal validation (Supplementary Notes 6.2). The second strategy is more risky, where we first select the one exhibiting the highest average generative likelihood, then further redesign the CDRs based on this candidate to get candidates with higher likelihood for experimental validation.

### Synthesis of the Designed Small Molecules

All the small molecules targeting KRAS G12D or PCSK9 were synthesized by WuXi AppTec Company, with detailed synthetic routes described in Supplementary Note 7. All synthesized molecules are visualized in Supplementary Fig. 4a and 11f for KRAS G12D and PCSK9, respectively. The IUPAC names and SMILES are provided in Supplementary Table 6.

### Peptide synthesis

Peptides were synthesized using standard Fmoc-based solid-phase peptide synthesis (SPPS). Synthesis was performed on the GenScript PepHTS high-throughput peptide synthesizer using 2-chlorotrityl chloride resin as the solid support. Protected amino acids were coupled sequentially from the C-terminus to the N-terminus. HBTU and DIEA were used as coupling reagents, DMF served as the solvent, and 20% piperidine in DMF was used for Fmoc deprotection.

Following chain assembly, the resin-bound peptide was cleaved using a trifluoroacetic acid (TFA)–containing cleavage cocktail. The resulting crude peptide was precipitated, collected, and purified by preparative reverse-phase high-performance liquid chromatography (RP-HPLC). The purity of the final peptide was assessed by analytical RP-HPLC, and the molecular weight was confirmed by electrospray ionization mass spectrometry (ESI-MS). All peptide products showed > 95% purity, and the observed molecular masses were consistent with theoretical values. All synthesized and tested peptides are summarized in Supplementary Table 7 and 8.

### Nanobody synthesis and purification

All nanobodies used in this study were manufactured by GenScript. DNA sequences encoding the target nanobodies were codon-optimized, synthesized, and subcloned into a mammalian expression vector. Transfection-grade plasmid DNA was prepared using a Maxi-prep kit. For expression, Chinese Hamster Ovary (CHO) cells were maintained in suspension culture at 37 °C with 5% CO_2_ on an orbital shaker. One day prior to transfection, cells were seeded to achieve the optimal density. On the day of transfection, the plasmid DNA and transfection reagent were mixed at an optimized ratio and added to the culture. A feed supplement was added 24 hours post-transfection to support cell growth and protein production. The sequences of synthesized nanobodies are included in Supplementary Table 9.

The cell culture supernatant was clarified by centrifugation and subsequent filtration to remove cell debris. The clarified supernatant was loaded onto an affinity purification column at a controlled flow rate. Following washing steps to remove non-specific impurities, the target protein was eluted and buffer-exchanged into the final formulation buffer. The physicochemical properties of the purified nanobodies were characterized by SDS-PAGE and size-exclusion chromatography (SEC-HPLC) to assess molecular weight and monomeric purity. Protein concentrations were determined by measuring the UV absorbance at 280 nm.

### HTRF binding assay for KRAS G12D

All test compounds were prepared as a 3 -fold, 10-point dilution series in Echo qualified LDV plates (Labcyte, Cat#LP-0200) and transferred at 100 nL of a 20 mM stock per reaction, resulting in a final concentration of 100 µM in the assay. The reaction buffer contained 50 mM HEPES (pH 7.5), 5 mM MgCl_2_, and 1 mM DTT. Recombinant human KRAS Protein, Human, Recombinant His & Avi-tag Biotinylated was purchased from SinoBiological, Cat#12259-H56E-B, corresponding to amino acids 1-169, expressed in *E. coli* with a C-terminal Avi biotinylated tag (MW=22 kDa), Lyophilized from sterile 50 mM Tris (pH8.0). 5 µL of the prepared 5nM KRAS G12D solution were dispensed into the assay wells. Subsequently, 5 µL of a 100nM Tracer (Compound 45^22^) solution was added into the wells. To initiate the reaction, 10 µL of a 0.125 nM Terbium-Streptavidin (Tb-SA; Cisbio, 610SATLF) solution was dispensed into the assay plates. The mixture was incubated at 25 °C for 1 hour prior to HTRF signal detection. The HTRF signal was measured using a PHERAstar reader in time-resolved fluorescence mode (dual emission: EX337nm, EM665nm/EM620nm) according to manufacturer protocols. The HTRF ratio is given by:

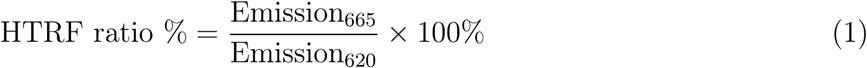

Data were fitted to the Hill equation (with **a** fixed Hill coefficient of 1) using nonlinear regression in Xlfit software (IDBS) to determine IC50 values :

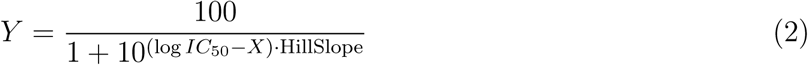

The analysis was performed using the "log(inhibitor) vs. normalized response - Variable slope" model, X represents the log of the inhibitor concentration, Y represents the normalized response (ranging from 100 to 0%), and the Hill slope was initialized at -1. All experiments were conducted in duplicate using biologically independent samples.

### Bio-layer interferometry (BLI) assay for KRAS G12D

Bio-layer interferometry (BLI) assays were performed using an Octet R8 system (Sartorius) at 30°C. Biotinylated KRAS-G12D protein (obtained from WuXi AppTec) was immobilized on Super Streptavidin (SSA) biosensors (Sartorius). The sensors were pre-hydrated in the assay buffer for at least 10 min prior to use. For immobilization, the protein was diluted to 50 µg/mL and loaded on the sensors for 600 s.

For nanobody binding kinetics, experiments were performed in PBST (without DMSO). Nanobodies were analyzed as a 2-fold dilution series. The assay cycle comprised a baseline step (60 s), followed by an association phase (300 s) and a dissociation phase (300 s).

Data were processed using the Octet Analysis Studio 13.0 software. Sensorgrams were corrected by double referencing, subtracting signals from a reference sensor and a blank buffer injection. The corrected binding data were globally fitted to a 1:1 Langmuir binding model to determine the equilibrium dissociation constant *K_d_*, the association rate *k_on_*, and the dissociation rate *k_off_*.

### Surface plasmon resonance (SPR) assay for KRAS G12D

SPR measurements were performed using a Biacore 8K system (Cytiva) in PBS-P buffer(pH 7.4, 0.05% (v/v) Surfactant P20, GE Healthcare). The KRAS G12D proteins (SinoBiological, Cat#12259-H56E-B) were immobilized on a CM5 chip (GE Healthcare) through amide coupling in 10 mM NaOAc (pH 4.5) for 420 s at a flow rate of 10 µl/min. Designed binders were injected as analytes at a single 150 nM concentration during binder pre-screening or in serial dilutions to assess binding kinetics. Injections were performed at a flow rate of 30 µl/min with an association phase of 120 s and a dissociation phase of 600 s. The chip surface was regenerated after each injection using 3 mM NaOH for 30 s at a flow rate of 30 µl/min. Binding curves were fitted to a 1:1 Langmuir binding model using the Biacore 8K analysis software.

### Surface plasmon resonance (SPR) assay for PCSK9

Surface Plasmon Resonance (SPR) experiments were performed on a Biacore 8K (Cytiva) at 25 °C. Biotinylated PCSK9 (AcroBio, PC9-H5223) was captured on a Series S Sensor Chip SA (Streptavidin-coated, Cytiva). Flow cell 1 served as the reference surface (blank immobilization), while flow cell 2 was immobilized with the target protein. The immobilization levels achieved for the active channels ranged from approximately 4570 to 4715 response units (RU). The running buffer consisted of 10 mM HEPES (pH 7.4), 150 mM NaCl, 0.2% P20, and 5% DMSO.

Analytes were prepared in the running buffer to ensure a constant DMSO concentration. To account for bulk refractive index variations, solvent correction was performed using a series of DMSO standards. Binding kinetics were determined via single-cycle kinetics (SCK). Analytes were injected as a 2-fold dilution series (nine concentrations) sequentially over the reference and active flow cells. Injections were performed at a flow rate of 30 µL/min with a contact time of 120 s per concentration, followed by a final dissociation phase of 1200 s.

Data analysis was performed using the Biacore 8K Evaluation Software. Sensorgrams were double-referenced by subtracting the responses from the reference flow cell and the blank buffer injections. After solvent correction, the kinetic profiles were fitted to a 1:1 binding model to determine the equilibrium dissociation constant *K_d_*, the association rate *k_on_*, and the dissociation rate *k_o_*_ff_.

### LDLR ELISA assay

HepG2 cells (ATCC, HB-8065) were maintained in MEM (Gibco) supplemented with 10% fetal bovine serum (FBS) and 100 U/mL penicillin-streptomycin at 37 °C in a humidified atmosphere containing 5% CO_2_. Cells were seeded at 100 µL/well of cell suspension containing 25000 cells into a 96-well culture plate, with MEM Assay medium containing 10% lipid-depleted FBS (v/v, Lonsera, U412-001) and 100 unit/mL penicillin-streptomycin for overnight culture at 37 °C under 5% CO_2_. The following day, the culture medium was discarded, replaced with 100 µL fresh assay medium containing serial diluted compounds. For the LDLR assay, 100 µL fresh assay medium containing 2 µg/ml recombinant human protein PCSK9 (Sino, 29698-H08H) was added and returned to incubation at 37 °C under 5% CO_2_ for 48 h. Following incubation, the medium was removed, and cells were lysed in 100 µL of RIPA lysis buffer containing a 1.5 × protease and phosphatase inhibitor cocktail. The plates were incubated for 60 min at room temperature with shaking at 300 rpm. The cell plates were centrifuged to transfer supernatant into a fresh plate with 4-fold dilution with Calibrator Diluent buffer. LDLR levels were quantified using a human LDLR ELISA kit (R&D Systems, DLDLR0) according to the manufacturer’s instructions. Relative LDLR expression was calculated by normalizing test samples to the vehicle control (VC):

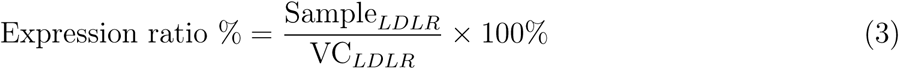

### PCSK9 secretion AlphaLISA assay

HepG2 cells (ATCC, HB-8065) were maintained in MEM (Gibco) supplemented with 10% FBS and 100 U/mL penicillin-streptomycin at 37 °C in a humidified atmosphere containing 5% CO_2_. Cells were seeded into 96-well plates at a density of 2.5 × 10^4^ cells/well in 100 µL of assay medium (MEM supplemented with 10% lipid-depleted FBS (Lonsera, U412-001) and 100 U/mL penicillinstreptomycin) and incubated overnight. The following day, the culture medium was replaced with 200 µL of assay medium containing serially diluted compounds, and plates were incubated for 48 h. Following incubation, plates were centrifuged at 1200 rpm for 5 min. Supernatants (100 µL/well) were collected and transferred to a fresh plate. Samples (4.5 µL) were transferred to a 384-well plate and analyzed using the PCSK9 AlphaLISA kit (Revvity, AL270F) according to the manufacturer’s instructions.

### Protein crystallization and structure determination

The PCSK9–compound 3 complex (1:3 molar ratio) was crystallized at 10.3 mg/mL using sitting-drop vapor diffusion at 20 °C. The reservoir solution consisted of 0.1 M HEPES (pH 7.5) and 20% (w/v) polyethylene glycol (PEG) 10,000. Crystals were soaked with 2 mM Compound 3 for 18 h at 20 °C, cryoprotected in reservoir solution supplemented with 25% glycerol, and flash-cooled in liquid nitrogen. Diffraction data were collected at beamline BL10U2 of the Shanghai Synchrotron Radiation Facility (SSRF) at 100 K. Data were processed using XDS and Aimless from the CCP4 suite. Phases were determined by molecular replacement using Phaser. Model building and refinement were performed using Coot and Refmac5. The quality of the final model was validated using MolProbity.

## Data Availability

All datasets used for model training are publicly available. BioLiP2 can be accessed at https://aideepmed.com/BioLiP/download.html, which provides official scripts for downloading the structural database. CrossDocked2020 with splits established by literature^54^ is available via Google Drive (https://drive.google.com/drive/folders/1CzwxmTpjbrt83z_wBzcQncq84OVDPurM). PDBbind v2020 is accessible from its official website (https://www.pdbbind-plus.org.cn/). SIU is hosted on the Hugging Face dataset platform (https://huggingface.co/datasets/bgao95/SIU). PepBench and ProtFrag are deposited on Zenodo (https://zenodo.org/records/13373108). SAbDab can be downloaded from its official website (https://opig.stats.ox.ac.uk/webapps/sabdab-sabpred/sabdab/). All additional structures analyzed in this study are identified by PDB accession codes and are available from the Protein Data Bank (https://www.wwpdb.org/). Atomic coordinates and structure factors of the X-ray structure for PCSK9 in complex with PCSK9-compound-3 reported here have been deposited in the PDB and is currently under review. The accession code will be released upon publication.

## Code Availability

The codes of AnewOmni are publicly available at github.com/bytedance/AnewOmni. We have also provided a platform for ease of use at https://anewbt-mind.healthybaike.com.

## Author contributions

X.K., Ziting Zhang (Zt.Z.), J.M., K.L., and Y.L. conceived and designed the generative model with contributions from J.C., G.L., R.J., Z.Y. and W.H. for theoretical derivations and engineering. X.K., J.C., and Zt.Z. collected data and established developing benchmarks, with helps from T.Z., Q.W, and Zishen Zhang (Zs.Z.). X.K. and J.C. trained and ablated the model. J.C., X.K., Zt.Z. and L.W. conducted the computational experiments and analyzed the results. G.L. and J.M. organized the synthesis of designed small molecules, peptides, and nanobodies. G.L., Q.Z., Y.S., X.K., W.D., and Xiaolun Wang (Xl.W.) designed and conducted the wet-lab experiments on KRAS G12D and PCSK9, with helps from J.Z. on developing assays. J.C., G.L., M.L., Z.G., W.L. and J.Z. helped construct physical-based models for filtering designs. G.L., Xl.W. and Q.H. conducted the crystal structure determination. X.K., J.C., Zt.Z., G.L., L.W., Zs.Z., W.T, R.J., Z.Y., L.Z., W.L. drafted the manuscript, with insights from Xiaowo Wang (Xw.W.) and Y.S. during discussion. All authors read and contributed to the manuscript. The study was supervised by Y.L., K.L. and J.M.

