## supplementary information for "Programming Biomolecular Interactions with All-Atom Generative Model"

[1]Department of Computer Science and Technology, Tsinghua University, Beijing, China
[2]Institute for AI Industry Research (AIR), Tsinghua University, Beijing, China
[3]ByteDance AI Drug Discovery, Shanghai, China
[4]Anew Therapeutics, North Tower, Singapore
[5]MOE Key Laboratory of Bioinformatics and Bioinformatics Division of BNRIST, Department of Automation, Tsinghua University, Beijing, China
[6]Department of Pharmaceutical and Artificial-Intelligence Sciences, Shanghai, China
[7]Institute of Artificial Intelligence for Medicine, Shanghai Jiao Tong University School of Medicine, Shanghai, China
[8]Department of Chemistry, Tsinghua University, Beijing, China
[9]Laboratory of Protein Design and Immunoengineering, École Polytechnique Fédérale de Lausanne and Swiss Institute of Bioinformatics, Lausanne, Switzerland
[10]Aithyra Research Institute for Biomedical Artificial Intelligence, Austrian Academy of Sciences, Vienna, Austria
[11]Gaoling School of Artificial Intelligence, Renmin University of China, Beijing, China
[12]Beijing Key Laboratory of Research on Large Models and Intelligent Governance, Beijing, China
[13]Department of Electronic Engineering, Tsinghua University, Beijing, China
[+]these authors contributed equally to this work

#### List of Algorithms

1 Subgraph-Level Decomposition . . . 2
2 Architecture of the Geometric Equivariant Transformer . . . 7
3 Architecture of the Graph-based Adapter. . . . 8
4 Architecture of the Adapted Geometric Equivariant Transformer . . . 9
5 Training Algorithm of the Iterative Full-Atom Autoencoder . . . 12
6 Training Algorithm of Conditional Latent Diffusion Model . . . 13
7 Sampling Algorithm of the Conditional Latent Diffusion Model . . . 13
8 Sampling Algorithm of the Iterative Full-Atom Autoencoder . . . 14
9 Normalized Log-Likelihood Computation . . . 17
10 Training Algorithm of Confidence Model . . . 18
11 Inference Algorithm of Confidence Model . . . 19

### 1 Supplementary Methods

#### 1.1 Notations

In the follow-up sections, we describe the algorithms of each module with the subsequent notations. We denote the training dataset as $\mathcal{S}$, and $\mathbb{I}$ stands for the node index list. Each molecule in data-space is represented as an all-atom geometric graph $\mathcal{G} = (\mathcal{V}, \mathcal{E})$. Specifically for a molecule with $N$ atoms and $M$ building blocks, the node set is defined as $\mathcal{V} = (\boldsymbol{A}, \vec{\boldsymbol{X}}) = (\bigcup_{i=1}^{M} \boldsymbol{A}_i, \bigcup_{i=1}^{M} \vec{\boldsymbol{X}}_i)$, where $\boldsymbol{A} \in \mathbb{Z}^N$ and $\vec{\boldsymbol{X}} \in \mathbb{R}^{N\times 3}$ denote the element types and coordinates of $N$ atoms, respectively, while $\boldsymbol{A}_i = \{a_{ij}\}_{j=1}^{n_i} \in \mathbb{Z}^{n_i}$ and $\vec{\boldsymbol{X}}_i = \{\vec{\boldsymbol{x}}_{ij}\}_{j=1}^{n_i} \in \mathbb{R}^{n_i\times 3}$ denote the corresponding features of block $i$ with $n_i$ atoms. The edge set is defined as $\mathcal{E} = \{\boldsymbol{B}_i\}_{i=1}^{M} \cup \{\boldsymbol{B}_{ij} \mid i \neq j\}_{i,j=1}^{M}$, where $\boldsymbol{B}_i \in \mathbb{Z}^{n_i\times n_i}$ and $\boldsymbol{B}_{ij} \in \mathbb{Z}^{n_i\times n_j}$ denote intra-block and inter-block chemical bonds, respectively. Note that for simplicity, we use matrix notation here, but the atoms within each block are unordered. We denote the corresponding 2D topological graph as $\mathcal{G}_{2D} = (\boldsymbol{A}, \mathcal{E}_{2D})$, where $\mathcal{E}_{2D} \subseteq \mathcal{E}$ represents the chemical bond connections.

We denote $\mathbb{V} = \mathbb{V}_{AA} \cup \mathbb{V}_{NB} \cup \mathbb{V}_{PS}$ as the building block vocabulary consisting of amino acids $\mathbb{V}_{AA}$, nucleotide bases $\mathbb{V}_{NB}$, and principle subgraphs $\mathbb{V}_{PS}$, and $s_i \in \mathbb{V}$ denote the building block type of block $i$. We denote the counter $\mathcal{C}$ as the frequencies for learned principal subgraphs. We further denote $\mathcal{G}_z = (\mathcal{V}_z, \mathcal{E}_z)$ as the latent graph of blocks encoded by an all-atom encoder, and hereinafter simplify the latent node feature as $\mathcal{V}_z = \mathcal{Z}$. Specifically, $\mathcal{Z} = \{(\boldsymbol{z}_i, \vec{\boldsymbol{z}}_i)\}$ contains the latent states $\boldsymbol{z}_i \in \mathbb{R}^d$ ($d = 8$ in this paper) and coordinates $\vec{\boldsymbol{z}}_i \in \mathbb{R}^3$ of block $i$ in $\mathcal{G}_z$, and $\mathcal{E}_z \in \mathbb{Z}^{M\times M}$ denote the edge feature in the latent space.

We split $\mathcal{G}_x$, $\mathcal{G}_y$ to represent the geometric graphs corresponding to the data-space of the binder and the context environment (e.g. the binding sites, the framework regions of antibodies), while $\mathcal{Z}_x, \mathcal{Z}_y$ denote the latent node feature of the binder and the environment. We denote $\boldsymbol{H}, \vec{\boldsymbol{V}}$ as the matrix of invariant and equivariant features inside the backbone models. If required, the layer index is specified as a superscript wrapped by a bracket (e.g. $\boldsymbol{H}^{(l)}$). $\vec{\boldsymbol{u}}_i^t = (\boldsymbol{z}_i^t, \vec{\boldsymbol{z}}_i^t)$ is the time-dependent latent representation of block $i$, timestep $t$ during the latent diffusion process. To incorporate conditions into the model, we denote the condition graph as $\mathcal{G}_c = (\mathcal{V}_c, \mathcal{E}_c)$ with $\mathcal{V}_c = (\boldsymbol{H}_c, \vec{\boldsymbol{X}}_c)$. Here $\boldsymbol{H}_c = \boldsymbol{H}_r \cup \boldsymbol{H}_d$ represents the invariant features of real nodes $\boldsymbol{H}_r$ and dummy nodes $\boldsymbol{H}_d$, and $\vec{\boldsymbol{X}}_c = \vec{\boldsymbol{X}}_r \cup \vec{\boldsymbol{X}}_d$ denotes the real coordinates $\vec{\boldsymbol{X}}_r$ and dummy coordinates $\vec{\boldsymbol{X}}_d$. We denote $\boldsymbol{M}_{2D}, \boldsymbol{M}_{3D} \in \{0, 1\}^M$ as the controlling mask vectors on 2D and 3D information based on each specific task.

For model architectures, $\mathcal{E}_\phi$, $\mathcal{D}_\xi$, $\boldsymbol{\epsilon}_\theta$, $\mathcal{A}_\eta$, and $\mathcal{C}_\psi$ denote the learnable all-atom encoder, decoder, the unconditional part of the latent diffusion model, the conditional adapter, and the confidence model, parameterized by $\phi$, $\xi$, $\theta$, $\eta$, and $\psi$ respectively. We use $\varphi$ (with proper subscripts) to denote Multi-Layer Perceptrons (MLPs), RBF, Softmax, LN, and Lookup denote radial basis functions, softmax activation, layer normalization, and lookup table retrieval, respectively. The loss terms CE, MSE, and $D_{\mathrm{KL}}$ denote Cross Entropy, Mean Squared Error, and Kullback–Leibler divergence. Finally, $\|\cdot\|_2$ denotes the $\ell_2$-norm of a vector, $|\cdot|$ denotes the cardinality of a set, $\mathrm{sum}(\cdot)$ denotes the summation of all elements in a vector, $\mathbf{U}(a, b)$ denotes the uniform distribution on $[a, b]$, and $\mathcal{N}(\mu, \sigma^2)$ denotes the Gaussian distribution with mean $\mu$ and variance $\sigma^2$. $\odot$ stands for the Hadamard product, and $\|$ stands for concatenation.

#### 1.2 Data Processing

##### 1.2.1 Frequent Molecular Fragment Mining

**Algorithm 1** Subgraph-Level Decomposition

```
def SubgraphDecomposition(𝒢_2D, 𝕍_PS, 𝒞):
    𝒢′ ← 𝒢_2D
    while True do
        freq ← −1; ℱ ← None
        for ⟨ℱ_i, ℱ_j, ℰ_ij⟩ in 𝒢′ do
#           Get neighboring fragments in the same ring with fragments i and j
            𝔽_i, 𝔽_j ← RingPrior(ℱ_i), RingPrior(ℱ_j)
            if (ℱ_j ∉ 𝔽_i and 𝔽_i ≠ ∅) xor (ℱ_i ∉ 𝔽_j and 𝔽_j ≠ ∅) then
#               If one of the fragment has a neighbor in a same ring while the other does not, discard
                this pair
                continue
            end if
#           Merge neighboring fragments into a new fragment
            ℱ′ ← Merge(⟨ℱ_i, ℱ_j, ℰ_ij⟩)
#           Convert a graph to SMILES representation
            s ← GraphToSMILES(ℱ′)
            if s ∈ 𝕍_PS and 𝒞[s] > freq  then
                freq ← 𝒞[s]
                ℱ ← ℱ′
            end if
        end for
        if freq = −1 then
#           No further merging needed
            break
        else
#           Update the graph representation
            𝒢′ ← MergeSubGraph(𝒢′, ℱ)
        end if
    end while
    return 𝒢′
```

Algorithm 1 outlines the block-level molecular decomposition procedure, adapted from the principal subgraph algorithm[1]. The method takes as input a 2D atom-level molecular graph $\mathcal{G}_{2D}$, a pre-defined set of principal subgraphs $\mathbb{V}_{PS}$, and their corresponding occurrence counts $\mathcal{C}$ obtained from the prior extraction step. The core process involves iteratively detecting and combining adjacent principal subgraph pairs whose union exhibit the highest frequency within the vocabulary. This iterative merging halts when no further adjacent subgraph pairs present in the vocabulary can be found. The `MergeSubGraph` function operates on the input graph $\mathcal{G}_{2D}$ and the selected highest-frequency fragment $\mathcal{F}$. Whenever $\mathcal{F}$ is identified within $\mathcal{G}_{2D}$, the function consolidates the adjoining nodes (*e.g.*, $i$ and $j$) that constitute $\mathcal{F}$ into a single new node, which now represents the combined fragment $\mathcal{F} = \mathcal{G}_i \cup \mathcal{G}_j \cup \mathcal{E}_{ij}$. Furthermore, we have adapted the original algorithm to prioritize the merging of adjacent fragments that belong to the same ring structure. This adjustment is implemented via the `RingPrior` function, which locates neighboring fragments that share a ring with the input fragment in the molecule. For any pair of neighboring fragments $i$ and $j$ under consideration for merging, if one fragment has a neighbor in the same ring while the other does not, the pair is discarded to prioritize merging fragments within the same ring. In the current study, we employ a vocabulary comprising 300 principal subgraphs. These were extracted from the ChEMBL database[2] using kekulized molecular representations.

###### *1.2.2 Datasets*

We compiled a comprehensive multi-modal dataset for training AnewOmni, covering different modalities including small molecules, peptides, and antibodies.

For biocomplexes concerning small molecules, we first integrated data from three primary sources: BioLip2 (up to 2024-06-30)[3], PDBbind v2020[4], and CrossDocked2020[5]. We implemented a rigorous data processing pipeline to ensure high data quality as follows:

1. Remove all entries overlapping with the test set and duplicate items (identified by PDB code) across the three source datasets.

2. Filter out and drop invalid ligands based on a customized exclusion list derived from the Chemical Component Dictionary (CCD), which removes common solvents, ions, and nonspecific ligands that would bias model during the generation phase (e.g., ATP and its analogs, glycan, Long-chain aliphatic compounds, Supplementary Table 3)

3. The majority ligand distributions within each dataset were analyzed, and manual inspection was performed to identify and exclude non-drug-like ligands.

Following this filtration pipeline, the final training corpus consisted of 14,200 entries from PDBBind, 18,012 entries from BioLip2, and 85,938 complexes from CrossDocked2020. To further enhance the model's foundation capability of accurate 3D pose generation and diverse atomic interaction understanding between ligand and protein, we additionally integrated SIU[6], a large-scale dataset comprising (5,408,415) synthetic protein-small molecule complex. During training, 5% of the entire dataset was sampled as a part of training data per epoch.

For peptides, we utilized the PepBench and ProtFrag datasets[7,8], comprising 4,157 protein-peptide complexes and 70,498 synthetic samples for training, and 114 complexes for validation. Testing was conducted on the LNR dataset which consists of 93 protein-peptide complexes with peptide lengths ranging from 4 to 25 residues[7]. A sequence identify cutoff of 40% was enforced between the training, the validation, and the test set to prevent data leakage.

For antibodies, in line with previous literature[9,10], we utilized SAbDab entries deposited prior to 2024-09-24 for training and validation[11], while 60 antigen-antibody complexes from RAbD[12] were reserved for testing. To prevent data leakage, the training data was filtered to ensure a sequence identity below 40% against the test set, resulting in final datasets of 9,473 entries for training and 400 entries for validation.

#### 1.3 Model Architecture

###### *1.3.1 All-atom Variational Auto-Encoder*

We introduce an iterative full-atom variational autoencoder (VAE) composed of an encoder $\mathcal{E}_\phi$ and a decoder $\mathcal{D}_\xi$[13], which is designed to learn the correspondence between detailed all-atom geometries and a low-dimensional latent representation.

**Encoder** The encoder $\mathcal{E}_\phi$ maps the binder graph $\mathcal{G}_x$ and the binding-site graph $\mathcal{G}_y$ into their respective latent point clouds $\mathcal{Z}_x$ and $\mathcal{Z}_y$:

$$\mathcal{Z}_x = \mathcal{E}_\phi(\mathcal{G}_x), \qquad \mathcal{Z}_y = \mathcal{E}_\phi(\mathcal{G}_y), \tag{4}$$

where $\mathcal{Z}_x = (\boldsymbol{z}_i, \vec{\boldsymbol{z}}_i)$ denotes the collection of latent features associated with block $i$ in $\mathcal{G}_x$, consisting of an invariant latent vector $\boldsymbol{z}_i \in \mathbb{R}^d$ (with $d = 8$ in this work) and a geometric latent coordinate $\vec{\boldsymbol{z}}_i \in \mathbb{R}^3$. The variables $\boldsymbol{z}_i$ and $\vec{\boldsymbol{z}}_i$ are drawn from the encoder-predicted Gaussian distributions $\mathcal{N}(\boldsymbol{z}_i; \boldsymbol{\mu}_i, \boldsymbol{\sigma}_i)$ and $\mathcal{N}(\vec{\boldsymbol{z}}_i; \vec{\boldsymbol{\mu}}_i, \vec{\boldsymbol{\sigma}}_i)$, respectively, via the reparameterization trick[14]. An analogous construction is applied to obtain the latent representation $\mathcal{Z}_y$ for the binding site. Importantly, the binder and binding site are processed by the encoder in a decoupled manner, with no mutual information exchange. This design is consistent with inference-time usage, where generation is conditioned solely on the binding site $\mathcal{G}_y$ in the absence of an observed binder $\mathcal{G}_x$. During training, we additionally perturb $\vec{\boldsymbol{z}}_i$ with random noise sampled from $\mathcal{N}(\mathbf{0}, \mathbf{I})$ before feeding it into the decoder. This enforced perturbation is essential because the diffusion module operates in latent space and may not generate coordinates perfectly matched to the distribution defined by the autoencoder, leading to deviations during generation. Introducing noise in advance during training enables the decoder to accommodate such imperfections and improves its robustness.

**Decoder** The decoder $\mathcal{D}_\xi$ recovers the full-atom structure of the binder through a two-step procedure consisting of block-type decoding followed by atomic coordinate reconstruction. In the first step, the decoder infers the block identities $s_i$ for the latent binder points by jointly conditioning on $\mathcal{Z}_x$ and $\mathcal{Z}_y$. The resulting block types are then mapped to atom types and intra-block chemical bonds via a vocabulary lookup:

$$s_i = \mathcal{D}_{\xi_1}(\mathcal{Z}_x, \mathcal{Z}_y), \qquad \boldsymbol{A}_i, \boldsymbol{B}_i = \mathrm{Lookup}(s_i, \mathbb{V}), \tag{5}$$

where $s_i$ specifies the decoded block type, while $\boldsymbol{A}_i$ and $\boldsymbol{B}_i$ correspond to the atom types and chemical bond patterns associated with block $i$, respectively. Subsequently, a structure module reconstructs atomic coordinates through an iterative update scheme, analogous to a lightweight flow-matching formulation[15]:

$$\{(\boldsymbol{H}_i^t, \vec{\boldsymbol{V}}_i^t)\} = \mathcal{D}_{\xi_2}(\{(\boldsymbol{A}_i, \boldsymbol{B}_i, \vec{\boldsymbol{X}}_i^t)\}, \mathcal{Z}_x, \mathcal{Z}_y, \mathcal{G}_y, t), \tag{6}$$

$$\vec{\boldsymbol{X}}_i^{t-\Delta t} = \vec{\boldsymbol{X}}_i^t + \Delta t \vec{\boldsymbol{V}}_i^t, \tag{7}$$

where $\boldsymbol{H}_i^t$ and $\vec{\boldsymbol{V}}_i^t$ denote the per-atom hidden representations and the corresponding vector fields for block $i$ at time $t$. During training, $t$ is sampled from $\mathbf{U}(0, 1)$, which enables arbitrary settings for time schedule during sampling. For sampling, we found a good balance between efficiency and performance by decreasing the time variable $t$ from 1.0 to 0.0 using a fixed step size $\Delta t = 0.1$, resulting in a total of 10 iterative updates. At the initial step $t = 1.0$, the atomic coordinates $\vec{\boldsymbol{X}}_i^1$ for each block $i$ are initialized by sampling from a Gaussian distribution $\mathcal{N}(\vec{\boldsymbol{z}}_i, \boldsymbol{I})$. In addition, inter-block chemical bonds are predicted based on local spatial proximity, considering only atom pairs whose distances fall below 3.5Å:

$$b_{ij,pq}^t = \mathrm{Softmax}(\mathrm{MLP}(\boldsymbol{h}_{i,p}^t + \boldsymbol{h}_{j,q}^t)), \quad i \neq j, \tag{8}$$

where $b_{ij,pq}^t$ indicates the predicted bond type (e.g., single, double, triple, or none) between the $p$-th atom of block $i$ and the $q$-th atom of block $j$. Here, $\boldsymbol{h}_{i,p}^t = \boldsymbol{H}_i^t[p]$ and $\vec{\boldsymbol{x}}_{i,p}^t = \vec{\boldsymbol{X}}_i^t[p]$ represent the hidden feature and spatial coordinate of the $p$-th atom in block $i$, respectively. The function MLP refers to a three-layer multilayer perceptron with SiLU activation[16], whose input is defined as the sum of $\boldsymbol{h}_{i,p}^t$ and $\boldsymbol{h}_{j,q}^t$ to enforce commutativity. Bond inference is further constrained to atom pairs within the 3.5Å neighborhood, as determined by the Euclidean distance $d_{ij,pq}^t = \|\vec{\boldsymbol{x}}_{i,p}^t - \vec{\boldsymbol{x}}_{j,q}^t\|_2$. Although discrepancies between predicted bond types and geometric distances may arise, such cases can be readily identified and filtered out (Supplementary Note 1.7).

Both the encoder and decoder are implemented using an equivariant transformer architecture[17], with the full procedure summarized in Algorithm 2. The optimization objective for training the VAE is provided in Supplementary Notes 1.4.1.

###### 1.3.2 Latent Diffusion Model

The autoencoder abstracts the high-dimensional molecular structures into compact latent variables, which facilitates learning the conditional distribution $p(\mathcal{Z}_x \mid \mathcal{Z}_y)$ using a diffusion-based framework. Before initiating the diffusion procedure, the latent coordinates are standardized by removing the center of mass of the binding-site latent point cloud $\mathcal{Z}_y$, followed by rescaling with an empirical factor of 10.0[18]. The forward diffusion process progressively injects Gaussian noise into the latent variables from $t = 0$ to $t = T$, ultimately mapping the data distribution to an isotropic Gaussian prior $\mathcal{N}(\mathbf{0}, \boldsymbol{I})$. Conversely, the reverse process incrementally eliminates the noise, recovering the original latent representations from $t = T$ back to $t = 0$. Let $\vec{\boldsymbol{u}}_i^t = [\boldsymbol{z}_i^t, \vec{\boldsymbol{z}}_i^t]$ denote the latent state of block $i$ at diffusion step $t$; the forward transitions are defined as:

$$q(\vec{\boldsymbol{u}}_i^t \mid \vec{\boldsymbol{u}}_i^{t-1}) = \mathcal{N}(\vec{\boldsymbol{u}}_i^t; \sqrt{1-\beta^t} \cdot \vec{\boldsymbol{u}}_i^{t-1}, \beta^t \boldsymbol{I}), \tag{9}$$

$$q(\vec{\boldsymbol{u}}_i^t \mid \vec{\boldsymbol{u}}_i^0) = \mathcal{N}(\vec{\boldsymbol{u}}_i^t; \sqrt{\bar{\alpha}^t} \cdot \vec{\boldsymbol{u}}_i^0, (1-\bar{\alpha}^t)\boldsymbol{I}), \tag{10}$$

where $\bar{\alpha}^t = \prod_{s=1}^{s=t}(1-\beta^s)$, and $\beta^t$ follows the cosine noise schedule[19]. Conditioned on the initial latent $\vec{\boldsymbol{u}}_i^0$, the latent state at an arbitrary timestep $t$ admits a closed-form sampling expression:

$$\vec{\boldsymbol{u}}_i^t = \sqrt{\bar{\alpha}^t}\vec{\boldsymbol{u}}_i^0 + \sqrt{1-\bar{\alpha}^t}\boldsymbol{\epsilon}_i, \tag{11}$$

with $\boldsymbol{\epsilon}_i \sim \mathcal{N}(\mathbf{0}, \boldsymbol{I})$. The reverse diffusion dynamics reconstruct the data distribution by successively removing noise[20], and are modeled as:

$$p_\theta(\vec{\boldsymbol{u}}_i^{t-1} \mid \mathcal{Z}_x^t, \mathcal{Z}_y) = \mathcal{N}(\vec{\boldsymbol{u}}_i^{t-1}; \vec{\boldsymbol{\mu}}_\theta(\mathcal{Z}_x^t, \mathcal{Z}_y), \beta^t \boldsymbol{I}), \tag{12}$$

$$\vec{\boldsymbol{\mu}}_\theta(\mathcal{Z}_x^t, \mathcal{Z}_y) = \frac{(\vec{\boldsymbol{u}}_i^t - \frac{\beta^t}{\sqrt{1-\bar{\alpha}^t}}\boldsymbol{\epsilon}_\theta(\mathcal{Z}_x^t, \mathcal{Z}_y, \mathcal{A}_\eta(\mathcal{G}_c), t)[i])}{\sqrt{\alpha^t}}, \tag{13}$$

where $\alpha^t = 1 - \beta^t$, and $\boldsymbol{\epsilon}_\theta$ denotes the denoising network, which is instantiated as an equivariant transformer[17] (Algorithm 2) augmented with a graph-based adapter for condition integration (Algorithm 3). The corresponding training objective is detailed in Supplementary Notes 1.4.2.

Notably, performing diffusion directly in the latent space avoids repeated operations on full-atom geometries, thereby substantially lowering the computational burden during both training and inference.

###### 1.3.3 Programmable Generation via Classifier-Free Guidance

To achieve flexible and programmable generation that enables arbitrary control on chemical composition, topology, and 3D geometry within a single framework, we formulate conditional generation using a disentangled graph-based conditioning mechanism paired with Classifier-Free Guidance (CFG)[21].

**Unified Condition Representation.** We represent user-specified constraints as a heterogeneous condition graph $\mathcal{G}_c = (\mathcal{V}_c, \mathcal{E}_c)$ that interacts with the evolving latent states. The conditioning information is explicitly decoupled into topological constraints (e.g., connectivity) and geometrical constraints (e.g., 3D coordinates), controlled by binary masks $\boldsymbol{M}_{2D}, \boldsymbol{M}_{3D} \in \{0,1\}^M$.

As defined in Algorithm 3, the condition node set $\mathcal{V}_c$ comprises invariant scalar features $\boldsymbol{H}_c$ and coordinate features $\vec{\boldsymbol{X}}_c$. These features integrate information from real nodes $\boldsymbol{H}_r, \vec{\boldsymbol{X}}_r$ and auxiliary dummy nodes $\boldsymbol{H}_d, \vec{\boldsymbol{X}}_d$, where the latter serve as global geometric priors or anchors. This formulation enables a unified interface for various scenarios such as docking conformation generation ($\boldsymbol{M}_{2D} = \mathbf{1}, \boldsymbol{M}_{3D} = \mathbf{0}$), inverse folding ($\boldsymbol{M}_{2D} = \mathbf{0}, \boldsymbol{M}_{3D} = \mathbf{1}$), and motif scaffolding via partial masking.

**Conditional Graph Adapter.** To incorporate heterogeneous conditions into the diffusion process, we introduce a learnable structural adapter module interleaved with the backbone layers. At layer $l$ of the denoising network $\epsilon_\theta$, denoted as $\epsilon_\theta^{(l)}$, we dynamically construct a condition graph that connects the evolving latent states $(\boldsymbol{H}^{(l)}, \vec{\boldsymbol{V}}^{(l)}, \vec{\boldsymbol{X}}^{(l)})$ with the condition nodes $\mathcal{V}_c$.

As detailed in Algorithm 3, the condition graph is built by concatenating real latent nodes with auxiliary dummy nodes representing conditional information, and by introducing edges between dummy nodes and real nodes to encode conditional constraints. The adapter $\mathcal{A}_\eta(\cdot)$ then performs equivariant message passing over this condition graph to aggregate conditional signals and compute guidance updates for the latent representation. These updates are injected back into the main denoising stream through a residual connection, ensuring that each layer of the diffusion model is softly steered by the conditions while preserving the original backbone dynamics. Through this mechanism, the generative trajectory is gradually biased toward regions of high probability under the conditional distribution, without disrupting the pretrained unconditional prior.

**Training with Condition Sampler.** To equip the model with programmable capabilities of handling hybrid control, we adopt a heterogeneous training strategy by stochastically sampling mask configurations $(\boldsymbol{M}_{2D}, \boldsymbol{M}_{3D})$ from a predefined task distribution. Specifically, we set the probability of unconditional generation to 50%, critical for enabling effective Classifier-Free Guidance[21] (CFG). For the other 50%, which are conditional generations, we assign 15% probability to giving full 2D chemistries ($\boldsymbol{M}_{2D} = \mathbf{1}, \boldsymbol{M}_{3D} = \mathbf{0}$), 10% to giving full 3D geometries by providing the center of mass for each block ($\boldsymbol{M}_{2D} = \mathbf{0}, \boldsymbol{M}_{3D} = \mathbf{1}$), and 10% to inpainting, where a subset of blocks provides both 2D and 3D context. Additionally, 15% of samples are trained with partial 2D chemical constraints, while the remaining budget is allocated to random mixed conditions comprising arbitrary combinations of 2D and 3D masks. This diverse sampling schedule ensures the model learns robust conditional distributions $p_t(\vec{\boldsymbol{u}}^t | \mathcal{G}_c)$ across the entire spectrum of constraints.

**Denoising with Guidance.** During training and inference, we control the trade-off between sample diversity and fidelity to the conditions via a guidance scale parameter $w$. The modified noise prediction $\tilde{\boldsymbol{\epsilon}}_\theta$ is computed as a linear combination of the conditional and unconditional outputs of the network $\boldsymbol{\epsilon}_\theta$:

$$\tilde{\boldsymbol{\epsilon}}_\theta(\mathcal{Z}_x^t, \mathcal{Z}_y, \mathcal{A}_\eta(\mathcal{G}_c), t) = (1+w)\,\boldsymbol{\epsilon}_\theta(\mathcal{Z}_x^t, \mathcal{Z}_y, \mathcal{A}_\eta(\mathcal{G}_c), t) - w\,\boldsymbol{\epsilon}_\theta(\mathcal{Z}_x^t, \mathcal{Z}_y, \mathcal{A}_\eta(\emptyset), t). \quad (14)$$

where $\tilde{\boldsymbol{\epsilon}}_\theta$ encapsulates both the invariant noise prediction $\tilde{\boldsymbol{\epsilon}}$ and equivariant noise prediction $\tilde{\vec{\boldsymbol{\epsilon}}}$. By adjusting $w$, we can programmatically strengthen the guidance, ensuring the generated molecules strictly adhere to the provided 2D or 3D partial structures while exploring the chemical space for the unconstrained regions.

##### *1.3.4 Algorithm for Geometric Equivariant Transformer*

**Algorithm 2** Architecture of the Geometric Equivariant Transformer

**def** GraphInitialization($\boldsymbol{H}, \vec{\boldsymbol{X}}$):

### *Single-layer message passing for feature initialization*

$\boldsymbol{h}_i, \boldsymbol{h}_j \leftarrow \boldsymbol{H}[i], \boldsymbol{H}[j]$

$\vec{\boldsymbol{x}}_i, \vec{\boldsymbol{x}}_j \leftarrow \vec{\boldsymbol{X}}[i], \vec{\boldsymbol{X}}[j]$

$\boldsymbol{m}_{ij} \leftarrow [\boldsymbol{h}_i, \boldsymbol{h}_j, \mathrm{RBF}(\|\vec{\boldsymbol{x}}_i - \vec{\boldsymbol{x}}_j\|_2)]$

$\boldsymbol{h}_i \leftarrow \varphi_h\left(\boldsymbol{h}_i, \sum_{j\in\mathcal{N}(i)} \varphi_s(\boldsymbol{m}_{ij}) \cdot \boldsymbol{h}_j\right)$

$\vec{\boldsymbol{v}}_i \leftarrow \sum_{j\in\mathcal{N}(i)} \varphi_v(\boldsymbol{m}_{ij}) \cdot (\vec{\boldsymbol{x}}_i - \vec{\boldsymbol{x}}_j)$

$\boldsymbol{H}, \vec{\boldsymbol{V}} \leftarrow \{\boldsymbol{h}_i\}_{i=1}^{N}, \{\vec{\boldsymbol{v}}_i\}_{i=1}^{N}$

**return** $\boldsymbol{H}, \vec{\boldsymbol{V}}$

**def** EquivariantSelfAttention($\boldsymbol{H}, \vec{\boldsymbol{V}}, \vec{\boldsymbol{X}}$):

### *Compute queries, keys, and values for head s*

$\boldsymbol{Q}_s, \boldsymbol{K}_s \leftarrow \boldsymbol{H}\boldsymbol{W}_s^{Q}, \boldsymbol{H}\boldsymbol{W}_s^{K}$

$\boldsymbol{V}_s \leftarrow [\boldsymbol{H}\boldsymbol{W}_s^{Vh}, \vec{\boldsymbol{V}}\boldsymbol{W}_s^{Vv}]$

### *Distance-aware attention bias*

$\boldsymbol{D} \leftarrow \{\vec{\boldsymbol{X}}[i] - \vec{\boldsymbol{X}}[j]\}_{i,j=1}^{N}$

$\boldsymbol{R} \leftarrow \{\varphi_r(\mathrm{RBF}(\boldsymbol{D}_{ij}))\}_{j\in\mathcal{N}(i)}$

$\boldsymbol{H}_s, \vec{\boldsymbol{V}}_s \leftarrow \mathrm{Softmax}\left(\frac{\boldsymbol{Q}_s^{\top}\boldsymbol{K}_s}{2\sqrt{h_s}} - \boldsymbol{D} + \boldsymbol{R}\right)\boldsymbol{V}_s$

### *Head aggregation and projection*

$\boldsymbol{H}, \vec{\boldsymbol{V}} \leftarrow \sum_s \boldsymbol{H}_s\boldsymbol{W}_s^{Oh}, \sum_s \vec{\boldsymbol{V}}_s\boldsymbol{W}_s^{Ov}$

**return** $\boldsymbol{H}, \vec{\boldsymbol{V}}$

**def** EquivariantFFN($\boldsymbol{H}, \vec{\boldsymbol{V}}$):

### *GVP-style channel mixing*

$\vec{\boldsymbol{V}}_1, \vec{\boldsymbol{V}}_2 \leftarrow \vec{\boldsymbol{V}}\boldsymbol{W}_1, \vec{\boldsymbol{V}}\boldsymbol{W}_2$

$\boldsymbol{H}, \boldsymbol{U} \leftarrow \varphi_{\mathrm{FFN}}(\boldsymbol{H}, \|\vec{\boldsymbol{V}}_1\|_2)$

$\vec{\boldsymbol{V}} \leftarrow \mathrm{LN}(\boldsymbol{U}) \odot \vec{\boldsymbol{V}}_2$

**return** $\boldsymbol{H}, \vec{\boldsymbol{V}}$

**def** GeometricEquivariantTransformer($\boldsymbol{H}, \vec{\boldsymbol{X}}$):

### *Initial feature construction*

$\boldsymbol{H}^{(0)}, \vec{\boldsymbol{V}}^{(0)} \leftarrow \mathrm{GraphInitialization}(\boldsymbol{H}, \vec{\boldsymbol{X}})$

### *Stack of L equivariant transformer layers*

**for** $l = 0, 1, \ldots, L-1$ **do**

$[\boldsymbol{H}^{(l)}, \vec{\boldsymbol{V}}^{(l)}] \leftarrow [\boldsymbol{H}^{(l)}, \vec{\boldsymbol{V}}^{(l)}] + \mathrm{EquivariantSelfAttention}(\mathrm{LN}(\boldsymbol{H}^{(l)}), \vec{\boldsymbol{V}}^{(l)}, \vec{\boldsymbol{X}})$

$[\boldsymbol{H}^{(l+1)}, \vec{\boldsymbol{V}}^{(l+1)}] \leftarrow [\boldsymbol{H}^{(l)}, \vec{\boldsymbol{V}}^{(l)}] + \mathrm{EquivariantFFN}(\mathrm{LN}(\boldsymbol{H}^{(l)}), \vec{\boldsymbol{V}}^{(l)})$

**end for**

**return** $LN(\boldsymbol{H}^{(L)}), \vec{\boldsymbol{V}}^{(L)}$

##### 1.3.5 Algorithm for Graph-Based Adapter

**Algorithm 3** Architecture of the Graph-based Adapter.

**def** AdapterGNNLayer($\boldsymbol{H}_c, \vec{\boldsymbol{V}}_c, \vec{\boldsymbol{X}}_c, \mathcal{E}_c, \boldsymbol{e}_{ij}$):
 # *Retrieve edge indices*
 $i, j \leftarrow Indices(\mathcal{E}_c)$
 $\boldsymbol{h}_i, \boldsymbol{h}_j, \vec{\boldsymbol{v}}_i, \vec{\boldsymbol{v}}_j, \vec{\boldsymbol{x}}_i, \vec{\boldsymbol{x}}_j \leftarrow \boldsymbol{H}_c[i], \boldsymbol{H}_c[j], \vec{\boldsymbol{V}}_c[i], \vec{\boldsymbol{V}}_c[j], \vec{\boldsymbol{X}}_c[i], \vec{\boldsymbol{X}}_c[j]$
 $\boldsymbol{d}_{ij} \leftarrow \mathrm{RBF}(\|\vec{\boldsymbol{x}}_i - \vec{\boldsymbol{x}}_j\|_2)$
 # *Vector Projection*
 $\vec{\boldsymbol{v}}_k \leftarrow \varphi_p(\vec{\boldsymbol{V}}_c), \forall k \in \{1, 2, 3\}$
 $p_i \leftarrow \sum(\vec{\boldsymbol{v}}_1 \odot \vec{\boldsymbol{v}}_2)$
 # *Message Construction and Aggregation*
 $\boldsymbol{m}_h, \boldsymbol{m}_v, \boldsymbol{m}_x \leftarrow \varphi_{msg}([\boldsymbol{h}_i, \boldsymbol{h}_j, \boldsymbol{d}_{ij}, \sum(\vec{\boldsymbol{v}}_{1,i} \odot \vec{\boldsymbol{v}}_{2,j}), \boldsymbol{e}_{ij}])$
 $\boldsymbol{h}_{\mathrm{aggr}} \leftarrow \sum_{j \in \mathcal{N}(i)} \boldsymbol{m}_h$
 $\vec{\boldsymbol{v}}_{\mathrm{aggr}} \leftarrow \sum_{j \in \mathcal{N}(i)} (\vec{\boldsymbol{v}}_{3,j} \cdot \boldsymbol{m}_v + (\vec{\boldsymbol{x}}_i - \vec{\boldsymbol{x}}_j) \cdot \boldsymbol{m}_x)$
 # *Output Projection*
 $\boldsymbol{o}_1, \boldsymbol{o}_2, \boldsymbol{o}_3 \leftarrow \varphi_{out}(\boldsymbol{h}_{\mathrm{aggr}})$
 $\Delta \boldsymbol{H} \leftarrow p_i \odot \boldsymbol{o}_2 + \boldsymbol{o}_3$
 $\Delta \vec{\boldsymbol{V}} \leftarrow \vec{\boldsymbol{v}}_{3,i} \cdot \boldsymbol{o}_1 + \vec{\boldsymbol{v}}_{\mathrm{aggr}}$
 **return** $\Delta \boldsymbol{H}, \Delta \vec{\boldsymbol{V}}$

**def** ConstructGraphEdges($\boldsymbol{M}_{2D}, \boldsymbol{M}_{3D}, M$):
 # *Topological edges from 2D molecule structure*
 $\mathcal{E}_{topo} \leftarrow \{(i, j) \mid \boldsymbol{M}_{2D}[i] = 1,\ \boldsymbol{M}_{2D}[j] = 1,\ i \neq j,\ \|\boldsymbol{B}_{ij}\|_0 > 0\}$
 # *Construct virtual edges linking real nodes and dummy nodes*
 $\mathcal{E}_{anchor} = \bigcup_{i:\boldsymbol{M}_{3D}[i]=1} \left\{(i, i'), (i', i) \mid i' = M + \sigma(i), \sigma(i) = \left(\sum_{j=1}^{i} \boldsymbol{M}_{3D}[j]\right) - 1\right\}$
 $\boldsymbol{e}_{ij} \leftarrow [\varphi_{edge}(\mathcal{E}_{topo}), \varphi_{type}(\mathcal{E}_{anchor})]$
 **return** $\mathcal{E}_{topo} \cup \mathcal{E}_{anchor}, \boldsymbol{e}_{ij}$

**def** GraphAdapter($\boldsymbol{H}_r, \vec{\boldsymbol{V}}_r, \vec{\boldsymbol{X}}_r, \boldsymbol{H}_d, \vec{\boldsymbol{X}}_d, \boldsymbol{M}_{2D}, \boldsymbol{M}_{3D}, l$):
 **if** $\sum \boldsymbol{M}_{2D} = 0$ **and** $\sum \boldsymbol{M}_{3D} = 0$ **then**
  # *Unconditional case*
  **return** $\boldsymbol{0}, \vec{\boldsymbol{0}}$
 **end if**
 # *Construct Condition Graph*
 $\boldsymbol{H}_c \leftarrow \varphi_{in}([\boldsymbol{H}_r, \boldsymbol{H}_d])$
 $\vec{\boldsymbol{V}}_c \leftarrow [\vec{\boldsymbol{V}}_r, \vec{\boldsymbol{0}}]$
 $\vec{\boldsymbol{X}}_c \leftarrow [\vec{\boldsymbol{X}}_r, \vec{\boldsymbol{X}}_d]$
 # *Number of real nodes*
 $M \leftarrow |\boldsymbol{H}_r|$
 $\mathcal{E}_c, \boldsymbol{e}_{ij} \leftarrow \mathrm{ConstructGraphEdges}(\boldsymbol{M}_{2D}, \boldsymbol{M}_{3D}, M)$
 $\Delta \boldsymbol{H}, \Delta \vec{\boldsymbol{V}} \leftarrow \mathrm{AdapterGNNLayer}[l](\boldsymbol{H}_c, \vec{\boldsymbol{V}}_c, \vec{\boldsymbol{X}}_c, \mathcal{E}_c, \boldsymbol{e}_{ij})$
 $\boldsymbol{M}_{upd} \leftarrow \boldsymbol{M}_{2D} \vee \boldsymbol{M}_{3D}$
 # *Only maintain real nodes*
 **return** $\boldsymbol{M}_{upd} \odot (\Delta \boldsymbol{H}[: M]), \boldsymbol{M}_{upd} \odot (\Delta \vec{\boldsymbol{V}}[: M])$

##### 1.3.6 Algorithm for Adapted Geometric Equivariant Transformer

**Algorithm 4** Architecture of the Adapted Geometric Equivariant Transformer

**def** AdaptedGeometricEquivariantTransformer($\boldsymbol{H}, \vec{\boldsymbol{X}}, \mathcal{G}_c, \boldsymbol{M}_{2D}, \boldsymbol{M}_{3D}$):
### *Initial feature construction*
  $\boldsymbol{H}_c^{(0)}, \vec{\boldsymbol{V}}_c^{(0)} \leftarrow \text{GraphInitialization}(\boldsymbol{H}_c, \vec{\boldsymbol{X}}_c)$
### *Collect Conditions*
  $\mathcal{V}_c, \mathcal{E}_c \leftarrow \mathcal{G}_c$
  $[\boldsymbol{H}_r, \vec{\boldsymbol{X}}_r], [\boldsymbol{H}_d, \vec{\boldsymbol{X}}_d] \leftarrow \mathcal{V}_c$
### *Stack of L adapted equivariant transformer layers*
  **for** $l = 0, 1, \ldots, L-1$ **do**
    $[\boldsymbol{H}^{(l)}, \vec{\boldsymbol{V}}^{(l)}] \leftarrow [\boldsymbol{H}^{(l)}, \vec{\boldsymbol{V}}^{(l)}] + \text{EquivariantSelfAttention}(\text{LN}(\boldsymbol{H}^{(l)}), \vec{\boldsymbol{V}}^{(l)}, \vec{\boldsymbol{X}})$
    $[\boldsymbol{H}^{(l+1)}, \vec{\boldsymbol{V}}^{(l+1)}] \leftarrow [\boldsymbol{H}^{(l)}, \vec{\boldsymbol{V}}^{(l)}] + \text{EquivariantFFN}(\text{LN}(\boldsymbol{H}^{(l)}), \vec{\boldsymbol{V}}^{(l)})$
    $[\boldsymbol{H}_r^{(l+1)}, \vec{\boldsymbol{X}}_r^{(l+1)}] \leftarrow [\boldsymbol{H}^{(l+1)} \| \boldsymbol{H}_r, \vec{\boldsymbol{X}}]$
    $[\boldsymbol{H}_d^{(l+1)}, \vec{\boldsymbol{X}}_d^{(l+1)}] \leftarrow [\boldsymbol{M}_{3D} \odot \boldsymbol{H}^{(l+1)} \| \boldsymbol{H}_d, \vec{\boldsymbol{X}}_d]$
    $[\Delta \boldsymbol{H}^{(l+1)}, \Delta \vec{\boldsymbol{V}}^{(l+1)}] \leftarrow \text{GraphAdapter}(\boldsymbol{H}_r^{(l+1)}, \vec{\boldsymbol{V}}_r^{(l+1)}, \vec{\boldsymbol{X}}_r^{(l+1)}, \boldsymbol{H}_d^{(l+1)}, \vec{\boldsymbol{X}}_d^{(l+1)}, \boldsymbol{M}_{2D}, \boldsymbol{M}_{3D}, l)$
    $[\boldsymbol{H}^{(l+1)}, \vec{\boldsymbol{V}}^{(l+1)}] \leftarrow [\Delta \boldsymbol{H}^{(l+1)}, \Delta \vec{\boldsymbol{V}}^{(l+1)}]$
  **end for**
  **return** $LN(\boldsymbol{H}^{(L)}), \vec{\boldsymbol{V}}^{(L)}$

#### 1.4 Training Objectives

##### 1.4.1 Losses of All-atom Variational Auto-Encoder

**KL divergence regularization.** To regularize the scale of the latent states $\boldsymbol{z}_i$ and coordinates $\vec{\boldsymbol{z}}_i$, we implement Kullback–Leibler (KL) divergence constraints relative to the prior distributions $\mathbb{N}(\boldsymbol{0}, \boldsymbol{I})$ and $\mathbb{N}(\vec{\boldsymbol{r}}_i, \boldsymbol{I})$[22], where $\vec{\boldsymbol{r}}_i$ represents the center of mass for all atoms in block $i$:

$$\mathcal{L}_{KL}(i) = \lambda_1 \cdot D_{\text{KL}}(\mathcal{N}(\boldsymbol{0}, \boldsymbol{I}) \| \mathcal{N}(\boldsymbol{\mu}_i, \text{diag}(\boldsymbol{\sigma}_i))) + \lambda_2 \cdot D_{\text{KL}}(\mathcal{N}(\vec{\boldsymbol{r}}_i, \boldsymbol{I}) \| \mathcal{N}(\vec{\boldsymbol{\mu}}_i, \text{diag}(\vec{\boldsymbol{\sigma}}_i))), \tag{15}$$

where $D_{\text{KL}}$ denotes the KL divergence, $\lambda_1$ and $\lambda_2$ control the strength of regularization for the latent states and coordinates, respectively. During training, we further perturb $\vec{\boldsymbol{z}}_i$ by adding random noise sampled from $\mathcal{N}(\boldsymbol{0}, \boldsymbol{I})$ before feeding it into the decoder. This necessity arises from the inherent difficulty the diffusion module faces when generating accurate positions within the latent domain, resulting in coordinate discrepancies throughout the generative phase. By proactively integrating stochastic noise into the training stage, the decoder acquires the capability to manage such inaccuracies with significantly enhanced robustness.

**Reconstruction loss.** The reconstruction loss includes cross entropy (CE) for block types and bond types, as well as the mean square error (MSE) for vector fields:

$$\mathcal{L}_{\text{rec}}(i) = \text{CE}(p(\hat{s}_i), p(s_i)) + \mathbb{E}_{t \sim U(0,1)} \left[ \sum_{j,p,q} \text{CE}(p(\hat{b}_{ij,pq}^t), p(b_{ij,pq}^t)) \right] + \mathbb{E}_{t \sim U(0,1)} \left[ \text{MSE}(\hat{\vec{\boldsymbol{V}}}_i^t, \vec{\boldsymbol{V}}_i^t) \right], \tag{16}$$

where $p(\hat{s}_i), p(\hat{b}_{ij,pq}^t)$, and $\hat{\vec{\boldsymbol{V}}}_{i,\text{gt}}^t$ denote the ground-truth block types, bond types, and vector fields, respectively.

**Pair-wise distance loss.** To facilitate superior modeling of local geometric structures and atomic interactions, a pairwise distance penalty is incorporated within the immediate vicinity of every atom. Given that structural configurations remain highly disordered at elevated time scales, this particular loss term is exclusively activated for intervals where :

$$\mathcal{L}_{\text{dist}}(i,t) = \frac{\sum_{p \in \mathcal{A}_i,(j,q)\in\mathcal{N}_i(p)} |\hat{d}^0_{ij,pq} - d^0_{ij,pq}|}{\sum_{p\in\mathcal{A}_i} |\mathcal{N}_i(p)|},\ \mathcal{L}_{\text{dist}}(i) = \mathbb{E}_{t\sim U(0,1)}[\mathbb{I}_{t<0.25} \cdot \mathcal{L}_{\text{dist}}(i,t)], \tag{17}$$

where $\mathcal{N}_i(p) = (j,q) \mid \hat{d}^0_{ij,pq} < 6.0$ denotes all atoms within 6Å distance to the $p$-th atom in block $i$ in the ground-truth structure.

**Overall autoencoder objective.** To enhance the information density within the binding site's latent point cloud $\mathcal{G}_y$, our approach incorporates KL divergence alongside reconstruction loss applied to a 5% subset of the binding site residues. Letting $\tilde{\mathcal{G}}_y$ represent this randomly sampled group of residues, the total objective function governing the autoencoder is defined as:

$$\mathcal{L}_{\text{AE}} = \sum_{i\in\mathcal{G}_x\cup\tilde{\mathcal{G}}_y} \frac{\mathcal{L}_{\text{KL}}(i) + \mathcal{L}_{\text{rec}}(i) + \lambda_{\text{dist}}\mathcal{L}_{\text{dist}}(i)}{|\mathcal{G}_x \cup \tilde{\mathcal{G}}_y|}, \tag{18}$$

where $\lambda_{\text{dist}}$ balances the contribution of the distance loss.

**Training details.** Teacher forcing is utilized throughout the training phase, during which the ground-truth atom categories $\hat{\boldsymbol{A}}_i$ and intra-block linkages $\hat{\boldsymbol{B}}_i$ are provided to the structural module; additionally, inter-block connectivity $\hat{\boldsymbol{B}}_{ij}$ is supplied with a 50% probability. In the inference stage, the model concurrently produces complete atomic configurations and inter-block connections, followed by a refinement process via a secondary encoding-decoding iteration that incorporates the predicted inter-block bonds as auxiliary inputs.

For every training iteration, time steps $t$ are selected through random sampling, where the input spatial coordinates $\vec{\boldsymbol{X}}_i^t$ are derived by interpolating between the reference ground-truth positions $\vec{\boldsymbol{X}}_i^0$ and the starting Gaussian-distributed coordinates $\vec{\boldsymbol{X}}_i^1$. The comprehensive procedures for training and sample generation are detailed in Algorithm 5 and 8, respectively.

###### *1.4.2 Losses of Conditional Latent Diffusion Model*

The training objective for the diffusion model minimizes the MSE between the predicted noise and the actual noise added during the forward process (Eq. 11), which is given as:

$$\mathcal{L}_{LDM} = \mathbb{E}_{t\sim \text{U}(1...T)}[\frac{\sum_i \|\boldsymbol{\epsilon}_i - \boldsymbol{\epsilon}_\theta(\mathcal{Z}_x^t, \mathcal{Z}_y, \mathcal{A}_\eta(\mathcal{G}_c), t)[i]\|^2}{|\mathcal{Z}_x^t|}], \tag{19}$$

where $|\mathcal{Z}_x^t|$ indicates the number of nodes in the latent point cloud $\mathcal{Z}_x^t$. The overall training and sampling algorithms are included in Algorithm 6, 7.

##### 1.5 Training Regimen

###### *1.5.1 Dynamic Batching for Training*

As all model components are instantiated using the geometric equivariant transformer (Algorithm 2), the computational and memory costs scale quadratically with input length. To accommodate the quadratic computational complexity of such attention-based architectures, we adopt a dynamic batching strategy during training to improve computational efficiency by avoiding paddings. Specifically, samples are incrementally added to a batch until the constraint $\sum_i n_i^2 \leq C$ is reached, where $n_i$ denotes the length of the $i$-th sample and $C$ is a predefined capacity threshold. Batch construction terminates when inclusion of the next sample would violate this constraint. The definition of input length depends on the module being trained. For the all-atom variational autoencoder and the supervised confidence module, $n_i$ corresponds to the number of atoms, as these components operate directly on atom-level geometries. For the latent diffusion model, $n_i$ is defined as the number of building blocks, reflecting the block-level latent representation. The capacity thresholds $C$ used for different datasets and training stages are summarized in Supplementary Table 4. These thresholds were determined empirically by testing each dataset separately, progressively increasing the capacity limit and training for one epoch until GPU memory limits were reached.

##### 1.5.2 Algorithm for Training All-atom Variational Auto-Encoder

**Algorithm 5** Training Algorithm of the Iterative Full-Atom Autoencoder

**def** Encode($\mathcal{E}_\phi$, $\mathcal{G}$):
### *Encode input graph*
  $\{(\boldsymbol{\mu}_i, \boldsymbol{\sigma}_i, \vec{\boldsymbol{\mu}}_i, \vec{\boldsymbol{\sigma}}_i)\} \leftarrow \mathcal{E}_\phi(\mathcal{G})$
### *Sample latent variables via reparameterization*
  $\{(\boldsymbol{\epsilon}_i, \vec{\boldsymbol{\epsilon}}_i)\} \sim \mathcal{N}(\mathbf{0}, \boldsymbol{I})$
  $\mathcal{Z} \leftarrow \{(\boldsymbol{\mu}_i + \boldsymbol{\epsilon}_i \odot \boldsymbol{\sigma}_i, \vec{\boldsymbol{\mu}}_i + \vec{\boldsymbol{\epsilon}}_i \odot \vec{\boldsymbol{\sigma}}_i)\}$
  **return** $\mathcal{Z}$

**def** TrainIterativeAutoencoder($\mathcal{S}$):
  Initialize $\mathcal{E}_\phi$, $\mathcal{D}_\xi$
  **while** $\phi, \xi$ have not converged **do**
    Sample $(\mathcal{G}_x, \mathcal{G}_y) \sim \mathcal{S}$
#   *Encode binder and binding site separately*
    $\mathcal{Z}_x, \mathcal{Z}_y \leftarrow \text{Encode}(\mathcal{E}_\phi, \mathcal{G}_x), \text{Encode}(\mathcal{E}_\phi, \mathcal{G}_y)$
#   *Sample 5% binding-site residues for reconstruction*
    Sample $\tilde{\mathbb{I}}_y \subseteq \mathbb{I}_y$
#   *Define nodes to be reconstructed*
    $\mathbb{I} \leftarrow \tilde{\mathbb{I}}_y \cup \mathbb{I}_x$
#   *Predict block-level types*
    $\{s_i \mid i \in \mathbb{I}\} \leftarrow \mathcal{D}_{\xi_1}(\mathcal{Z}_x, \mathcal{Z}_y)$
#   *Sample time and initial atomic coordinates at $t = 1$*
    Sample $t \sim U(0,1), \vec{\boldsymbol{X}}_i^1 \sim \mathcal{N}(\vec{\boldsymbol{z}}_i, \boldsymbol{I})$
#   *Compute ground-truth atomic displacements*
    $\hat{\vec{\boldsymbol{V}}}_i^t \leftarrow \vec{\boldsymbol{X}}_i^0 - \vec{\boldsymbol{X}}_i^1$
#   *Interpolate atomic coordinates at time $t$*
    $\vec{\boldsymbol{X}}_i^t \leftarrow t \cdot \vec{\boldsymbol{X}}_i^1 + (1-t) \cdot \vec{\boldsymbol{X}}_i^0$
#   *Reveal inter-block bonds with 50% probability*
    $\tilde{\boldsymbol{B}}_{ij} \leftarrow \phi$ if $p < 0.5$ else $\hat{\boldsymbol{B}}_{ij}$, $p \sim U(0,1)$
#   *Decode atoms and motions with teacher forcing*
    $\{(\boldsymbol{H}_i^t, \vec{\boldsymbol{V}}_i^t) \mid i \in \mathbb{I}\} = \mathcal{D}_{\xi_2}(\{(\hat{\boldsymbol{A}}_i, \hat{\boldsymbol{B}}_i \cup \tilde{\boldsymbol{B}}_{ij}, \vec{\boldsymbol{X}}_i^t)\}, \mathcal{Z}_x, \mathcal{Z}_y, \mathcal{G}_y, t)$
#   *Predict bond types*
    $b_{ij,pq}^t = \text{Softmax}(\text{MLP}(\boldsymbol{h}_{i,p}^t + \boldsymbol{h}_{j,q}^t)), i \neq j$
#   *Compute autoencoder objective*
    $\mathcal{L}_{\text{AE}} = \sum_{i \in \mathbb{I}} (\mathcal{L}_{\text{KL}}(i) + \mathcal{L}_{\text{rec}}(i) + \lambda_{\text{dist}} \mathcal{L}_{\text{dist}}(i)) / |\mathbb{I}|$
    $\phi, \xi \leftarrow \text{optimizer}(\mathcal{L}_{AE}; \phi, \xi)$
  **end while**
  **return** $\mathcal{E}_\phi$, $\mathcal{D}_\xi$

##### 1.5.3 Algorithm for Training Conditional Latent Diffusion Model

**Algorithm 6** Training Algorithm of Conditional Latent Diffusion Model

**def** TrainLatentDiffusion($\mathcal{S}$, $\mathcal{E}_\phi$, $\mathcal{D}_\xi$):
\# *Fix parameters $\phi$ and $\xi$*
Initialize $\tilde{\boldsymbol{\epsilon}}_\theta, \mathcal{A}_\eta$
**while** $\theta$ have not converged **do**
Sample $(\mathcal{G}_x, \mathcal{G}_y) \sim \mathcal{S}$
Sample $\mathcal{G}_c$ by conditional sampler
\# *Encoding the binder and context*
$\mathcal{Z}_x, \mathcal{Z}_y \leftarrow \text{Encode}(\mathcal{E}_\phi, \mathcal{G}_x), \text{Encode}(\mathcal{E}_\phi, \mathcal{G}_y)$
\# *Sample noises*
$t \sim \mathbf{U}(1, T)$, $\{(\boldsymbol{\epsilon}_i, \vec{\boldsymbol{\epsilon}}_i)\} \sim \mathcal{N}(\mathbf{0}, \boldsymbol{I})$
\# *Sample intermediate states at time t*
$\mathcal{Z}_x^t \leftarrow \{(\boldsymbol{z}_i^t, \vec{\boldsymbol{z}}_i^t) \mid i \in \mathbb{I}_x, [\boldsymbol{z}_i^t, \vec{\boldsymbol{z}}_i^t] = \sqrt{\bar{\alpha}^t}[\boldsymbol{z}_i^0, \vec{\boldsymbol{z}}_i^0] + \sqrt{1-\bar{\alpha}^t}[\boldsymbol{\epsilon}_i, \vec{\boldsymbol{\epsilon}}_i]\}$
$\mathcal{L}_{LDM}^t = \sum_i \|[\boldsymbol{\epsilon}_i, \vec{\boldsymbol{\epsilon}}_i] - \tilde{\boldsymbol{\epsilon}}_\theta(\mathcal{Z}_x^t, \mathcal{Z}_y, \mathcal{A}_\eta(\mathcal{G}_c), t)[i]\|^2 / |\mathcal{Z}_x^t|$
$\theta, \eta \leftarrow \text{optimizer}(\mathcal{L}_{LDM}^t; \theta, \eta)$
**end while**
**return** $\tilde{\boldsymbol{\epsilon}}_\theta, \mathcal{A}_\eta$

#### 1.6 Inference Regimen

##### 1.6.1 Algorithm for Sampling from Conditional Latent Diffusion Model

**Algorithm 7** Sampling Algorithm of the Conditional Latent Diffusion Model

**def** SamplefromConditionalLatentDiffusion($\mathcal{E}_\phi$, $\mathcal{D}_\xi$, $\boldsymbol{\epsilon}_\theta$, $\mathcal{A}_\eta$, $\mathcal{G}_c$, $\mathcal{G}_y$, $T$, $N$):
\# *Encode the conditioning binding site into the latent space*
$\mathcal{Z}_y \leftarrow \text{Encode}(\mathcal{E}_\phi, \mathcal{G}_y)$
\# *Initialize the latent variables with Gaussian noise*
$\{(\boldsymbol{z}_i^T, \vec{\boldsymbol{z}}_i^T)\} \sim \mathcal{N}(\mathbf{0}, \boldsymbol{I})$
**for** $t$ in $T$, $T-1, \cdots, 1$ **do**
\# *Iterative reverse diffusion for the molecular binder's latent nodes*
$\mathcal{Z}_x^t \leftarrow \{(\boldsymbol{z}_i^t, \vec{\boldsymbol{z}}_i^t) \mid i \in \mathbb{I}_x\}$
$\vec{\boldsymbol{u}}_i^t \leftarrow [\boldsymbol{z}_i^t, \vec{\boldsymbol{z}}_i^t]$
$\boldsymbol{\varepsilon}_i = [\boldsymbol{\epsilon}_i, \vec{\boldsymbol{\epsilon}}_i] \sim \mathcal{N}(\mathbf{0}, \boldsymbol{I})$
\# *Compute the denoising update for the latent point cloud*
$\vec{\boldsymbol{u}}_i^{t-1} \leftarrow \frac{1}{\sqrt{\alpha^t}}(\vec{\boldsymbol{u}}_i^t - \frac{\beta^t}{\sqrt{1-\bar{\alpha}^t}}\tilde{\boldsymbol{\epsilon}}_\theta(\mathcal{Z}_x^t, \mathcal{Z}_y, \mathcal{A}_\eta(\mathcal{G}_c), t)[i]) + \sqrt{\beta^t}\boldsymbol{\varepsilon}_i$
$[\boldsymbol{z}_i^{t-1}, \vec{\boldsymbol{z}}_i^{t-1}] \leftarrow \vec{\boldsymbol{u}}_i^{t-1}$
**end for**
$\mathcal{Z}_x \leftarrow \{(\boldsymbol{z}_i^0, \vec{\boldsymbol{z}}_i^0) \mid i \in \mathbb{I}_x\}$
\# *Reconstruct the molecular graph via the iterative decoder*
$\mathcal{G}_x \leftarrow \text{Decode}(\mathcal{E}_\phi, \mathcal{D}_\xi, \mathcal{G}_y, \mathcal{Z}_x, \mathcal{Z}_y, N)$
**return** $\mathcal{G}_x$

##### 1.6.2 Algorithm for Sampling from All-atom Variational Auto-Encoder

**Algorithm 8** Sampling Algorithm of the Iterative Full-Atom Autoencoder

**def** DecodeStruct($\mathcal{D}_\xi$, $\{(\boldsymbol{A}_i, \boldsymbol{B}_i, \boldsymbol{B}_{ij})\}$, $\mathcal{Z}_x$, $\mathcal{Z}_y$, $\mathcal{G}_y$):
  $\Delta t \leftarrow 1.0/N$
  # *Initialize atomic coordinates with Gaussian noise*
  $\vec{\boldsymbol{X}}_i^0 \sim \mathcal{N}(\vec{\boldsymbol{z}}_i, \boldsymbol{I})$
  **for** $n$ in $0, 1, \cdots, N-1$ **do**
    $t \leftarrow 1.0 - n \cdot \Delta t$
    $\{(\boldsymbol{H}_i^t, \vec{\boldsymbol{V}}_i^t) \mid i \in \mathbb{I}_x\} \leftarrow \mathcal{D}_{\xi_2}(\{(\boldsymbol{A}_i, \boldsymbol{B}_i \cup \boldsymbol{B}_{ij}, \vec{\boldsymbol{X}}_i^t)\}, \mathcal{Z}_x, \mathcal{Z}_y, \mathcal{G}_y, t)$
    # *Update atomic coordinates by integration*
    $\vec{\boldsymbol{X}}_i^{t-\Delta t} \leftarrow \vec{\boldsymbol{X}}_i^t + \Delta t \vec{\boldsymbol{V}}_i^t$
    $\boldsymbol{H}_i^{t-\Delta t} \leftarrow \boldsymbol{H}_i^t$
  **end for**
  # *Predict inter-block bonds from final decoded features*
  $\boldsymbol{B}_{ij} \leftarrow \{\text{Argmax}(\text{MLP}(\boldsymbol{h}_{i,p}^0 + \boldsymbol{h}_{j,q}^0)) \mid i \neq j, d_{ij,pq}^0 < 3.5\text{Å}\}$ if $\boldsymbol{B}_{ij} = \emptyset$ else $\boldsymbol{B}_{ij}$
  **return** $\{(\vec{\boldsymbol{X}}_i^0, \boldsymbol{B}_{ij}) \mid i \in \mathbb{I}_x\}$

**def** SamplefromIterativeAutoencoder($\mathcal{E}_\phi$, $\mathcal{D}_\xi$, $\mathcal{G}_y$, $\mathcal{Z}_x$, $\mathcal{Z}_y$, $N$):
  # *Predict block-level types for binder nodes*
  $\{s_i \mid i \in \mathbb{I}_x\} \leftarrow \mathcal{D}_{\xi_1}(\mathcal{Z}_x, \mathcal{Z}_y)$
  # *Retrieve atom types and intra-block bonds from the vocabulary*
  $\boldsymbol{A}_i, \boldsymbol{B}_i \leftarrow \text{Lookup}(\mathrm{s_i}, \mathbb{V})$
  # *Decode atomic coordinates and inter-block bonds*
  $\{(\vec{\boldsymbol{X}}_i, \boldsymbol{B}_{ij}) \mid i \in \mathbb{I}_x\} \leftarrow \text{DecodeStruct}(\mathcal{D}_\xi, \{(\boldsymbol{A}_i, \boldsymbol{B}_i, \emptyset)\}, \mathcal{Z}_x, \mathcal{Z}_y, \mathcal{G}_y)$
  $\mathcal{G}_x \leftarrow (\{(\boldsymbol{A}_i, \vec{\boldsymbol{X}}_i) \mid i \in \mathbb{I}_x\}, \{(\boldsymbol{B}_i, \boldsymbol{B}_{ij}) \mid i \in \mathbb{I}_x, i \neq j\})$
  # *Re-encode the generated binder graph*
  $\mathcal{Z}_x \leftarrow \text{Encode}(\mathcal{E}_\phi, \mathcal{G}_x)$
  # *Refine geometry with fixed inter-block bonds*
  $\{(\vec{\boldsymbol{X}}_i, \boldsymbol{B}_{ij}) \mid i \in \mathbb{I}_x\} \leftarrow \text{DecodeStruct}(\mathcal{D}_\xi, \{(\boldsymbol{A}_i, \boldsymbol{B}_i, \boldsymbol{B}_{ij})\}, \mathcal{Z}_x, \mathcal{Z}_y, \mathcal{G}_y)$
  $\mathcal{G}_x \leftarrow (\{(\boldsymbol{A}_i, \vec{\boldsymbol{X}}_i) \mid i \in \mathbb{I}_x\}, \{(\boldsymbol{B}_i, \boldsymbol{B}_{ij}) \mid i \in \mathbb{I}_x, i \neq j\})$
  **return** $\mathcal{G}_x$

#### 1.7 Soft Physical Corrections on VAE Reconstruction

To improve the physical realism of the generated conformations, additional constraints can be integrated into the proposed framework. In the following, we describe two representative strategies already implemented for alleviating steric clashes and refining local structural geometries.

##### 1.7.1 Clashes Avoidance

Clashes frequently occur in structures generated by deep learning models. In AnewOmni, we disentangle the modeling of global arrangement and local atomic geometry via latent diffusion coupled with a full-atom variational autoencoder. This design enables explicit treatment of atomic overlaps during the decoding phase by injecting repulsive interactions into the predicted vector field (Eq.7) throughout the iterative reconstruction process. Concretely, at each decoding step, we evaluate pairwise distances between the generated atoms and those in the surrounding context (i.e., the binding site). When the overlap between their van der Waals radii exceeds a predefined tolerance $\delta$[23], a corrective force is applied; in this work, $\delta$ is set to 0.3Å.

Formally, let $r_i$ and $r_j$ denote the van der Waals radii of atoms $i$ and $j$, and let $\vec{x}_i^t$ and $\vec{x}_j^t$ represent their positions at time step $t$. The strength of the repulsive interaction is given by:

$$f_{ij} = \begin{cases} r_i + r_j - \delta - \|\vec{x}_i^t - \vec{x}_j^t\|, & \|\vec{x}_i^t - \vec{x}_j^t\| < r_i + r_j - \delta, \\ 0, & else, \end{cases} \tag{20}$$

During each decoding iteration, steric conflicts are mitigated by further adjusting the atomic coordinates according to $\vec{x}_i^t = \vec{x}_i^t + \sum_j \frac{\vec{x}_i^t - \vec{x}_j^t}{\|\vec{x}_i^t - \vec{x}_j^t\|} \cdot f_{ij}$, where the summation over $j$ spans all atoms in the fixed contextual environment.

##### 1.7.2 Bond Validity

Prior work commonly adopts OpenBabel[24] to infer chemical bonds from pairwise interatomic distances. By contrast, our unified framework resolves bonding by combining block-specified intra-block connections with inter-block bonds predicted by the model. Although this prediction-based strategy offers increased flexibility, enabling, for example, straightforward extension to non-canonical amino acids—it can also lead to challenges such as valency violations and mismatches between the inferred 2D topology and the realized 3D geometry (e.g., overly long bonds or abnormal bond angles). To mitigate valency errors, predicted bonds are sorted according to their confidence scores and incorporated incrementally, discarding any bond that would violate valency limits. To enforce 2D–3D consistency, we estimate empirical distributions of bond lengths and angles, where bond lengths are binned from 1.1Å to 1.7Å with a resolution of 0.005Å, and angles are discretized from 0° to 180° in 2° intervals. A generated molecule is deemed inconsistent and rejected if more than 10

#### 1.8 Re-ranking Regimen

##### 1.8.1 Algorithm for Diffusion Likelihood Calculation

Algorithm 9 outlines the procedure for computing the diffusion likelihood of the latent features for the generated binder $\mathcal{Z}_x^0 = \{(\boldsymbol{z}_i^0, \vec{\boldsymbol{z}}_i^0) \mid i \in \mathbb{I}_x\}$. We begin with the Probability Flow ODE (PF ODE) associated with DDPM[20]:

$$\frac{\mathrm{d}\vec{\boldsymbol{u}}_i^t}{\mathrm{d}t} = \tilde{\boldsymbol{f}}_\theta(\vec{\boldsymbol{u}}_i^t, t) \equiv -\frac{1}{2}\beta^t\Big(\vec{\boldsymbol{u}}_i^t - \frac{1}{\sqrt{(1-\bar{\alpha}^t)}}\boldsymbol{\epsilon}_\theta(\mathcal{Z}_x^t, \mathcal{Z}_y, \mathcal{A}(\mathcal{G}_c), t)\Big). \tag{21}$$

Following previous literature[25], the log-likelihood of latent features $\vec{\boldsymbol{u}}_i^0 = [\boldsymbol{z}_i^0, \vec{\boldsymbol{z}}_i^0]$ is given by:

$$\log p_0(\vec{\boldsymbol{u}}_i^0) = \log p_T(\vec{\boldsymbol{u}}_i^T) + \int_0^T \nabla \cdot \tilde{\boldsymbol{f}}_\theta(\vec{\boldsymbol{u}}_i^t, t)\,\mathrm{d}t, \tag{22}$$

where $\nabla\cdot$ denotes the divergence operation, and $\{\vec{\boldsymbol{u}}_i^t\}_{t=0}^T$ can be obtained by solving Equation 21 with the initial condition $\vec{\boldsymbol{u}}_i^0$ at $t = 0$. However, numerically solving Equation 21 can be unstable in practice. We therefore compute the likelihood directly using trajectories generated by DDPM. Empirically, this approach yields correlations with experimental metrics comparable to those obtained using DDIM, as shown in Supplementary Fig. 13.

Since directly computing the divergence $\nabla \cdot \tilde{\boldsymbol{f}}_\theta(\vec{\boldsymbol{u}}_i^t, t)$ is expensive, we adopt the Skilling-Hutchinson trace estimator[26,27] to compute the term:

$$\nabla \cdot \tilde{\boldsymbol{f}}_\theta(\vec{\boldsymbol{u}}_i^t, t) = \mathbb{E}_{p(\boldsymbol{\varepsilon})}\left[\boldsymbol{\varepsilon}^\top \nabla \tilde{\boldsymbol{f}}_\theta(\vec{\boldsymbol{u}}_i^t, t)\boldsymbol{\varepsilon}\right], \tag{23}$$

where $p(\boldsymbol{\varepsilon}) = \mathcal{N}(\boldsymbol{0}, \boldsymbol{I})$, and $\nabla\tilde{\boldsymbol{f}}_\theta$ denotes the Jacobian of $\tilde{\boldsymbol{f}}_\theta$. Specifically, the vector-Jacobian product $\boldsymbol{\varepsilon}^\top \nabla \tilde{\boldsymbol{f}}_\theta(\vec{\boldsymbol{u}}_i^t, t)$ can be efficiently computed through automatic differentiation. In practice, the time integration of Equation 22 is discretized into $T$ (the trajectory length for solving Equation 21, which is 100 in this paper) uniform bins, and the expectation of Equation 23 is estimated using $L$ independent runs for computation.

Next, we aggregate the computed log-likelihoods for the latent states $\boldsymbol{z}_i^t$ and the coordinates $\vec{\boldsymbol{z}}_i^t$, averaging over their respective dimensions to prevent either term from dominating the total. As a result, we obtain a unique score, denoted by $w$, for each generated binder after aggregation.

Finally, to eliminate variations in the magnitude of the computed score induced by the number of atoms, we apply the following normalization with the number of latent points and a proper scaling factor $\alpha \in \mathbb{R}_+$:

$$\hat{w} = \alpha \cdot w / \operatorname{card}(\mathbb{I}_x) \tag{24}$$

The normalized score $\hat{w}$ serves as a computational efficient metric for ranking generated binders with varying number of atoms.

**Algorithm 9** Normalized Log-Likelihood Computation

**def** ComputeNormalizedLogLikelihood($\boldsymbol{\epsilon}_\theta, \{\mathcal{Z}_x^t\}_{t=0}^T, \mathcal{Z}_y, \mathcal{A}_\eta, \mathcal{G}_c, T, L, \alpha$):
  $w \leftarrow 0$
  **for** $t$ in $T, T-1, \cdots, 1$ **do**
    $\{(\boldsymbol{z}_i^t, \vec{\boldsymbol{z}}_i^t) \mid i \in \mathbb{I}_x\} \leftarrow \mathcal{Z}_x^t$
    $\vec{\boldsymbol{u}}_i^t \leftarrow [\boldsymbol{z}_i^t, \vec{\boldsymbol{z}}_i^t]$
### *Enabling Gradients for Backpropagation*
    $\vec{\boldsymbol{u}}_i^t \leftarrow \text{EnableGrad}(\vec{\boldsymbol{u}}_i^t)$
    $\{(\boldsymbol{\epsilon}_i, \vec{\boldsymbol{\epsilon}}_i) \mid i \in \mathbb{I}_x\} \leftarrow \boldsymbol{\epsilon}_\theta(\mathcal{Z}_x^t, \mathcal{Z}_y, \mathcal{A}_\eta(\mathcal{G}_c), t)$
### *Computing the Drift Term of the Probability Flow ODE*
    $\vec{\boldsymbol{f}}_i^t \leftarrow -\frac{\beta^t}{2}(\vec{\boldsymbol{u}}_i^t - \frac{1}{\sqrt{1-\bar{\alpha}^t}}[\boldsymbol{\epsilon}_i, \vec{\boldsymbol{\epsilon}}_i])$
### *Computing the Divergence via Skilling-Hutchinson Trace Estimator*
    **for** $k$ in $1, \cdots, L$ **do**
      $[\boldsymbol{\varepsilon}_i, \vec{\boldsymbol{\varepsilon}}_i] \leftarrow \mathcal{N}(\mathbf{0}, \boldsymbol{I})$
      $a_i^t \leftarrow \text{sum}([\boldsymbol{\varepsilon}_i, \vec{\boldsymbol{\varepsilon}}_i] \odot \vec{\boldsymbol{f}}_i^t)$
### *Computing the Vector-Jacobian Product through Automatic Differentiation*
      $[\boldsymbol{\delta}_i^t, \vec{\boldsymbol{\delta}}_i^t] \leftarrow \partial a_i^t / \partial \vec{\boldsymbol{u}}_i^t$
### *Aggregating Likelihoods for Features and Coordinates, Averaging over Their Respective Dimensions to Avoid Dominance by Either Term*
      $w \leftarrow w + \frac{1}{TL}\sum_{i\in\mathbb{I}_x}\left(\frac{1}{d}\text{sum}(\boldsymbol{\varepsilon}_i \odot \boldsymbol{\delta}_i^t) + \frac{1}{3n_i}\text{sum}(\vec{\boldsymbol{\varepsilon}}_i \odot \vec{\boldsymbol{\delta}}_i^t)\right)$
    **end for**
  **end for**
### *Eliminating the Dependence of the Number of latent points, and scaling by $\alpha$ to obtain a more readable scale*
  $\hat{w} \leftarrow \alpha \cdot w / \text{card}(\mathbb{I}_x)$
  **return** $\hat{w}$

##### 1.8.2 Confidence Assessment via Pairwise Distance Error Prediction

To evaluate the reliability of the generated molecules without relying on ground-truth references, we introduce a confidence model $\mathcal{C}_\psi$, parameterized by $\psi$. This auxiliary module estimates the structural quality of the generated binder $\hat{\mathcal{G}}_x$ within the context of the environment $\mathcal{G}_y$. Specifically, it predicts the Pairwise Distance Error (PDE) distribution for interactions within the ligand (intra-molecular) and between the ligand and the environment (inter-molecular).

We define the ground-truth PDE for an atom pair $(i, j)$ as the absolute difference between the distances in the generated conformation $\hat{\vec{\boldsymbol{X}}}$ and the reference $\vec{\boldsymbol{X}}$: $\boldsymbol{E}_{ij} = |\|\hat{\vec{\boldsymbol{x}}}_i - \hat{\vec{\boldsymbol{x}}}_j\|_2 - \|\vec{\boldsymbol{x}}_i - \vec{\boldsymbol{x}}_j\|_2|$. Then, we discretize the error space into $K$ bins $\{b_k\}_{k=1}^K$ and treat the prediction as a classification task. The confidence model predicts a probability distribution $\boldsymbol{P}_{ij} \in \Delta^{K-1}$ indicating the likelihood of $\boldsymbol{E}_{ij}$ falling into each bin. From this, we derive two primary quality metrics:

1. **Ligand PDE** ($\text{PDE}_{\text{lig}}$): The expected error of intra-ligand pairs, reflecting internal structural consistency.

2. **Complex PDE** ($\text{PDE}_{\text{cplx}}$): The expected error of pairs between the ligand and the binding pocket, measuring the accuracy of the binding pose.

Architecturally, $\mathcal{C}_\psi$ consists of a encoder which is instantiated as an equivariant transformer[17] and a pairwise product head. The encoder refines the invariant latent features $\hat{\boldsymbol{H}}$ from the base model, incorporating the generated coordinates $\hat{\vec{\boldsymbol{X}}}$. Subsequently, the node embeddings are projected into $2K$ subspaces, representing $K$ error bins for inter and intra-molecular errors, and the logits $\hat{\boldsymbol{L}}_{ij}$ for each pair $(i, j)$ are computed via the dot product of their projected features.

The training objective minimizes the cross-entropy loss between the predicted distributions $\hat{\boldsymbol{P}}_{ij} = \mathrm{Softmax}(\hat{\boldsymbol{L}}_{ij})$ and the discretized ground-truth errors. We identify the sets of valid atom pairs for intra-ligand interactions ($\boldsymbol{M}_{\text{lig}}$) and ligand-context interactions ($\boldsymbol{M}_{\text{cplx}}$). The total confidence loss $\mathcal{L}_{\text{conf}}$ is formulated as:

$$\mathcal{L}_{\text{conf}} = \mathcal{L}_{\text{lig}} + \mathcal{L}_{\text{cplx}} \tag{25}$$

where each term averages the cross-entropy over the corresponding set of pairs:

$$\mathcal{L}_{\text{type}} = \frac{1}{|\boldsymbol{M}_{\text{type}}|} \sum_{(i,j)\in \boldsymbol{M}_{\text{type}}} \mathrm{CE}(\boldsymbol{P}_{ij}, \mathbf{Y}_{ij}) \tag{26}$$

Here, $\mathbf{Y}_{ij}$ represents the ground-truth bin index for error $\boldsymbol{E}_{ij}$.

During inference, the final confidence score $s_{\text{conf}}$ is computed as a weighted average of $\text{PDE}_{\text{lig}}$ and $\text{PDE}_{\text{cplx}}$, where a lower score indicates higher predicted structural quality. The detailed training and inference procedures are provided in Algorithm 10 and Algorithm 11.

**Algorithm 10** Training Algorithm of Confidence Model

**def** TrainConfidence($\mathcal{S}, \mathcal{E}_\phi, \mathcal{D}_\xi, \boldsymbol{\epsilon}_\theta$):
  Initialize confidence model $\mathcal{C}_\psi$
  **while** $\psi$ have not converged **do**
    Sample $(\mathcal{G}_x, \mathcal{G}_y) \sim \mathcal{S}$
### *Generate structure with frozen base model*
    $\hat{\mathcal{G}}_x, \hat{\boldsymbol{H}} \leftarrow \mathrm{Sample}(\mathcal{E}_\phi, \mathcal{D}_\xi, \boldsymbol{\epsilon}_\theta, \mathcal{G}_y)$
### *Compute ground-truth PDE matrix and labels*
    $\boldsymbol{E}_{ij} \leftarrow \big|\|\hat{\vec{\boldsymbol{x}}}_i - \hat{\vec{\boldsymbol{x}}}_j\|_2 - \|\vec{\boldsymbol{x}}_i - \vec{\boldsymbol{x}}_j\|_2\big|$
    $\mathbf{Y}_{ij} \leftarrow \mathrm{Binning}(\boldsymbol{E}_{ij}, \{b_k\})$
### *Identify valid atom pairs*
    $\boldsymbol{M}_{\text{lig}}, \boldsymbol{M}_{\text{cplx}} \leftarrow \mathrm{GetMasks}(\hat{\mathcal{G}}_x, \mathcal{G}_y)$
### *Predict probability distributions*
    $\hat{\boldsymbol{L}}_{ij} \leftarrow \mathcal{C}_\psi(\hat{\mathcal{G}}_x, \hat{\boldsymbol{H}}, \mathcal{G}_y)$
### *Optimize via Cross Entropy*
    $\mathcal{L}_{\text{lig}} \leftarrow \mathrm{CE}(\hat{\boldsymbol{L}}[\boldsymbol{M}_{\text{lig}}], \boldsymbol{Y}[\boldsymbol{M}_{\text{lig}}])$
    $\mathcal{L}_{\text{cplx}} \leftarrow \mathrm{CE}(\hat{\boldsymbol{L}}[\boldsymbol{M}_{\text{cplx}}], \boldsymbol{Y}[\boldsymbol{M}_{\text{cplx}}])$
    $\psi \leftarrow \mathrm{optimizer}(\mathcal{L}_{\text{lig}} + \mathcal{L}_{\text{cplx}}; \psi)$
  **end while**
  **return** $\mathcal{C}_\psi$

**Algorithm 11** Inference Algorithm of Confidence Model

**def** InferenceConfidence($\hat{\mathcal{G}}_x, \hat{\boldsymbol{H}}, \mathcal{G}_y, \mathcal{C}_\psi$):
### *Predict logits and probabilities*
$\hat{\boldsymbol{L}}_{ij} \leftarrow \mathcal{C}_\psi(\hat{\mathcal{G}}_x, \hat{\boldsymbol{H}}, \mathcal{G}_y)$
$\hat{\boldsymbol{P}}_{ij} \leftarrow \mathrm{Softmax}(\hat{\boldsymbol{L}}_{ij})$
### *Compute expected PDE matrix*
$\hat{\boldsymbol{E}}_{ij} \leftarrow \sum_k \hat{\boldsymbol{P}}_{ijk} \cdot b_k$
### *Aggregate errors for reliability scores*
$\boldsymbol{M}_{\text{lig}}, \boldsymbol{M}_{\text{cplx}} \leftarrow \mathrm{GetMasks}(\hat{\mathcal{G}}_x, \mathcal{G}_y)$
$\mathrm{PDE}_{\text{lig}} \leftarrow \frac{1}{|\boldsymbol{M}_{\text{lig}}|} \sum_{(i,j)\in \boldsymbol{M}_{\text{lig}}} \hat{\boldsymbol{E}}_{ij}$
$\mathrm{PDE}_{\text{cplx}} \leftarrow \frac{1}{|\boldsymbol{M}_{\text{cplx}}|} \sum_{(i,j)\in \boldsymbol{M}_{\text{cplx}}} \hat{\boldsymbol{E}}_{ij}$
$s_{\text{conf}} \leftarrow 0.5 \cdot (\mathrm{PDE}_{\text{lig}} + \mathrm{PDE}_{\text{cplx}})$
**return** $\mathrm{PDE}_{\text{lig}}, \mathrm{PDE}_{\text{cplx}}, s_{\text{conf}}$

#### 2 *In Silico* Results on Established Benchmarks

For solid assessment, the performance of the proposed model architecture as well as the mutual benefits of cross-modality training should be evaluated against literature baselines under a consistent and fair setting using the same public benchmarks. To this end, we collected representative benchmarks from well-established fields, including small molecules, peptides, and antibodies, for *in silico* evaluation. The detailed implementations and results are included below for completeness of this work, aligned with previous literature[10].

##### 2.1 Small Molecule

For small molecules, we employ the Crossdocked2020 dataset following the established splitting protocol[5]. The training set comprises 99,900 complexes, while 100 complexes are reserved for testing. An additional 100 complexes are randomly sampled from the training pool to serve as the validation set.

Performance evaluation is conducted using the CBGBench framework[28], which provides a comprehensive assessment across four primary dimensions: substructure, chemical properties, geometry, and binding interactions. Specifically, **substructure** fidelity is quantified by calculating the Jensen-Shannon Divergence (JSD) and Mean Absolute Error (MAE) between the generated and reference distributions of atoms, rings, and functional groups. **Chemical property** assessment encompasses drug-likeness (QED), synthetic accessibility (SA), octanol-water partition coefficient (LogP), and Lipinski's Rule of Five (LPSK) compliance. To evaluate structural realism, **geometry** metrics analyze the JSD of bond lengths and angles, alongside atomic and molecular clash rates. Furthermore, **interaction** quality is measured via Vina energy scores across scoring, minimization, and docking modes, as well as the distribution of interaction types between ligands and target proteins. Within each category, models are assigned a rank that is subsequently converted into a score using the formula ($N-\mathrm{rank}$), where $N$ denotes the total number of models. The final evaluation metric is defined as a weighted sum of these categorical scores, where higher values indicate superior overall performance. We refer to the original paper of this benchmark[28] as well as its official repository (`https://github.com/EDAPINENUT/CBGBench`) for detailed definition and implementation of each metric.

The resulting performance metrics and multi-dimensional rankings for small molecule *de novo* design are organized as follows: Supplementary Table 10 provides the overall scores and rankings, while Supplementary Tables 11, 12, 13 and 14 detail the substructure fidelity, chemical properties, geometric validity, and interaction analysis, respectively.

#### 2.2 Peptide

For peptides, our training pipeline utilizes the PepBench and ProtFrag datasets[22], which provide a robust foundation of 4,157 protein-peptide complexes and 70,498 synthetic samples. Model selection is performed using a validation set of 114 complexes. Following the methodology in[22], we conduct performance evaluation on the LNR dataset, a curated benchmark of 93 protein-peptide complexes[29] with peptide lengths ranging from 4 to 25 residues.

We evaluate model performance using the following set of metrics. **Amino Acid Recovery (AAR)** quantifies the fraction of residues in the generated peptide that match those in the reference sequence. Although several variants of AAR have been proposed in prior work[18,22,30,31], we follow the formulation most commonly adopted in bioinformatics[32,33]. Since AAR has been shown to be an unreliable indicator of peptide quality[22], we report it only for completeness. **Complex RMSD** measures the root-mean-square deviation of $C_\alpha$ atoms between generated and reference peptides after alignment based on the target protein. **Peptide RMSD** is defined analogously, but with alignment performed on the peptide itself. $\boldsymbol{\Delta G}$ and $\boldsymbol{\Delta\Delta G}$ evaluate binding-related properties computed using pyRosetta[34], including the predicted absolute binding free energy and the gains of $\boldsymbol{\Delta G}$ compared to the native binders. **Intra-Peptide Clash** and **Interface Clash** capture residue-level steric clashes within the peptide and between the peptide and the target protein, respectively. A clash is defined when the distance between any pair of $C_\alpha$ atoms from non-adjacent residues on the sequence falls below 3.6574 Å[35]. For each target protein in the test set, we generate 100 peptides per model for evaluation. Detailed benchmarking results for peptide design are organized in Supplementary Fig. 1a.

#### 2.3 Antibody

For antibodies, we follow standard practices in the literature[18,30] by utilizing SAbDab[11] entries deposited prior to September 24, 2021, for training and validation. To ensure rigorous evaluation, we hold out 60 antigen-antibody complexes from the RAbD dataset[36] for testing. After applying a 40% sequence identity filter relative to the test set to prevent data leakage[30], the final training and validation sets consist of 9,473 and 400 entries, respectively.

For evaluation, we focus on the complementarity-determining regions (CDRs), which play a central role in antigen recognition. Consistent with prior work[18], we define the Complementarity-Determining Regions (CDRs) using the Chothia numbering system[37]. Unlike the conventional sequence-based IMGT system[38], the Chothia system is based on structural alignment statistics, offering better robustness against sequence-level biases. Specifically, the IMGT system is susceptible to simple unigram patterns that can artificially inflate Amino Acid Recovery (AAR) to 39%; in contrast, the same patterns yield only 23% AAR under the Chothia system, effectively alleviating this issue[39]. For performance evaluation, we employ the previously defined metrics, including **AAR**, **Complex RMSD**, $\boldsymbol{\Delta G}$ and $\boldsymbol{\Delta\Delta G}$, alongside rationality indicators such as **Intra-CDR Clash** and **Interface Clash**. Detailed benchmarking results for antibody design are visualized in Supplementry Fig. 1b.

##### 2.4 Overall Weighted Score

Because each field employs multiple metrics that characterize different aspects of design quality, we derived an aggregated weighted score to enable direct and comprehensive comparison of different models. For small molecules, the benchmark itself provides an overall ranking scheme. Therefore, we use the final score normalized by its theoretical upper bound to obtain a value in the range $[0, 1]$. For peptides and antibodies, each metric is first normalized by projecting values to $[0, 1]$ as follows:

$$v_i = \begin{cases} (v_i - v_{\min})/(v_{\max} - v_{\min}), & \text{if larger values indicate better performance} \\ 1.0 - (v_i - v_{\min})/(v_{\max} - v_{\min}), & \text{otherwise,} \end{cases} \tag{27}$$

where $v_{\max}$ and $v_{\min}$ denote the maximum and minimum values of the metric after excluding outliers beyond $\mu \pm 3\sigma$, with $\mu$ and $\sigma$ representing the mean and standard deviation, respectively. The normalized metric values are then averaged to yield a single overall score.

#### 3 Implementation Details for *In Silico* Analysis

##### 3.1 Selection of Targets for Cross-Modality Interaction Analysis

To ensure a distinct spectrum of biocomplexes for deep analysis and case study, we selected entries released recently between *July 1, 2024* (the cutoff date for the training set), and *November 20, 2025*. Subsequently, the filtering protocol on these data was performed following the steps below:

1. We restricted the dataset to complexes containing one single target chain to facilitate subsequent computational analysis, since many conventional tools only accept single-chain target.
2. To mitigate potential data leakage against training set, we dropped entries with target chain shared greater than 40% sequence identity with any chain in the training set. The sequence identities were calculated by MMSeq2[40].
3. For entries derived from BioLiP2, we applied an extra ligand exclusion filter (please refer to Supplementary Table 3 for the full list). An entry was discarded if the ligand was present on this list or exhibited a *Tanimoto similarity* greater than 0.3 with any ligand on the list.
4. Next, we performed interaction verification to exclude complexes stabilized by solely hydrophobic intractions, which might be too biased to single interaction type. Specifically, interactions between the reference ligand and the target protein were calculated, and only complexes exhibiting at least one polar interaction were retained.
5. Finally, we manually inspected the dataset to remove those containing co-factors ligands, as well as redundant items with highly similar or homologous targets.

Finally, eight target-ligand complexes passed all the filters for analysis, including PDB entries: 9CSD, 9DMV, 9G05, 9GMU, 9IT0, 9L4D, 9MEV, and 9MQ7. Specifically, 9GMU, 9MEV, 9MQ7, and 9DMV involve antibody ligands.

##### 3.2 Selection of RNA/DNA targets for *De Novo* Design

The nucleic acid dataset used for de novo design was obtained from XDock[41]. Specifically, the dataset contained 139 targets, which include 17 DNA and 122 RNA structures with diverse molecular size. For each target, we generated 100 samples for both peptides and small molecules and applied XDock[41] to score the generated pose in minimized mode. The PDB ID used for generation can be found at Supplementary Table 15.

##### 3.3 Zero-Shot RNA-Small Molecule Binding Prediction via Likelihoods

For zero-shot RNA-small molecule binding prediction, we obtained the testing set from SMRTnet[42]. A total of 376 compounds were experimentally tested for binding with MYC IRES, a special RNA functional element that is located in the 5' UTR of MYC mRNA. Among these compounds, 15 were identified as positive binders based on Microscale thermophoresis assay[43], a suiltable classifcation task to test the zero-shot binding prediction ability of AnewOmni.

We used Protenix to predict the 3D structure of the target. The predicted structure aligned with the secondary structure provided in the original paper. Then we performed molecular docking by prompting our model with the chemical structure given the smiles of 376 compounds. For each compound, 100 poses were generated, and a maximum timeout was set at 1h. The mean value of likelihood from the generated poses was considered as binding score from AnewOmni.

##### 3.4 Implementation of Generalization Test on Glycans and Phosphorylation Sites

For post-translational modification target, we chose phosphorylated Smad2 protein as demonstrated in BoltzDesign[44]. The serine residues at positions 465 and 467 can be specifically phosphorylated by transforming growth factor-$\beta$ (T$\beta$RI). This process is crucial for Smad2 dissociating from the receptor and binding to Smad4, a key step in TGF-$\beta$ signaling[45][46].

For glycan targets, we focused on the H type-1 antigen, an important type of blood group antigen that serves as one of the precursor structures for the ABO blood system. It also can be exploited as target for certain pathogens as mediators of infections[47][48]. Specifically, we extracted structure of H type-1 antigen from its complex structure with human rotavirus and selected the same epitope for AnewOmni generation.

For each target, we generated 1000 samples for both peptides and small molecules and applied XDock[41] to score the generated pose in minimized mode. The PDB ID used for generation can be found at Supplementary Table 15.

#### 4 Implementation of Molecular Dynamics Simulations

##### 4.1 Complexes between Target Protein and Peptides

All simulations were performed in Amber20 CUDA version of particle-mesh Ewald molecular dynamics (PMEMD) [49] using the AMBER ff14SB force field for proteins and peptides [50]. Protonation states of titratable residues at pH 7.0 were assigned with PropKa via the PDB2PQR pipeline[49]. Each complex was solvated in a TIP3P [51] water box with 10 Å buffer and neutralized with 150 mM NaCl using PACKMOL-Memgen [52]. Following a 10-step minimization and relaxation protocol [53,54], systems were equilibrated in the NPT ensemble at 300 K and 1 bar, regulated by Langevin and Berendsen coupling, respectively. Next, production simulations were then run for 100 ns per complex using a 4.0 fs timestep enabled by hydrogen mass repartitioning [55]. Long-range electrostatic interactions were treated using the Particle Mesh Ewald (PME) method, while the Lennard-Jones potentials were truncated at a cutoff distance of 9.0 Å. Covalent bonds involving hydrogens were constrained with SHAKE [56]. The snapshots were collected every 20 ps, and the final 40 ns of each simulation were subjected to MM/PBSA analysis [57]. The polar solvation component was calculated using an internal dielectric constant of 2.0 and an external constant of 80.0. The nonpolar solvation energy was estimated as a linear function of the SASA[58].

##### 4.2 Complexes between Target Protein and Small Molecules

All-atom molecular dynamics (MD) simulations were performed using the GROMACS 2023 software package[59]. The ligand-protein complexes were prepared by placing the molecules at the center of triclinic boxes with a minimum distance of 1.0 nm to the box boundary. The TIP3P water model[51] was used to solvate the complexes. The system was neutralized and brought to a physiological ionic strength of 0.15 M by adding sodium and chloride ions[60]. The proteins were parameterized with Amber ff14SB[50] and ligands were parameterized with the GAFF2 force field[61]. The relaxation of each system started with energy minimization of 50,000 steps, followed by 400 ps relaxation in the NVT ensemble with a 1fs time step to increase the temperature to 298.15K and 100 ps relaxation in the NPT ensemble at 298.15 K and a pressure of 1 bar. The protein backbone atoms and the ligand heavy atoms were restrained by harmonic position restraints with a force constant of 1000 kJ/mol/nm$^2$ during relaxation. Finally, the systems were further equilibrated without any restraints for 2 ns in the NPT ensemble with a 2 fs time step before collecting 10 ns production data with saving trajectory every 20 ps. The temperature in the simulations was controlled using a Langevin integrator[62] with the friction coefficient set to 0.5 ps$^{-1}$. The stochastic cell rescaling algorithm[63] was applied for pressure coupling with a time constant of 2 ps. All the bonds with hydrogen atoms were constrained using the LINCS algorithm[64]. Periodic boundary conditions were applied, and the particle mesh-Ewald method[65] was used to treat long-range electrostatic interactions with a direct space cutoff of 1.0 nm. The Lennard-Jones interactions were switched off at 0.8 nm and a dispersion correction was applied for energy and pressure. The analysis of MD trajectories was performed using MDAnalysis[66,67] and ProLIF[68].

### 5 Interaction Definitions for Small Molecules Pipeline

#### 5.1 KRAS G12D

Targeting KRAS G12D is challenging because the binding site lacks a deep hydrophobic pocket. Interaction with ASP12 is obviously critical, as this residue distinguishes the G12D mutant from other KRAS variants and is strongly associated with biological activity. ASP69, located at the deepest region of the pocket, acts together with ASP12 to form two spatially distant "anchors" in the pocket. We observed that ligands capable of simultaneously engaging both residues typically exceed the size of conventional small molecules. Thus, during the interaction-filtering stage, ASP12 and ASP69 were defined as key interaction constraints, enabling rapid and efficient reduction of the candidate chemical space (Supplementary Fig. 4d). To further encourage full occupancy of the pocket, we considered the distal residue GLU62. Together, these three residues span the binding pocket in a triangular arrangement. The final interaction score composition used in the molecular dynamics simulations is defined as follows:

$$\text{Interaction Score} = \text{IL}_{\text{ASP12}} + \text{IL}_{\text{GLU62}} + \text{IL}_{\text{ASP69}}, \tag{28}$$

where IL refers to interaction lifetime.

#### 5.2 PCSK9

Our filter pipeline targeted the allosteric site of PCSK9, which was enriched within the Protenix-predicted structures by 1000 different random seeds. We observed a specific hydrogen bond network on the complex of PCSK9 and AZD0780 in the predicted structure. The pyrimidine motif of AZD0780 functions as a hydrogen bond acceptor, with its two nitrogen atoms engaging VAL589 and SER636, respectively. The bridging secondary amide which connecting the pyrimidine and the aliphatic ring acts as a hydrogen bond donor to the backbone carbonyl of VAL589. Thus, The filter pipeline prioritizes interactions with these two key residues on the interaction filter phase and post-MD analysis phase. specifically, a strict filter was applied on the interaction profile for VAL589 during the interaction filter, requiring the ligand to act simultaneously as a hydrogen bond donor and acceptor. The interaction score is defined as follows:

$$\text{Interaction Score} = \text{IL}_{\text{VAL589}} + \text{IL}_{\text{SER636}}, \tag{29}$$

where IL refers to interaction lifetime.

### 6 Metrics in Antibody Design Pipeline

#### 6.1 Implementation Details

We define a set of metrics to rank candidates in the iterative antibody design pipeline that combines AnewOmni with Protenix[69]. These metrics are organized into two groups: (i) Structural plausibility metrics, which enforce alignment with the desired structural objective and are assigned high weights, and (ii) confidence and consistency metrics, which refine the ranking among candidates that satisfy the primary criteria.

**Binding Site Overlap** (weight 10.0) To ensure epitope-specific antibody design, the highest priority is to generate candidates that engage the intended target binding site (epitope). For each designed complex, we define the epitope $\mathbb{P}$ as the set of target residues whose heavy atoms lie within 5 Å of any heavy atom of the antibody. The reference epitope, denoted as $\hat{\mathbb{P}}$, can be specified manually or computed analogously from a reference binder. Then we compute the fraction of residues in the reference epitope that are recovered by the designed candidate:

$$\text{Binding Site Overlap} = \frac{\text{card}(\mathbb{P} \cap \hat{\mathbb{P}})}{\text{card}(\hat{\mathbb{P}})}. \tag{30}$$

**Ratio of Contacts by CDRs** (weight 5.0) A desirable interaction pattern is expected to be dominated by complementarity-determining regions (CDRs), whereas framework residues are typically not involved in target-specific binding, as the frameworks are usually selected by developabilities that are not relevant to binding. We first identify all antibody residues whose heavy atoms lie within 5 Å of any heavy atom of the target. These contacting residues are then partitioned into those belonging to CDRs ($\mathbb{C}_{\text{CDR}}$) and those belonging to the framework ($\mathbb{C}_{\text{FR}}$). The ratio of contacts contributed by CDRs is computed as:

$$\text{Ratio of Contacts by CDRs} = \frac{\text{card}(\mathbb{C}_{\text{CDR}})}{\text{card}(\mathbb{C}_{\text{CDR}} \cup \mathbb{C}_{\text{FR}})}. \tag{31}$$

The following five metrics are used to refine the ranking among candidates that satisfy the primary structural plausibility criteria. These metrics collectively emphasize structural confidence, interface quality, and generative consistency. Their weights sum to 1.0.

**ipTM** (weight 0.3) The interface predicted template modeling score (ipTM) between the target and the antibody as reported by Protenix, reflecting confidence in the predicted interchain interface.

**Antibody pLDDT** (weight 0.2) The predicted local distance difference test (pLDDT) score averaged over all residues of the antibody, measuring the internal structural confidence of the designed antibody.

**ipAE** (weight 0.1) The interface predicted aligned error (ipAE) computed over interchain residue pairs by Protenix. During score aggregation, ipAE is normalized to the range $[0, 1]$, with larger values indicating higher confidence:

$$\text{Normalized ipAE} = 1.0 - \frac{\text{ipAE}}{31.0} \tag{32}$$

**Likelihood Robustness** (weight 0.2) We compute the average generative likelihood of subsequent CDR designs conditioned on a given candidate. This metric reflects whether the docking orientation provides a favorable context for further CDR generation, with higher values indicating more robust and consistent generative behavior.

**scRMSD** (weight 0.2) Self-consistent root mean square deviation (scRMSD) is defined as the RMSD between the $C_\alpha$ atoms of the designed antibody structure and the predicted structure, after alignment on the target protein. For aggregation, scRMSD is normalized as:

$$\text{Normalized scRMSD} = 1.0 - \frac{\text{scRMSD}}{40.0}, \tag{33}$$

where higher values indicate greater consistency between the designed and predicted structures.

#### 6.2 Analysis

For the KRAS G12D case, we performed 60 iterative rounds of antibody design, yielding a total of 3,000 candidates. As binding site overlap and the ratio of contacts contributed by CDRs are assigned the highest weights, the iterative process progressively enriches candidates with higher overall scores, increased epitope coverage, and stronger CDR-dominated interactions among the top-ranked designs (Supplementary Fig. 8d–f). Notably, the desired epitope on the KRAS switch II pocket is not a favorable binding region in the learned distribution of Protenix (Supplementary Fig. 8b), which we attribute to the lack of solved KRAS-antibody complex structures. As a consequence, inter-chain confidence metrics, including ipTM and ipAE, as well as self-consistency, remain relatively low and exhibit substantial variability throughout the iterations (Supplementary Fig. 8g–i). In contrast, the pLDDT of the designed antibodies increases steadily, accompanied by a consistent improvement in generative likelihood (Supplementary Fig. 8j–k), indicating that the generated CDR sequences become increasingly compatible with their respective frameworks. To further characterize the relationships among ranking criteria, we performed a pairwise correlation analysis of all metrics (Supplementary Fig. 8l). As expected, cofolding-derived confidence metrics such as ipTM, ipAE, and pLDDT show medium mutual correlations. Interestingly, binding site overlap exhibits a negative correlation with scRMSD, suggesting that candidates engaging the desired epitope tend to yield more consistent structure predictions across subsequent rounds. These phenomenon are the primary reasons for why we chose scRMSD as an orthogonal validation in the experimental selection strategy while ignoring other confidences.

### 7 Synthesis of Small Molecules

#### 7.1 Designs for KRAS G12D

##### *7.1.1 KRAS G12D-compound-1*

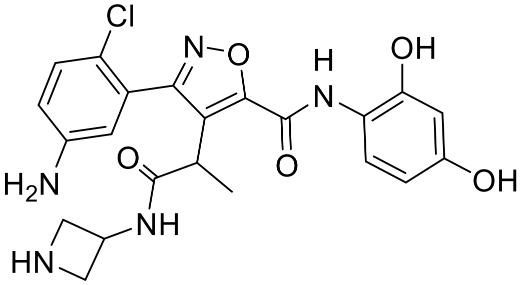

**KRAS G12D-compound-1**. 3-(5-amino-2-chlorophenyl)-4-(1-(azetidin-3-ylamino)-1-oxopropan-2-yl)-N-(2,4-dihydroxyphenyl)isoxazole-5-carboxamide

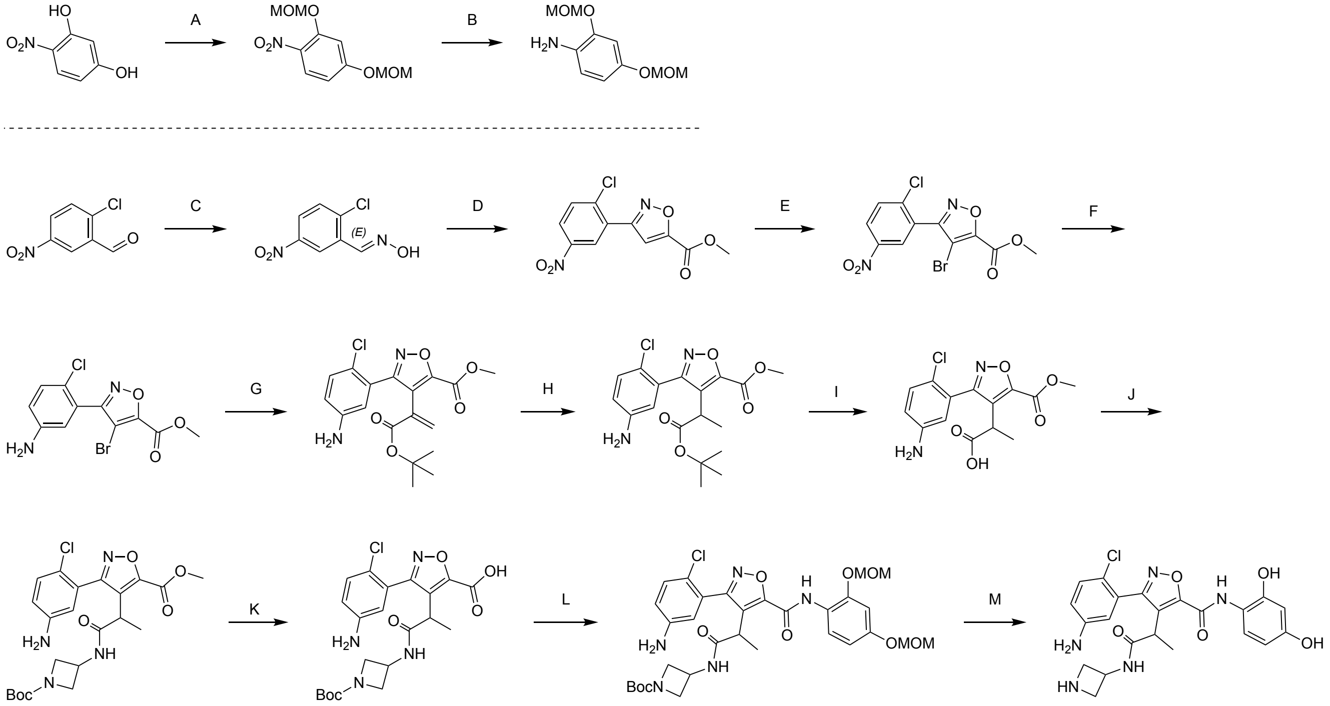

**KRAS G12D-compound-1**

Reagents and conditions: A. bromo(methoxy)methane, DIEA, DMAP, DCM, 0 to 40 °C, 46%; B. 1% Pt/C, MeOH, $H_2$, 1.5 MPa, 60 °C, 86%; C. hydroxylamine hydrochloride, MeOH, 25 °C, 33%; D. methyl propiolate, iodanediyl diacetate, ACN, $H_2O$, 0 to 25 °C, 27%; E. NBS, TFA, 80 °C, 88%; F. Fe, $NH_4Cl$, EtOH, $H_2O$, 80 °C, 69%; G. tert-butyl 2-(tributylstannyl)acrylate, $Pd(PPh_3)_4$, toluene, 110 °C, 32%; H. $PtO_2$, EA, 20 °C, 15 Psi, 48%; I, HCl/dioxane, DCM, 35 °C, 99%; J. HATU, DIEA, DMF, 25 °C, 76%; K. $Na_2CO_3$, THF, MeOH, $H_2O$, 25 °C, 32%; L. TCFH, NMI, ACN, 25 °C, 34%; M. HCl/MeOH, 25 °C, 75%.

**Synthesis of KRAS G12D-compound-1**

Step A: 2,4-bis(methoxymethoxy)-1-nitrobenzene. To a solution of 4-nitrobenzene-1,3-diol (900 mg, 5.80 mmol, 1.0 equiv), N-ethyl-N-isopropylpropan-2-amine (4.50 g, 34.8 mmol, 6.0 equiv) and N,N-dimethylpyridin-4-amine (70.9 mg, 580 $\mu$mol, 0.1 equiv) in dichloromethane (10.0 mL) was added bromo(methoxy)methane (3.63 g, 29.0 mmol, 5.0 equiv) at 0 °C The reaction was stirred at

40 °C for 16 hours. After completion of the reaction, the mixture was cooled to 25 °C. The mixture was purified by column chromatography ($SiO_2$, Petroleum ether: Ethyl acetate = 100/1 to 4/1) to afford 2,4-bis(methoxymethoxy)-1-nitrobenzene (650 mg, 46% yield) as a yellow oil; $^1$H NMR (400 MHz, CHLOROFORM-d) $\delta$ = 7.93 (d, $J$ = 9.2 Hz, 1H), 6.94 (d, $J$ = 2.4 Hz, 1H), 6.74 (dd, $J$ = 2.4, 9.2 Hz, 1H), 5.30 (s, 2H), 5.23 (s, 2H), 3.55 (s, 3H), 3.50 (s, 3H).

Step B: 2,4-bis(methoxymethoxy)aniline. Solution 1: 2,4-bis(methoxymethoxy)-1-nitrobenzene (200 mg, 822 $\mu$mol, 1.0 equiv) in methanol (10.0 mL). The fixed bed (named FLR1, volume 5 mL) was completely packed with granular catalyst {WXSC1017 (1% Pt/C), 1.40 g}. The hydrogen back pressure regulator was adjusted to 1.5 Mpa, and the hydrogen flow rate was {15.0 equiv, 12.1 sccm}. The solution 1 was pumped by Pump 1 {S1, P1, 0.4 mL/min} to flow reactor 1 {FLR1, SS, Fixed bed, 6.350(1/4") mm, 5.00 mL, 60 °C}. The reaction mixture was continuously collected from the reactor outlet into the container. After completion of the reaction, the mixture was concentrated under reduced pressure to afford 2,4-bis(methoxymethoxy)aniline (150 mg, 86% yield) as a brown oil; LCMS-Method 15 $[M+1]^+$ = 214.0.

Step C: (E)-2-chloro-5-nitrobenzaldehyde oxime. To a solution of 2-chloro-5-nitrobenzaldehyde (15.0 g, 80.8 mmol, 1.0 equiv) in methanol (60.0 mL) was added hydroxylamine hydrochloride (5.90 g, 84.9 mmol, 1.05 equiv). The reaction was stirred at 25 °C for 16 hours. After completion of the reaction, the reaction mixture was purified by column chromatography ($SiO_2$, Petroleum ether: Ethyl acetate = 100/1 to 5/1) to afford (E)-2-chloro-5-nitrobenzaldehyde oxime (8.50 g, 33% yield) as a white solid; $^1$H NMR (400 MHz, CHLOROFORM-d) $\delta$ = 8.73 (d, $J$ = 2.8 Hz, 1H), 8.56 (s, 1H), 8.17 (dd, $J$ = 2.8, 8.8 Hz, 1H), 8.06 (br s, 1H), 7.58 (d, $J$ = 8.8 Hz, 1H).

Step D: methyl 3-(2-chloro-5-nitrophenyl)isoxazole-5-carboxylate. To a solution of (E)-2-chloro-5-nitrobenzaldehyde oxime (10.0 g, 49.9 mmol, 1.0 equiv) and methyl propiolate (6.29 g, 74.8mmol, 1.5 equiv) in acetonitrile (400.0 mL) and water (100.0 mL) was added phenyl-l3-iodanediyl diacetate (17.7 g, 54.8 mmol, 1.1 equiv) in acetonitrile (400.0 mL) at 0 °C. The reaction was stirred at 25 °C for 16 hours. After completion of the reaction, the mixture was filtered and the filtrate was concentrated under reduced pressure to give a residue. The residue was purified by column chromatography ($SiO_2$, Petroleum ether: Ethyl acetate = 100/1 to 5/1 to 2/3) to afford methyl 3-(2-chloro-5-nitrophenyl)isoxazole-5-carboxylate (3.80 g, 27% yield) as a white solid; $^1$H NMR (400 MHz, CHLOROFORM-d) $\delta$ = 8.70 (d, $J$ = 2.8 Hz, 1H), 8.30 (dd, $J$ = 2.8, 9.2 Hz, 1H), 7.73 (d, $J$ = 8.8 Hz, 1H), 7.49 (s, 1H), 4.04 (s, 3H); LCMS-Method 15 $[M+1]^+$ = 283.0.

Step E: methyl4-bromo-3-(2-chloro-5-nitrophenyl)isoxazole-5-carboxylate. To a solution of methyl 3-(2-chloro-5-nitrophenyl)isoxazole-5-carboxylate (3.40 g, 12.0 mmol, 1.0 equiv) in 2,2,2-trifluoroacetic acid (30.0 mL) was added 1-bromopyrrolidine-2,5-dione (6.42 g, 36.1 mmol, 3.0 equiv). The reaction was stirred at 80 °C for 36 hours. After completion of the reaction, the mixture was cooled to 25 °C. The mixture was adjusted pH to 7 with sodium hydrogen carbonate aqueous solution, extracted with ethyl acetate (3 × 100 mL). The combined organic layers were washed with brine (100 mL), dried over anhydrous sodium sulfate, filtered and concentrated under reduced pressure to give a residue. The residue was purified by column chromatography ($SiO_2$, Petroleum ether: Ethyl acetate = 100/1 to 5/1) and then triturated with Petroleum ether: Ethyl acetate = 5:1 (2 × 10 ml) to afford methyl 4-bromo-3-(2-chloro-5-nitrophenyl)isoxazole-5-carboxylate (4.00 g, 88% yield) as a yellow solid; $^1$H NMR (400 MHz, CHLOROFORM-d) $\delta$ = 8.40 - 8.36 (m, 1H), 8.36 - 8.34 (m, 1H), 7.77 (d, $J$ = 8.8 Hz, 1H), 4.07 (s, 3H).

Step F: methyl3-(5-amino-2-chlorophenyl)-4-bromoisoxazole-5-carboxylate. To a solution of methyl 4-bromo-3-(2-chloro-5-nitrophenyl)isoxazole-5-carboxylate (2.80 g, 7.74 mmol, 1.0 equiv) and ammonia hydrochloride (2.07 g, 38.7 mmol, 5.0 equiv) in ethanol (50.0 mL) and water (10.0

mL) was added iron (2.16 g, 38.7 mmol, 5.0 equiv). The reaction was stirred at 80 °C for 2 hours. After completion of the reaction, the mixture was cooled to 25 °C. The mixture was diluted with water (200 mL) and extracted with ethyl acetate (3 × 100 mL). The combined organic layers were washed with brine (100 mL), dried over anhydrous sodium sulfate, filtered and concentrated under reduced pressure to give a residue. The residue was purified by column chromatography ($SiO_2$, Petroleum ether: Ethyl acetate = 100/1 to 2/1) to afford methyl 3-(5-amino-2-chlorophenyl)-4-bromoisoxazole-5-carboxylate (2.00 g, 69% yield) as a yellow oil; LCMS-Method 15 $[M+1]^+$ = 330.9.

Step G: methyl3-(5-amino-2-chlorophenyl)-4-(3-(tert-butoxy)-3-oxoprop-1-en-2-yl)isoxazole-5-carboxylate. A mixture of methyl 3-(5-amino-2-chlorophenyl)-4-bromoisoxazole-5-carboxylate (540 mg, 1.63 mmol, 1.0 equiv), tert-butyl 2-(tributylstannyl)acrylate (1.02 g, 2.44 mmol, 1.5 equiv), tetrakis(triphenylphosphine) palladium(0) (188 mg, 163 $\mu$mol, 0.1 equiv) in toluene (10.0 mL) was degassed and purged with nitrogen for 3 times, and then the reaction was stirred at 110 °C for 20 hours under nitrogen atmosphere. After completion of the reaction, the mixture was cooled to 25 °C. The mixture was quenched by addition potassium fluoride solvent (20 mL), extracted with ethyl acetate (3 × 20 mL). The combined organic layers were washed with brine (20 mL), dried over anhydrous sodium sulfate, filtered and concentrated under reduced pressure to give a residue. The residue was purified by column chromatography ($SiO_2$, Petroleum ether: Ethyl acetate = 100/1 to 2/1) and reversed-phase flash (Sfar C18 40 g D Duo 30 $\mu$m; mobile phase: [water (FA)-ACN]; gradient: 33% - 57% B over 11 min) to afford methyl 3-(5-amino-2-chlorophenyl)-4-(3-(tert-butoxy)-3-oxoprop-1-en-2-yl)isoxazole-5-carboxylate (210 mg, 32% yield) as a yellow solid; LCMS-Method 15 $[M+1]^+$ = 379.1.

Step H: methyl3-(5-amino-2-chlorophenyl)-4-(1-(tert-butoxy)-1-oxopropan-2-yl)isoxazole-5-carboxylate. To a solution of methyl 3-(5-amino-2-chlorophenyl)-4-(3-(tert-butoxy)-3-oxoprop-1-en-2-yl)isoxazole-5-carboxylate (400 mg, 1.06 mmol, 1.0 equiv) in ethyl acetate (20.0 mL) was added platinum(IV) oxide (24.0 mg, 106 $\mu$mol, 0.1 equiv) nitrogen atmosphere. The suspension was degassed and purged with hydrogen for 3 times. The reaction was stirred under hydrogen (15.0 Psi) at 20 °C for 6 hours. After completion of the reaction, the mixture was filtered and the filtrate was purified by column chromatography ($SiO_2$, Petroleum ether: Ethyl acetate = 100/1 to 2/1) to afford methyl 3-(5-amino-2-chlorophenyl)-4-(1-(tert-butoxy)-1-oxopropan-2-yl)isoxazole-5-carboxylate (200 mg, 48% yield) as a white solid; LCMS-Method 15 $[M+1]^+$ = 381.2.

Step I: 2-(3-(5-amino-2-chlorophenyl)-5-(methoxycarbonyl)isoxazol-4-yl)propanoicacid. To a solution of methyl 3-(5-amino-2-chlorophenyl)-4-(1-(tert-butoxy)-1-oxopropan-2-yl)isoxazole-5-carboxylate (190 mg, 499 $\mu$mol, 1.0 equiv) in dichloromethane (1.0 mL) was added hydrogen chloride/dioxane (10.0 mL, 2.0 M, 40.1 equiv). The reaction was stirred at 35 °C for 4 hours. After completion of the reaction, the mixture was cooled to 25 °C. The mixture was concentrated under reduced pressure to afford 2-(3-(5-amino-2-chlorophenyl)-5-(methoxycarbonyl)isoxazol-4-yl)propanoic acid (160 mg, 99% yield) as a white solid; LCMS-Method 1 $[M+1]^+$ = 325.1.

Step J: methyl3-(5-amino-2-chlorophenyl)-4-(1-((1-(tert-butoxycarbonyl)azetidin-3-yl)amino)-1-oxopropan-2-yl)isoxazole-5-carboxylate. To a solution of 2-(3-(5-amino-2-chlorophenyl)-5-(methoxycarbonyl)isoxazol-4-yl)propanoic acid (160 mg, 493 $\mu$mol, 1.0 equiv), tert-butyl 3-aminoazetidine-1-carboxylate (93.4 mg, 542 $\mu$mol, 1.1 equiv), N-ethyl-N-isopropylpropan-2-amine (191 mg, 1.48 mmol, 3.0 equiv) in N,N-dimethylformamide (2.0 mL) was added 2-(3H-[1,2,3]triazolo[4,5-b]pyridin-3-yl)-1,1,3,3-tetramethyluroniumhexafluorophosphate(V) (187 mg, 493 $\mu$mol, 1.0 equiv). The reaction was stirred at 25 °C for 1 hour. After completion of the reaction, the mixture was purified by reversed-phase flash (Sfar C18 40 g D Duo 30 $\mu$m; mobile phase: [water

(FA)-ACN]; gradient: 37%-55% B over 11 mins) to afford methyl 3-(5-amino-2-chlorophenyl)-4-(1-((1-(tert-butoxycarbonyl)azetidin-3-yl)amino)-1-oxopropan-2-yl)isoxazole-5-carboxylate (200 mg, 76% yield) as a white solid; LCMS-Method 1 $[M+1]^+ = 479.3$.

Step K: 3-(5-amino-2-chlorophenyl)-4-(1-((1-(tert-butoxycarbonyl)azetidin-3-yl)amino)-1-oxopropan-2-yl)isoxazole-5-carboxylicacid. To a solution of methyl 3-(5-amino-2-chlorophenyl)-4-(1-((1-(tert-butoxycarbonyl)azetidin-3-yl)amino)-1-oxopropan-2-yl)isoxazole-5-carboxylate (100 mg, 209 μmol, 1.0 equiv) in tetrahydrofuran (1.0 mL) and methanol (1.0 mL) was added sodium carbonate (22.1 mg, 209 μmol, 1.0 equiv) in water (0.5 mL). The reaction was stirred at 25 °C for 1 hour. After completion of the reaction, the mixture was adjusted pH to 2 with hydrogen chloride (1.0 M). The mixture was purified by reversed-phase flash (Sfar C18 40 g D Duo 30 μm; mobile phase: [water (HCl)-ACN]; gradient: 38%-60% B over 11 min) to afford 3-(5-amino-2-chlorophenyl)-4-(1-((1-(tert-butoxycarbonyl)azetidin-3-yl)amino)-1-oxopropan-2-yl)isoxazole-5-carboxylic acid (31.0 mg, 32% yield) as a white solid; LCMS-Method 1 $[M+1]^+ = 465.1$.

Step L: tert-butyl3-(2-(3-(5-amino-2-chlorophenyl)-5-((2,4-bis(methoxymethoxy)phenyl)carbamoyl)isoxazol-4-yl)propanamido)azetidine-1-carboxylate. To a solution of 3-(5-amino-2-chlorophenyl)-4-(1-((1-(tert-butoxycarbonyl)azetidin-3-yl)amino)-1-oxopropan-2-yl)isoxazole-5-carboxylic acid (25.0 mg, 53.8 μmol, 1.0 equiv), 2,4-bis(methoxymethoxy)aniline (34.4 mg, 161 μmol, 3.0 equiv) and 1-methyl-1H-imidazole (9.27 mg, 112 μmol, 2.1 equiv) in acetonitrile (1.0 mL) as added N-(chloro(dimethylamino)methylene)-N-methylmethanaminium hexafluorophosphate(V) (18.1 mg, 64.5 μmol, 1.2 equiv). The reaction was stirred at 25 °C for 1 hour. After completion of the reaction, the mixture was purified by prep-HPLC (Sfar C18 12 g D Duo 30 μm; mobile phase: [water(FA)-ACN]; gradient: 33%-57% B over 15 min) to afford tert-butyl3-(2-(3-(5-amino-2-chlorophenyl)-5-((2,4-bis(methoxymethoxy)phenyl)carbamoyl)isoxazol-4-yl)propanamido)azetidine-1-carboxylate (13.0 mg, 34% yield) as a white solid; LCMS-Method 1 $[M+1]^+ = 660.3$.

Step M: KRAS G12D-compound-1. To a solution of tert-butyl3-(2-(3-(5-amino-2-chlorophenyl)-5-((2,4-bis(methoxymethoxy)phenyl)carbamoyl)isoxazol-4-yl)propanamido)azetidine-1-carboxylate (13.0 mg, 19.7 μmol, 1.0 equiv) in methanol (2.0 mL) was added hydrogen chloride/methanol (3.70 mL, 2 M, 377 equiv). The reaction was stirred at 25 °C for 1 hour. After completion of the reaction, the mixture was purified by prep-HPLC (Sfar C18 12 g D Duo 30 μm; mobile phase: [water(FA)-ACN]; gradient: 15%-30% B over 11 min) to afford 3-(5-amino-2-chlorophenyl)-4-(1-(azetidin-3-ylamino)-1-oxopropan-2-yl)-N-(2,4-dihydroxyphenyl)isoxazole-5-carboxamide (8.04 mg, 75% yield) as a white solid; $^1$H NMR (400 MHz, METHANOL-d4) $\delta$ = 7.60 (d, $J$ = 8.8 Hz, 1H), 7.23 (d, $J$ = 8.8 Hz, 1H), 6.82 (dd, $J$ = 2.8, 8.8 Hz, 1H), 6.70 (d, $J$ = 2.8 Hz, 1H), 6.42 (d, $J$ = 2.8 Hz, 1H), 6.33 (dd, $J$ = 2.8, 8.8 Hz, 1H), 4.53 (quin, $J$ = 7.2 Hz, 1H), 4.18 - 4.01 (m, 4H), 3.92 (q, $J$ = 7.2 Hz, 1H), 1.45 (d, $J$ = 7.6 Hz, 3H); LCMS $[M+1]^+ = 472.2$; HPLC Rt = 0.895 min;

**Analytical method by SFC:**

Column: Chiralcel OD-3R 100 × 4.6 mm I.D., 3 μm;

Mobile phase: Phase A for Water with 0.04% TFA, and Phase B for ACN with 0.02% TFA;

Gradient elution: 10% B in A;

Flow rate: 1 mL/min; Detector: PDA;

Column Temp: 35 °C;

Retention time A: 5.771 min;

Retention time B: 6.895 min.

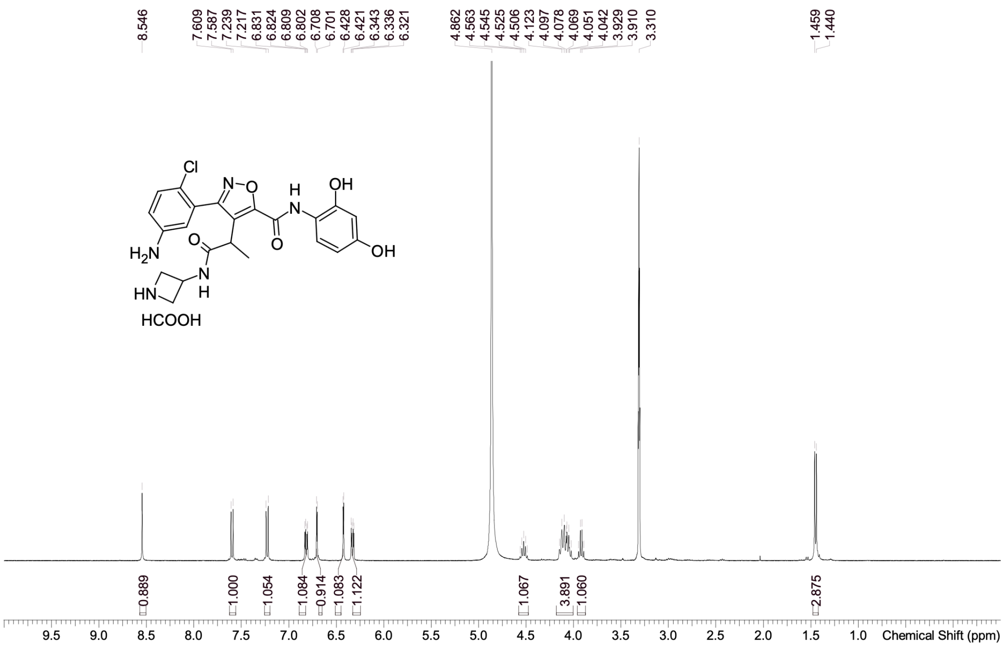

| | |
|---|---|
| Acquisition Time (sec) | 1.9999 |
| Comment | EW53498-1150-P1N MeOD BRUKER_03_017_400MHz |
| Date | 01 Dec 2025 10:50:16 (GMT+08:00) |
| Frequency (MHz) | 400.1500 |
| Nucleus | 1H |
| Number of Transients | 1 |
| Origin | Avance |
| Original Points Count | 16393 |
| Owner | nmrsu |
| Points Count | 65536 |
| Pulse Sequence | zg |
| Receiver Gain | 18.00 |
| SW(cyclical) (Hz) | 8196.72 |
| Solvent | METHANOL-d4 |
| Spectrum Offset (Hz) | 2463.4827 |
| Spectrum Type | standard |
| Sweep Width (Hz) | 8196.60 |
| Temperature (degree C) | 25.706 |

Confidential. For research information only

$^{1}$H NMR (400 MHz, METHANOL-d4) δ = 7.60 (d, *J* = 8.8 Hz, 1H), 7.23 (d, *J* = 8.8 Hz, 1H), 6.82 (dd, *J* = 2.8, 8.8 Hz, 1H), 6.70 (d, *J* = 2.8 Hz, 1H), 6.42 (d, *J* = 2.8 Hz, 1H), 6.33 (dd, *J* = 2.8, 8.8 Hz, 1H), 4.53 (quin, *J* = 7.2 Hz, 1H), 4.18 - 4.01 (m, 4H), 3.92 (q, *J* = 7.2 Hz, 1H), 1.45 (d, *J* = 7.6 Hz, 3H)

#### NMR Spectrum of KRAS G12D-compound-1

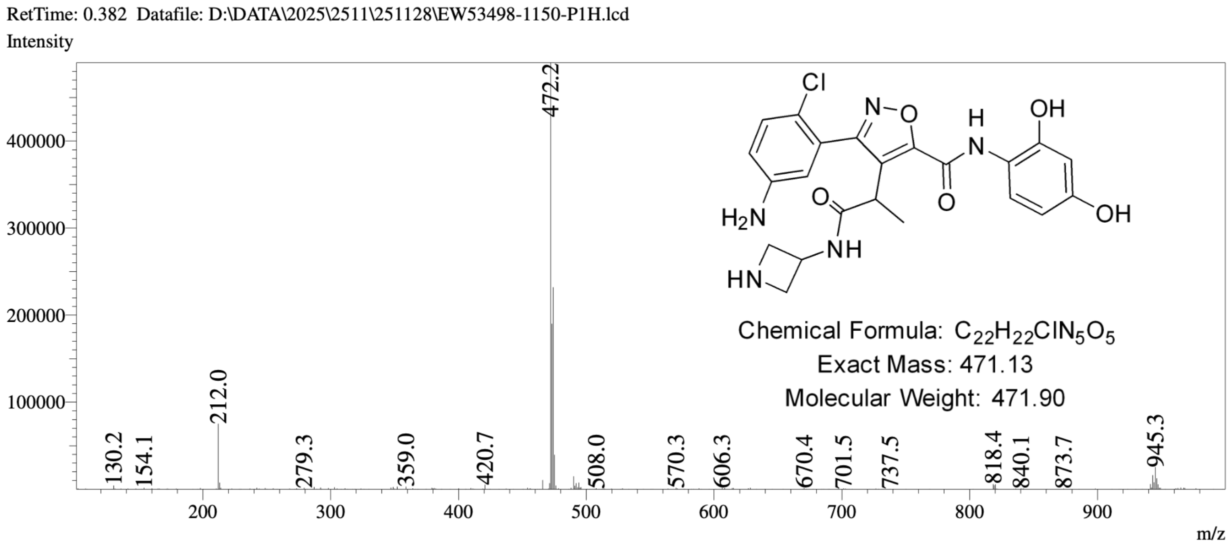

#### MS Spectrum of KRAS G12D-compound-1

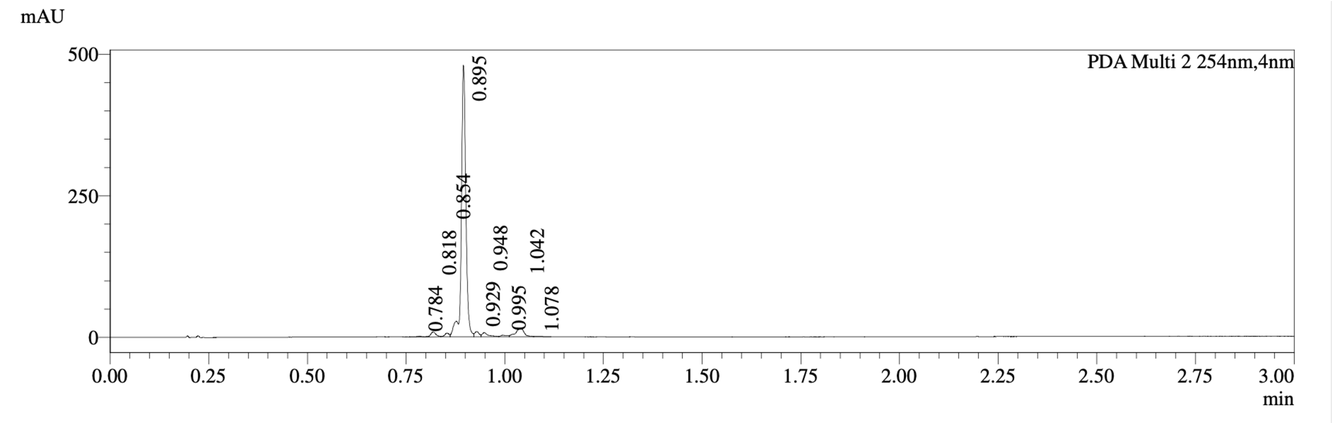

| PDA Ch2 254nm Peak# | Ret. Time | Width | Height | Height% | Area | Area% |
|---|---|---|---|---|---|---|
| 1 | 0.784 | 0.022 | 1160 | 0.221 | 1183 | 0.271 |
| 2 | 0.818 | 0.023 | 8736 | 1.666 | 8392 | 1.920 |
| 3 | 0.854 | 0.025 | 6501 | 1.240 | 5314 | 1.216 |
| 4 | 0.895 | 0.019 | 473393 | 90.296 | 379257 | 86.794 |
| 5 | 0.929 | 0.026 | 9423 | 1.797 | 7398 | 1.693 |
| 6 | 0.948 | 0.025 | 7448 | 1.421 | 8354 | 1.912 |
| 7 | 0.995 | 0.072 | 2783 | 0.531 | 3511 | 0.804 |
| 8 | 1.042 | 0.033 | 13976 | 2.666 | 22518 | 5.153 |
| 9 | 1.078 | 0.078 | 847 | 0.161 | 1037 | 0.237 |

**HPLC Chromatogram of KRAS G12D-compound-1**

##### *7.1.2 KRAS G12D-compound-2*

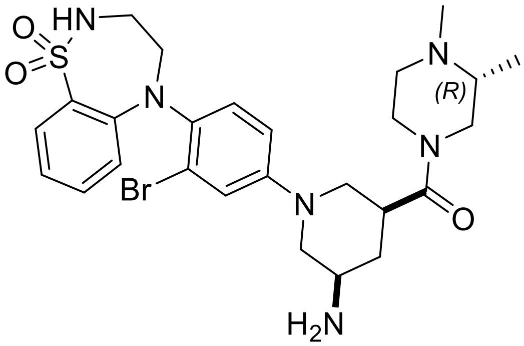

**KRAS G12D-compound-2**. cis-(5-amino-1-(3-bromo-4-(1,1-dioxido-3,4-dihydrobenzo[f][1,2,5]thiadiazepin-5(2H)-yl)phenyl)piperidin-3-yl)((R)-3,4-dimethylpiperazin-1-yl)methanone

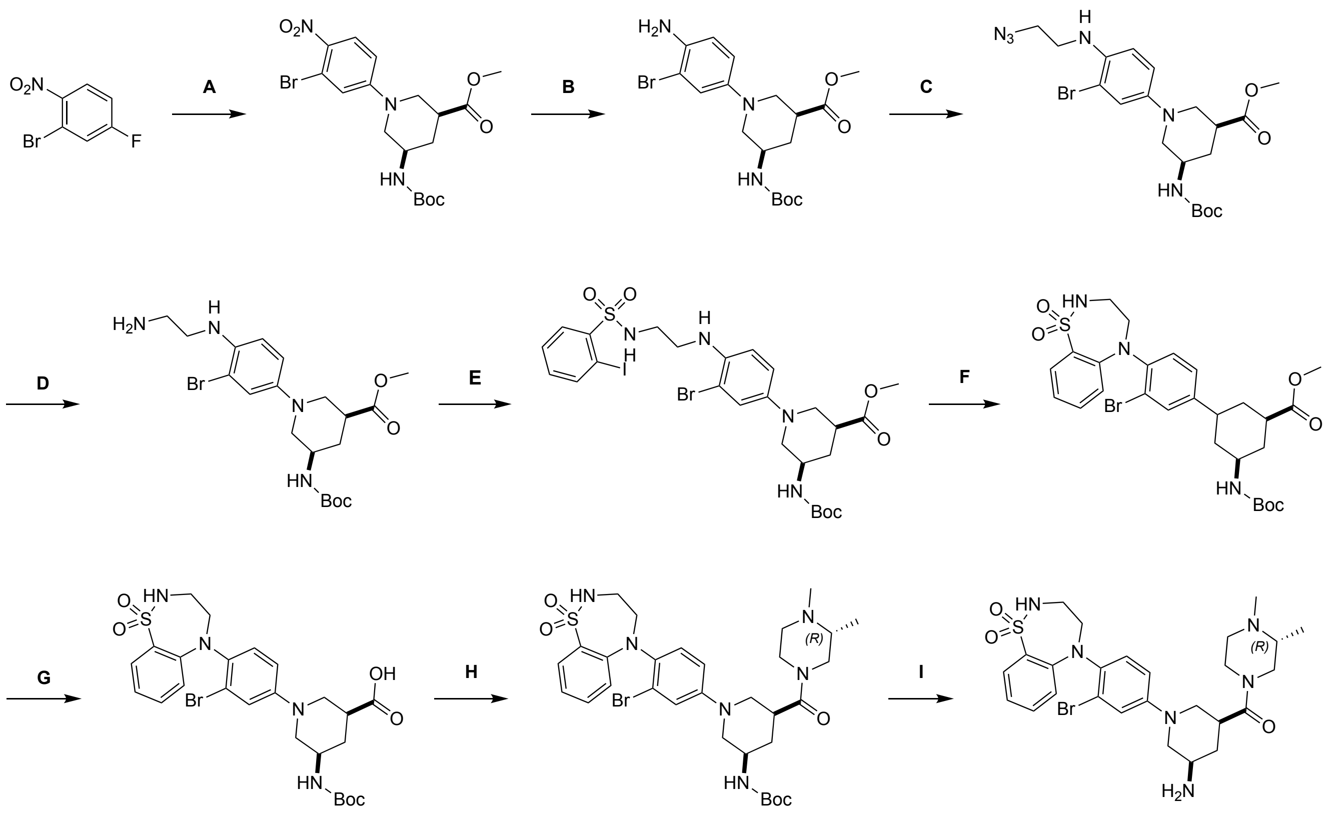

KRAS G12D-compound-2

Reagents and conditions: A. NMP, DIEA, 90 °C, 38%; B, Fe, $NH_4Cl$, EtOH, $H_2O$, 80 °C, 93%; C. $Cs_2CO_3$, KI, Acetone, $H_2O$, 90 °C, 72%; D. $PPh_3$, THF, $H_2O$, 20 °C, 93%; E. 2-iodobenzenesulfonyl chloride, DIEA, DCM, 0 °C, 91%; F. Cu, CuI, $K_2CO_3$, DMF, 100 °C, 74%; G. LiOH, THF, MeOH, 20 °C, 96%; H. EDCI, Pridine, 20 °C, 73%; I. HCl/dioxane, MeOH, 20 °C, 77%.

#### Synthesis of KRAS G12D-compound-2

Step A: cis-methyl1-(3-bromo-4-nitrophenyl)-5-((tert-butoxycarbonyl)amino)piperidine-3-carboxylate. To a solution of 2-bromo-4-fluoro-1-nitrobenzene (750 mg, 3.41 mmol, 1.0 equiv) and methyl 5-((tert-butoxycarbonyl)amino)piperidine-3-carboxylate (881 mg, 3.41 mmol, 1.0 equiv) in 1-methylpyrrolidin-2-one (10.0 mL) was added N-ethyl-N-isopropylpropan-2-amine (881 mg, 6.82 mmol, 2.0 equiv). The reaction mixture was stirred at 90 °C for 2 hours. After completion of the reaction, the reaction mixture was cool to room temperature. The mixture was diluted with water (70 mL), extracted with ethyl acetate (40 mL × 2). The combined organic layers were washed with brine (50 mL × 2), dried over anhydrous sodium sulfate, filtered and concentrated under reduced pressure to give a residue. The residue was purified by prep-HPLC (column: Waters X bridge Prep OBD C18 150*40mm*10$\mu$m; mobile phase: [$H_2O$(10mM $NH_4HCO_3$)-ACN]; gradient : 45%-75% B over 18.0 minutes) to afford cis-methyl 1-(3-bromo-4-nitrophenyl)-5-((tert-butoxycarbonyl)amino)piperidine-3-carboxylate (630 mg, 38% yield) as yellow solid; $^1$H NMR (400 MHz, CHLOROFORM-d) $\delta = 8.01$ (t, $J = 3.2$ Hz, 1H), 7.14 (s, 1H), 6.89 -6.87 (m, 1H), 4.73 (s, 1H), 3.99 - 3.93 (m, 2H), 3.75 (s, 3H), 3.74 - 3.6 (m, 1H), 3.31 - 3.26 (m, 1H), 2.91 (t, $J = 4.0$ Hz, 1H), 2.73 (s, 1H), 2.34 (t, $J = 2.8$ Hz, 1H), 1.66 (t, $J = 3.6$ Hz, 1H), 1.47 (s, 9H); LCMS-Method 27 $[M+1]^+ = 458.0$.

Step B: cis-methyl1-(4-amino-3-bromophenyl)-5-((tert-butoxycarbonyl)amino)piperidine-3-carboxylate: To a solution of cis-methyl 1-(3-bromo-4-nitrophenyl)-5-((tert-butoxycarbonyl)a mino)piperidine-3-carboxylate (580 mg, 1.27 mmol, 1.0 equiv) in ethanol (6.0 mL) were added iron powder (353 mg, 6.33 mmol, 5.0 equiv) and ammonium chloride (677 mg, 12.7 mmol, 10.0 equiv) in water (1.5 mL). The mixture was stirred at 80 °C for 3 hours. After completion of the reaction, the reaction mixture was cool to room temperature. The reaction mixture was diluted with water (50 mL) and extracted with dichloromethane (40 mL × 2). The combined organic layers were washed with brine (70 mL × 2), dried over anhydrous sodium sulfate, filtered and concentrated under reduced pressure to give a residue. The residue was purified by column chromatography ($SiO_2$, Commercial hexanes: Ethyl acetate = 10/1 to 1/1) to afford cis-methyl 1-(4-amino-3-bromophenyl)-5-((tert-butoxycarbonyl)amino)piperidine-3-carboxylate (532 mg, 93% yield) as yellow oil; LCMS-Method 1 $[M+1]^+$ = 428.2.

Step C: cis-methyl1-(4-((2-azidoethyl)amino)-3-bromophenyl)-5-((tert-butoxycarbonyl)amino)pi peridine-3-carboxylate: A mixture of cis-methyl 1-(4-amino-3-bromophenyl)-5-((tert-butoxycarbon yl)amino)piperidine-3-carboxylate (300 mg, 700 $\mu$mol, 1.0 equiv), calcium carbonate (210 mg, 2.10 mmol, 3.0 equiv) and potassium iodide (233 mg, 1.40 mmol, 2.0 equiv) in acetone (3.0 mL) and water (3.0 mL) was added 2-azidoethyl 4-methylbenzenesulfonate (338 mg, 1.40 mmol, 2.0 equiv). The reaction was stirred at 90 °C for 12 hours. After completion of the reaction, the reaction mixture was cool to room temperature. The mixture was filtered and the filtrate was concentrated under reduced pressure to give a residue. The residue was purified by reversed-phase flash (column: Sfar C18 20 g D Duo 30um; mobile phase: [water(FA)-ACN]; gradient: 30%-80% B over 15 minutes) to afford cis-methyl 1-(4-((2-azidoethyl)amino)-3-bromophenyl)-5-((tert-butoxycarbonyl)amino)piperidine-3-carboxylate (250 mg, 72% yield) as yellow solid; LCMS-Method 1 $[M+1, M+3]^+$ = 497.2, 499.2.

Step D: cis-methyl1-(4-((2-aminoethyl)amino)-3-bromophenyl)-5-((tert-butoxycarbonyl)ami no)piperidine-3-carboxylate: To a solution of cis-methyl1-(4-((2-azidoethyl)amino)-3-bromophe nyl)-5-((tert-butoxycarbonyl)amino)piperidine-3-carboxylate (250 mg, 502 $\mu$mol, 1.0 equiv) and triphenylphosphane (264 mg, 1.01 mmol, 2.0 equiv) in tetrahydrofuran (5.0 mL) and water (1.0 mL). The reaction was stirred at 20 °C for 2 hours. After completion of the reaction, the mixture was diluted with water (10 mL) and extracted with dichloromethane (3 × 10 mL). The combined organic layers were washed with brine (20 mL), dried over anhydrous sodium sulfate, filtered and concentrated under reduced pressure to give a residue. The residue was purified by reversed-phase flash (column: Sfar C18 20 g D Duo 30$\mu$m; mobile phase: [water(FA)-ACN]; gradient: 20%-60% B over 10 minutes) to afford cis-methyl1-(4-((2-aminoethyl)amino)-3-bromophenyl)-5-((tert-butoxyca rbonyl)amino)piperidine-3-carboxylate (220 mg, 93% yield) as yellow solid; $^1$H NMR (400 MHz, CHLOROFORM-d) $\delta$ = 7.12 (d, $J$ = 2.8 Hz, 1H), 6.89 (dd, $J$ = 2.4, 8.8 Hz, 1H), 6.62 (d, $J$ = 8.8 Hz, 1H), 4.78 - 4.65 (m, 1H), 3.88 - 3.76 (m, 1H), 3.72 (s, 3H), 3.55 - 3.37 (m, 2H), 3.23 (t, $J$ = 5.6 Hz, 2H), 3.03 - 2.94 (m, 2H), 2.90 - 2.81 (m, 1H), 2.81 - 2.72 (m, 1H), 2.51 - 2.41 (m, 1H), 2.30 - 2.19 (m, 1H), 1.62 - 1.50 (m, 2H), 1.46 (s, 9H).

Step E: cis-methyl1-(3-bromo-4-((2-((2-iodophenyl)sulfonamido)ethyl)amino)phenyl)-5-((tert-bu toxycarbonyl)amino)piperidine-3-carboxylate: To a solution of cis-methyl1-(4-((2-aminoethyl)amin o)-3-bromophenyl)-5-((tert-butoxycarbonyl)amino)piperidine-3-carboxylate (210 mg, 445 $\mu$mol, 1.0 equiv) and N-ethyl-N-isopropylpropan-2-amine (173 mg, 1.34 mmol, 3.0 equiv) in dichloromethane (2.0 mL) was added 2-iodobenzenesulfonyl chloride (135 mg, 445 $\mu$mol, 1.0 equiv) at 0 °C. The reaction was stirred at 0 °C for 30 minutes. After completion of the reaction, the reaction mixture was diluted with water (10 mL) and extracted with dichloromethane (3 × 10 mL). The combined organic layers were washed with brine (20 mL), dried over anhydrous sodium sulfate, filtered and concentrated under reduced pressure to give a residue. The residue was purified by reversed-phase flash (column: Sfar C18 40 g D Duo 30um; mobile phase: [water(FA)-ACN]; gradient: 30%-80% B over 15 minutes) to afford cis-methyl1-(3-bromo-4-((2-((2-iodophenyl)sulfonamido)ethyl)amino)phenyl)-5-((tert-butoxycarbonyl)amino)piperidine-3-carboxylate (300 mg, 91% yield) as yellow solid; $^1$H NMR (400 MHz, DMSO-d6) $\delta$ = 8.12 (dd, $J$ = 1.2, 7.6 Hz, 1H), 7.97 (dd, $J$ = 1.6, 8.0 Hz, 1H), 7.96 - 7.91 (m, 1H), 7.60 - 7.49 (m, 1H), 7.28 (dt, $J$ = 1.6, 7.6 Hz, 1H), 7.03 (d, $J$ = 2.4 Hz, 1H), 6.94 (br d, $J$ = 7.6 Hz, 1H), 6.83 (dd, $J$ = 2.4, 8.8 Hz, 1H), 6.51 (d, $J$ = 9.2 Hz, 1H), 4.85 (t, $J$ = 6.0 Hz, 1H), 3.62 (s, 3H), 3.59 - 3.53 (m, 1H), 3.52 - 3.37 (m, 4H), 3.15 (q, J = 6.0 Hz, 2H), 3.00 (q, $J$ = 6.0 Hz, 2H), 2.75 - 2.66 (m, 1H), 2.32 - 2.22 (m, 1H), 2.12 - 2.00 (m, 1H), 1.39 (s, 9H); LCMS-Method 1 $[M+1]^+$ = 737.1.

Step F: cis-methyl3-(3-bromo-4-(1,1-dioxido-3,4-dihydrobenzo[f][1,2,5]thiadiazepin-5(2H)-yl)phenyl)-5-((tert-butoxycarbonyl)amino)cyclohexane-1-carboxylate: To a solution of cis-methyl1-(3-bromo-4-((2-((2-iodophenyl)sulfonamido)ethyl)amino)phenyl)-5-((tert-butoxycarbonyl)amino)piperidine-3-carboxylate (260 mg, 353 $\mu$mol, 1.0 equiv) in N,N-dimethyl formamide (2.0 mL) were added copper (11.2 mg, 176 $\mu$mol, 0.50 equiv), copper(l) iodide (67.2 mg, 353 $\mu$mol, 1.0 equiv) and potassium carbonate (97.5 mg, 705 $\mu$mol, 2.0 equiv). The reaction was stirred at 100 °C for 1 hour under nitrogen. After completion of the reaction, the reaction mixture was cool to room temperature. The reaction mixture was filtered and the filtrate was concentrated under reduced pressure to give a residue. The residue was purified by reversed-phase flash (column: Sfar C18 20 g D Duo 30$\mu$m; mobile phase: [water($NH_3 \cdot H_2O$)-ACN]; gradient: 20%-80% B over 20 minutes) to afford cis-methyl3-(3-bromo-4-(1,1-dioxido-3,4-dihydrobenzo[f][1,2,5]thiadiazepin-5(2H)-yl)phenyl)-5-((tert-butoxycarbonyl)amino)cyclohexane-1-carboxylate (160 mg, 74 % yield) as yellow solid; LCMS-Method 4 $[M+1, M+3]^+$ = 609.0, 611.0.

Step G: cis-1-(3-bromo-4-(1,1-dioxido-3,4-dihydrobenzo[f][1,2,5]thiadiazepin-5(2H)-yl)phenyl)-5-((tert-butoxycarbonyl)amino)piperidine-3-carboxylicacid: To a solution of cis-methyl3-(3-bromo-4-(1,1-dioxido-3,4-dihydrobenzo[f][1,2,5]thiadiazepin-5(2H)-yl)phenyl)-5-((tert-butoxycarbonyl)amino)cyclohexane-1-carboxylate (150 mg, 246 $\mu$mol, 1.0 equiv) in tetrahydrofuran (2.0 mL) and methanol (1.0 mL) was added lithium hydroxide hydrate (1 M, 492 $\mu$L, 2.0 equiv). The reaction was stirred at 20 °C for 0.5 hour. After completion of the reaction, the mixture was adjusted pH to 6 with HCl (1N). Then the mixture was diluted with water (10 mL) and extracted with ethyl acetate (20 mL × 3). The combined organic layers were washed with brine (10 mL), dried over anhydrous sodium sulfate, filtered and concentrated under reduced pressure to afford cis-1-(3-bromo-4-(1,1-dioxido-3,4-dihydrobenzo[f][1,2,5]thiadiazepin-5(2H)-yl)phenyl)-5-((tert-butoxycarbonyl)amino)piperidine-3-carboxylicacid (140 mg, 96% yield) as yellow solid; LCMS-Method 1 $[M+1]^+$ = 595.1.

Step H: cis-tert-butyl(1-(3-bromo-4-(1,1-dioxido-3,4-dihydrobenzo[f][1,2,5]thiadiazepin-5(2H)-yl)phenyl)-5-((R)-3,4-dimethylpiperazine-1-carbonyl)piperidin-3-yl)carbamate: To a solution of cis-1-(3-bromo-4-(1,1-dioxido-3,4-dihydrobenzo[f][1,2,5]thiadiazepin-5(2H)-yl)phenyl)-5-((tert-butoxycarbonyl)amino)piperidine-3-carboxylicacid (130 mg, 218 $\mu$mol, 1.0 equiv) and (R)-1,2-dimethylpiperazine (49.3 mg, 327 $\mu$mol, 1.50 equiv, HCl salt) in pyridine (2.0 mL) was added 3-(((ethylimino)methylene)amino)-N,N-dimethylpropan-1-amine hydrochloride (62.8 mg, 327 $\mu$mol, 1.50 equiv). The reaction was stirred at 20 °C for 0.5 hour. After completion of the reaction, the mixture was concentrated under reduced pressure to give a residue. The residue was purified by reversed-phase flash (column: Sfar C18 40 g D Duo 30um; mobile phase: [water($NH_3 \cdot H_2O$)-ACN]; gradient: 20%-80% B over 10 minutes) to afford cis-tert-butyl(1-(3-bromo-4-(1,1-dioxido-3,4-dihydrobenzo[f][1,2,5]thiadiazepin-5(2H)-yl)phenyl)-5-((R)-3,4-dimethylpiperazine-1-carbonyl)piperidin-3-yl)carbamate (110 mg, 73% yield) as yellow solid; $^1$H NMR (400 MHz, DMSO-d6) $\delta$ = 8.01 - 7.77

(m, 1H), 7.71 (d, $J$ = 8.0 Hz, 1H), 7.35 - 7.29 (m, 1H), 7.28 - 7.20 (m, 2H), 7.08 - 6.91 (m, 3H), 6.45 (d, $J$ = 8.4 Hz, 1H), 4.22 - 4.01 (m, 1H), 3.94 - 3.68 (m, 4H), 3.62 - 3.48 (m, 1H), 3.24 - 3.10 (m, 3H), 3.06 - 2.78 (m, 3H), 2.77 - 2.65 (m, 2H), 2.47 - 2.30 (m, 1H), 2.22 - 2.13 (m, 3H), 2.05 - 1.77 (m, 3H), 1.61 - 1.49 (m, 1H), 1.40 (s, 9H), 1.05 - 0.93 (m, 3H); LCMS-Method 4 $[M+1, M+3]^+$ = 691.1, 693.1.

Step I: KRAS G12D-compound-2: To a solution of cis-tert-butyl(1-(3-bromo-4-(1,1-dioxido-3,4-dihydrobenzo[f][1,2,5]thiadiazepin-5(2H)-yl)phenyl)-5-((R)-3,4-dimethylpiperazine-1-carbon yl)piperidin-3-yl)carbamate (100 mg, 145 $\mu$mol, 1.0 equiv) in methanol (1.0 mL) was dropwise hydrochloric acid/dioxane (2 M, 1.0 mL, 13.8 equiv). The reaction was stirred at 20 °C for 1 hour. After completion of the reaction, the mixture was concentrated under reduced pressure to give a residue. The residue was purified by reversed-phase flash (column: Sfar C18 40 g D Duo 30$\mu$m; mobile phase: [water(FA)-ACN]; gradient: 10%-50% B over 10 minutes) to afford cis-(5-amino-1-(3-bromo-4-(1,1-dioxido-3,4-dihydrobenzo[f][1,2,5]thiadiazepin-5(2H)-yl)phenyl)piperidin-3-yl)((R)-3,4-dimethylpiperazin-1-yl)methanone (67.85 mg, 77% yield) as off-white solid; $^1$H NMR (400 MHz, CHLOROFORM-d) $\delta$ = 7.88 (d, $J$ = 8.0 Hz, 1H), 7.26 - 7.16 (m, 3H), 6.97 (br t, $J$ = 8.0 Hz, 1H), 6.90 (dd, $J$ = 2.7, 8.8 Hz, 1H), 6.58 - 6.47 (m, 1H), 5.24 - 4.99 (m, 1H), 4.53 - 4.27 (m, 1H), 4.22 - 3.83 (m, 2H), 3.82 - 3.70 (m, 2H), 3.69 - 3.56 (m, 2H), 3.49 - 3.27 (m, 3H), 3.07 - 2.98 (m, 2H), 2.96 (br s, 3H), 2.59 - 2.51 (m, 1H), 2.32 (s, 3H), 2.27 - 2.15 (m, 1H), 2.15 - 1.97 (m, 3H), 1.61 - 1.46 (m, 1H), 1.16 - 1.06 (m, 3H); LCMS $[M+1, M+3]^+$ = 591.2, 593.2; HPLC Rt = 0.808 min.

**Analytical method by SFC:**

Column: Chiralpak IE-3 50×4.6mm I.D., 3$\mu$m;

Mobile phase: Phase A for $CO_2$, and Phase B for EtOH+ACN (0.05% DEA);

Gradient elution: 60% EtOH+ACN(0.05%DEA) in $CO_2$;

Flow rate: 3 mL/min; Detector: PDA;

Colum Temp:35°C; Back Pressure: 100 Bar;

Retention time A: 1.104 min;

Retention time B: 1.424 min.

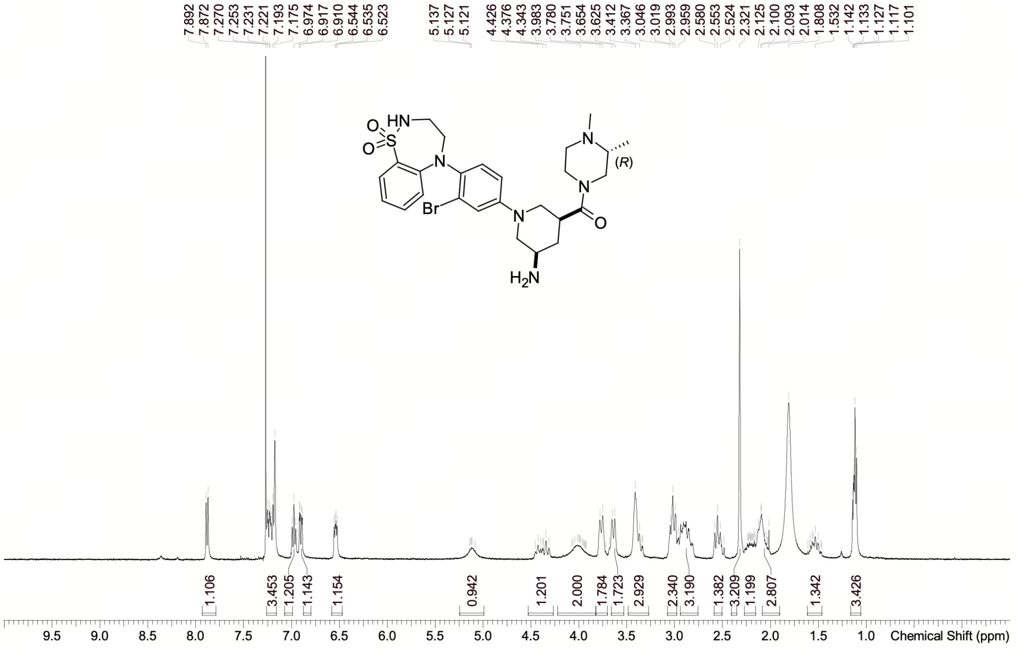

| | |
|---|---|
| Acquisition Time (sec) | 1.9999 |
| Comment | EW53194-1085-P1D CDCl3 BRUKER_03_K_400 MHz |
| Date | 01 Sep 2025 11:29:36 (GMT+08:00) |
| Frequency (MHz) | 400.2800 |
| Nucleus | 1H |
| Number of Transients | 1 |
| Origin | Avance neo400 |
| Original Points Count | 16393 |
| Owner | nmrsu |
| Points Count | 65536 |
| Pulse Sequence | zg |
| Receiver Gain | 18.00 |
| SW(cyclical) (Hz) | 8196.72 |
| Solvent | CHLOROFORM-d |
| Spectrum Offset (Hz) | 2465.3264 |
| Spectrum Type | standard |
| Sweep Width (Hz) | 8196.60 |
| Temperature (degree C) | 24.568 |

Confidential. For research information only

$^1$H NMR (400 MHz, CHLOROFORM-d) δ = 7.88 (d, *J* = 8.0 Hz, 1H), 7.26 - 7.16 (m, 3H), 6.97 (br t, *J* = 8.0 Hz, 1H), 6.90 (dd, *J* = 2.7, 8.8 Hz, 1H), 6.58 - 6.47 (m, 1H), 5.24 - 4.99 (m, 1H), 4.53 - 4.27 (m, 1H), 4.22 - 3.83 (m, 2H), 3.82 - 3.70 (m, 2H), 3.69 - 3.56 (m, 2H), 3.49 - 3.27 (m, 3H), 3.07 - 2.98 (m, 2H), 2.96 (br s, 3H), 2.59 - 2.51 (m, 1H), 2.32 (s, 3H), 2.27 - 2.15 (m, 1H), 2.15 - 1.97 (m, 3H), 1.61 - 1.46 (m, 1H), 1.16 - 1.06 (m, 3H)

#### NMR Spectrum of KRAS G12D-compound-2

RetTime: 0.378 Datafile: D:\DATA\2025\2509\250901\EW53194-1085-P1L.lcd

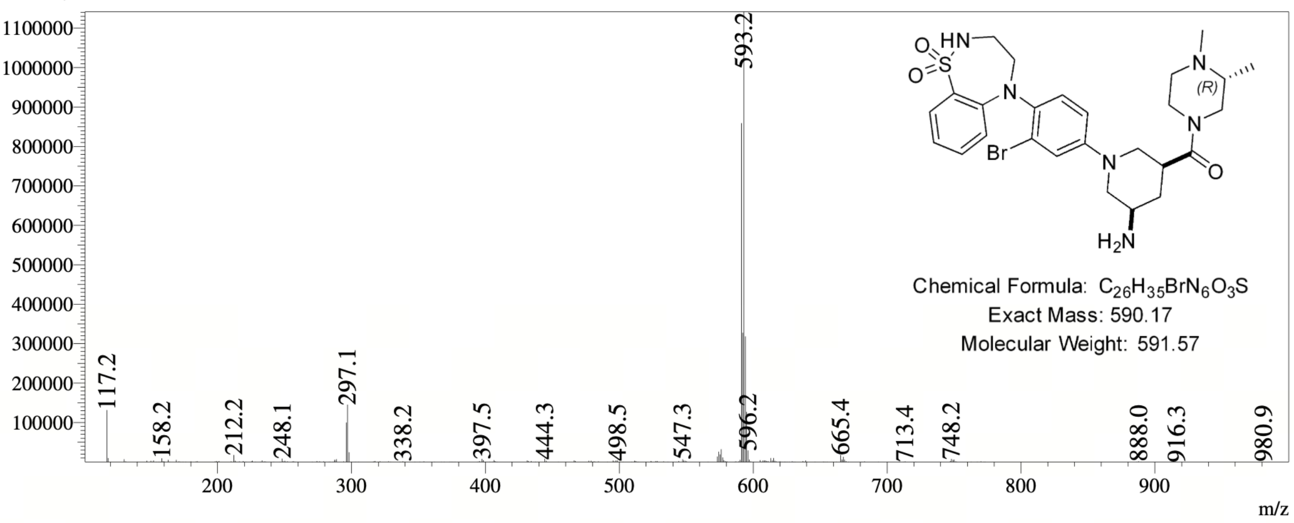

#### MS Spectrum of KRAS G12D-compound-2

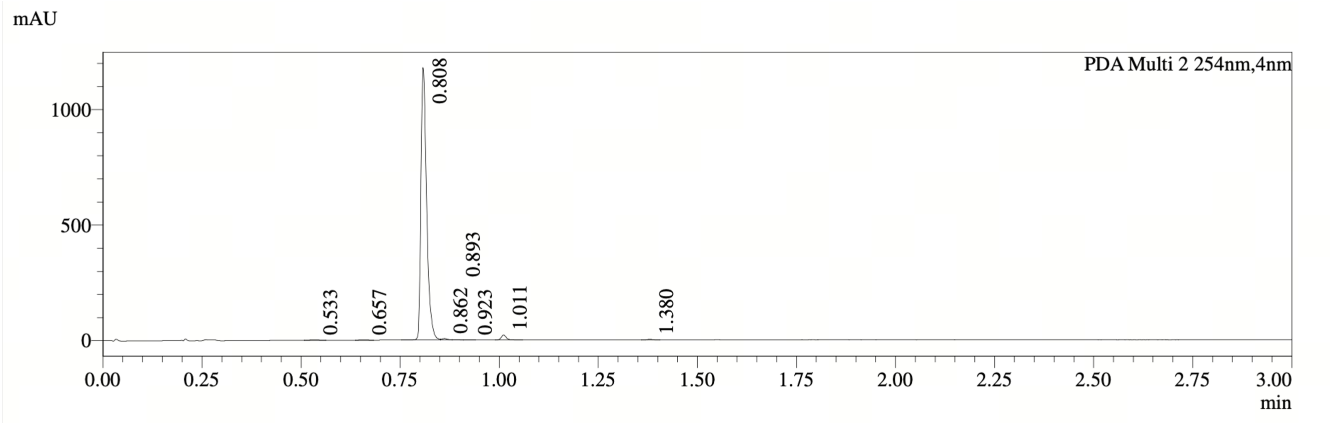

PDA Ch2 254nm

| Peak# | Ret. Time | Width | Height | Height% | Area | Area% |
|---|---|---|---|---|---|---|
| 1 | 0.533 | 0.035 | 1783 | 0.147 | 2311 | 0.191 |
| 2 | 0.657 | 0.027 | 1456 | 0.120 | 1422 | 0.118 |
| 3 | 0.808 | 0.027 | 1179003 | 96.993 | 1173579 | 97.028 |
| 4 | 0.862 | 0.036 | 6357 | 0.523 | 6534 | 0.540 |
| 5 | 0.893 | 0.045 | 1838 | 0.151 | 2094 | 0.173 |
| 6 | 0.923 | 0.054 | 888 | 0.073 | 1222 | 0.101 |
| 7 | 1.011 | 0.025 | 21493 | 1.768 | 19736 | 1.632 |
| 8 | 1.380 | 0.027 | 2737 | 0.225 | 2633 | 0.218 |

**HPLC Chromatogram of KRAS G12D-compound-2**

##### *7.1.3 KRAS G12D-compound-3*

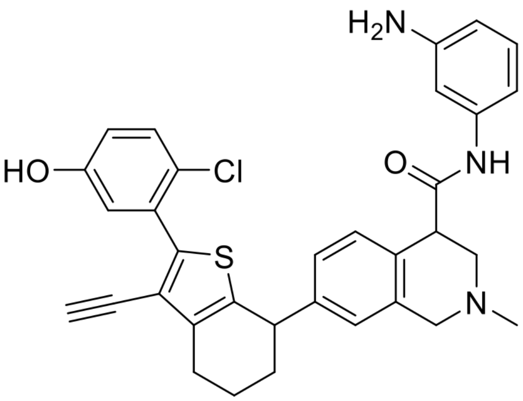

**KRAS G12D-compound-3**. N-(3-aminophenyl)-7-(2-(2-chloro-5-hydroxyphenyl)-3-ethynyl-4,5,6,7-tetrahydrobenzo[b]thiophen-7-yl)-2-methyl-1,2,3,4-tetrahydroisoquinoline-4-carboxamide

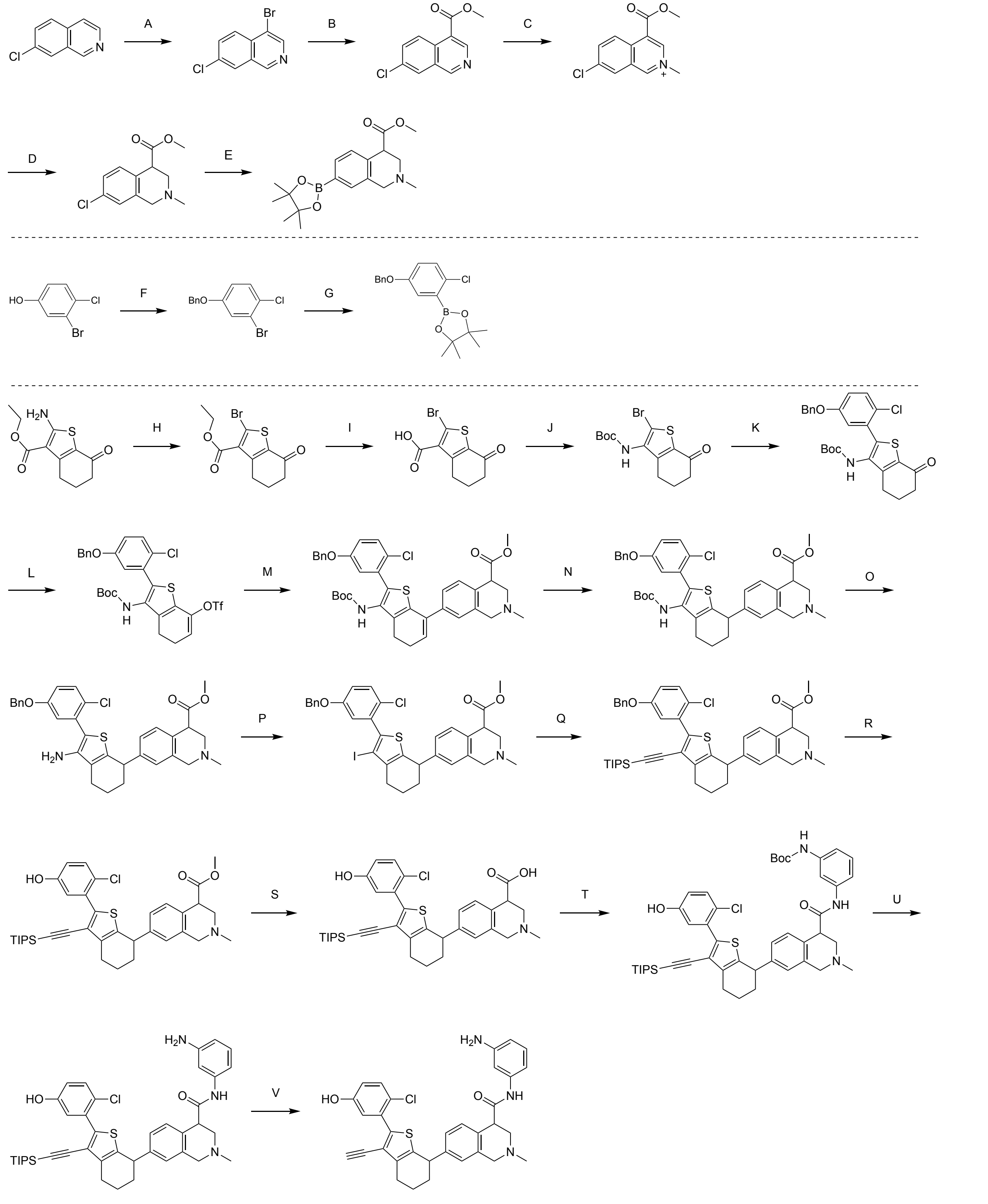

**KRAS G12D-compound-3**

Reagents and conditions: A. KBr, phenyl-$I_3$-iodanediyl diacetate, DCE, 70 °C, 34%; B. WXPS1001, TMEDA, CO, MeOH, THF, 3.5 MPa, 120 °C, 85%; C. MeI, ACN, THF, 40 °C, 99%; D. $NaBH(CN)_3$, EtOH, 25 °C, 95%; E. $Pin_2B_2$, AcOK, $Pd(OAc)_2$, $PCy_3$, dioxane, 110 °C, 15%; F, BnBr, $K_2CO_3$, KI, acetone, 60 °C, 30%; G. $Pin_2B_2$, AcOK, $Pd(dppf)Cl_2$, dioxane, 90 °C, 95%; H. t-BuONO, $CuBr_2$, ACN, 0 - 25 °C, 57%; I. LiOH, THF, EtOH, $H_2O$, 25 °C, 73%; J. DPPA, DIEA, t-BuOH, toluene, 100 °C, 82%; K. $Pd(PPh_3)_4$, $Na_2CO_3$, dioxane, $H_2O$, 100 °C, 85%; L. N-(5-chloropyridin-2-yl)-1,1,1-trifluoro-N-((trifluoromethyl)sulfonyl)methanesulfonamide, LiHMDS, THF, -78 - 25 °C, 89%; M. $Pd(PPh_3)_4$, $Na_2CO_3$, dioxane, $H_2O$, 70 °C, 72%; N. 3% $Pt/Al_2O_3$, $H_2$, THF, 1 MPa, 50 °C, 78%; O. TFA, DCM, 25 °C, 85%; P. $MeI_2$, isopentyl nitrite, ACN, 70 °C, 16%; Q. $Pd(PPh_3)_2Cl_2$, TEA, THF, 70 °C, 76%; R. $BCl_3$, DCM, -78 - 20 °C, 33%; S. LiOH, THF, 40 °C, 99%; T. EDCI, HOBt, pyridine, 30 °C, 50%; U. TFA, DCM, 25 °C, 99%; V. CsF, DMF, 40 °C, 71%.

#### Synthesis of KRAS G12D-compound-3

Step A: 4-bromo-7-chloroisoquinoline. To a solution of 7-chloroisoquinoline (9.00 g, 55.0 mmol, 1.0 equiv) in 1,2-dichloroethane (100 mL) were added potassium bromide (32.7 g, 275 mmol, 5.0 equiv) and phenyl-l3-iodanediyl diacetate (26.6 g, 82.5 mmol, 1.5 equiv) under nitrogen atmosphere. The reaction was stirred at 70 °C for 36 hours. After completion of the reaction, the mixture was cooled to 25 °C . The mixture was filtered and the filtrate was concentrated under reduce pressure. The residue was purified by column chromatography ($SiO_2$, Petroleum ether: Ethyl acetate = 100/1 to 10/1) to afford 4-bromo-7-chloroisoquinoline (4.67 g, 34% yield) as a yellow solid; $^1$H NMR (400 MHz, CHLOROFORM-d) $\delta$= 9.11 (s, 1H), 8.74 (s, 1H), 8.14 (d, $J$ = 8.8 Hz, 1H), 7.99 (d, $J$ = 2.0 Hz, 1H), 7.77 (dd, $J$ = 2.0, 9.2 Hz, 1H).

Step B: methyl 7-chloroisoquinoline-4-carboxylate. Solution 1: 4-bromo-7-chloroisoquinoline (4.50 g, 18.6 mmol, 1.0 equiv) and N1,N1,N2,N2-tetramethylethane-1,2-diamine (6.47 g, 55.7 mmol, 3.0 equiv) were dissolved in methanol (23.0 mL) and tetrahydrofuran (23.0 mL). The resulting solution was delivered by a plunger pump at a flow rate of 3.03 mL/min. CO (from a high-pressure cylinder) was introduced via a mass flow controller (MFC) at 14.4 sccm (1.1 equiv) and the system was maintained at 3.5 MPa. The liquid and gas streams were mixed with a T-shaped mixer and the combined solution was passed into a stainless-steel (SS) fixed-bed reactor (Outer diameter: 12.700 mm (1/2 inch); Internal volume: 100.00 mL) packed with 38.0 g of WXPS1001 (3% Pd@POL)-4 and held at 120 °C . After completion of the reaction, the mixture was cooled to 25 °C . The mixture was concentrated under reduced pressure to give a residue. The residue was diluted with water (50 mL) and extracted with ethyl acetate (3 × 30 mL). The combined organic layers were washed with brine (50 mL), dried over anhydrous sodium sulfate, filtered and concentrated under reduced pressure to afford methyl 7-chloroisoquinoline-4-carboxylate (3.70 g, 85% yield) as a yellow solid; LCMS-Method 1 [M+1] $^+$= 222.0.

Step C: 7-chloro-4-(methoxycarbonyl)-2-methylisoquinolin-2-ium. To a solution of methyl 7-chloroisoquinoline-4-carboxylate (3.70 g, 16.7 mmol, 1.0 equiv) in acetonitrile (40.0 mL) and tetrahydrofuran (10.0 mL) was added iodomethane (21.3 g, 150 mmol, 9.0 equiv). The reaction was stirred at 40 °C for 16 hours. After completion of the reaction, the mixture was concentrated under reduced pressure to afford 7-chloro-4-(methoxycarbonyl)-2-methylisoquinolin-2-ium (3.90 g, 99% yield) as a yellow solid; LCMS-Method 1 [M] $^+$= 236.1.

Step D: methyl 7-chloro-2-methyl-1,2,3,4-tetrahydroisoquinoline-4-carboxylate. To a solution of sodium cyanotrihydroborate (5.99 g, 95.3 mmol, 6.0 equiv) in ethanol (150 mL) was added 7-chloro-4-(methoxycarbonyl)-2-methylisoquinolin-2-ium (3.76 g, 15.9 mmol, 1.0 equiv). The reaction was stirred at 25 °C for 2 hours. After completion of the reaction, the mixture was quenched with hydrogen chloride (50.0 mL, 1.0 M) aqueous solution at 0 °C and adjusted pH to 8 with sodium hydrogen carbonate aqueous solution. The mixture was extracted with ethyl acetate (3 × 50 mL). The combined organic layers were washed with brine (50 mL), dried over anhydrous sodium sulfate, filtered and concentrated under reduced pressure to give a residue. The residue was purified by column chromatography ($SiO_2$, Petroleum ether: Ethyl acetate = 100/1 to 1/1) to afford methyl 7-chloro-2-methyl-1,2,3,4-tetrahydroisoquinoline-4-carboxylate (3.80 g, 95% yield) as a white solid; LCMS-Method 27 [M+1] $^+$= 239.9.

Step E: methyl2-methyl-7-(4,4,5,5-tetramethyl-1,3,2-dioxaborolan-2-yl)-1,2,3,4-tetrahydroisoquinoline-4-carboxylate. A mixture of methyl 7-chloro-2-methyl-1,2,3,4-tetrahydroisoquinoline-4-carboxylate (3.00 g, 12.5 mmol, 1.0 equiv), 4,4,4',4',5,5,5',5'-octamethyl-2,2'-bi(1,3,2-dioxaborolane) (12.7 g, 50.1 mmol, 4.0 equiv), palladium diacetate (281 mg, 1.25 mmol, 0.1 equiv), tricyclohexylphosphine (702 mg, 2.50 mmol, 0.2 equiv) and potassium acetate (3.68 g, 37.6 mmol, 3.0 equiv) in dioxane (70.0 mL) was degassed and purged with nitrogen for three times, and then the reaction was stirred at 110 °C for 16 hours under nitrogen atmosphere. After completion of the reaction, the mixture was cooled to 25 °C . The mixture was filtered and the filtrate was purified by column chromatography ($SiO_2$, Petroleum ether: Ethyl acetate = 100/1 to 5/1 to 1/1) and reversed-phase flash (Sfar C18 300 g D Duo 30 $\mu$m; mobile phase: [water-ACN]; gradient: 33%-57% B over 11 min) to afford methyl 2-methyl-7-(4,4,5,5-tetramethyl-1,3,2-dioxaborolan-2-yl)-1,2,3,4-tetrahydroisoquinoline-4-carboxylate (600 mg, 15% yield) as a white solid; LCMS-Method 4 [M+1] $^{+}$= 332.1.

Step F: 4-(benzyloxy)-2-bromo-1-chlorobenzene. To a solution of 3-bromo-4-chlorophenol (10.0 g, 48.2 mmol, 1.0 equiv) in acetone (100 mL) were added potassium carbonate (20.0 g, 145 mmol, 3.0 equiv), potassium iodide (800 mg, 4.82 mmol, 0.1 equiv) and (bromomethyl)benzene (14.4 g, 84.2 mmol, 1.75 equiv). The reaction was stirred at 60 °C for 12 hours under nitrogen atmosphere. After completion of the reaction, the mixture was cool to 25 °C . The mixture was diluted with water (100 mL) and extracted with ethyl acetate (3 × 50 mL). The combined organic layers were washed with brine (50 mL), dried over anhydrous sodium sulfate, filtered and concentrated under reduced pressure to give a residue. The residue was purified by prep-HPLC(column: Phenomenex luna C18 250mm*100mm*10um;mobile phase: [$H_2O$(0.225% FA)-ACN]; gradient:65%-85% B over 20.0 min) to afford 4-(benzyloxy)-2-bromo-1-chlorobenzene (4.30 g, 30% yield) as yellow oil; $^1$H NMR (400 MHz, CHLOROFORM-d) $\delta$= 7.41 (d, $J$ = 4.4 Hz, 4H), 7.39 - 7.35 (m, 1H), 7.34 (d, $J$ = 8.8 Hz, 1H), 7.26 (d, $J$ = 2.8 Hz, 1H), 6.88 (dd, $J$ = 2.8, 8.8 Hz, 1H), 5.04 (s, 2H).

Step G: 2-(5-(benzyloxy)-2-chlorophenyl)-4,4,5,5-tetramethyl-1,3,2-dioxaborolane. To a solution of 4-(benzyloxy)-2-bromo-1-chlorobenzene (4.00 g, 13.4 mmol, 1.0 equiv) and 4,4,4',4',5,5,5',5'-octamethyl-2,2'-bi(1,3,2-dioxaborolane) (5.12 g, 20.2 mmol, 1.5 equiv) in dioxane (50 mL) were added potassium acetate (2.64 g, 26.88 mmol, 2.0 equiv) and Pd(dppf)$Cl_2$ (984 mg, 1.34 mmol, 0.1 equiv). The reaction was degassed and purged with nitrogen for three times, and then stirred at 90 °C for 5 hours under nitrogen atmosphere. After completion of the reaction, the mixture was cool to 25 °C . The mixture was diluted with water (50 mL) and extracted with ethyl acetate (3 × 50 mL). The combined organic layers were washed with brine (50 mL), dried over anhydrous sodium sulfate, filtered and concentrated under reduced pressure to give a residue. The residue was purified by flash silica gel chromatography (ISCO®; 80 g SepaFlash® Silica Flash Column, Eluent of 0$\tilde{5}$% Ethyl acetate/Commercial hexanes gradient @ 100 mL/min) to afford 2-(5-(benzyloxy)-2-chlorophenyl)-4,4,5,5-tetramethyl-1,3,2-dioxaborolane (5.50 g, 95% yield) as a white solid; $^1$H NMR (400 MHz, CHLOROFORM-d) $\delta$= 7.46 - 7.30 (m, 6H), 7.25 (d, $J$ = 8.8 Hz, 1H), 6.94 (dd, $J$ = 3.2, 8.8 Hz, 1H), 5.06 (s, 2H), 1.38 (s, 12H).

Step H: ethyl 2-bromo-7-oxo-4,5,6,7-tetrahydrobenzo[b]thiophene-3-carboxylate. To a solution of copper(II) bromide (12.4 g, 55.6 mmol, 1.4 equiv) in acetonitrile (200 mL) was added dropwise tert-butyl nitrite (4.91 g, 47.6 mmol, 1.2 equiv) at 0 °C . After addition, the mixture was stirred at 0 °C for 1 hour, and then ethyl 2-amino-7-oxo-4,5,6,7-tetrahydrobenzo[b]thiophene-3-carboxylate (9.50 g, 39.7 mmol, 1.0 equiv) in acetonitrile (200 mL) was added dropwise at 0 °C . The reaction was stirred at 25 °C for 15 hours. After completion of the reaction, the mixture was diluted with water (200 mL) and extracted with ethyl acetate (3 × 100 mL). The combined organic layers were washed with brine (100 mL), dried over anhydrous sodium sulfate, filtered and concentrated under reduced pressure to give a residue. The residue was purified by column chromatography ($SiO_2$, Petroleum ether: Ethyl acetate = 100/1 to 5/1) to afford ethyl 2-bromo-7-oxo-4,5,6,7-tetrahydrobenzo[b]thiophene-3-carboxylate (9.30 g, 57% yield) as a white solid; LCMS-Method 1 [M+1, M+3] $^{+}$= 302.9, 304.9.

Step I: 2-bromo-7-oxo-4,5,6,7-tetrahydrobenzo[b]thiophene-3-carboxylic acid. To a solution of ethyl 2-bromo-7-oxo-4,5,6,7-tetrahydrobenzo[b]thiophene-3-carboxylate (9.00 g, 29.7 mmol, 1.0 equiv) in ethanol (30.0 mL) and tetrahydrofuran (30.0 mL) was added lithium hydroxide hydrate (2.49 g, 59.4 mmol, 2.0 equiv) in water (10.0 mL). The reaction was stirred at 25 °C for 16 hours. After completion of the reaction, the mixture was adjusted pH to 2 with hydrogen chloride (1.0 M) aqueous solution. The mixture was diluted with water (200 mL) and extracted with ethyl acetate (3 × 100 mL). The combined organic layers were washed with brine (100 mL), dried over anhydrous sodium sulfate, filtered and concentrated under reduced pressure to give a residue. The residue was purified by prep-HPLC (column: Phenomenex luna C18 250 × 50 mm, 10 $\mu$m; mobile phase: [$H_2O$ (0.05% HCl)-ACN]; gradient: 15% - 45% B over 25.0 min) to afford 2-bromo-7-oxo-4,5,6,7-tetrahydrobenzo[b]thiophene-3-carboxylic acid (6.00 g, 73% yield) as a white solid; LCMS-Method 1 [M+1, M+3] $^+$= 274.9, 276.9.

Step J: tert-butyl (2-bromo-7-oxo-4,5,6,7-tetrahydrobenzo[b]thiophen-3-yl)carbamate. To a solution of 2-bromo-7-oxo-4,5,6,7-tetrahydrobenzo[b]thiophene-3-carboxylic acid (3.00 g, 10.9 mmol, 1.0 equiv), 4A molecular sieves (3.00 g) and N-ethyl-N-isopropylpropan-2-amine (5.64 g, 43.6 mmol, 4.0 equiv) in 2-methylpropan-2-ol (30.0 mL) and toluene (60.0 mL) was added dropwise diphenyl phosphoryl azide (3.60 g, 13.1 mmol, 1.2 equiv) in toluene (30.0 mL). The reaction was stirred at 100 °C for 16 hours. After completion of the reaction, the mixture was cooled to 25 °C . The mixture was diluted with water (100 mL) and extracted with ethyl acetate (3 × 100 mL). The combined organic layers were washed with brine (100 mL), dried over anhydrous sodium sulfate, filtered and concentrated under reduced pressure to give a residue. The residue was purified by prep-HPLC (column: Phenomenex luna C18 (250 × 70 mm, 10 $\mu$m); mobile phase: [$H_2O$ (0.05% HCl) -ACN]; gradient: 45%-75% B over 20.0 min) to afford tert-butyl (2-bromo-7-oxo-4,5,6,7-tetrahydrobenzo[b]thiophen-3-yl)carbamate (3.20 g, 82% yield) as a white solid; LCMS-Method 1 [M+1, M+3] $^+$= 345.9, 347.9.

Step K: tert-butyl (2-(5-(benzyloxy)-2-chlorophenyl)-7-oxo-4,5,6,7-tetrahydrobenzo[b]thiophen-3-yl)carbamate. A mixture of tert-butyl (2-bromo-7-oxo-4,5,6,7-tetrahydrobenzo[b]thiophen-3-yl)carbamate (3.10 g, 8.95 mmol, 1.0 equiv), 2-(5-(benzyloxy)-2-chlorophenyl)-4,4,5,5-tetramethyl-1,3,2-dioxaborolane (3.24 g, 9.40 mmol, 1.05 equiv), tetrakis(triphenylphosphine) palladium(0) (1.03 g, 895 $\mu$mol, 0.1 equiv) and sodium carbonate (2.85 g, 26.9 mmol, 3.0 equiv) in dioxane (40.0 mL) and water (4.0 mL) was degassed and purged with nitrogen for 3 times, and then the reaction was stirred at 100 °C for 5 hours under nitrogen atmosphere. After completion of the reaction, the mixture was cooled to 25 °C . The mixture was filtered and the filtrate was concentrated under reduced pressure to give a residue. The residue was purified by prep-HPLC (column: Phenomenex luna C18 (250 × 70 mm, 10 $\mu$m); mobile phase: [$H_2O$ (0.05% HCl)-ACN]; gradient: 50%-95% B over 30.0 min) to afford tert-butyl (2-(5-(benzyloxy)-2-chlorophenyl)-7-oxo-4,5,6,7-tetrahydrobenzo[b]thiophen-3-yl)carbamate (3.67 g, 85% yield) as a white solid; LCMS-Method 1 [M+1] $^+$= 484.2.

Step L: 2-(5-(benzyloxy)-2-chlorophenyl)-3-((tert-butoxycarbonyl)amino)-4,5-dihydroben zo[b]thiophen-7-yltrifluoromethanesulfonate. To a solution of tert-butyl (2-(5-(benzyloxy)-2-chlorophenyl)-7-oxo-4,5,6,7-tetrahydrobenzo[b]thiophen-3-yl)carbamate (2.00 g, 4.13 mmol, 1.0 equiv) in tetrahydrofuran (160.0 mL) was added dropwise lithium bis(trimethylsilyl)amide (10.3 mL, 1.0 M, 2.5 equiv) at -78 °C . After addition, the mixture was stirred at -78 °C for 0.5 hour, and then N-(5-chloropyridin-2-yl)-1,1,1-trifluoro-N-((trifluoromethyl)sulfonyl)methanesulfonamide (2.43 g, 6.20 mmol, 1.5 equiv) in tetrahydrofuran (40.0 mL) was added dropwise at -78 °C . The reaction was stirred at 25 °C for 1.5 hours under nitrogen atmosphere. After completion of the reaction, the mixture was quenched with saturated ammonium chloride aqueous solution

(20 mL) at 0 °C . The mixture was diluted with water (200 mL) and extracted with ethyl acetate (3 × 100 mL). The combined organic layers were washed with brine (30 mL), dried over anhydrous sodium sulfate, filtered and concentrated under reduced pressure to give a residue. The residue was purified by column chromatography ($SiO_2$, Petroleum ether: Ethyl acetate = 100/1 to 5/1) to afford 2-(5-(benzyloxy)-2-chlorophenyl)-3-((tert-butoxycarbonyl)amino)-4,5-dihydrobenzo[b]thiophen-7-yl trifluoromethanesulfonate (2.30 g, 89% yield) as a yellow solid; LCMS-Method 1 $[M+23]^{+}$= 638.2.

Step M: methyl7-(2-(5-(benzyloxy)-2-chlorophenyl)-3-((tert-butoxycarbonyl)amino)-4,5-dihydrobenzo[b]thiophen-7-yl)-2-methyl-1,2,3,4-tetrahydroisoquinoline-4-carboxylate. A mixture of 2-(5-(benzyloxy)-2-chlorophenyl)-3-((tert-butoxycarbonyl)amino)-4,5-dihydrobenzo[b]thiophen-7-yl trifluoromethanesulfonate (1.05 g, 1.70 mmol, 1.0 equiv), methyl 2-methyl-7-(4,4,5,5-tetramethyl-1,3,2-dioxaborolan-2-yl)-1,2,3,4-tetrahydroisoquinoline-4-carboxylate (565 mg, 1.70 mmol, 1.0 equiv), tetrakis (triphenylphosphine) palladium(0) (197 mg, 170 $\mu$mol, 0.1 equiv) and sodium carbonate (542 mg, 5.11 mmol, 3.0 equiv) in dioxane (50.0 mL) and water (1.0 mL) was degassed and purged with nitrogen for 3 times, and then the reaction was stirred at 70 °C for 3 hours under nitrogen atmosphere. After completion of the reaction, the mixture was cooled to 25 °C . The mixture was diluted with water (100 mL) and extracted with ethyl acetate (3 × 50 mL). The combined organic layers were washed with brine (50 mL), dried over anhydrous sodium sulfate, filtered and concentrated under reduced pressure to give a residue. The residue was purified by column chromatography ($SiO_2$, Petroleum ether: Ethyl acetate = 100/1 to 1/2) and reversed-phase flash Sfar (C18 300 g D Duo 30 $\mu$m; mobile phase: [water ($NH_3.H_2O$)-ACN]; gradient: 33%-57% B over 11 min) to afford methyl 7-(2-(5-(benzyloxy)-2-chlorophenyl)-3-((tert-butoxycarbonyl)amino)-4,5-dihydrobenzo[b]thiophen-7-yl)-2-methyl-1,2,3,4-tetrahydroisoquinoline-4-carboxylate (900 mg, 72% yield) as a white solid; LCMS-Method 1 $[M+1]^{+}$= 671.3.

Step N: methyl7-(2-(5-(benzyloxy)-2-chlorophenyl)-3-((tert-butoxycarbonyl)amino)-4,5,6,7-tetrahydrobenzo[b]thiophen-7-yl)-2-methyl-1,2,3,4-tetrahydroisoquinoline-4-carboxylate. Solution 1: methyl7-(2-(5-(benzyloxy)-2-chlorophenyl)-3-((tert-butoxycarbonyl)amino)-4,5-dihydrobenzo[b]thiophen-7-yl)-2-methyl-1,2,3,4-tetrahydroisoquinoline-4-carboxylate (600 mg, 894 $\mu$mol, 1.0 equiv) was dissolved in tetrahydrofuran (60.0 mL). The resulting solution was delivered by a plunger pump at a flow rate of 0.40 mL/min. hydrogen (from a high-pressure cylinder) was introduced via a mass flow controller (MFC) at 11.3 sccm (80.0 equiv) and the system was maintained at 1.0 MPa. The liquid and gas streams were mixed with a T-shaped mixer and the combined solution was passed into a stainless-steel(SS) fixed-bed reactor (Outer diameter: 6.350 mm (1/4 inch); Internal volume: 5.00 mL) packed with 1.0 g of WXSC1064 (3% $Pt/Al_2O_3$) and held at 50 °C . Upon complete injection of the reaction solution, the solvent feed was switched to methanol at 0.4 mL/min and the reactor was flushed for 1 hour. After completion of the reaction, the mixture was concentrated under reduced pressure to give a residue. The residue was purified by column chromatography ($SiO_2$, Petroleum ether: Ethyl acetate = 100/1 to 1/2) to afford methyl 7-(2-(5-(benzyloxy)-2-chlorophenyl)-3-((tert-butoxycarbonyl)amino)-4,5,6,7-tetrahydrobenzo[b]thiophen-7-yl)-2-methyl-1,2,3,4-tetrahydroisoquinoline-4-carboxylate (470 mg, 78% yield) as a yellow solid; LCMS-Method 15 $[M+1]^{+}$= 673.3.

Step O: methyl7-(3-amino-2-(5-(benzyloxy)-2-chlorophenyl)-4,5,6,7-tetrahydrobenzo[b]thiophen-7-yl)-2-methyl-1,2,3,4-tetrahydroisoquinoline-4-carboxylate. To a solution of methyl7-(2-(5-(benzyloxy)-2-chlorophenyl)-3-((tert-butoxycarbonyl)amino)-4,5,6,7-tetrahydrobenzo[b]thiophen-7-yl)-2-methyl-1,2,3,4-tetrahydroisoquinoline-4-carboxylate (470 mg, 698 $\mu$mol, 1.0 equiv) in dichloromethane (3.0 mL) was added 2,2,2-trifluoroacetic acid (3.07 g, 26.9 mmol, 38.6 equiv).

The reaction was stirred at 25 °C for 16 hours. After completion of the reaction, the mixture was adjusted pH to 8 with sodium hydrogen carbonate aqueous solution. The mixture was extracted with ethyl acetate (3 × 30 mL) and the combined organic layers were washed with brine (30 mL), dried over anhydrous sodium sulfate, filtered and concentrated under reduced pressure to afford methyl 7-(3-amino-2-(5-(benzyloxy)-2-chlorophenyl)-4,5,6,7-tetrahydrobenzo[b]thiophen-7-yl)-2-methyl-1,2,3,4-tetrahydroisoquinoline-4-carboxylate (340 mg, 85% yield) as a yellow solid; LCMS-Method 15 $[M+1]^+$= 573.2.

Step P: methyl 7-(2-(5-(benzyloxy)-2-chlorophenyl)-3-iodo-4,5,6,7-tetrahydrobenzo[b]thiophen-7-yl)-2-methyl-1,2,3,4-tetrahydroisoquinoline-4-carboxylate. To a solution of isopentyl nitrite (79.7 mg, 680 $\mu$mol, 1.3 equiv) in acetonitrile (10.0 mL) was added diiodomethane (210 mg, 785 $\mu$mol, 1.5 equiv). After addition, the mixture was stirred at 70 °C for 0.5 hour, and then methyl 7-(3-amino-2-(5-(benzyloxy)-2-chlorophenyl)-4,5,6,7-tetrahydrobenzo[b]thiophen-7-yl)-2-methyl-1,2,3,4-tetrahydroisoquinoline-4-carboxylate (300 mg, 523 $\mu$mol, 1.0 equiv) in acetonitrile (2.0 mL) was added dropwise at 70 °C . The reaction was stirred at 70 °C for 3.5 hours. After completion of the reaction, the mixture was cooled to 25 °C . The mixture was diluted with water (50 mL) and extracted with ethyl acetate (3 × 30 mL). The combined organic layers were washed with brine (30 mL), dried over anhydrous sodium sulfate, filtered and concentrated under reduced pressure to give a residue. The residue was purified by reversed-phase flash (Sfar C18 80 g D Duo 30 $\mu$m; mobile phase: [water (FA)-ACN]; gradient: 45% - 57% B over 11 min) to afford methyl 7-(2-(5-(benzyloxy)-2-chlorophenyl)-3-iodo-4,5,6,7-tetrahydrobenzo[b]thiophen-7-yl)-2-methyl-1,2,3,4-tetrahydroisoquinoline-4-carboxylate (110 mg, 16% yield) as a yellow solid; LCMS-Method 31 $[M+1]^+$= 684.2.

Step Q: methyl7-(2-(5-(benzyloxy)-2-chlorophenyl)-3-((triisopropylsilyl)ethynyl)-4,5,6,7-tetr ahydrobenzo[b]thiophen-7-yl)-2-methyl-1,2,3,4-tetrahydroisoquinoline-4-carboxylate. A mixture of methyl 7-(2-(5-(benzyloxy)-2-chlorophenyl)-3-iodo-4,5,6,7-tetrahydrobenzo[b]thiophen-7-yl)-2-methyl-1,2,3,4-tetrahydroisoquinoline-4-carboxylate (100 mg, 146 $\mu$mol, 1.0 equiv), ethynyltriisopropylsilane (107 mg, 585 $\mu$mol, 4.0 equiv), bis-triphenylphosphine-palladium(II) chloride (51.3 mg, 73.1 $\mu$mol, 0.5 equiv) and copper(I) iodide (27.8 mg, 146 $\mu$mol, 1.0 equiv) in tetrahydrofuran (5.0 mL) and triethylamine (5.0 mL) was degassed and purged with nitrogen for 3 times, and then the reaction was stirred at 70 °C for 3 hours under nitrogen atmosphere. After completion of the reaction, the mixture was cooled to 25 °C . The mixture was diluted with water (50 mL) and extracted with ethyl acetate (3 × 30 mL). The combined organic layers were washed with brine (50 mL), dried over anhydrous sodium sulfate, filtered and concentrated under reduced pressure to give a residue. The residue was purified by prep-TLC ($SiO_2$, Petroleum ether: Ethyl acetate = 0/1) to afford methyl 7-(2-(5-(benzyloxy)-2-chlorophenyl)-3-((triisopropylsilyl)ethynyl)-4,5,6,7-tetrahydrobenzo[b]thiophen-7-yl)-2-methyl-1,2,3,4-tetrahydroisoquinoline-4-carboxylate (100 mg, 76% yield) as a yellow solid; LCMS-Method 31 $[M+1, M+3]^+$= 738.4, 740.1.

Step R: methyl7-(2-(2-chloro-5-hydroxyphenyl)-3-((triisopropylsilyl)ethynyl)-4,5,6,7-tetrahydrob enzo[b]thiophen-7-yl)-2-methyl-1,2,3,4-tetrahydroisoquinoline-4-carboxylate. To a solution of methyl 7-(2-(5-(benzyloxy)-2-chlorophenyl)-3-((triisopropylsilyl)ethynyl)-4,5,6,7-tetrahydrobenzo[b]thiophen-7-yl)-2-methyl-1,2,3,4-tetrahydroisoquinoline-4-carboxylate (98.0 mg, 133 $\mu$mol, 1.0 equiv) in dichloromethane (20.0 mL) was added trichloroborane (664 $\mu$L, 1.0 M, 5.0 equiv) at -78 °C . The mixture was stirred at 20 °C for 2 hours. After completion of the reaction, the mixture was quenched with sodium hydrogen carbonate aqueous solution (2 mL) at 0 °C . The mixture was purified by reversed-phase flash (Sfar C18 30 g D Duo 30 $\mu$m; mobile phase: [water (FA) - ACN]; gradient: 53%-80% B over 11 min) to afford methyl 7-(2-(2-chloro-5-hydroxyphenyl)-3-((triisopropylsilyl)ethynyl)-4,5,6,7-

tetrahydrobenzo[b]thiophen-7-yl)-2-methyl-1,2,3,4-tetrahydroisoquinoline-4-carboxylate (34.0 mg, 33% yield) as a yellow solid; LCMS-Method 31 $[M+1]^+$= 648.3.

Step S: 7-(2-(2-chloro-5-hydroxyphenyl)-3-((triisopropylsilyl)ethynyl)-4,5,6,7-tetrahydrobenzo[b]thiophen-7-yl)-2-methyl-1,2,3,4-tetrahydroisoquinoline-4-carboxylicacid. To a solution of methyl 7-(2-(2-chloro-5-hydroxyphenyl)-3-((triisopropylsilyl)ethynyl)-4,5,6,7-tetrahydrobenzo[b]thiophen-7-yl)-2-methyl-1,2,3,4-tetrahydroisoquinoline-4-carboxylate (32.0 mg, 49.4 $\mu$mol, 1.0 equiv) in tetrahydrofuran (3.0 mL) was added lithium hydroxide hydrate (247 $\mu$L, 2.0 M, 10.0 equiv). The reaction was stirred at 40 °C for 8 hours. After completion of the reaction, the mixture was cooled to 25 °C . The mixture was concentrated under reduced pressure to afford 7-(2-(2-chloro-5-hydroxyphenyl)-3-((triisopropylsilyl)ethynyl)-4,5,6,7-tetrahydrobenzo[b]thiophen-7-yl)-2-methyl-1,2,3,4-tetrahydroisoquinoline-4-carboxylicacid (31.0 mg, 99% yield) as a yellow solid; LCMS-Method 15 $[M+1]^+$= 634.3.

Step T: tert-butyl(3-(7-(2-(2-chloro-5-hydroxyphenyl)-3-((triisopropylsilyl)ethynyl)-4,5,6,7-tetrahydrobenzo[b]thiophen-7-yl)-2-methyl-1,2,3,4-tetrahydroisoquinoline-4-carboxamido)phenyl)carbamate. To a solution of 7-(2-(2-chloro-5-hydroxyphenyl)-3-((triisopropylsilyl)ethynyl)-4,5,6,7-tetrahydrobenzo[b]thiophen-7-yl)-2-methyl-1,2,3,4-tetrahydroisoquinoline-4-carboxylic acid (27.0 mg, 42.6 $\mu$mol, 1.0 equiv), tert-butyl (3-aminophenyl)carbamate (88.6 mg, 426 $\mu$mol, 10.0 equiv) and 4A molecular sieves (50 mg) in pyridine (2.0 mL) were added 1-ethyl-(3-(3-dimethylamino)propyl)-carbodiimide hydrochloride (81.6 mg, 426 $\mu$mol, 10.0 equiv) and 1H-benzo[d][1,2,3]triazol-1-ol (57.5 mg, 426 $\mu$mol, 10.0 equiv). The reaction was stirred at 30 °C for 1 hour. After completion of the reaction, the mixture was cooled to 25 °C . The mixture was purified by reversed-phase flash (Sfar C18 80 g D Duo 30 $\mu$m; mobile phase: [water (FA)-ACN]; gradient: 20% - 80% B over 11 min) to afford tert-butyl(3-(7-(2-(2-chloro-5-hydroxyphenyl)-3-((triisopropylsilyl)ethynyl)-4,5,6,7-tetrahydrobenzo[b]thiophen-7-yl)-2-methyl-1,2,3,4-tetrahydroisoquinoline-4-carboxamido)phenyl)carbamate (20.0 mg, 50% yield) as a yellow solid; LCMS-Method 15 $[M+1]^+$= 824.3.

Step U: N-(3-aminophenyl)-7-(2-(2-chloro-5-hydroxyphenyl)-3-((triisopropylsilyl)ethynyl)-4,5,6,7-tetrahydrobenzo[b]thiophen-7-yl)-2-methyl-1,2,3,4-tetrahydroisoquinoline-4-carboxamide. To a solution of tert-butyl(3-(7-(2-(2-chloro-5-hydroxyphenyl)-3-((triisopropylsilyl)ethynyl)-4,5,6,7-tetrahydrobenzo[b]thiophen-7-yl)-2-methyl-1,2,3,4-tetrahydroisoquinoline-4-carboxamido)phenyl)carbamate (16.0 mg, 19.4 $\mu$mol, 1.0 equiv) in dichloromethane (3.0 mL) was added 2,2,2-trifluoroacetic acid (1.34 g, 11.8 mmol, 605 equiv). The reaction was stirred at 25 °C for 1 hour. After completion of the reaction, the mixture was adjusted pH to 8 with sodium hydrogen carbonate aqueous solution. The mixture was extracted with dichloromethane (3 × 30 mL). The combined organic layers were washed with brine (30 mL), dried over anhydrous sodium sulfate, filtered and concentrated under reduced pressure to afford N-(3-aminophenyl)-7-(2-(2-chloro-5-hydroxyphenyl)-3-((triisopropylsilyl)ethynyl)-4,5,6,7-tetrahydrobenzo[b]thiophen-7-yl)-2-methyl-1,2,3,4-tetrahydroisoquinoline-4-carboxamide (14.0 mg, 99% yield) as a yellow solid; LCMS-Method 15 $[M+1]^+$= 724.3.

Step V: KRAS G12D-compound-3. To a solution of N-(3-aminophenyl)-7-(2-(2-chloro-5-hydroxyphenyl)-3-((triisopropylsilyl)ethynyl)-4,5,6,7-tetrahydrobenzo[b]thiophen-7-yl)-2-methyl-1,2,3,4-tetrahydroisoquinoline-4-carboxamide (13.0 mg, 17.9 $\mu$mol, 1.0 equiv) in N,N-dimethylformamide (2.0 mL) was added cesium fluoride (27.3 mg, 179 $\mu$mol, 10.0 equiv). The reaction was stirred at 40 °C for 1 hour. After completion of the reaction, the mixture was filtered and the filtrate was purified by reversed-phase flash (Sfar C18 30 g D Duo 30 $\mu$m; mobile phase: [water ($NH_3{\cdot}H_2O$)-ACN]; gradient: 33% - 57% B over 11 min) to afford N-(3-aminophenyl)-7-(2-(2-chloro-5-hydroxyphenyl)-3-ethynyl-4,5,6,7-tetrahydrobenzo[b]thiophen-7-yl)-2-methyl-1,2,3,4-tetrahydroisoquinoline-4-carboxamide (7.34 mg, 71% yield) as an off-white solid; $^1$H NMR (400 MHz, DMSO-d6) $\delta$= 10.23 (d, $J$ = 3.6 Hz, 1H),

9.83 (d, $J$ = 6.8 Hz, 1H), 7.30 (d, $J$ = 8.4 Hz, 1H), 7.16 (d, $J$ = 7.6 Hz, 1H), 7.08 - 6.99 (m, 2H), 6.92 - 6.83 (m, 3H), 6.80 (dt, $J$ = 1.6, 8.4 Hz, 1H), 6.65 - 6.60 (m, 1H), 6.24 (br d, $J$ = 8.4 Hz, 1H), 5.03 (br s, 2H), 4.19 (s, 1H), 4.17 - 4.10 (m, 1H), 3.78 (br s, 1H), 3.70 - 3.61 (m, 1H), 3.56 - 3.45 (m, 1H), 2.98 - 2.87 (m, 1H), 2.85 - 2.76 (m, 1H), 2.68 - 2.62 (m, 2H), 2.42 (s, 3H), 2.20 - 2.07 (m, 1H), 2.00 - 1.87 (m, 1H), 1.86 - 1.69 (m, 2H); LCMS [M+1] $^{+}$= 568.2; HPLC Rt = 3.002 min;

Analytical method by SFC:

Column: Chiralpak IG-3 50 × 4.6 mm I.D., 3 $\mu$m;

Mobile phase: Phase A for $CO_2$, and Phase B for IPA+ACN (0.05%DEA);

Gradient elution: IPA+ACN (0.05%DEA) from 20% to 60% in $CO_2$;

Flow rate: 3 mL/min; Detector: PDA;

Column Temp: 35 °C ; Back Pressure: 100 Bar;

Retention time A: 2.246 min;

Retention time B: 2.965 min;

Retention time C: 3.225 min;

Retention time D: 3.463 min.

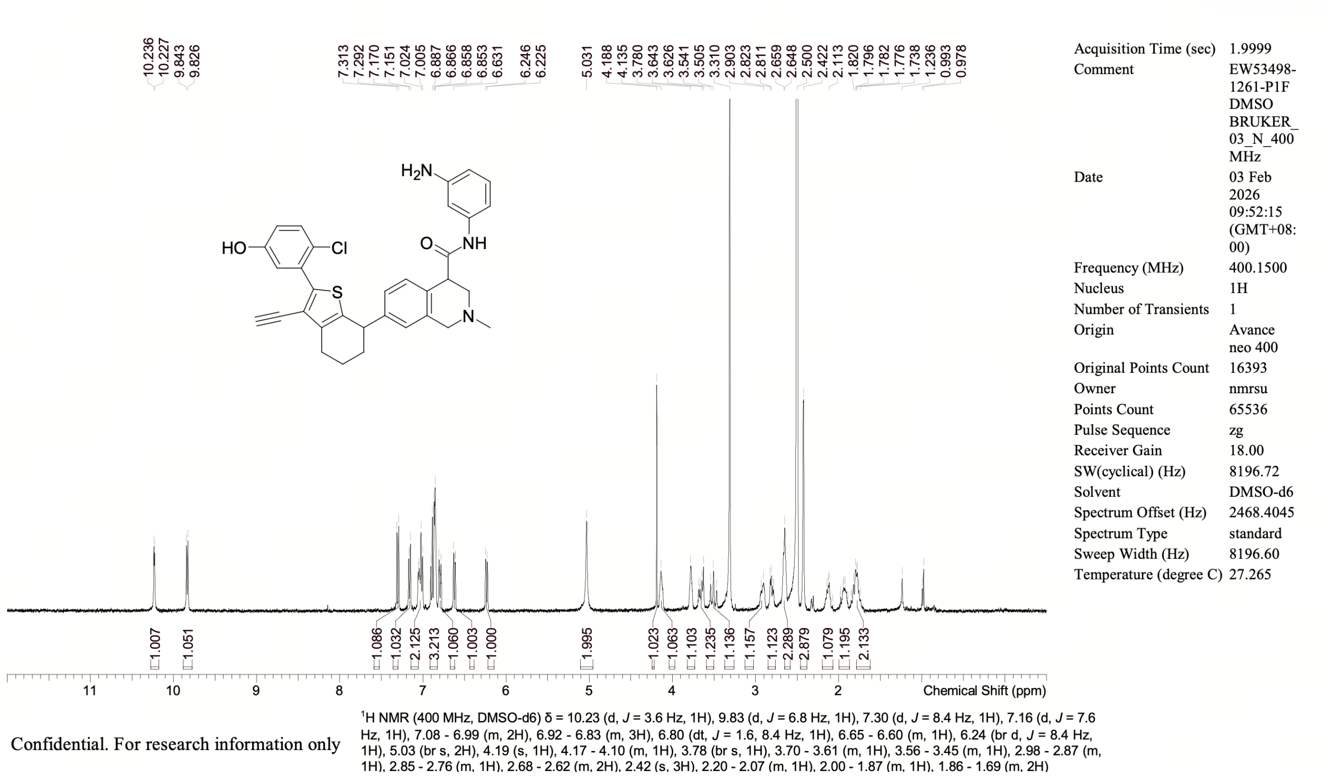

**NMR Spectrum of KRAS G12D-compound-3**

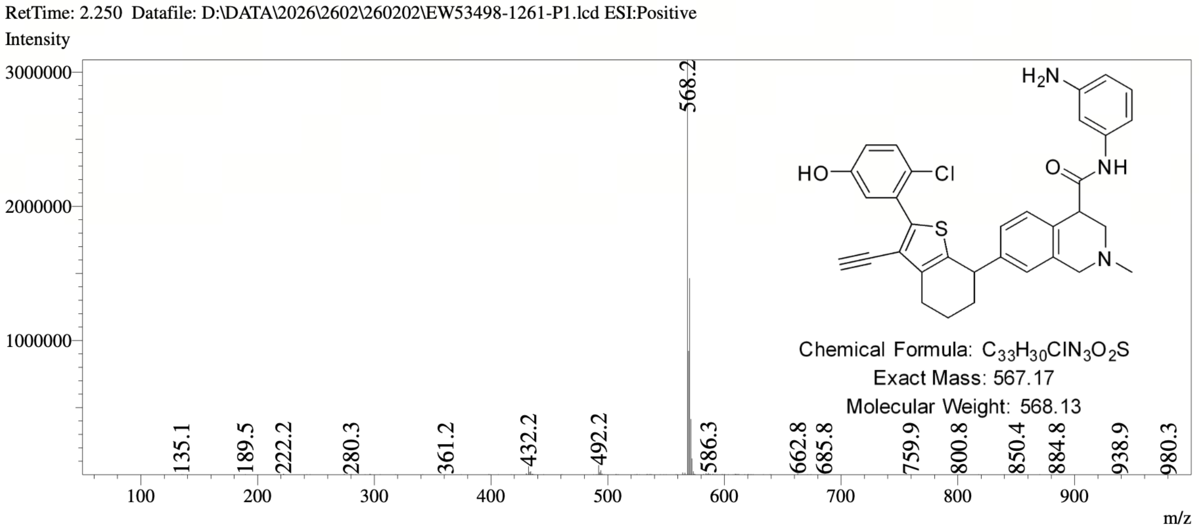

**MS Spectrum of KRAS G12D-compound-3**

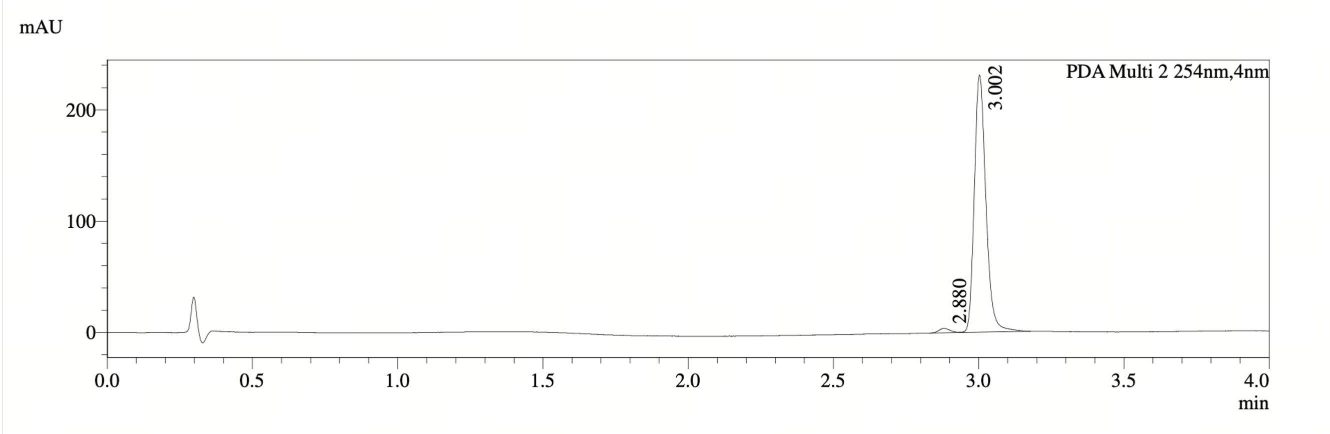

PDA Ch2 254nm

| Peak# | Ret. Time | Width | Height | Height% | Area | Area% |
|---|---|---|---|---|---|---|
| 1 | 2.880 | 0.072 | 4006 | 1.704 | 10683 | 1.639 |
| 2 | 3.002 | 0.072 | 231058 | 98.296 | 640943 | 98.361 |

**HPLC Chromatogram of KRAS G12D-compound-3**

#### 7.2 Designs for PCSK9

##### *7.2.1 PCSK9-compound-1*

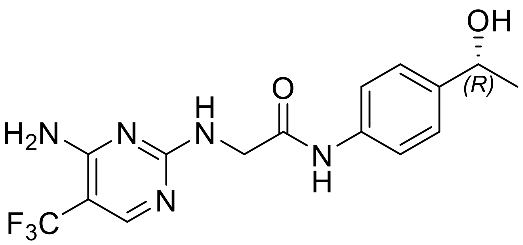

**PCSK9-compound-1**. (R)-2-((4-amino-5-(trifluoromethyl)pyrimidin-2-yl)amino)-N-(4-(1-hydroxyethyl)phenyl)acetamide

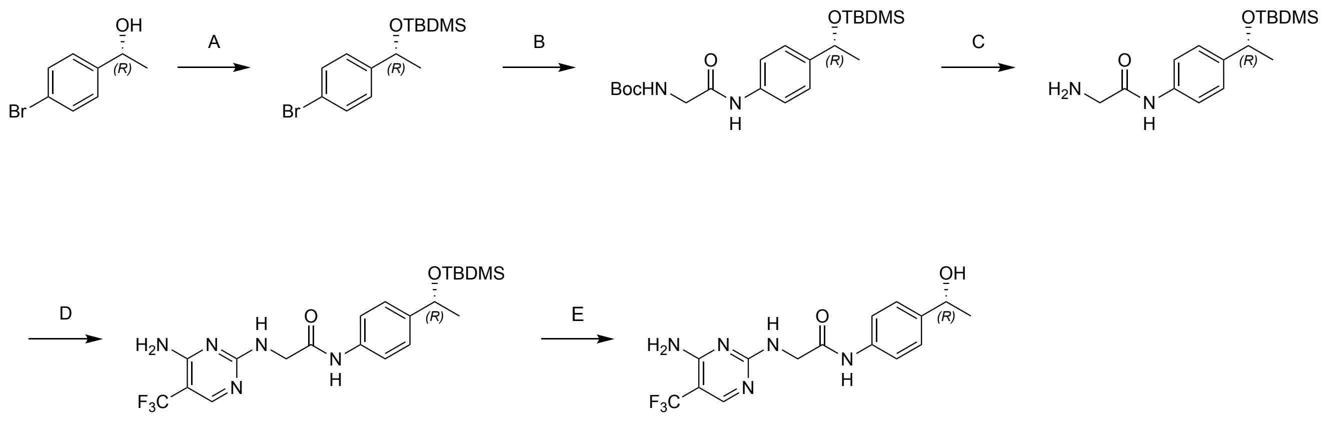

Reagents and conditions: A. TBDMSOTf, 2,6- Lutidine , DCM, 25 °C, 80%; B. Xantphos Pd $G_3$, $Cs_2CO_3$, dioxane, 100 °C, 38%; C. TFA, DCM, 20 °C, 89%; D. 2-chloro-5-(trifluoromethyl)pyrimidin-4-amine, DIEA, THF, 50 °C, 53%; E. TBAF, THF, 0 to 20 °C, 19%.

#### Synthesis of PCSK9-compound-1

Step A: (R)-(1-(4-bromophenyl)ethoxy)(tert-butyl)dimethylsilane: To a solution of (R)-1-(4-bromophenyl)ethan-1-ol (1.50 g, 7.46 mmol, 1.0 equiv) in dichloromethane (15.0 mL) was added 2,6-dimethylpyridine (1.60 g, 14.9 mmol, 1.74 mL, 2.0 equiv) and then dropwise added tert-butyldimethylsilyl trifluoromethanesulfonate (2.96 g, 11.2 mmol, 2.57 mL, 1.5 equiv) in dichloromethane (10.0 mL). The reaction was stirred at 25 °C for 1 hour. After completion of the reaction, the reaction mixture was quenched with water (30.0 mL) and extracted with dichloromethane (30.0 mL × 2). The combined organic layers were washed with brine (30.0 mL), dried over anhydrous sodium sulfate, filtered and concentrated under reduced pressure. The residue was purified by flash silica gel chromatography (ISCO®; 20.0 g SepaFlash® Silica Flash Column, Eluent of 0~2% Ethyl acetate/Commercial hexanes gradient @ 20 mL/min) to afford (R)-(1-(4-bromophenyl)ethoxy)(tert-butyl)dimethylsilane (1.90 g, 80.8% yield) as a colorless oil; LCMS-Method 1 $[M-131]^+ = 184.9$.

Step B: tert-butyl(R)-(2-((4-(1-((tert-butyldimethylsilyl)oxy)ethyl)phenyl)amino)-2-oxoethyl)carbamate: To a solution of (R)-(1-(4-bromophenyl)ethoxy)(tert-butyl)dimethylsilane (1.20 g, 3.81 mmol, 1.0 equiv) in dioxane (10.0 mL) were added tert-butyl N-(2-amino-2-oxo-ethyl)carbamate (1.19 g, 6.85 mmol, 1.8 equiv), cesium carbonate (1.61 g, 4.95 mmol, 1.3 equiv) and Methanesulfonato[9,9-dimethyl-4,5-bis(diphenylphosphino)xanthene][2-amino-1,1-biphenyl]palladium(II) (108 mg, 114 $\mu$mol, 0.03 equiv). The reaction was stirred at 100 °C for 12 hours under nitrogen. After completion of the reaction, the reaction mixture was quenched with water (10.0 mL) and extracted with ethyl acetate (20.0 mL × 2). The combined organic layers were washed with brine (20 mL), dried over anhydrous sodium sulfate, filtered and concentrated under reduced pressure to give a residue. The residue was purified by flash silica gel chromatography (ISCO®; 12.0 g SepaFlash® Silica Flash Column, Eluent of 0~50% Ethyl acetate/Commercial hexanesgradient @ 12 mL/min) to afford tert-butyl (R)-(2-((4-(1-((tert-butyldimethylsilyl)oxy)ethyl)phenyl)amino)-2-oxoethyl)carbamate (600 mg, 38.6% yield) as a brown oil; LCMS-Method 1 $[M+23]^+ = 431.2$.

Step C: (R)-2-amino-N-(4-(1-((tert-butyldimethylsilyl)oxy)ethyl)phenyl)acetamide:Toasolutionoftert-butyl(R)-(2-((4-(1-((tert-butyldimethylsilyl)oxy)ethyl)phenyl)amino)-2-oxoethyl)carbamate (400 mg, 979 $\mu$mol, 1.0 equiv) in dichloromethane (8.0 mL) were added trimethylsilyl trifluoromethanesulfonate (870 mg, 3.92 mmol, 707 $\mu$L, 4.0 equiv) and 2,6-dimethylpyridine (524 mg, 4.89 mmol, 570 $\mu$L, 5.0 equiv) at 0 °C. The reaction was stirred at 20 °C for 30 mins. After completion of the reaction, the reaction mixture quenched with ammonium chloride solution (15.0 mL) and extracted with ethyl acetate (15.0 mL× 2). The combined organic layers were washed with sodium bicarbonate (15.0 mL), dried over anhydrous sodium sulfate, filtered and concentrated under reduced pressure to give a residue. The residue was purified by flash silica gel chromatography (ISCO®; 12.0 g SepaFlash® Silica Flash Column, Eluent of 0~30% THF/hexanes gradient @ 20 mL/min) to afford (R)-2-amino-N-(4-(1-((tert-butyldimethylsilyl)oxy)ethyl)phenyl)acetamide (270 mg, 89.4% yield) as a white solid; LCMS-Method 1 $[M+1]^+$ = 309.2.

Step D: (R)-2-((4-amino-5-(trifluoromethyl)pyrimidin-2-yl)amino)-N-(4-(1-((tert-butyldimethylsilyl)oxy)ethyl)phenyl)acetamide:Toasolutionof(R)-2-amino-N-(4-(1-((tert-butyldimethylsilyl)oxy)ethyl)phenyl)acetamide(270mg,875μmol,1.0equiv)intetrahydrofuran (5.0 mL) were added Diisopropylethylamine (1.00 g, 7.74 mmol, 1.35 mL, 8.84 equiv) and 2-chloro-5-(trifluoromethyl)pyrimidin-4-amine (173 mg, 875 μmol, 1.0 equiv). The reaction was stirred at 50 °C for 12 hours. After completion of the reaction, the reaction mixture was concentrated under reduced pressure to remove tetrahydrofuran to give a residue. The residue was triturated with dichloromethane (5.0 mL) at 20 °C for 1 hours, filtered and the filter cake was dried under reduced pressure to afford (R)-2-((4-amino-5-(trifluoromethyl)pyrimidin-2-yl)amino)-N-(4-(1-((tert-butyldimethylsilyl)oxy)ethyl)phenyl)acetamide (220 mg, 53.5% yield) as a white solid; LCMS-Method 1 $[M+1]^+$ = 470.2.

Step E. PCSK-compound-1: To a solution of (R)-2-((4-amino-5-(trifluoromethyl)pyrimidin-2-yl)amino)-N-(4-(1-((tert-butyldimethylsilyl)oxy)ethyl)phenyl)acetamide (40.0 mg, 85.2 μmol, 1.0 equiv) in tetrahydrofuran (1.00 mL) was added tetrabutylammonium fluoride (1 M, 2.00 mL, 23.5 equiv) at 0 °C, the reaction was stirred at 20 °C for 2 hours. After completion of the reaction, the reaction mixture was quenched with sodium bicarbonate solution (5.00 mL) and extracted with ethyl acetate (10.0 mL × 2). The combined organic layers were washed with brine (10.0 mL), dried over anhydrous sodium sulfate, filtered and concentrated under reduced pressure to give a residue. The residue was purified by prep-HPLC (neutral condition; column: Waters Xbridge C18 150*25mm*5μm; mobile phase: [$H_2O$(10mM $NH_4HCO_3$)-ACN]; gradient: 10%-40% B over 10.0 min) to afford (R)-2-((4-amino-5-(trifluoromethyl)pyrimidin-2-yl)amino)-N-(4-(1-hydroxyethyl)phenyl)acetamide (6.00 mg, 19.7% yield) as a white solid; $^1$H NMR (400 MHz, MeOD) $\delta$ = 8.06 (s, 1H), 7.51 (d, $J$ = 8.6 Hz, 2H), 7.31 (d, $J$ = 8.6 Hz, 2H), 4.81 - 4.76 (m, 1H), 4.14 (s, 2H), 1.42 (d, $J$ = 6.4 Hz, 3H), LCMS $[M+1]^+$ = 356.1; HPLC Rt = 1.477 min.

**Analytical method by SFC:**

Column: Chiralcel OJ-3 50*4.6mm I.D., 3 μm;
Mobile phase: Phase A for $CO_2$, and Phase B for IPA (0.05%DEA);
Gradient elution: IPA (0.05%DEA) in $CO_2$ from 5% to 40%;
Flow rate:4 mL/min; Detector: PDA;
Colum Temp: 35 °C; Back Pressure: 100 Bar;
Retention time: 1.152 min.

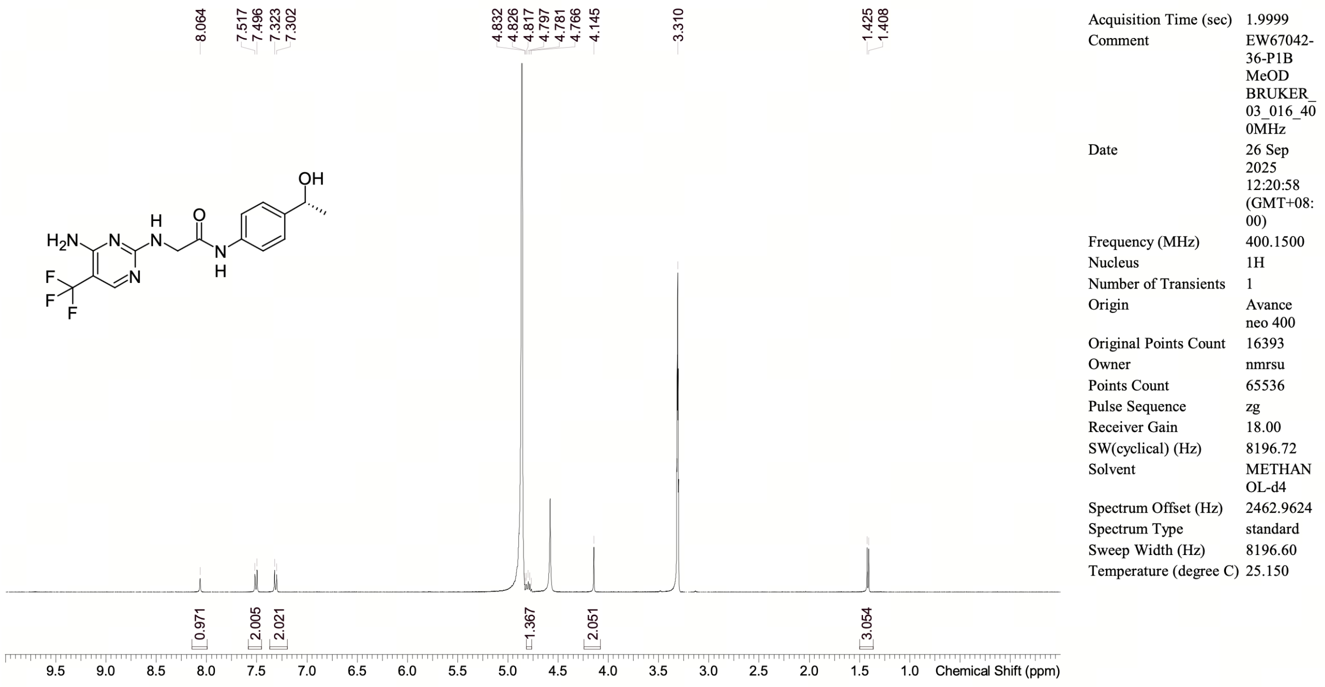

Confidential. For research information only

[1]H NMR (400 MHz, CD3OD) $\delta$ (ppm) = 8.06 (s, 1H), 7.51 (d, $J$ = 8.6 Hz, 2H), 7.31 (d, $J$ = 8.6 Hz, 2H), 4.81 - 4.76 (m, 1H), 4.14 (s, 2H), 1.42 (d, $J$ = 6.4 Hz, 3H)

Operator: Date:

**NMR Spectrum of PCSK9-compound-1**

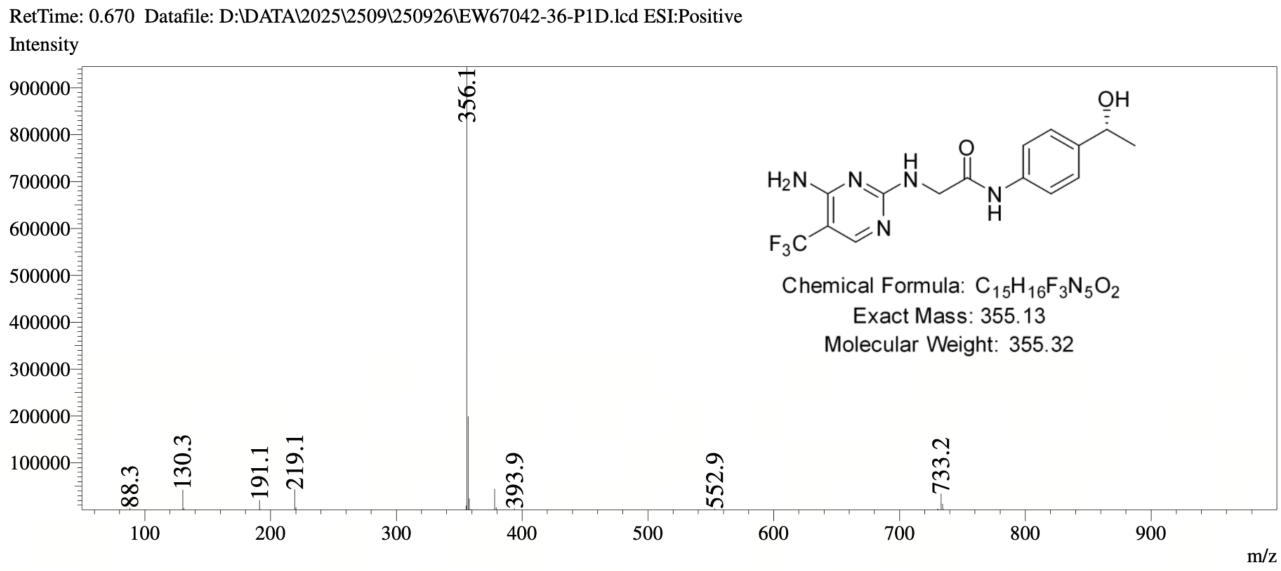

**MS Spectrum of PCSK9-compound-1**

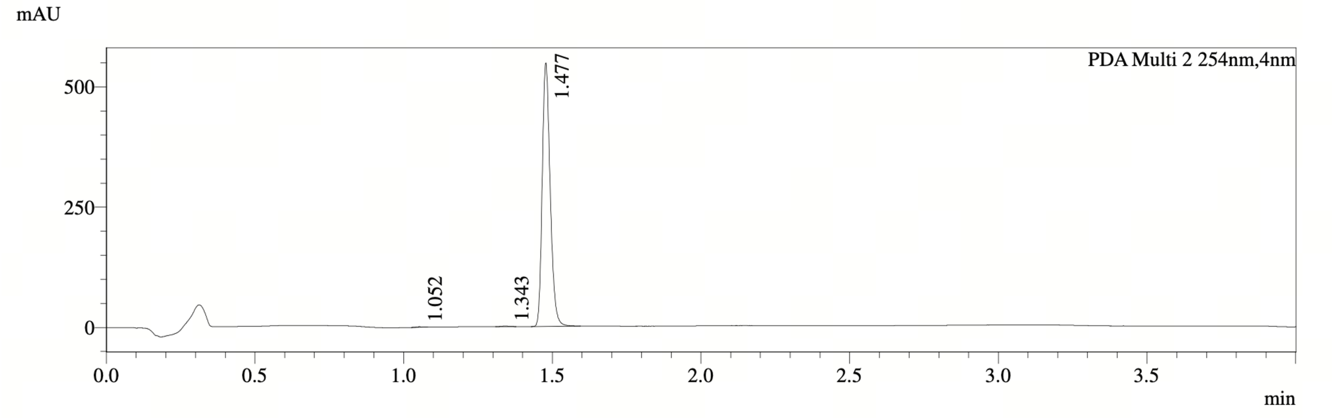

PDA Ch2 254nm

| Peak# | Ret. Time | Width | Height | Height% | Area | Area% |
|---|---|---|---|---|---|---|
| 1 | 1.052 | 0.043 | 957 | 0.174 | 1498 | 0.146 |
| 2 | 1.343 | 0.053 | 985 | 0.179 | 1929 | 0.188 |
| 3 | 1.477 | 0.050 | 547442 | 99.646 | 1020352 | 99.665 |

**HPLC Chromatogram of PCSK9-compound-1**

##### *7.2.2 PCSK9-compound-2*

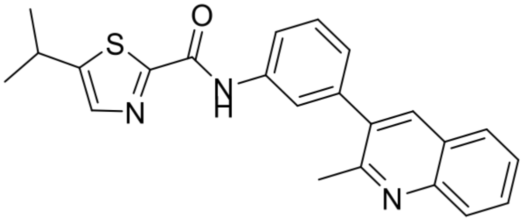

**PCSK9-compound-2**. 5-isopropyl-N-(3-(2-methylquinolin-3-yl)phenyl)thiazole-2-carboxamide

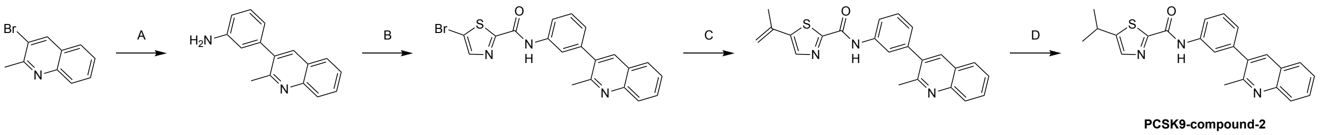

Reagents and conditions: (A) (3-aminophenyl)boronic acid, Pd(dppf)$Cl_2$ $CH_2Cl_2$, $K_2CO_3$, DMTF, $H_2O$, 90 °C; (B) 5-bromothiazole-2-carboxylic acid, TATU, TEA, THF, 25 °C; (C) Isopropenylboronic acid pinacol ester, Pd(dppf)$Cl_2$.$CH_2Cl_2$, $K_2CO_3$, ACN, $H_2O$, 80 °C, 73.4%; (D) $PtO_2$, $H_2$(15 Psi), THF, 25 °C, 41.5%.

**Synthesis of PCSK9-compound-2**

Step A: 3-(2-methylquinolin-3-yl)aniline:Toasolutionof3-bromo-2-methylquinoline(200mg,901$\mu$mol,1.00equiv)and(3-aminophenyl)boronicacid (250 mg, 1.44 mmol, 1.60 equiv) in 2-methyltetrahydrofuran (6.0 mL) were added a solution of potassium carbonate (249 mg, 1.80 mmol, 2.00 equiv) in water (1.50 mL) and Dichloro[1,1-bis(diphenylphosphino)ferrocene]palladium(II) (52.0 mg, 45.0 $\mu$mol, 0.05 equiv). The mixture was stirred at 90 °C for 12 hours. After completion of the reaction, the mixture was quenched with water (10.0 mL) and extracted with ethyl acetate (15.0 mL ×2). The combined organic layers were washed with brine (10 mL), dried over anhydrous sodium sulfate, filtered and concentrated under reduced pressure to give a residue. The residue was purified by flash silica gel chromatography (ISCO®; 12.0 g SepaFlash® Silica Flash Column, Eluent of 5∼30% Ethyl acetate/Commercial hexanes gradient at 15 mL/min) to afford 3-(2-methylquinolin-3-yl)aniline (300 mg, crude) as a brown oil; LCMS-Method 1 $[M+1]^+ = 235.1$.

Step B: 5-bromo-N-(3-(2-methylquinolin-3-yl)phenyl)thiazole-2-carboxamide: To a solution of 3-(2-methylquinolin-3-yl)aniline (120 mg, 512 $\mu$mol, 1.00 equiv) and 5-bromothiazole-2-carboxylic acid (120 mg, 579 $\mu$mol, 1.13 equiv) in tetrahydrofuran (2.00 mL) were added triethylamine (77.7 mg, 768 $\mu$mol, 106 $\mu$L, 1.50 equiv) and 2-(7-Azabenzotriazole-1-yl)-1,1,3,3-tetramethyluronium tetrafluoroborate (195 mg, 512 $\mu$mol, 1.00 equiv). The mixture was stirred at 25 °C for 12 hours. After completion of the reaction, the mixture was quenched with methanol (1.00 mL) and water (10 mL), and extracted with ethyl acetate (15.0 mL). The combined organic layers were washed with brine (10 mL), dried over anhydrous sodium sulfate, filtered and concentrated under reduced pressure to give a residue. The residue was purified by prep-HPLC (TFA condition; column: Phenomenex Luna C18 150*25mm*10$\mu$m; mobile phase: [$H_2O$ (0.1% TFA)-ACN]; gradient: 18%-48% B over 13.0 min) to afford 5-bromo-N-(3-(2-methylquinolin-3-yl)phenyl)thiazole-2-carboxamide (220 mg, crude) as a colorless oil; LCMS-Method 1 $[M+1, M+3]^+ = 424.0, 426.0$.

Step C: N-(3-(2-methylquinolin-3-yl)phenyl)-5-(prop-1-en-2-yl)thiazole-2-carboxamide: To a solution of 5-bromo-N-(3-(2-methylquinolin-3-yl)phenyl)thiazole-2-carboxamide (150 mg, 354 $\mu$mol, 1.00 equiv) and 2-isopropenyl-4,4,5,5-tetramethyl-1,3,2-dioxaborolane (89.1 mg, 530 $\mu$mol, 1.50 equiv) in acetonitrile (6.00 mL) and water (0.60 mL) were added potassium carbonate (97.7 mg, 707 $\mu$mol, 2.00 equiv) and Dichloro[1,1-bis(diphenylphosphino)ferrocene]palladium(II) (25.9 mg, 35.4 $\mu$mol, 0.10 equiv). The reaction mixture was stirred at 80 °C for 12 hours under nitrogen. After completion of the reaction, the mixture was quenched with water (10 mL), and extracted with ethyl acetate (15.0 mL). The combined organic layers were washed with brine (10 mL), dried over anhydrous sodium sulfate, filtered and concentrated under reduced pressure to give a residue. The residue was purified by flash silica gel chromatography (ISCO®; 12.0 g SepaFlash® Silica Flash Column, Eluent of 5~50% Ethyl acetate/Commercial hexanes gradient @ 20 mL/min) to afford N-(3-(2-methylquinolin-3-yl)phenyl)-5-(prop-1-en-2-yl)thiazole-2-carboxamide (100 mg, 73.4% yield) as a brown solid; LCMS-Method 25 $[M+1]^+ = 386.2$.

Step D: PCSK9-compound-2: To a solution of N-(3-(2-methylquinolin-3-yl)phenyl)-5-(prop-1-en-2-yl)thiazole-2-carboxamide (70.0 mg, 182 $\mu$mol, 1.00 equiv) in tetrahydrofuran (10.0 mL) was added platinum dioxide (4.12 mg, 18.2 $\mu$mol, 0.10 eq) under nitrogen. The reaction mixture was stirred under hydrogen (15 psi) at 25 °C for 36 hours. After completion of the reaction, the reaction mixture was filtered and the filter was concentrated under reduced pressure to give a residue. The residue was purified by prep-HPLC (column: Phenomenex Luna C18 150*25mm*10$\mu$m; mobile phase: [$H_2O$ (0.1% TFA)-ACN]; gradient: 20%-50% B over 13.0 min) to afford 5-isopropyl-N-(3-(2-methylquinolin-3-yl)phenyl)thiazole-2-carboxamide (30.0 mg, 41.5% yield) as a brown solid; $^1H$ NMR (400 MHz, MeOD) $\delta$ = 8.95 (s, 1H), 8.35 - 8.27 (m, 1H), 8.24 - 8.17 (m, 1H), 8.16 - 8.09 (m, 1H), 8.09 - 8.03 (m, 1H), 7.97 - 7.89 (m, 1H), 7.88 - 7.82 (m, 1H), 7.80 - 7.74 (m, 1H), 7.61 (t, $J$ = 7.8 Hz, 1H), 7.37 (d, $J$ = 7.6 Hz, 1H), 3.39 - 3.35 (m, 1H), 2.94 (s, 3H), 1.41 (d, $J$ = 7.0 Hz, 6H); LCMS $[M+1]^+ = 388.1$; HPLC Rt = 2.317 min.

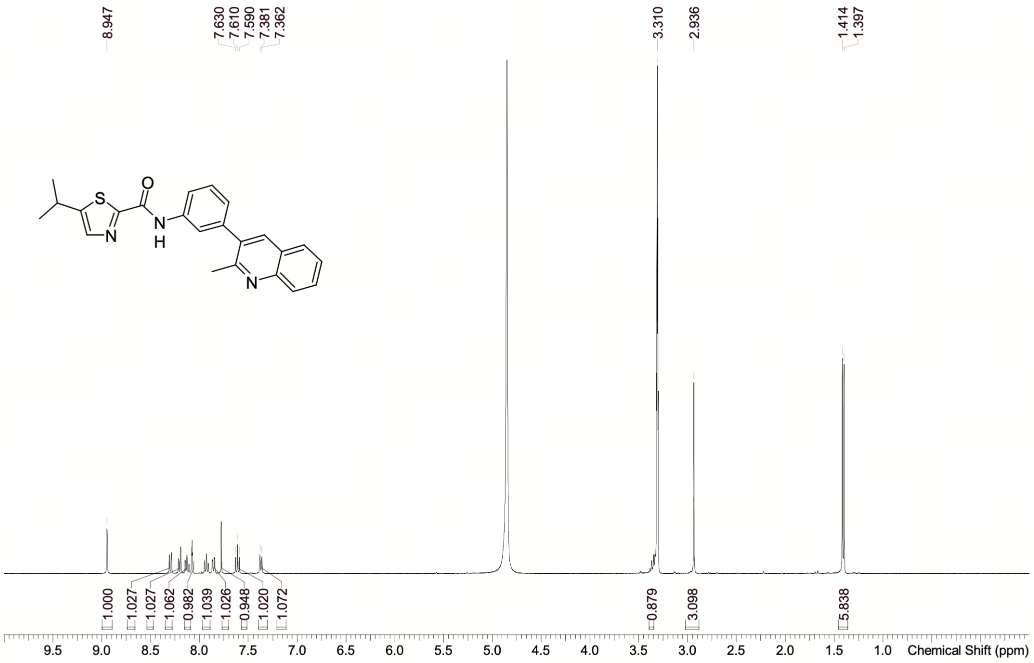

| | |
|---|---|
| Acquisition Time (sec) | 3.9977 |
| Comment | EW67042-44-P1C MeOD BRUKER_03_016_400MHz |
| Date | 20 Oct 2025 10:53:00 (GMT+08:00) |
| Frequency (MHz) | 400.1500 |
| Nucleus | 1H |
| Number of Transients | 8 |
| Origin | Avance neo 400 |
| Original Points Count | 32768 |
| Owner | nmrsu |
| Points Count | 131072 |
| Pulse Sequence | zg30 |
| Receiver Gain | 101.00 |
| SW(cyclical) (Hz) | 8196.72 |
| Solvent | METHANOL-d4 |
| Spectrum Offset (Hz) | 2393.0164 |
| Spectrum Type | standard |
| Sweep Width (Hz) | 8196.66 |
| Temperature (degree C) | 25.176 |

Confidential. For research information only

$^{1}$H NMR (400 MHz, CD3OD) $\delta$ (ppm) = 8.95 (s, 1H), 8.35 - 8.27 (m, 1H), 8.24 - 8.17 (m, 1H), 8.16 - 8.09 (m, 1H), 8.09 - 8.03 (m, 1H), 7.97 - 7.89 (m, 1H), 7.88 - 7.82 (m, 1H), 7.80 - 7.74 (m, 1H), 7.61 (t, $J$ = 7.6 Hz, 1H), 7.37 (d, $J$ = 7.6 Hz, 1H), 3.39 - 3.35 (m, 1H), 2.94 (s, 3H), 1.41 (d, $J$ = 6.8 Hz, 6H)

Operator: Date:

#### NMR Spectrum of PCSK9-compound-2

RetTime: 2.020 Datafile: D:\DATA\2025\2510\251020\EW67042-44-P1F2.lcd

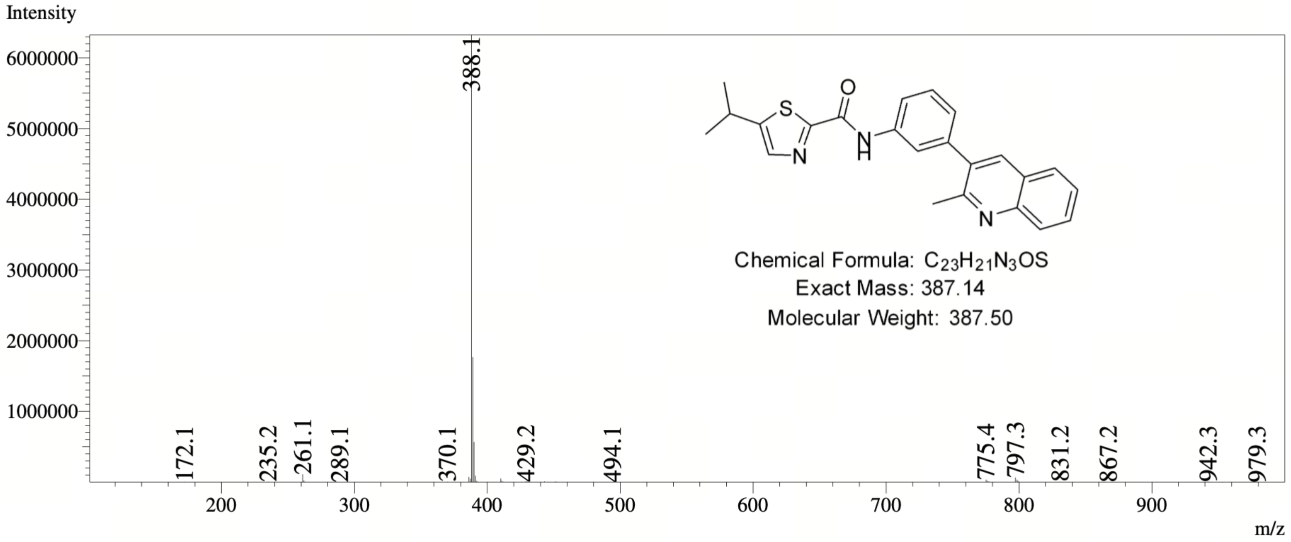

#### MS Spectrum of PCSK9-compound-2

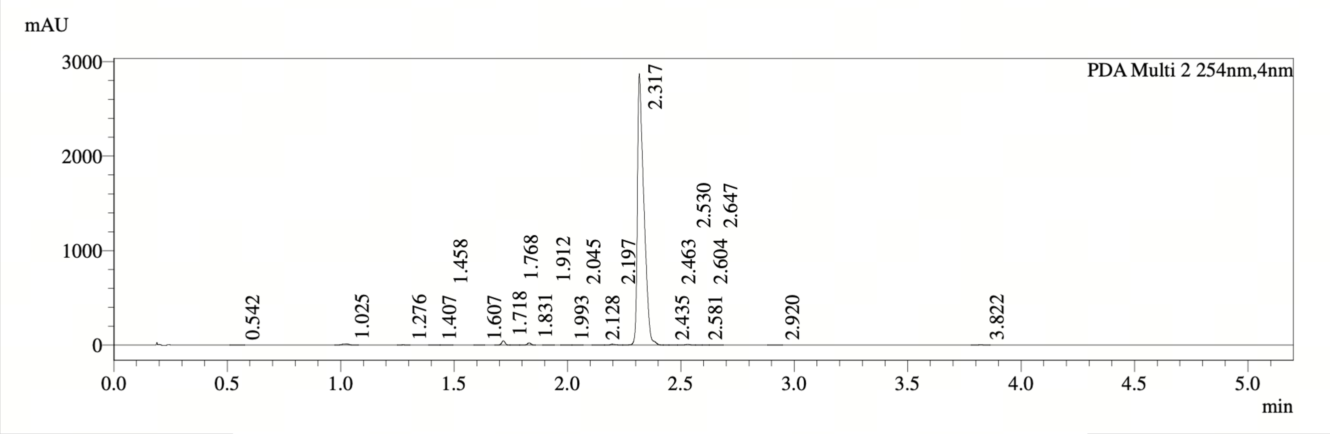

PDA Ch2 254nm

| Peak# | Ret. Time | Width | Height | Height% | Area | Area% |
|---|---|---|---|---|---|---|
| 1 | 0.542 | 0.046 | 690 | 0.023 | 1222 | 0.021 |
| 2 | 1.025 | 0.052 | 11455 | 0.383 | 23416 | 0.402 |
| 3 | 1.276 | 0.030 | 2557 | 0.086 | 2859 | 0.049 |
| 4 | 1.407 | 0.029 | 1507 | 0.050 | 1641 | 0.028 |
| 5 | 1.458 | 0.026 | 1418 | 0.047 | 1434 | 0.025 |
| 6 | 1.607 | 0.027 | 1827 | 0.061 | 1853 | 0.032 |
| 7 | 1.718 | 0.027 | 46080 | 1.543 | 48089 | 0.825 |
| 8 | 1.768 | 0.037 | 2283 | 0.076 | 3247 | 0.056 |
| 9 | 1.831 | 0.028 | 23916 | 0.801 | 27742 | 0.476 |
| 10 | 1.912 | 0.028 | 1768 | 0.059 | 1900 | 0.033 |
| 11 | 1.993 | 0.030 | 1207 | 0.040 | 1493 | 0.026 |
| 12 | 2.045 | 0.038 | 713 | 0.024 | 1030 | 0.018 |
| 13 | 2.128 | 0.051 | 1519 | 0.051 | 2960 | 0.051 |
| 14 | 2.197 | 0.037 | 9848 | 0.330 | 16176 | 0.278 |
| 15 | 2.317 | 0.055 | 2858516 | 95.696 | 5659447 | 97.095 |
| 16 | 2.435 | 0.129 | 1743 | 0.058 | 2355 | 0.040 |
| 17 | 2.463 | 0.040 | 2814 | 0.094 | 4201 | 0.072 |
| 18 | 2.530 | 0.037 | 7617 | 0.255 | 12663 | 0.217 |
| 19 | 2.581 | 0.159 | 1442 | 0.048 | 1940 | 0.033 |
| 20 | 2.604 | 0.041 | 1691 | 0.057 | 2085 | 0.036 |
| 21 | 2.647 | 0.046 | 1293 | 0.043 | 2299 | 0.039 |
| 22 | 2.920 | 0.036 | 2064 | 0.069 | 2941 | 0.050 |
| 23 | 3.822 | 0.050 | 3118 | 0.104 | 5809 | 0.100 |

**HPLC Chromatogram of PCSK9-compound-2**

##### *7.2.3 PCSK9-compound-3*

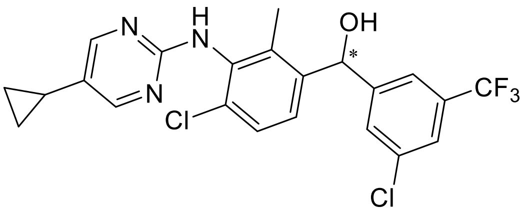

**PCSK9-compound-3**. (4-chloro-3-((5-cyclopropylpyrimidin-2-yl)amino)-2-methylphenyl)(3-chloro-5-(trifluoromethyl)phenyl)methanol

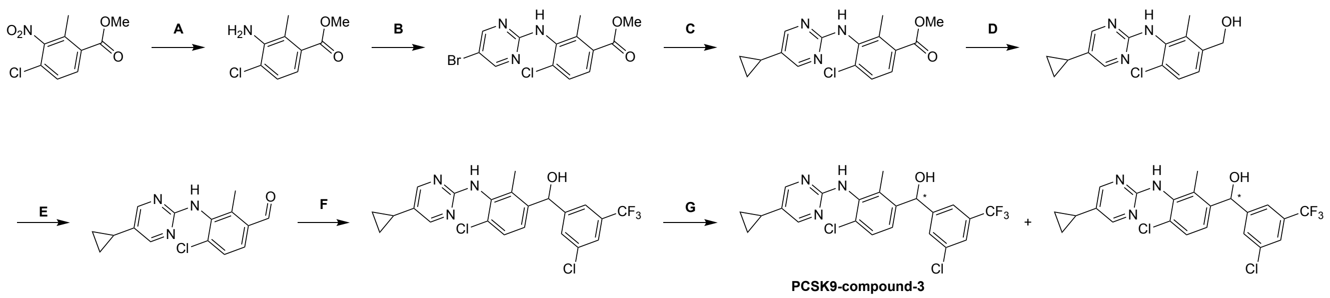

Reagents and conditions: (A) 1%Pt/C, $H_2$ (1.0 Mpa), THF, EtOAc, 45 °C, 92%; (B) NaHMDS, 5-bromo-2-methylsulfonyl-pyrimidine, THF, 0 – 25 °C; (C) cataCXium® A Pd $G_3$, cyclopropylboronic acid, $K_3PO_4$, toluene., $H_2O$; 90 °C, 42.6%; (D) DIBAL-H, DCM, 0 °C, 95.9%; (E) $MnO_2$, DCM, 95.9%; (F) *n*-BuLi, 4-chloro-3-((5-cyclopropylpyrimidin-2-yl)amino)-2-methylbenzaldehyde, THF, -50 °C; (G) chiral SFC.

#### Synthesis of PCSK-compound-3

Step A: methyl 3-amino-4-chloro-2-methylbenzoate: Solution 1: methyl 4-chloro-2-methyl-3-nitro-benzoate (5.00 g, 21.8 mmol, 1.0 equiv) in tetrahydrofuran (50.0 mL) and ethyl acetate (50.0 mL). The fixed bed (named FLR1, volume 5 mL) was completely packed with granular catalyst {WXSC1050(1%Pt/C), 2.53 g}. The hydrogen back pressure regulator was adjusted to 1.0 MPa, and the hydrogen flow rate was {9.0 equiv, 16.6 sccm}. The solution 1 was pumped by Pump 1 {S1, P1, 0.4 mL/min} to flow reactor 1 {FLR1, SS, Fixed bed, 6.350(1/4") mm, 5.00 mL, 45.0 °C}. The reaction mixture was continuously collected from the reactor outlet into the container, the mixture was collected with a bottle. After completion of the reaction, the reaction mixture was filtered and the filter liquor was concentrated under reduced pressure to afford methyl 3-amino-4-chloro-2-methyl-benzoate (4.00 g, 92.0% yield) as yellow solid.

Step B: methyl3-((5-bromopyrimidin-2-yl)amino)-4-chloro-2-methylbenzoate: To a solution of methyl 3-amino-4-chloro-2-methyl-benzoate (2.30 g, 11.5 mmol, 1.0 equiv) in tetrahydrofuran (20.0 mL) was added sodium bis(trimethylsilyl)amide (1 M, 23.0 mL, 2.0 equiv) at -70 °C, the mixture was stirred at 0 °C for 1 hour under nitrogen. And then a solution of 5-bromo-2-methylsulfonyl-pyrimidine (2.73 g, 11.5 mmol, 1.0 equiv) in tetrahydrofuran (5.0 mL) was added dropwise at 0 °C. The reaction was stirred at 25 °C for 2 hours under nitrogen. After completion of the reaction, the mixture was quenched with ammonium chloride solution (20.0 mL) and extracted with ethyl acetate (20.0 mL × 3). The combined organic layers were washed with brine (30 mL), dried over anhydrous sodium sulfate, filtered and concentrated under reduced pressure to give a residue. The residue was purified by column chromatography (silicon dioxide, Commercial hexanes: Ethyl acetate = 50/1 to 10/1) to afford methyl 3-((5-bromopyrimidin-2-yl)amino)-4-chloro-2-methylbenzoate (500 mg, crude) as yellow solid; LCMS-Method 1 $[M+1]^+$ = 357.8.

Step C: methyl4-chloro-3-((5-cyclopropylpyrimidin-2-yl)amino)-2-methylbenzoate: To a solution of methyl 3-((5-bromopyrimidin-2-yl)amino)-4-chloro-2-methylbenzoate (150 mg, 420 $\mu$mol, 1.0 equiv) in toluene (1.00 mL) and water (100 $\mu$L) were added cyclopropylboronic acid (72.3 mg, 841 $\mu$mol, 2.0 equiv), potassium phosphate (267 mg, 1.26 mmol, 3.0 equiv) and Methanesulfonato(diadamantyl-n-butylphosphino)-2-amino-1,1-biphenyl-2-yl)palladium(II) (30.6 mg, 42.1 $\mu$mol, 0.10 equiv). The reaction was stirred at 90 °C for 1 hour under nitrogen. After completion of the reaction, the mixture was quenched with water (10.0 mL) and extracted with ethyl acetate (10.0 mL × 2). The combined organic layers were washed with brine (10 mL), dried over anhydrous sodium sulfate, filtered and concentrated under reduced pressure to give a residue. The residue was purified by column chromatography (silicon dioxide, Commercial hexanes: Ethyl acetate = 20/1 to 10/1) to afford methyl 4-chloro-3-[(5-cyclopropylpyrimidin-2-yl)amino]-2-methyl-benzoate (90.0 mg, 42.6% yield) as yellow solid; LCMS-Method 1 $[M+1]^+$ = 317.9.

Step D: (4-chloro-3-((5-cyclopropylpyrimidin-2-yl)amino)-2-methylphenyl)methanol: To a solution of methyl 4-chloro-3-[(5-cyclopropylpyrimidin-2-yl)amino]-2-methyl-benzoate (80.0 mg, 251 $\mu$mol, 1.0 equiv) in dichloromethane (1.00 mL) was added Diisobutylaluminium hydrid (1 M, 1.01 mL, 4.0 equiv) at 0 °C, the reaction was stirred at 0 °C for 2 hours under nitrogen. After completion of the reaction, the reaction mixture was quenched with sodium sulfate decahydrate (10.0 mL), filtered and extracted with ethyl acetate (30.0 mL × 3). The combined organic layers were washed with brine (30 mL × 3), dried over anhydrous sodium sulfate, filtered and concentrated under reduced pressure to afford (4-chloro-3-((5-cyclopropylpyrimidin-2-yl)amino)-2-methylphenyl)methanol (70.0 mg, 95.9% yield) as yellow solid; LCMS-Method 1 $[M+1]^+$ = 290.0.

Step E: 4-chloro-3-((5-cyclopropylpyrimidin-2-yl)amino)-2-methylbenzaldehyde: To a solution of (4-chloro-3-((5-cyclopropylpyrimidin-2-yl)amino)-2-methylphenyl)methanol (70.0 mg, 241 $\mu$mol, 1.0 equiv) in dichloromethane (0.10 mL) was added manganese dioxide (126 mg, 1.45 mmol, 6.0 equiv), the reaction was stirred at 25 °C for 4 hours under nitrogen. After completion of the reaction, the reaction mixture was filtered and the filter liquor was concentrated under reduced pressure to afford 4-chloro-3-((5-cyclopropylpyrimidin-2-yl)amino)-2-methylbenzaldehyde (65.0 mg, 93.5% yield) as yellow solid; LCMS-Method 1 $[M+1]^+$ = 288.0.

Step F: (4-chloro-3-((5-cyclopropylpyrimidin-2-yl)amino)-2-methylphenyl)(3-chloro-5-(trifluoromethyl)phenyl)methanol: To a solution of 1-bromo-3-chloro-5-(trifluoromethyl)benzene (175 mg, 678 $\mu$mol, 3.0 equiv) in tetrahydrofuran (1.00 mL) was added n-butyllithium (1.60 M, 423 $\mu$L, 3.0 equiv), the mixture was stirred at -50 °C for 15 minutes under nitrogen, then 4-chloro-3-((5-cyclopropylpyrimidin-2-yl)amino)-2-methylbenzaldehyde (65.0 mg, 225 $\mu$mol, 1.00 equiv) in tetrahydrofuran (1.00 mL) was added dropwise, the reaction was stirred at -50 °C for 15 minutes under nitrogen. After completion of the reaction, the reaction mixture was quenched with ammonium chloride solution (5 mL) and extracted with ethyl acetate (10.0 mL × 2). The combined organic layers were washed with brine (10 mL), dried over anhydrous sodium sulfate, filtered and concentrated under reduced pressure. The residue was purified by prep-HPLC (column: Waters Xbridge C18 150*25mm*5um;mobile phase: [$H_2O$ (0.05% $NH_3H_2O$)-ACN];gradient:45%-75% B over 12.0 min) to afford (4-chloro-3-((5-cyclopropylpyrimidin-2-yl)amino)-2-methylphenyl)(3-chloro-5-(trifluoromethyl)phenyl)methanol (20 mg) as yellow solid.

Step G: PCSK9-compound-3: (4-chloro-3-((5-cyclopropylpyrimidin-2-yl)amino)-2-methylphenyl)(3-chloro-5-(trifluoromethyl)phenyl)methanol (20 mg) was separated via SFC (column: DAICEL CHIRALPAK IG (250mm*30mm,10um);mobile phase: [$CO_2$-IPA(0.1% $NH_3H_2O$)]; B%:30%, isocratic elution mode) to afford the two isomers which without absolute configuration identification. Isomer 1 (178B): (4-chloro-3-((5-cyclopropylpyrimidin-2-yl)amino)-2-methylphenyl)(3-chloro-5-(trifluoromethyl)phenyl)methanol (Peak 1 on SFC, Rt = 3.023 min, single diastereomer of unknown absolute configuration) (3.40 mg) as yellow solid; $^1$H NMR (400 MHz, CHLOROFORM-d) $\delta$ = 8.14 (s, 2H), 7.58 - 7.49 (m, 3H), 7.39 (d, $J$ = 8.0 Hz, 1H), 7.24 (s, 1H), 6.60 (s,1H), 6.07 (s, 1H), 2.60 (s, 1H), 2.23 (s, 3H), 1.77 - 1.74 (m, 1H), 0.98 - 0.92 (m, 2H), 0.67 - 0.61 (m, 2H); $^{19}$F NMR (400 MHz, CHLOROFORM-d) $\delta$ = -62.70; LCMS $[M+1]^+$ = 468.2; HPLC Rt = 2.702 min.

**Analytical method by SFC:**

Column: Chiralpak AD-3 100×4.6mm I.D., 3 $\mu$m;

Mobile phase: Phase A for $CO_2$, and Phase B for EtOH (0.05% DEA);

Gradient elution: B in A from 5% to 40%;

Flow rate: 2.5 mL/min; Detector: PDA;

Colum Temp:35°C; Back Pressure: 100 Bar;

Retention time: 3.023 min.

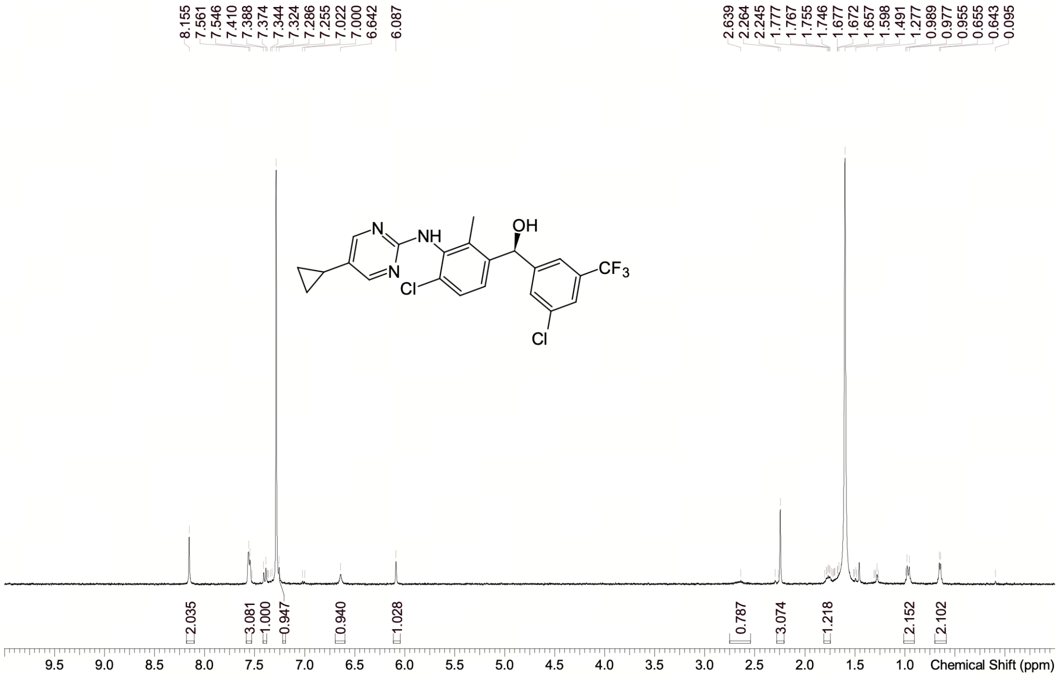

| | |
|---|---|
| Acquisition Time (sec) | 1.9999 |
| Comment | EW67290-30-P1A1 CDCl3 BRUKER_03_015_400MHz |
| Date | 26 Sep 2025 13:49:54 (GMT+08:00) |
| Frequency (MHz) | 400.1300 |
| Nucleus | 1H |
| Number of Transients | 1 |
| Origin | Avance neo 400 |
| Original Points Count | 16393 |
| Owner | nmrsu |
| Points Count | 65536 |
| Pulse Sequence | zg |
| Receiver Gain | 18.00 |
| SW(cyclical) (Hz) | 8196.72 |
| Solvent | CHLOROFORM-d |
| Spectrum Offset (Hz) | 2470.8015 |
| Spectrum Type | standard |
| Sweep Width (Hz) | 8196.60 |
| Temperature (degree C) | 23.347 |

Confidential. For research information only

$^{1}$H NMR (400 MHz, CHLOROFORM-$d$) δ 8.14 (s, 2 H), 7.49 - 7.58 (m, 3 H), 7.39 (d, J = 8.0 Hz, 1 H), 7.24 (s, 1 H), 6.60 (br s, 1 H), 6.07 (s, 1 H), 2.60 (br s, 1 H), 2.23 (s, 3 H), 1.74 - 1.77 (m, 1 H), 0.92 - 0.98 (m, 2 H), 0.61 - 0.67 (m, 2 H).

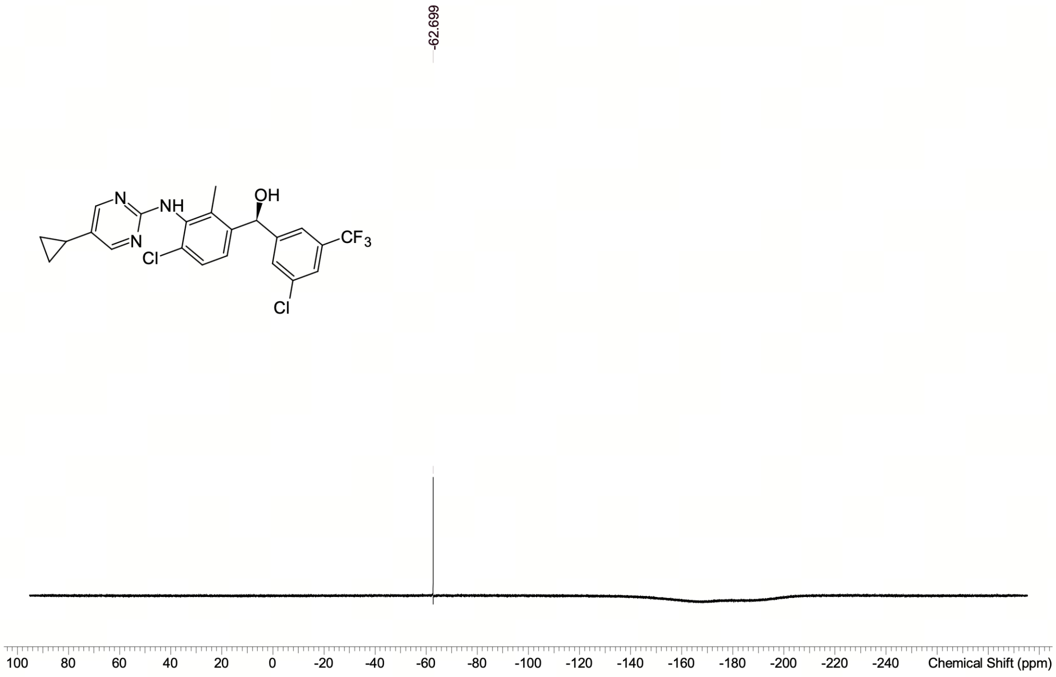

| | |
|---|---|
| Acquisition Time (sec) | 0.4456 |
| Comment | EW67290-30-P1A1 CDCl3 BRUKER_03_015_400MHz |
| Date | 26 Sep 2025 13:51:11 (GMT+08:00) |
| Frequency (MHz) | 376.4984 |
| Nucleus | 19F |
| Number of Transients | 8 |
| Origin | Avance neo 400 |
| Original Points Count | 65536 |
| Owner | nmrsu |
| Points Count | 65536 |
| Pulse Sequence | zgig |
| Receiver Gain | 101.00 |
| SW(cyclical) (Hz) | 147058.83 |
| Solvent | CHLOROFORM-d |
| Spectrum Offset (Hz) | -37649.8398 |
| Spectrum Type | standard |
| Sweep Width (Hz) | 147056.58 |
| Temperature (degree C) | 23.405 |

Confidential. For research information only

Operator: Date:

**NMR Spectra of PCSK9-compound-3**

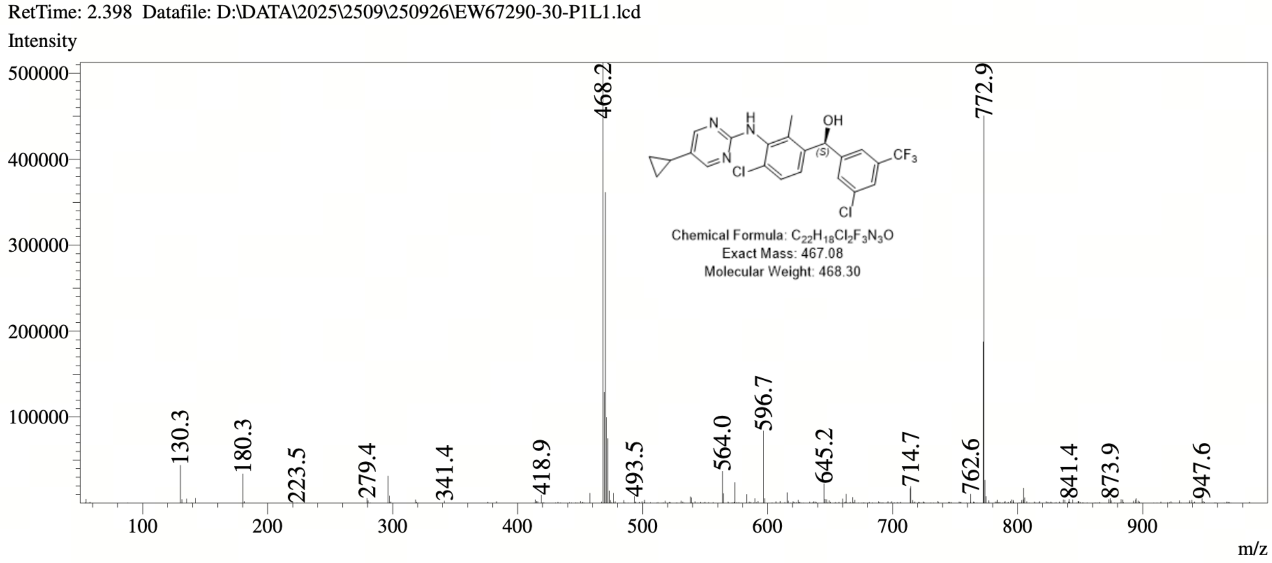

**MS Spectrum of PCSK9-compound-3**

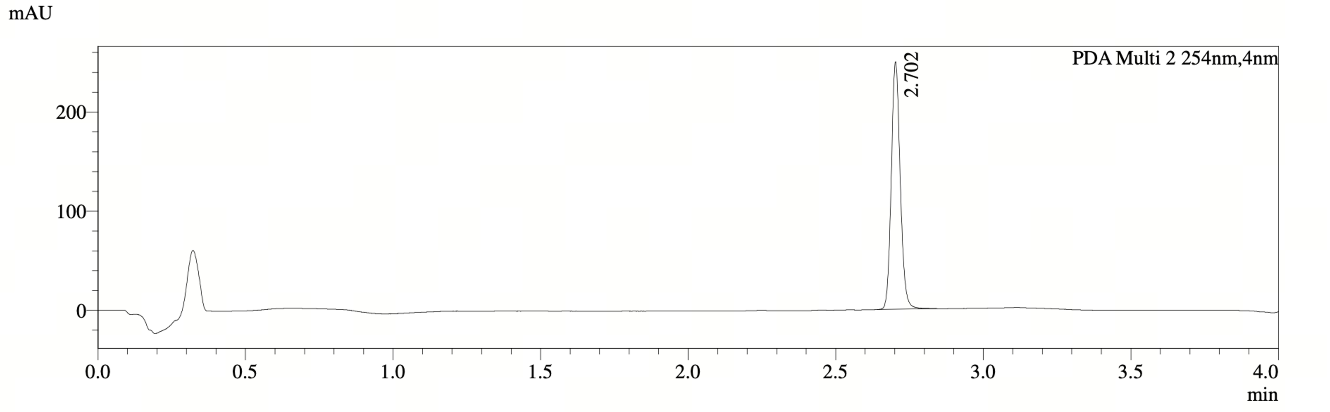

PDA Ch2 254nm

| Peak# | Ret. Time | Width | Height | Height% | Area | Area% |
|---|---|---|---|---|---|---|
| 1 | 2.702 | 0.056 | 249593 | 100.000 | 527480 | 100.000 |

**HPLC Chromatogram of PCSK9-compound-3**

##### *7.2.4 PCSK9-compound-4*

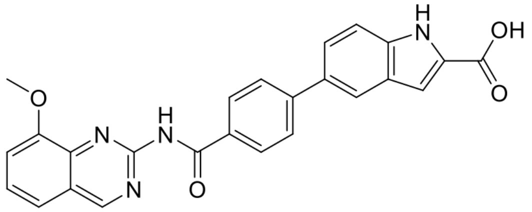

**PCSK9-compound-4**. 5-(4-((8-methoxyquinazolin-2-yl)carbamoyl)phenyl)-1H-indole-2-carboxy licacid

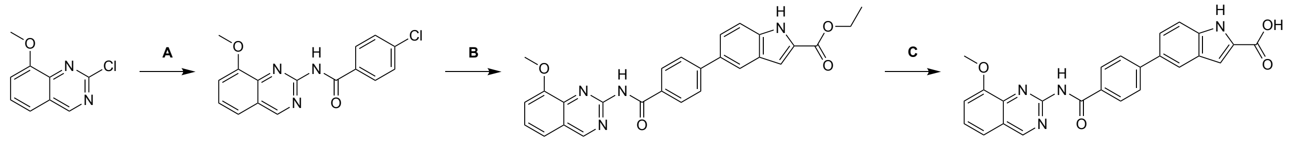

PCSK9-compound-4

Reagents and conditions: (A) 4-chlorobenzamide, $Pd(OAc)_2$, XantPhos, $K_2CO_3$, dioxane, 90 °C 84.8%; (B) ethyl 5-(4,4,5,5-tetramethyl-1,3,2-dioxaborolan-2-yl)-1H-indole-2-carboxylate, cataCXium® A Pd $G_3$, $K_3PO_4$, THF, $H_2O$, 60 °C, 67.3%; (C) NaOH, MeOH, $H_2O$, 25 °C, 12.7%.

#### Synthesis of PCSK-compound-4

Step A: 4-chloro-N-(8-methoxyquinazolin-2-yl)benzamide: To a solution of 2-chloro-8-methoxyquinazoline (500 mg, 2.57 mmol, 1.00 equiv) and Xantphos (150 mg, 259 $\mu$mol, 0.10 equiv) in dioxane (5.00 mL) were added potassium carbonate (710 mg, 5.14 mmol, 2.00 equiv), 4-chlorobenzamide (500 mg, 3.21 mmol, 1.25 equiv) and palladium acetate (50.0 mg, 223 $\mu$mol, 0.09 equiv), the reaction was stirred at 90 °C for 12 hours under nitrogen. After completion of the reaction, the reaction mixture was quenched with water (20 mL) and extracted with Ethyl acetate (30 mL × 3). The combined organic layers were washed with brine (20 mL), dried over anhydrous sodium sulfate, filtered and concentrated under reduced pressure to give a residue. The residue was purified by prep-HPLC (column: Waters Xbridge C18 150*50mm*10um; mobile phase: [$H_2O$ (0.225%FA)-ACN]; gradient: 32%-62% B over 15.0 min) to afford 4-chloro-N-(8-methoxyquinazolin-2-yl)benzamide (690 mg, 84.4% yield) as a yellow solid; LCMS-Method 1 $[M+1]^+ = 314.0$.

Step B: ethyl 5-[4-[(8-methoxyquinazolin-2-yl)carbamoyl]phenyl]-1H-indole-2-carboxylate: To a solution of 4-chloro-N-(8-methoxyquinazolin-2-yl)benzamide (65.0 mg, 207 $\mu$mol, 1.00 equiv) in water (0.30 mL) and tetrahydrofuran (0.50 mL) were added potassium phosphate (1.5 M, 3.00 equiv), ethyl 5-(4,4,5,5-tetramethyl-1,3,2-dioxaborolan-2-yl)-1H-indole-2-carboxylate (100 mg, 317 $\mu$mol, 1.53 equiv) and [2-(2-aminophenyl)phenyl]palladium(1+); bis(1-adamantyl)-butyl-phosphane;methanesulfonate (16.0 mg, 22.0 $\mu$mol, 0.10 equiv), the reaction was stirred at 60 °C for 3 hours under nitrogen. After completion of the reaction, the reaction mixture was quenched with water (10 mL) and extracted with Ethyl acetate (10mL × 3). The combined organic layers were washed with brine (10 mL), dried over anhydrous sodium sulfate, filtered and concentrated under reduced pressure to give a residue. The residue was purified by prep-HPLC (column: Waters Xbridge C18 150*50mm*10$\mu$m; mobile phase: [$H_2O$(0.225%FA)-ACN]; gradient: 35%-65% B over 30.0 min ) to afford ethyl 5-[4-[(8-methoxyquinazolin-2-yl)carbamoyl]phenyl]-1H-indole-2-carboxylate (130 mg, 67.3% yield) as a white solid; LCMS-Method 1 $[M+1]^+ = 467.2$.

Step C: PCSK9-compound-4: To a solution of ethyl 5-[4-[(8-methoxyquinazolin-2-yl)carbam oyl]phenyl]-1H-indole-2-carboxylate (100 mg, 214 $\mu$mol, 1.00 equiv) in methanol (1.00 mL) and water (1.00 mL) was added sodium hydroxide (25.0 mg, 625 $\mu$mol, 2.92 equiv), the reaction mixture was stirred at 25 °C for 2 hours under nitrogen. After completion of the reaction, the reaction mixture was quenched with water (20 mL) and filtered. The cake was collected and stirred at 25 °C with N,N-dimethylformamide (10 mL) for 30 minutes. Then the mixture was filtered and the cake was dried under reduced pressure to afford 5-[4-[(8-methoxyquinazolin-2-yl)carbamoyl]phenyl]-1H-indole-2-carboxylic acid (25.93 mg, 12.7% yield) as off-white solid; $^1H$ NMR (400 MHz, DMSO-d6) $\delta = 11.24$ (s, 1H), 10.90 (s, 1H), 9.54 (s, 1H), 8.09 (d, $J = 8.4$ Hz, 2H), 7.88 (s, 1H), 7.82 (d, $J = 8.4$ Hz, 2H), 7.66 (d, $J = 8.0$ Hz, 1H), 7.57 (t, $J = 8.0$ Hz, 1H), 7.48 - 7.38 (m, 3H), 6.57 (s, 1H), 3.99 (s, 3H); LCMS $[M+1]^+ = 439.2$; HPLC Rt = 1.274 min.

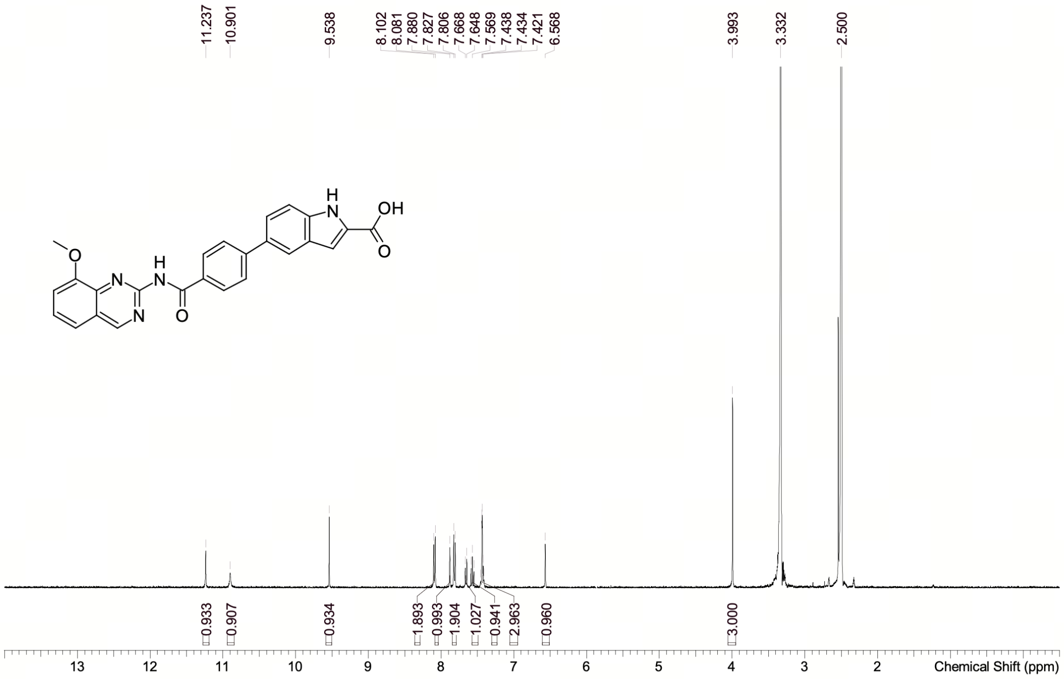

| | |
|---|---|
| Acquisition Time (sec) | 1.9999 |
| Comment | EW67056-14-P1A DMSO BRUKER_03_015_400MHz |
| Date | 16 Sep 2025 17:23:43 (GMT+08:00) |
| Frequency (MHz) | 400.1300 |
| Nucleus | 1H |
| Number of Transients | 1 |
| Origin | Avance neo 400 |
| Original Points Count | 16393 |
| Owner | nmrsu |
| Points Count | 65536 |
| Pulse Sequence | zg |
| Receiver Gain | 18.00 |
| SW(cyclical) (Hz) | 8196.72 |
| Solvent | DMSO-d6 |
| Spectrum Offset (Hz) | 2467.2405 |
| Spectrum Type | standard |
| Sweep Width (Hz) | 8196.60 |
| Temperature (degree C) | 24.264 |

Confidential. For research information only

$^1$H NMR (400 MHz, DMSO-$d_6$) δ = 11.24 (s, 1H), 10.90 (s, 1H), 9.54 (s, 1H), 8.09 (d, *J* = 8.4 Hz, 2H), 7.88 (s, 1H), 7.82 (d, *J* = 8.4 Hz, 2H), 7.66 (d, *J* = 8.0 Hz, 1H), 7.57 (t, *J* = 7.9 Hz, 1H), 7.48 - 7.38 (m, 3H), 6.57 (s, 1H), 3.99 (s, 3H)

Operator: Date:

#### NMR Spectrum of PCSK9-compound-4

RetTime: 0.458 Datafile: D:\DATA\2025\2509\250916\EW67056-14-P1D1.lcd

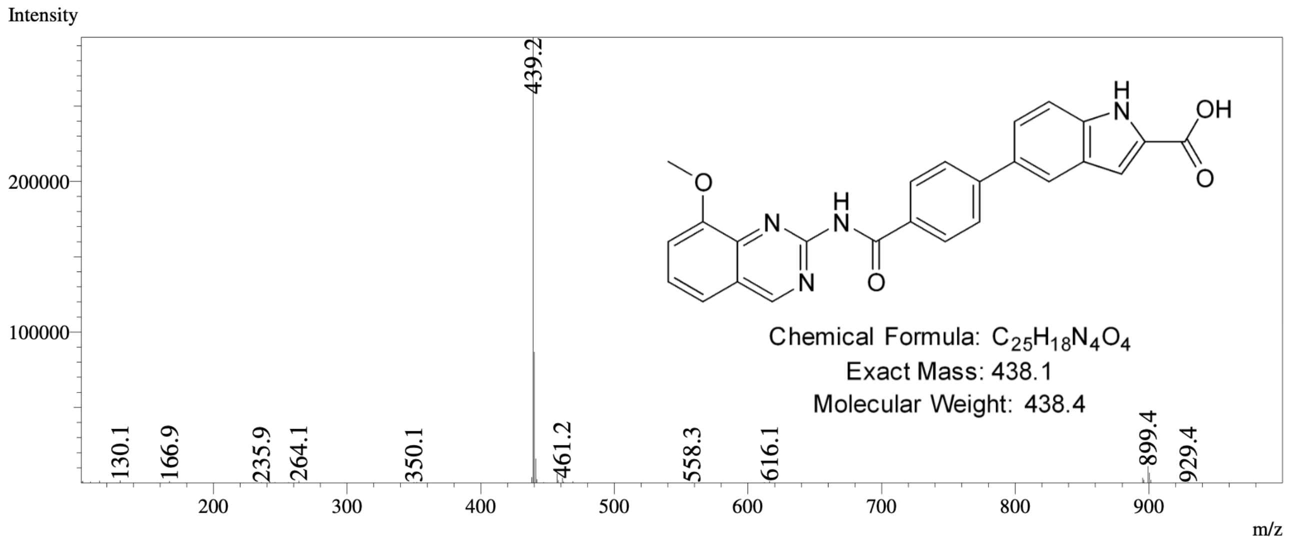

#### MS Spectrum of PCSK9-compound-4

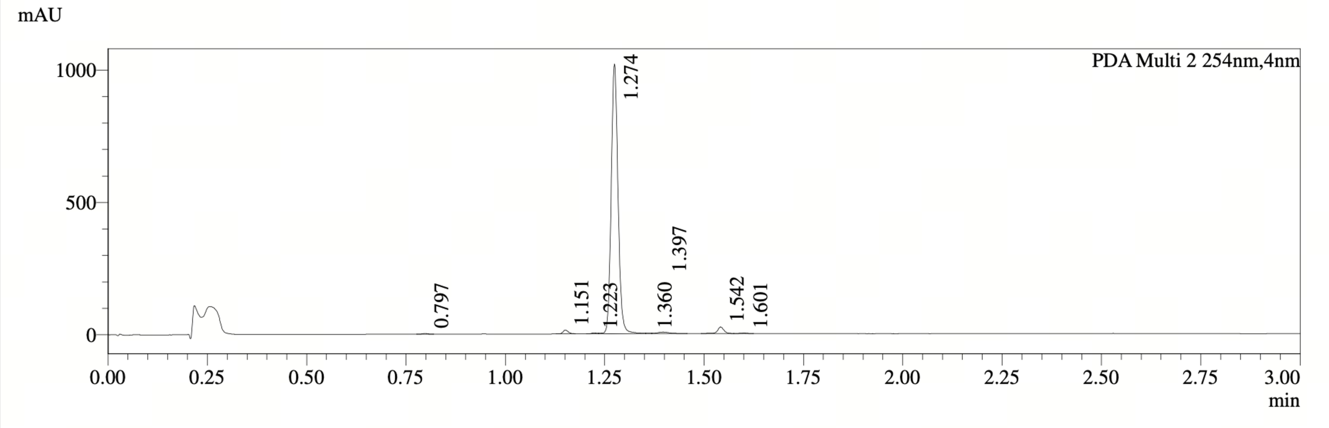

PDA Ch2 254nm

| Peak# | Ret. Time | Width | Height | Height% | Area | Area% |
|---|---|---|---|---|---|---|
| 1 | 0.797 | 0.026 | 2271 | 0.212 | 2019 | 0.158 |
| 2 | 1.151 | 0.027 | 14458 | 1.348 | 13875 | 1.083 |
| 3 | 1.223 | 0.033 | 3455 | 0.322 | 3371 | 0.263 |
| 4 | 1.274 | 0.033 | 1018533 | 94.959 | 1218193 | 95.092 |
| 5 | 1.360 | 0.282 | 1537 | 0.143 | 1303 | 0.102 |
| 6 | 1.397 | 0.054 | 5477 | 0.511 | 11703 | 0.914 |
| 7 | 1.542 | 0.029 | 24843 | 2.316 | 28022 | 2.187 |
| 8 | 1.601 | 0.032 | 2029 | 0.189 | 2587 | 0.202 |

**HPLC Chromatogram of PCSK9-compound-4**

##### *7.2.5 PCSK9-compound-5*

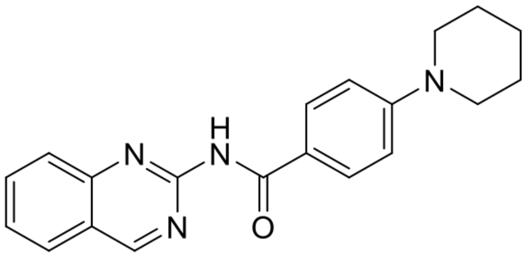

**PCSK9-compound-5**. 4-(piperidin-1-yl)-N-(quinazolin-2-yl)benzamide

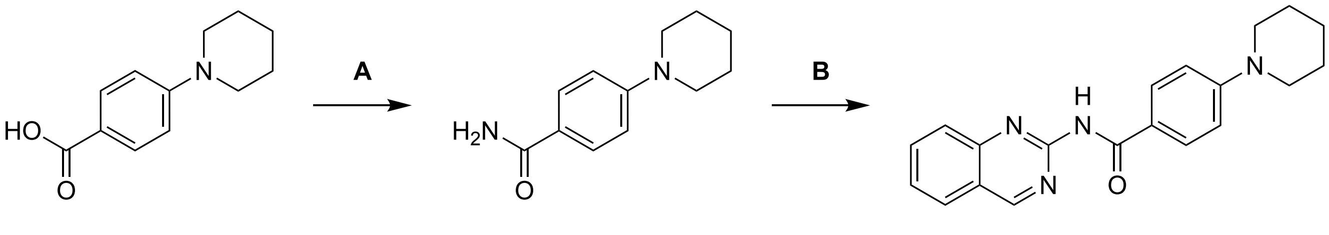

**PCSK9-compound-5**

Reagents and conditions: (A) $(COCl)_2$, THF, 25 °C, then $NH_3$/THF; (B) 4-(1-piperidyl)benzamide, $Pd(OAc)_2$, N-XantPhos, $K_2CO_3$, dioxane, 90 °C, 18.4%.

**Synthesis of PCSK-compound-5**

Step A: 4-(1-piperidyl)benzamide: To a solution of 4-(1-piperidyl)benzoic acid (100 mg, 1.00 equiv) and N,N-dimethylformamide (4.00 mg, 0.11 equiv) in tetrahydrofuran (2.00 mL) was added oxalyl dichloride (75.0 mg, 1.21 equiv). The mixture was stirred at 25 °C for 1 hour under nitrogen. The mixture was concentrated under reduced pressure to give a crude. Then ammonia /tetrahydrofuran (10 mL) was added to the above mixture and stirred at 25 °C for 11 hours under nitrogen. After completion of the reaction, the mixture was concentrated under reduced pressure to afford 4-(1-piperidyl)benzamide (100 mg, crude) as a yellow solid; LCMS-Method 1 $[M+1]^+ = 205.2$.

Step B: PCSK9-compound-5: To a solution of 2-chloroquinazoline (46.0 mg, 1.14 equiv) in dioxane (1.00 mL) were added 4-(1-piperidyl)benzamide (50.0 mg, 1.0 equiv), potassium carbonate (68.0 mg, 2.01 equiv), 4,6-bis(diphenylphosphanyl)-10H-phenoxazine (29.0 mg, 0.21 equiv) and palladium acetate (7.00 mg, 0.13 equiv). The mixture was stirred at 90 °C for 12 hours under nitrogen. After completion of the reaction, the mixture was quenched with water (10 mL) and extracted with ethyl acetate (10 mL × 3). The combined organic layers were washed with brine (10 mL), dried over anhydrous sodium sulfate, filtered and concentrated under reduced pressure to give a residue. The residue was purified by prep-HPLC (column: WePure PrePulite Platinum C18 150*25mm*7um; mobile phase: [$H_2O$ (0.225% FA)-ACN]; gradient: 23%-53% B over 15.0 min) to afford 4-(1-piperidyl)-N-quinazolin-2-yl-benzamide (31.32 mg, 18.4% yield) as a yellow solid; $^1H$ NMR (400 MHz, DMSO-d6) $\delta = 10.80$ (s, 1H), 9.54 (s, 1H), 8.10 (d, $J = 8.0$ Hz, 1H), 8.00 - 7.86 (m, 3H), 7.84 (d, $J = 8.4$ Hz, 1H), 7.62 (t, $J = 7.2$ Hz, 1H), 6.97 (d, $J = 8.8$ Hz, 2H), 3.34 - 3.32 (m, 4H), 1.60 (s, 6H); LCMS $[M+1]^+ = 333.1$; HPLC Rt = 1.208 min.

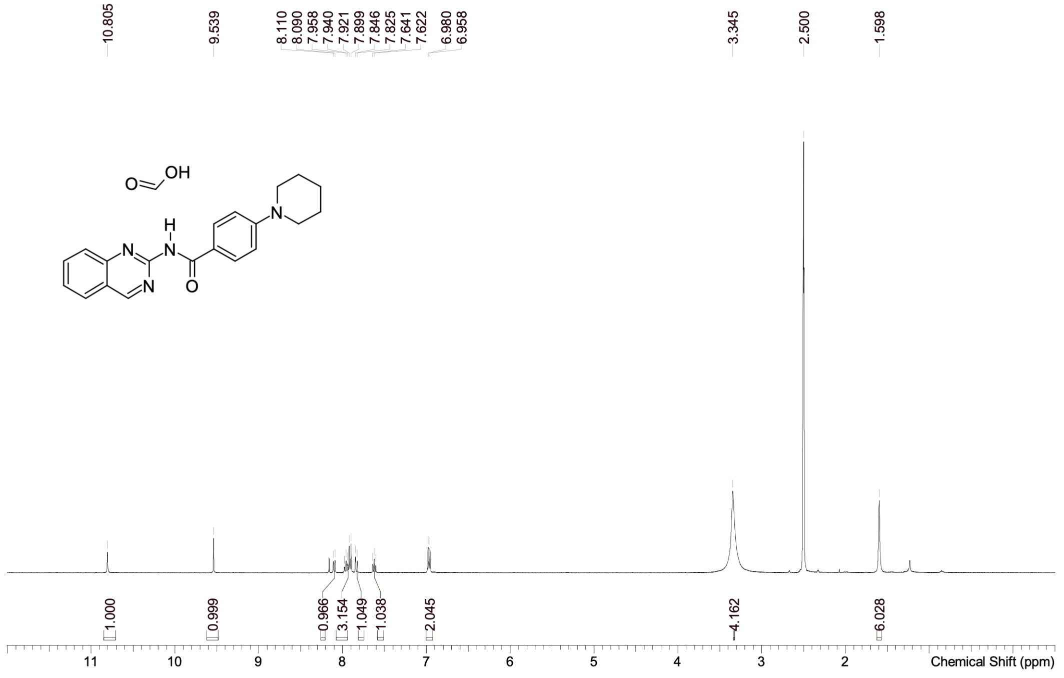

| | |
|---|---|
| Acquisition Time (sec) | 1.9999 |
| Comment | EW67056-7-P1A DMSO BRUKER_03_015_400MHz |
| Date | 29 Aug 2025 17:11:26 (GMT+08:00) |
| Frequency (MHz) | 400.1300 |
| Nucleus | 1H |
| Number of Transients | 1 |
| Origin | Avance neo 400 |
| Original Points Count | 16393 |
| Owner | nmrsu |
| Points Count | 65536 |
| Pulse Sequence | zg |
| Receiver Gain | 18.00 |
| SW(cyclical) (Hz) | 8196.72 |
| Solvent | DMSO-d6 |
| Spectrum Offset (Hz) | 2467.7205 |
| Spectrum Type | standard |
| Sweep Width (Hz) | 8196.60 |
| Temperature (degree C) | 24.351 |

Confidential. For research information only

$^1H$ NMR ( 400 MHz, DMSO-$d_6$) δ = 10.80 ( s, 1H) , 9.54 ( s, 1H) , 8.10 ( d, $J$ = 8.0 Hz, 1H) , 8.00 - 7.86 ( m, 3H) , 7.84 ( d, $J$ = 8.4 Hz, 1H) , 7.62 ( t, $J$ = 7.4 Hz, 1H) , 6.97 ( d, $J$ = 9.0 Hz, 2H) , 3.34 - 3.32 ( m, 4H) , 1.60 ( s, 6H)

**NMR Spectrum of PCSK9-compound-5**

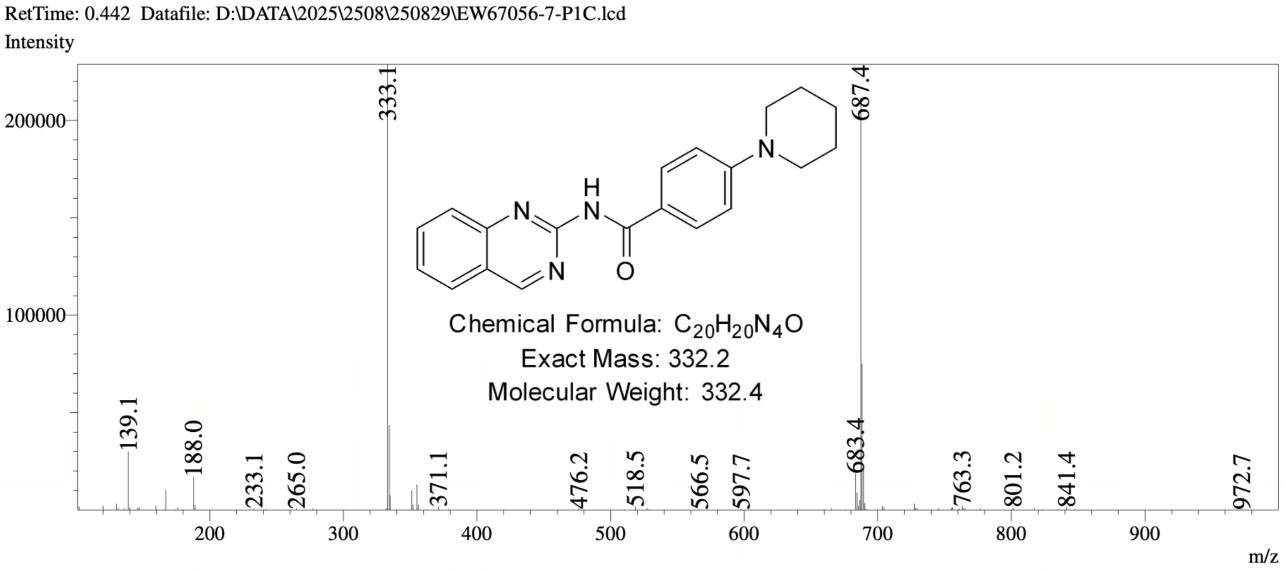

#### MS Spectrum of PCSK9-compound-5

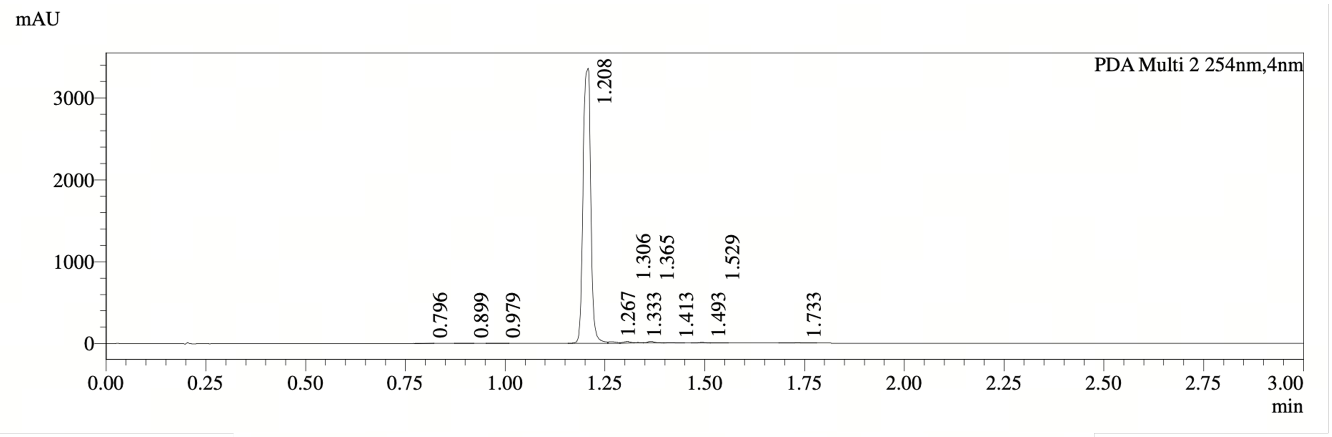

PDA Ch2 254nm

| Peak# | Ret. Time | Width | Height | Height% | Area | Area% |
|---|---|---|---|---|---|---|
| 1 | 0.796 | 0.040 | 714 | 0.021 | 1095 | 0.024 |
| 2 | 0.899 | 0.034 | 1590 | 0.046 | 1982 | 0.043 |
| 3 | 0.979 | 0.040 | 792 | 0.023 | 1195 | 0.026 |
| 4 | 1.208 | 0.033 | 3356686 | 97.437 | 4499672 | 97.475 |
| 5 | 1.267 | 0.067 | 18424 | 0.535 | 24069 | 0.521 |
| 6 | 1.306 | 0.038 | 19435 | 0.564 | 24775 | 0.537 |
| 7 | 1.333 | 0.059 | 7157 | 0.208 | 8212 | 0.178 |
| 8 | 1.365 | 0.033 | 21406 | 0.621 | 27386 | 0.593 |
| 9 | 1.413 | 0.036 | 5227 | 0.152 | 6768 | 0.147 |
| 10 | 1.493 | 0.031 | 7577 | 0.220 | 8693 | 0.188 |
| 11 | 1.529 | 0.056 | 2097 | 0.061 | 2899 | 0.063 |
| 12 | 1.733 | 0.058 | 3889 | 0.113 | 9504 | 0.206 |

#### HPLC Chromatogram of PCSK9-compound-5

##### 7.2.6 PCSK9-compound-6

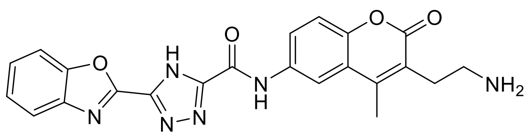

**PCSK9-compound-6**. N-(3-(2-aminoethyl)-4-methyl-2-oxo-2H-chromen-6-yl)-5-(benzo[d]oxazol-2-yl)-4H-1,2,4-triazole-3-carboxamide

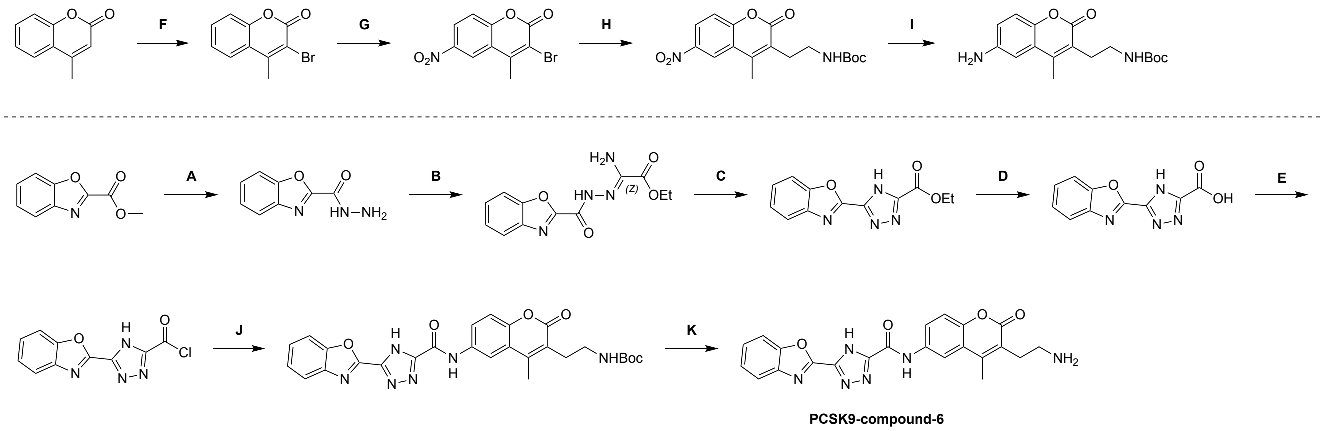

Reagents and conditions: (A) $NH_2NH_2 \cdot H_2O$, EtOH, 25 °C, 99.5%; (B) ethyl 2-amino-2-thioxo-acetate, EtOH, 80 °C, 44.9%; (C) diphenyl ether, 165 °C, 63.2%; (D) LiOH•$H_2O$, THF, $H_2O$, 25 °C, 53.8%; (E) $(COCl)_2$, DCM, DMF, 0 °C; (F) $Br_2$, AcOH, 25 – 60 °C, 93.8%; (G) $HNO_3$, $H_2SO_4$, 0 °C, 49.6%; (H) potassium;2-(tert-butoxycarbonylamino)ethyl-trifluoro-boranuide, $Pd(dba)_2$, RuPhos, $Cs_2CO_3$, toluene, $H_2O$, 90 °C, 72.8%; (I) Fe $NH_4Cl$, EtOH, $H_2O$, 60 °C, 20%; (J) tert-butyl N-[2-(6-amino-4-methyl-2-oxo-chromen-3-yl)ethyl]carbamate, TEA, DCM, 25 °C, 18%; (K) TFA, DCM, 25 °C, 29.5%.

**Synthesis of PCSK9-compound-6**

Step A: 1,3-benzoxazole-2-carbohydrazide:Toasolutionofmethyl1,3-benzoxazole-2-carboxylate (3.70 g, 20.9 mmol, 1.0 equiv) in ethanol (150 mL) was added hydrazine hydrate (1.32 g, 25.1 mmol, 1.28 mL, 95% purity, 1.2 equiv) at 25 °C. The reaction was stirred at 80 °C for 2 hours. After completion of the reaction, the reaction mixture was quenched by water (100 mL), extracted with ethyl acetate (100 mL × 2). The combined organic layers were washed with brine (100 mL), dried over anhydrous sodium sulfate, filtered and concentrated under reduced pressure to afford 1,3-benzoxazole-2-carbohydrazide (3.68 g, 99.5% yield) as yellow solid; LCMS-Method 1 $[M+1]^+$ = 178.1.

Step B: ethyl(2Z)-2-amino-2-(1,3-benzoxazole-2-carbonylhydrazono)acetate:Toasolutionof1,3-benzoxazole-2-carbohydrazide (1.00 g, 5.64 mmol, 1.0 equiv) in ethanol (10 mL) was added ethyl 2-amino-2-thioxo-acetate (1.50 g, 11.3 mmol, 2.0 equiv). The reaction was stirred at 80 °C for 3 hours. After completion of the reaction, the mixture was filtered and the filter cake was dried over under reduced pressure to afford ethyl (2Z)-2-amino-2-(1,3-benzoxazole-2-carbonylhydrazono)acetate (970 mg, 44.9% yield) as yellow solid; LCMS-Method 1 $[M+1]^+$ = 277.1.

Step C: ethyl5-(1,3-benzoxazol-2-yl)-4H-1,2,4-triazole-3-carboxylate:Amixtureofethyl(2Z)-2-amino-2-(1,3-benzoxazole-2-carbonylhydrazono)acetate (970 mg, 3.51 mmol, 1.0 equiv) in diphenyl ether (10.0 mL) was stirred at 165 °C for 3 hours. After completion of the reaction, the reaction mixture was quenched by water (50.0 mL) and extracted with ethyl acetate (50.0 mL × 2). The combined organic layers were washed with brine (50.0 mL), dried over anhydrous sodium sulfate, filtered and concentrated under reduced pressure to give a residue. The residue was purified by column chromatography (silicon dioxide, Commercial hexanes: Ethyl acetate = 100/1 to 0/1) to afford ethyl 5-(1,3-benzoxazol-2-yl)-4H-1,2,4-triazole-3-carboxylate (780 mg, 63.2% yield) as yellow solid; LCMS-Method 1 $[M+1]^+ = 259.1$.

Step D: 5-(benzo[d]oxazol-2-yl)-4H-1,2,4-triazole-3-carboxylicacid:Toasolutionofethyl5-(1,3-benz oxazol-2-yl)-4H-1,2,4-triazole-3-carboxylate (730 mg, 2.83 mmol, 1.0 equiv) in tetrahydrofuran (10.0 mL) and water (2.0 mL) was added lithium hydrate (593 mg, 14.13 mmol, 5.0 equiv). The reaction was stirred at 25 °C for 2 hours. After completion of the reaction, the mixture was washed with ethyl acetate (20.0 mL × 2). The aqueous layer was adjusted the pH to 3 with hydrochloric acid (1M) and extracted with ethyl acetate (20.0 mL × 3). The combined organic layers were washed with brine (20.0 mL), dried over anhydrous sodium sulfate, filtered and concentrated under reduced pressure to afford 5-(benzo[d]oxazol-2-yl)-4H-1,2,4-triazole-3-carboxylic acid (350 mg, 53.79% yield) as an off-white solid; LCMS-Method 1 $[M+1]^+ = 231.1$.

Step E: 5-(1,3-benzoxazol-2-yl)-4H-1,2,4-triazole-3-carbonylchloride:Toasolutionof5-(benzo[d]oxa zol-2-yl)-4H-1,2,4-triazole-3-carboxylicacid (100 mg, 434 $\mu$mol, 1.0 equiv) in dichloromethane (2.0 mL) were added oxalyl dichloride (66.2 mg, 521 $\mu$mol, 45.6 $\mu$L, 1.2 equiv) and N, N-dimethylformamide (3.18 mg, 43.4 $\mu$mol, 3.34 $\mu$L, 0.1 equiv) at 0 °C. The reaction was stirred at 25 °C for 2 hours. After completion of the reaction, the reaction mixture was concentrated under reduced pressure to afford 5-(1,3-benzoxazol-2-yl)-4H-1,2,4-triazole-3-carbonyl chloride (200 mg, crude) as white solid.

Step F: 3-bromo-4-methyl-chromen-2-one: To a solution of 4-methylchromen-2-one (1.00 g, 6.24 mmol, 1.0 equiv) in acetic acid (10.0 mL) was added dropwise bromine (1.24 g, 7.76 mmol, 0.4 mL, 1.24 equiv) at 25 °C. The reaction was stirred at 60 °C for 12 hours. After completion of the reaction, the mixture was quenched with water (50.0 mL) and extracted with ethyl acetate (50.0 mL × 2). The combined organic layers were washed with brine (50.0 mL), dried over anhydrous sodium sulfate, filtered and concentrated under reduced pressure to give a residue. The residue was purified by column chromatography (silicon dioxide, Commercial hexanes: Ethyl acetate = 100/1 to 0/1) to afford 3-bromo-4-methyl-chromen-2-one (1.40 g, 93.8% yield) as white solid; LCMS-Method 1 $[M+1]^+ = 239.0$.

Step G: 3-bromo-4-methyl-6-nitro-chromen-2-one: Solution 1: 3-bromo-4-methyl-chromen-2-one (1.0 g, 4.18 mmol, 1.0 equiv) was dissolved in sulfuric acid (10.0 mL). Solution 2: nitric acid (322 mg, 5.02 mmol, 230 $\mu$L, 98% purity, 1.20 equiv). Solution 1 and 2 were mixed with a T-shaped mixer at 0 °C and the combined solution was passed into a perfluoroalkoxy (PFA) Coils reactor (Outer diameter: 3.175 mm(1/8 inch); Internal volume: 10 mL) at 0 °C, with flow rates of 1.96 mL/min and 0.04 mL/min, respectively. The residence time of flow reactor was 20 mins. After completion of the reaction, the reactor effluent was collected in a flat-bottomed container (Quenched by 50.0 mL water at 15 °C) and extracted with dichloromethane (20.0 mL ×3). The combined organic layers were washed with brine (50.0 mL), dried over anhydrous sodium sulfate, filtered and concentrated under reduced pressure to give a residue. The residue was purified by Chromatography (Commercial hexanes: Ethyl acetate = 10/1) to afford 3-bromo-4-methyl-6-nitro-chromen-2-one (600 mg, 49.6% yield) as white solid; LCMS-Method 1 $[M+1,M+3]^+ = 283.8, 285.8$.

Step H: tert-butylN-[2-(4-methyl-6-nitro-2-oxo-chromen-3-yl)ethyl]carbamate: To a solution of 3-bromo-4-methyl-6-nitro-chromen-2-one (600 mg, 2.11 mmol, 1.0 equiv) and potassium;2-(tert-butoxycarbonylamino)ethyl-trifluoro-boranuide (636 mg, 2.53 mmol, 1.2 equiv) in toluene (6.00 mL) and water (2.0 mL) were added cesium carbonate (2.06 g, 6.34 mmol, 3.0 equiv), [2',6'-bis(propan-2-yloxy)-[1,1'-biphenyl]-2-yl]dicyclohexylphosphane (197 mg, 422 $\mu$mol, 0.20 equiv) and 1,5-diphenylpenta-1,4-dien-3-one palladium (121 mg, 211 $\mu$mol, 0.10 equiv). The reaction was stirred at 90 °C for 12 hours. After completion of the reaction, the reaction mixture was quenched with water (20.0 mL) and extracted with ethyl acetate (20.0 mL × 2). The combined organic layers were washed with brine (50.0 mL), dried over anhydrous sodium sulfate, filtered and concentrated under reduced pressure to give a residue. The residue was purified by Chromatography (Commercial hexanes: Ethyl acetate = 3/1) to afford tert-butyl N-[2-(4-methyl-6-nitro-2-oxo-chromen-3-yl)ethyl]carbamate (600 mg, 72.8% yield) as white solid; LCMS-Method 1 $[M-100]^+$ = 248.1.

Step I. tert-butylN-[2-(6-amino-4-methyl-2-oxo-chromen-3-yl)ethyl]carbamate: To a solution of tert-butyl N-[2-(4-methyl-6-nitro-2-oxo-chromen-3-yl)ethyl]carbamate (530 mg, 1.52 mmol, 1.0 equiv) in ethanol (6.00 mL) and water (3.00 mL) were added iron (424 mg, 7.61 mmol, 5.0 equiv) and ammonium chloride (813 mg, 15.2 mmol, 10.0 equiv) at 60 °C. The reaction was stirred at 60 °C for 2 hours. After completion of the reaction, the mixture was filtered and filtrate was concentrated under reduced pressure to give a residue. The residue was purified by column chromatography (silicon dioxide, Commercial hexanes: Ethyl acetate = 100/1 to 0/1) to afford tert-butyl N-[2-(6-amino-4-methyl-2-oxo-chromen-3-yl)ethyl]carbamate (120 mg, 20.0% yield) as yellow oil.

Step J: tert-butylN-[2-[6-[[5-(1,3-benzoxazol-2-yl)-4H-1,2,4-triazole-3-carbonyl]amino]-4-methyl-2-oxo-chromen-3-yl]ethyl]carbamate: To a solution of tert-butyl N-[2-(6-amino-4-methyl-2-oxo-chromen-3-yl)ethyl]carbamate (120 mg, 376.92 μmol, 1.0 equiv) in dichloromethane (2.00 mL) were added triethylamine (114 mg, 1.13 mmol, 157 μL, 3.0 equiv) and 5-(1,3-benzoxazol-2-yl)-4H-1,2,4-triazole-3-carbonyl chloride (200 mg, 804 μmol, 2.13 equiv). The reaction was stirred at 25 °C for 1 hour. After completion of the reaction, the reaction mixture was concentrated under reduced pressure to give a residue. The residue was purified by reversed-phase flash (0.1% FA condition) to afford tert-butyl N-[2-[6-[[5-(1,3-benzoxazol-2-yl)-4H-1,2,4-triazole-3-carbonyl]amino]-4-methyl-2-oxo-chromen-3-yl]ethyl]carbamate (38.0 mg, 18.0% yield) as yellow oil; LCMS-Method 1 $[M-99]^+$ = 431.2.

Step K: PCSK9-compound-6: To a solution of tert-butylN-[2-[6-[[5-(1,3-benzoxazol-2-yl)-4H-1,2,4-triazole-3-carbonyl]amino]-4-methyl-2-oxo-chromen-3-yl]ethyl]carbamate (38.0 mg, 71.6 μmol, 1.0 equiv) in dichloromethane (2.00 mL) was added trifluoroacetic acid (767 mg, 6.73 mmol, 94.0 equiv). The reaction was stirred at 25 °C for 20 mins. After completion of the reaction, the reaction mixture was concentrated under reduced pressure to give a residue. The residue was lyophilized to afford N-(3-(2-aminoethyl)-4-methyl-2-oxo-2H-chromen-6-yl)-5-(benzo[d]oxazol-2-yl)-4H-1,2,4-triazole-3-carboxamide (9.68 mg, 29.5% yield, trifluoroacetic acid salt) as brown solid; $^1$H NMR (400 MHz, DMSO-$d_6$) $\delta$ = 11.4 - 11.0 (m, 1H), 8.39 (d, $J$ = 2.4 Hz, 1H), 8.15 (dd, $J$ = 2.0, 8.8 Hz, 1H), 7.91 (t, $J$ = 6.4 Hz, 2H), 7.79 (brs, 3H), 7.57 - 7.45 (m, 3H), 3.02 - 2.91 (m, 4H), 2.47 (s, 3H); $^{19}$F NMR (400 MHz, DMSO-d6) $\delta$ = -74.05; LCMS $[M+1]^+$ = 431.1.

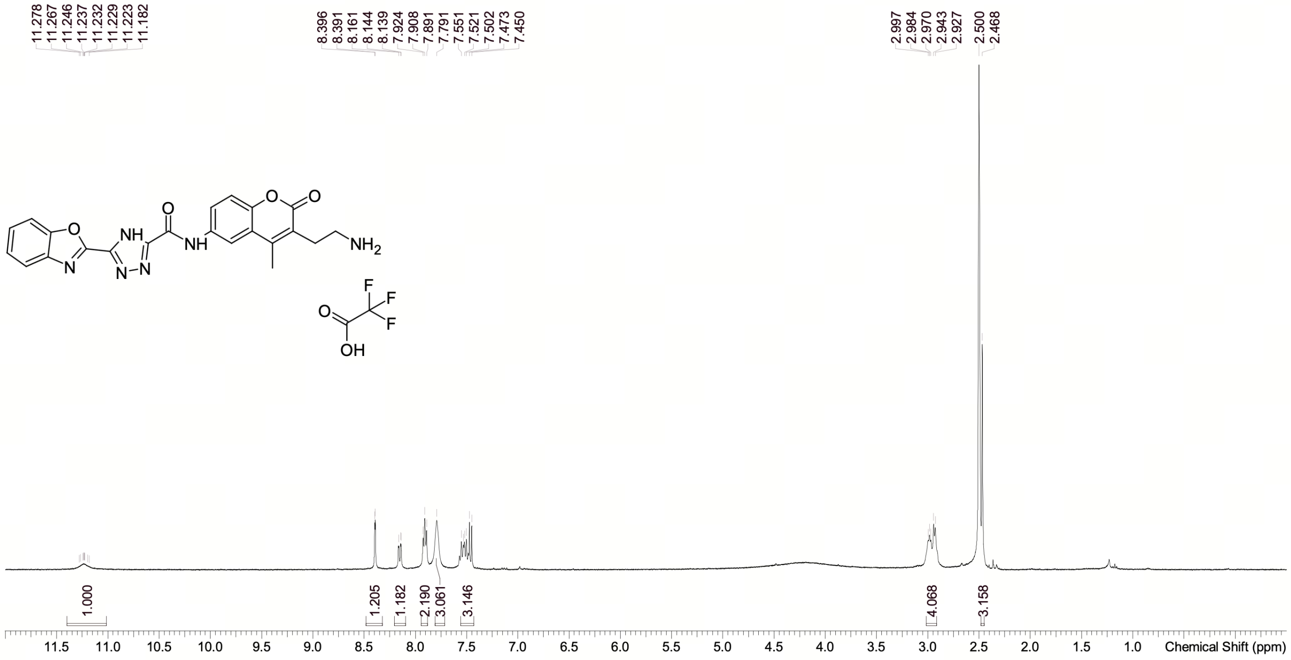

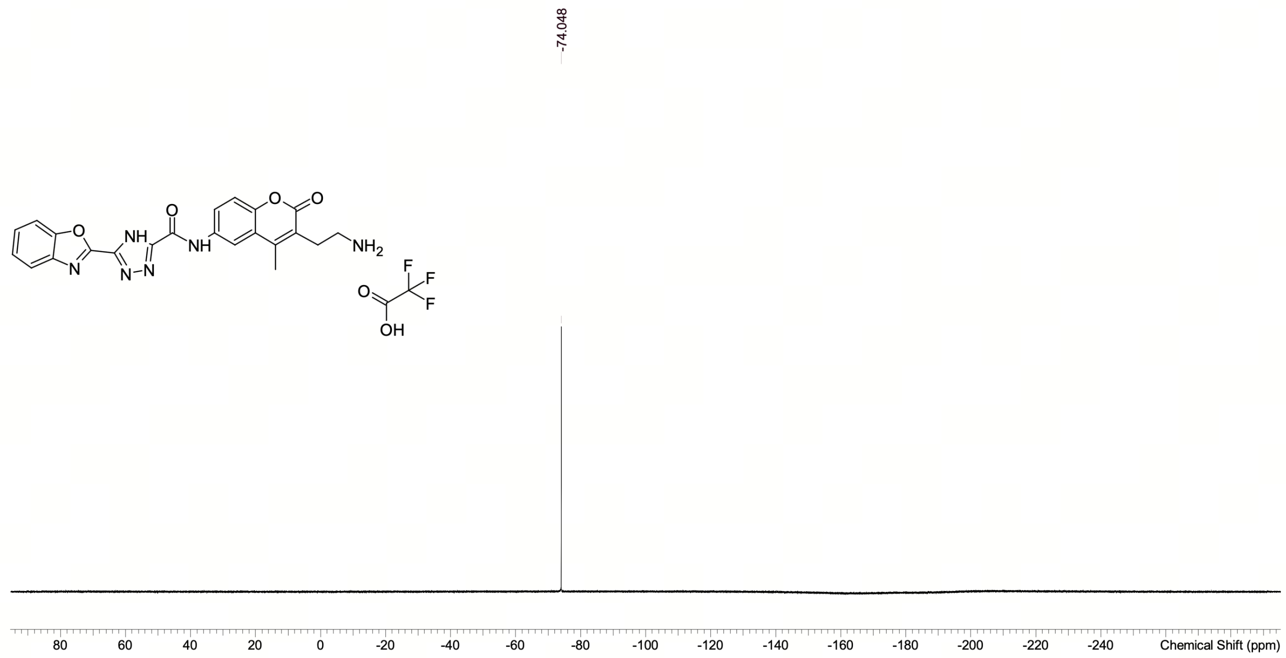

**NMR Spectra of PCSK9-compound-6**

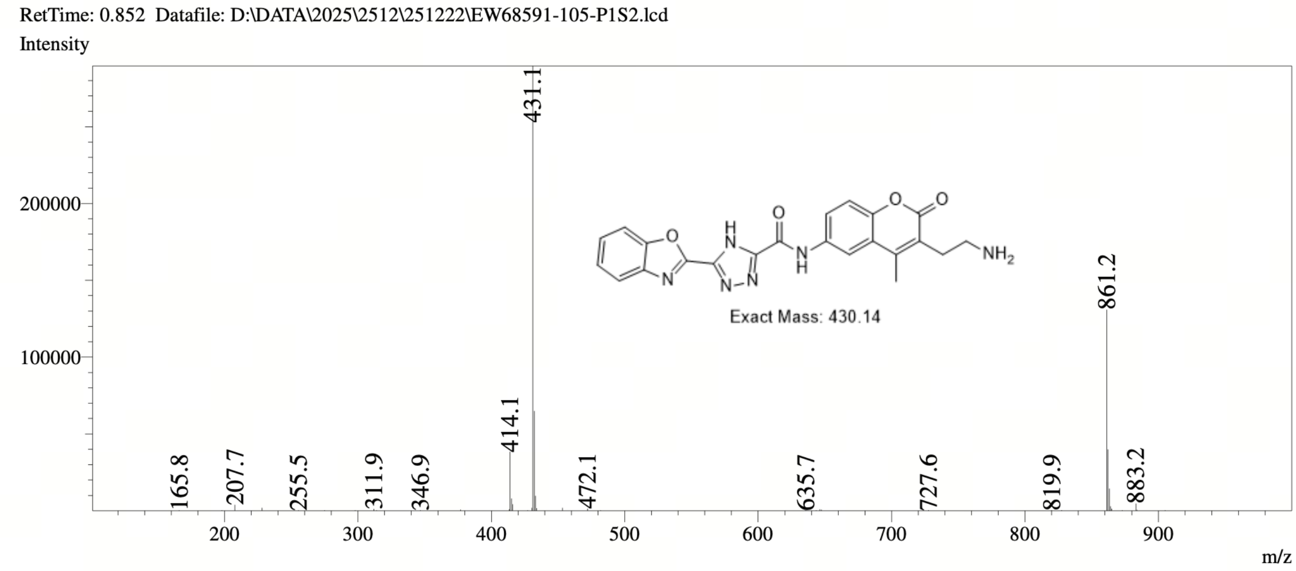

**MS Spectrum of PCSK9-compound-6**

##### *7.2.7 PCSK9-compound-7*

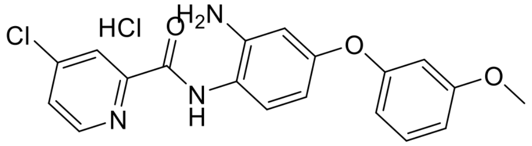

**PCSK9-compound-7**. N-[2-amino-4-(3-methoxyphenoxy)phenyl]-4-chloropyridine-2-carboxami dehydrochloride

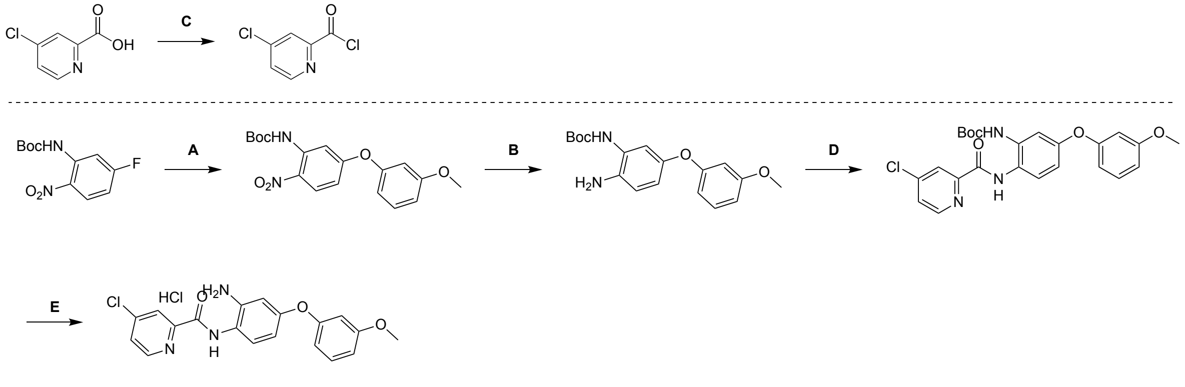

PCSK9-compound-7

Reagents and conditions: A. 3-methoxyphenol, $K_2CO_3$, DMF, 50 °C, 80%; B. $H_2$, THF, 1%Pt/C, 1.5 MPa; C. oxalyl chloride, DMF, THF, 0 to 25 °C; D. TEA, DCM, 0 to 25 °C, 46%; E. HCl/EA, 25 °C, 61%.

**Synthesis of PCSK9-compound-7**

Step A: tert-butyl N-[5-(3-methoxyphenoxy)-2-nitrophenyl]carbamate: To a solution of tert-butyl N-(5-fluoro-2-nitro-phenyl)carbamate (2.00 g, 7.81 mmol, 1.0 equiv) and 3-methoxyphenol (968 mg, 7.81 mmol, 1.0 equiv) in N,N-dimethylformamide (20.0 mL) was added potassium carbonate (2.16 g,

15.6 mmol, 2.0 equiv). The reaction mixture was stirred at 55 °C for 2 hours under nitrogen. After completion of the reaction, the reaction mixture was quenched by water (100 mL) and extracted with ethyl acetate (100 mL × 3). The combined organic layers were washed with brine (100 mL × 2), dried over anhydrous sodium sulfate, filtered and concentrated under reduced pressure to give a residue. The residue was purified by column chromatography (silicon dioxide, commercial hexanes/ethyl acetate = 1/0 to 0/1) to afford tert-butyl N-[5-(3-methoxyphenoxy)-2-nitro-phenyl]carbamate (2.40 g, 80.8% yield) as yellow solid; LCMS-Method 1 $[M-99]^+ = 261.1$.

Step B: tert-butyl N-[2-amino-5-(3-methoxyphenoxy)phenyl]carbamate: Solution 1: tert-butyl N-[5-(3-methoxyphenoxy)-2-nitro-phenyl]carbamate (100 mg, 277 μmol, 1.0 equiv) in tetrahydrofuran (20.0 mL). The fixed bed (named FLR1, volume 5.00 mL) was completely packed with granular catalyst WXSC1017(1%Pt/C), 1.00g. The hydrogen back pressure regulator was adjusted to 1.5 Mpa, and the hydrogen flow rate was 12.0 equiv, 13.7 sccm. The solution 1 was pumped by Pump 1 S1, P1, 0.4 mL/min to flow reactor 1 FLR1, SS, Fixed bed, 6.350(1/4") mm, 5.000 mL, 50.0 °C. The reaction mixture was continuously collected from the reactor outlet into the container. The mixture was collected with a bottle. After completion of the reaction, the mixture was concentrated under reduced pressure to afford tert-butyl N-[2-amino-5-(3-methoxyphenoxy)phenyl]carbamate (120 mg, crude) as yellow oil; LCMS-Method 1 $[M-55]^+ = 275.1$.

Step C: 4-chloropyridine-2-carbonyl chloride: To a solution of 4-chloropyridine-2-carboxylic acid (300 mg, 1.90 mmol, 1.0 equiv) in tetrahydrofuran (3.00 mL) were added N,N-dimethylformamide (6.96 mg, 95.2 μmol, 0.05 equiv) and oxalyl chloride (604 mg, 4.76 mmol, 2.5 equiv) at 0 °C. The reaction mixture was stirred at 25 °C for 0.5 hours under nitrogen. After completion of the reaction, the mixture was concentrated under reduced pressure to afford 4-chloropyridine-2-carbonyl chloride (335 mg, crude) as brown solid.

Step D: tert-butyl N-[2-(4-chloropyridine-2-amido)-5-(3-methoxyphenoxy)phenyl]carbamate: To a solution of tert-butyl N-[2-amino-5-(3-methoxyphenoxy)phenyl]carbamate (120 mg, 363 μmol, 1.0 equiv) in dichloromethane (1.20 mL) was added triethylamine (110 mg, 1.09 mmol, 3.0 equiv) at 0 °C and then a solution of 4-chloropyridine-2-carbonyl chloride (63.9 mg, 363 °mol, 1.0 equiv) in dichloromethane (0.60 mL) was added, the reaction mixture was stirred at 25 °C for 1.5 hours under nitrogen. After completion of the reaction, the reaction mixture was quenched by water (5.00 mL) and extracted with dichloromethane (3.00 mL × 3). The combined organic layers were washed with brine (5.00 mL), dried over anhydrous sodium sulfate, filtered and concentrated under reduced pressure to give a residue. The residue was purified by prep-TLC (Silica, commercial hexanes/ethyl acetate = 3/1) to afford tert-butyl N-[2-[(4-chloropyridine-2-carbonyl)amino]-5-(3-methoxyphenoxy)phenyl]carbamate (80.0 mg, 46.8% yield) as yellow solid; LCMS-Method 1 $[M+1]^+ = 470.2$.

Step E: PCSK9-compound-7: A solution of tert-butylN-[2-[(4-chloropyridine-2-carbonyl)amino]-5-(3-methoxyphenoxy)phenyl]carbamate (70.0 mg, 148 μmol, 1.0 equiv) in hydrochloric acid/ ethyl acetate (2 M, 3.50 mL) was stirred at 25 °C for 2 hours under nitrogen. After completion of the reaction, the mixture was concentrated under reduced pressure to give a residue. The residue was purified by prep-HPLC (column: Phenomenex luna C18 150*25mm*10μm; mobile phase: [water (0.05% HCl)-ACN]; gradient:38%-68% B over 13.0 min) to afford N-[2-amino-4-(3-methoxyphenoxy)phenyl]-4-chloropyridine-2-carboxamide hydrochloride (37.2 mg, , 61.2% yield) as yellow solid; $^1$H NMR (400 MHz, DMSO-d6) $\delta$ = 10.50 - 10.19 (m, 1H), 8.73 (d, $J$ = 5.2 Hz, 1H), 8.13 (d, $J$ = 2.0 Hz, 1H), 7.85 (dd, $J$ = 2.0 Hz, $J$ = 5.4 Hz, 1H), 7.52 - 7.41 (m, 1H), 7.31 (t, $J$ = 8.0 Hz, 1H), 6.85 - 6.56 (m, 5H), 3.75 (s, 3H); LCMS $[M+1]^+ = 370.1$; HPLC Rt = 1.912 min.

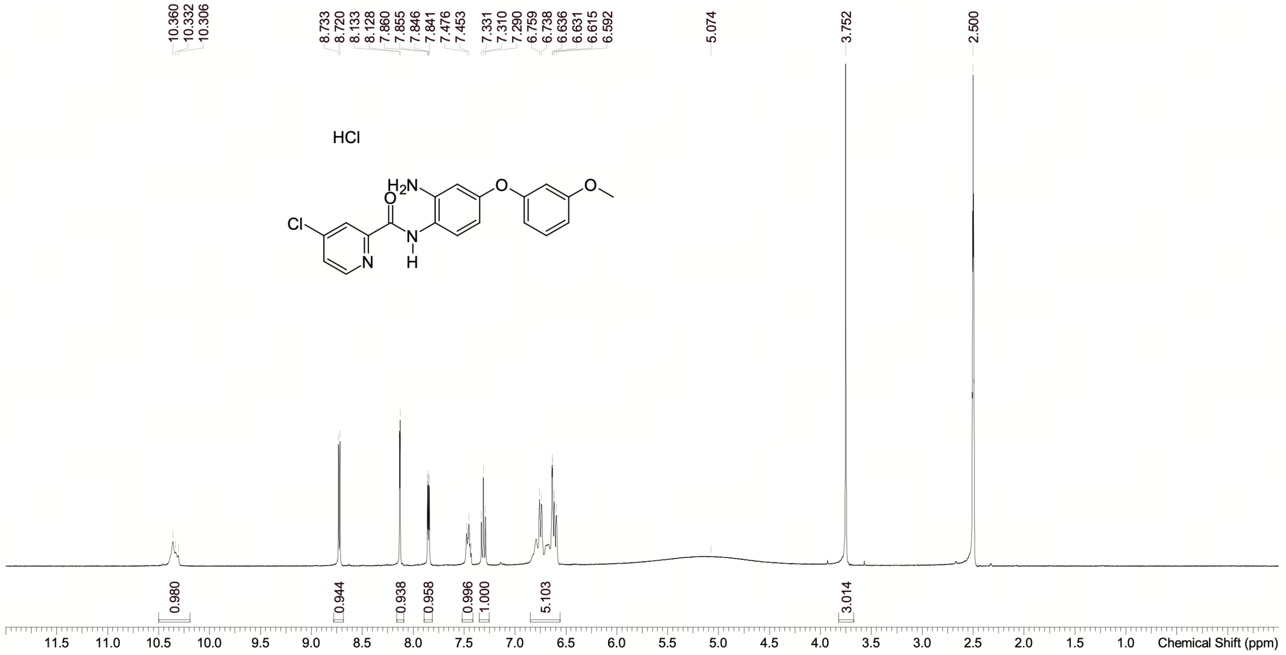

**NMR Spectrum of PCSK9-compound-7**

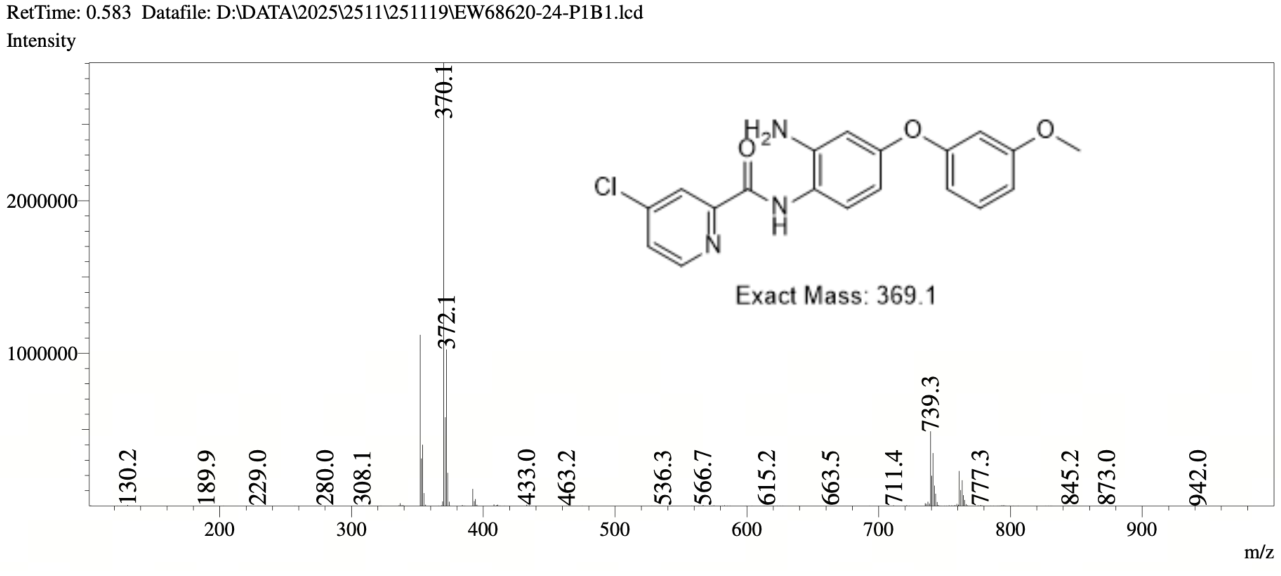

**MS Spectrum of PCSK9-compound-7**

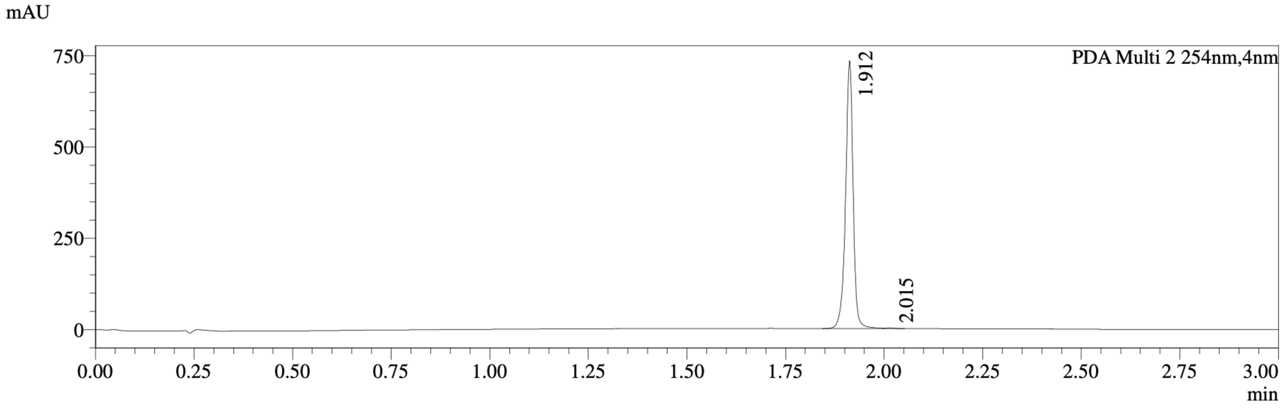

PDA Ch2 254nm

| Peak# | Ret. Time | Width | Height | Height% | Area | Area% |
|---|---|---|---|---|---|---|
| 1 | 1.912 | 0.035 | 733344 | 99.822 | 974225 | 99.794 |
| 2 | 2.015 | 0.076 | 1305 | 0.178 | 2007 | 0.206 |

**HPLC Chromatogram of PCSK9-compound-7**

##### *7.2.8 PCSK9-compound-8*

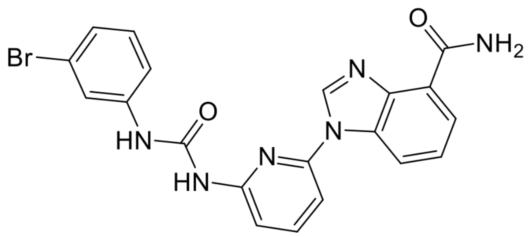

**PCSK9-compound-8**. 1-(6-(3-(3-bromophenyl)ureido)pyridin-2-yl)-1H-benzo[d]imidazole-4-car boxamide

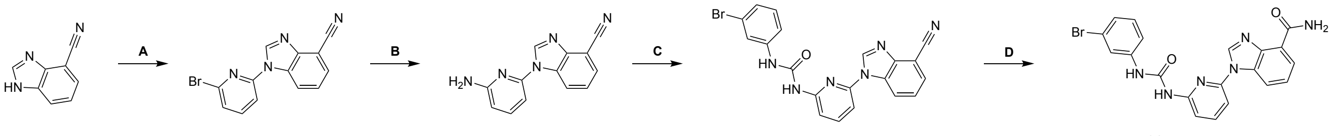

Reagents and conditions: A. 2-bromo-6-fluoro-pyridine, $K_2CO_3$, DMSO, 60 °C, 73%; B. diphenylmethanimine, t-BuONa, $Pd_2(dba)_3$, BINAP, toluene, 110 °C, 61%; C. 1-bromo-3-isocyanato-benzene, DCM, 25 °C, 79%; D. NaOH, THF, 80 °C.

**Synthesis of PCSK9-compound-8**

Step A: 1-(6-bromopyridin-2-yl)-1H-1,3-benzodiazole-4-carbonitrile: To a solution of 1H-benzimidazole-4-carbonitrile (1.00 g, 6.99 mmol, 1.0 equiv) and 2-bromo-6-fluoro-pyridine (1.84 g, 10.4 mmol, 1.5 equiv) in methylsulfinyl methane (10.0 mL) was added potassium carbonate (2.90 g, 20.9 mmol, 3.0 equiv), the reaction mixture was stirred at 60 °C for 12 hours under nitrogen. After completion of the reaction, the reaction mixture was filtered, and the filter cake was washed with ethyl acetate (100 mL) and water (200 mL). The filter cake was dried over under reduced pressure to afford 1-(6-bromopyridin-2-yl)-1H-benzo[d]imidazole-4-carbonitrile (1.60 g, 73.6% yield) as yellow solid; LCMS-Method 1 $[M+1, M+3]^+ = 299.0, 301.0$.

Step B: 1-(6-aminopyridin-2-yl)-1H-benzo[d]imidazole-4-carbonitrile: To a mixture of 1-(6-bromopyridin-2-yl)-1H-benzo[d]imidazole-4-carbonitrile (1.50 g, 5.01 mmol, 1.0 equiv), diphenylmethanimine (908 mg, 5.01 mmol, 1.0 equiv), Pd2(dba)3 (229 mg, 250 $\mu$mol, 0.050 equiv) and BINAP (312 mg, 501 $\mu$mol, 0.10 equiv) in in toluene (15.0 mL) was added sodium tert-butoxide (530 mg, 5.52 mmol, 1.1 equiv), the reaction mixture was stirred at 110 °C for 6 hours under nitrogen. After completion of the reaction, the reaction mixture was quenched with water (100 mL) and filtered through a pad of celite, the cake was washed with ethyl acetate (100 mL). The organic phases was washed with brine (100 mL $\times$ 2), dried over anhydrous sodium sulfate, filtered and concentrated under reduced pressure to give a residue. The residue was purified by reversed-phase flash (0.1% Formic acid condition) to afford 1-(6-aminopyridin-2-yl)-1H-benzo[d]imidazole-4-carbonitrile (720 mg, 61.0% yield) as yellow solid; LCMS-Method 1 $[M+1]^+ = 236.1$.

Step C: 1-(3-bromophenyl)-3-(6-(4-cyano-1H-benzo[d]imidazol-1-yl)pyridin-2-yl)urea: To a solution of 1-(6-aminopyridin-2-yl)-1H-benzo[d]imidazole-4-carbonitrile (200 mg, 850. $\mu$mol, 1.0 equiv) in dichloromethane (2.00 mL) was added 1-bromo-3-isocyanato-benzene (252 mg, 1.28 mmol, 1.5 equiv). The reaction mixture was stirred at 25 °C for 12 hours under nitrogen. After completion of the reaction, the mixture was filtered, the filter cake was washed with dichloromethane (30.0 mL) and the filter cake was dried under reduced pressure to afford 1-(3-bromophenyl)-3-(6-(4-cyano-1H-benzo[d]imidazol-1-yl)pyridin-2-yl)urea (338 mg, 79.5% yield) as off-white solid; LCMS-Method 1 $[M+1, M+3]^+ = 433.1, 435.1$.

Step D: PCSK9-compound-8: To a solution of 1-(3-bromophenyl)-3-(6-(4-cyano-1H-benzo[d]imidazol-1-yl)pyridin-2-yl)urea (160 mg, 369 $\mu$mol, 1.0 equiv) in tetrahydrofuran (1.60 mL) was added sodium hydroxide (118 mg, 25% purity, 2.0 equiv). The reaction mixture was stirred at 80 °C for 12 hours under nitrogen. After completion of the reaction, the mixture was quenched with water (20.0 mL) and extracted with ethyl acetate (50.0 mL). The combine organic layers were washed with brine (30.0 mL), dried over anhydrous sodium sulfate, filtered and concentrated under reduced pressure to a residue. The residue was purified by prep-HPLC (column: Phenomenex Luna C18 150*25mm*10$\mu$m;mobile phase: [EtOH(0.1% TFA)-ACN]; gradient:25%-55% B over 15.0 min) to afford 1-(6-(3-(3-bromophenyl)ureido)pyridin-2-yl)-1H-benzo[d]imidazole-4-carboxamide (10.5 mg) as off-white solid; 1H NMR (400 MHz, DMSO-d6) $\delta$ = 9.63 (s, 1H), 9.51 (s, 1H), 9.18 (s, 1H), 9.11 (br d, $J$ = 2.0 Hz, 1H), 8.57 (dd, $J$ = 1.2 Hz, $J$ = 8.0 Hz, 1H), 8.10 - 8.00 (m, 2H), 7.91 - 7.83 (m, 3H), 7.61 (d, $J$ = 7.6 Hz, 1H), 7.52 (t, $J$ = 7.6 Hz, 1H), 7.34 - 7.18 (m, 3H); LCMS $[M+1, M+3]^+ = 451.1, 453.1$; HPLC Rt = 1.731 min.

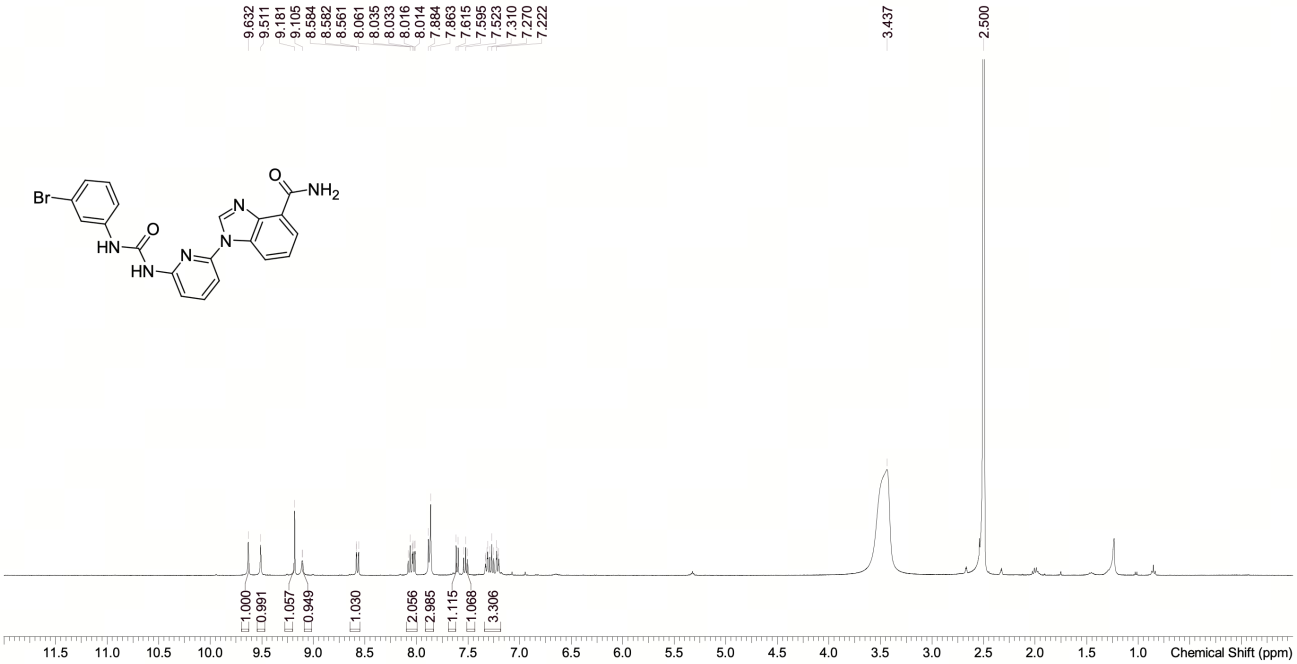

**NMR Spectrum of PCSK9-compound-8**

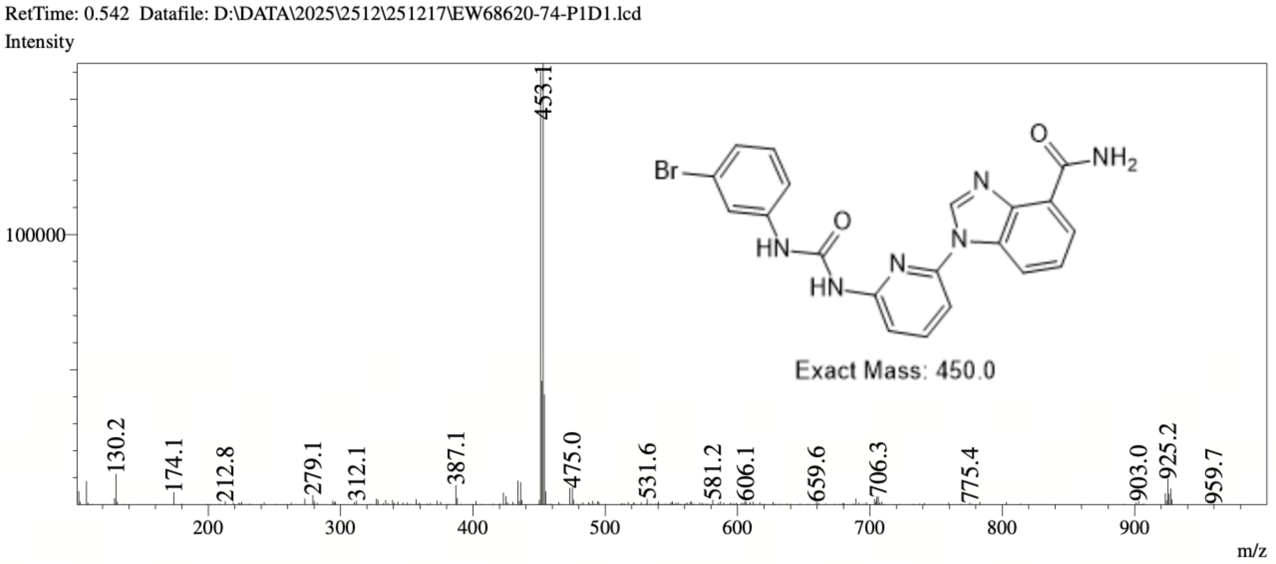

**MS Spectrum of PCSK9-compound-8**

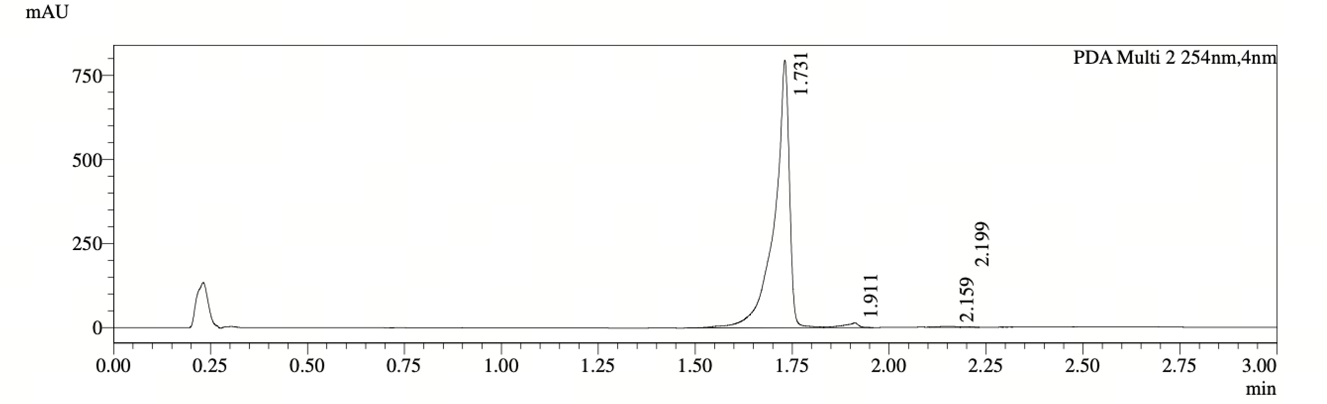

PDA Ch2 254nm

| Peak# | Ret. Time | Width | Height | Height% | Area | Area% |
|---|---|---|---|---|---|---|
| 1 | 1.731 | 0.051 | 794072 | 97.819 | 2135549 | 97.845 |
| 2 | 1.911 | 0.075 | 13771 | 1.696 | 36742 | 1.683 |
| 3 | 2.159 | 0.095 | 2538 | 0.313 | 7989 | 0.366 |
| 4 | 2.199 | 0.060 | 1400 | 0.172 | 2311 | 0.106 |

**HPLC Chromatogram of PCSK9-compound-8**

##### *7.2.9 PCSK9-compound-9*

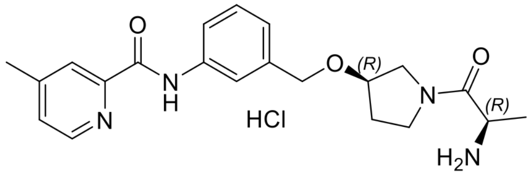

**PCSK9-compound-9**. N-[3-([(3R)-1-[(2R)-2-aminopropanoyl]pyrrolidin-3-yl]oxymethyl)phenyl]-4-methylpyridine-2-carboxamidehydrochloride

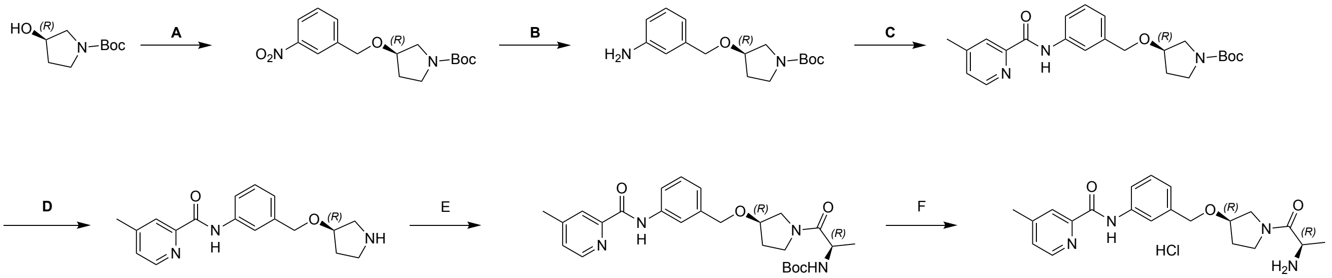

PCSK9-compound-9

Reagents and conditions: A. 1-(bromomethyl)-3-nitro-benzene, NaH, THF, 0 to 25 °C, 76%; B. 1%Pt/C, $H_2$, 1 MPa, 50 °C, 48%; C. 4-methylpyridine-2-carboxylic acid, EDCI, HOBt, TEA, DCM, 50 °C, 70%; D. TAF, DCM, 25 °C, 84%; E. (2R)-2-(tert-butoxycarbonylamino) propanoic acid, EDCI, HOBt, TEA, DCM, 50 °C, 75%; F. HCl/dioxane, 25 °C, 68%.

**Synthesis of PCSK9-compound-9**

Step A: (R)-tert-butyl3-((3-nitrobenzyl)oxy)pyrrolidine-1-carboxylate: To a solution of tert-butyl(3R)-3-hydroxypyrrolidine-1-carboxylate (1.00 g, 5.34 mmol, 1.0 equiv) in tetrahydrofuran (10.0 mL) was added sodium hydride (320 mg, 8.01 mmol, 60% purity, 1.5 equiv) at 0 °C, the mixture was stirred at 0 °C for 0.5 hours. Then a solution of 1-(bromomethyl)-3-nitro-benzene (1.38 g, 6.41 mmol, 1.2 equiv) in tetrahydrofuran (10.0 mL) was added to the above solution at 0 °C. The reaction mixture was stirred at 25 °C for 2 hours under nitrogen. After completion of the reaction, the mixture was quenched by ammonium chloride (20.0 mL) at 0 °C and extracted with ethyl acetate (50.0 mL × 2). The combined organic layers were washed with brine (50.0 mL × 2), dried over anhydrous sodium sulfate, filtered and concentrated under reduced pressure to give a residue. The residue was purified by column chromatography (silicon dioxide, commercial hexanes/ethyl acetate = 100/1 to 0/1) to afford (R)-tert-butyl 3-((3-nitrobenzyl)oxy)pyrrolidine-1-carboxylate (1.43 g, 76.2% yield) as yellow liquid; LCMS-Method 1 $[M-55]^+$ = 267.1.

Step B. tert-butyl(3R)-3-[(3-aminophenyl)methoxy]pyrrolidine-1-carboxylate: Solution 1: ( R)-tert-butyl3-((3-nitrobenzyl)oxy)pyrrolidine-1-carboxylate (1.20 g, 3.72 mmol, 1.0 equiv) in tetrahydrofuran (24.0 mL). The fixed bed (named FLR1, volume 5 mL) was completely packed with granular catalyst WXSC1017 (1%Pt/C), 2.68 g. The hydrogen back pressure regulator was adjusted to 1 MPa and the hydrogen flow rate was 9.0 equiv, 11.5 sccm. The solution 1 was pumped by Pump 1 S1, P1, 0.4 mL/min to flow reactor 1 FLR1, SS, Fixed bed, 6.35 (1/4") mm, 5.00 mL, 50 °C. The reaction mixture was continuously collected from the reactor outlet into the container. The mixture was collected with a bottle. After completion of the reaction, the mixture was concentrated under reduced pressure to afford tert-butyl (3R)-3-[(3-aminophenyl)methoxy]pyrrolidine-1-carboxylate e (880 mg, 48.5% yield) as yellow oil.

Step C: tert-butyl(3R)-3-[3-(4-methylpyridine-2-amido)phenyl]methoxypyrrolidine-1-carboxyl ate: To a solution of tert-butyl(3R)-3-[(3-aminophenyl)methoxy]pyrrolidine-1-carboxylate (880 mg, 3.01 mmol, 1.0 equiv) and 4-methylpyridine-2-carboxylic acid (412 mg, 3.01 mmol, 1.0 equiv) in dichloromethane (8.8 mL) were added 3-(((ethylimino)methylene)amino)-N,N-dimethylpropan-1-amine hydrochloride (865 mg, 4.51 mmol, 1.5 equiv), 1H-benzo[d][1,2,3]triazol-1-ol (488 mg, 3.61 mmol, 1.2 equiv) and triethylamine (609 mg, 6.02 mmol, 2.0 equiv). The reaction mixture was stirred at 50 °C for 2 hours under nitrogen. After completion of the reaction, the mixture was quenched with water (20.0 mL) and extracted with dichloromethane (25.0 mL × 3). The combined organic layers were washed with brine (25.0 mL × 2), dried over anhydrous sodium sulfate, filtered and concentrated under reduced pressure to give a residue. The residue was purified by column chromatography (silicon dioxide, commercial hexanes/ethyl acetate = 15/1 to 5/1) to afford tert-butyl (3R)-3-[3-(4-methylpyridine-2-amido)phenyl]methoxypyrrolidine-1-carboxylate (870 mg, 70.2% yield) as yellow oil; LCMS-Method 1 $[M+23]^+$ = 434.2.

Step D: 4-methyl-N-(3-[(3R)-pyrrolidin-3-yloxy]methylphenyl)pyridine-2-carboxamide: To a solution of tert-butyl (3R)-3-[3-(4-methylpyridine-2-amido)phenyl]methoxypyrrolidine-1-carboxylate (850 mg, 2.07 mmol, 1.0 equiv) in dichloromethane (8.50 mL) was added trifluoroacetic acid (4.25 mL). The reaction mixture was stirred at 25 °C for 2 hours under nitrogen. After completion of the reaction, the mixture was quenched with sodium hydrogen carbonate solution (50.0 mL) and extracted with dichloromethane (25.0 mL × 2). The combined organic layers were washed with brine (50.0 mL × 2), dried over anhydrous sodium sulfate, filtered and concentrated under reduced pressure to afford 4-methyl-N-(3-[(3R)-pyrrolidin-3-yloxy]methylphenyl)pyridine-2-carboxamide (550 mg, 84.4% yield) as yellow oil; LCMS-Method 1 $[M+1]^+$ = 312.1.

Step E: tert-butylN-[(2R)-1-[(3R)-3-[3-(4-methylpyridine-2-amido)phenyl]methoxypyrrolidin-1-y l]-1-oxopropan-2-yl]carbamate: To a solution of 4-methyl-N-(3-[(3R)-pyrrolidin-3-yloxy]methylphen yl)pyridine-2-carboxamide (100 mg, 321 $\mu$mol, 1 equiv) and (2R)-2-(tert-butoxycarbonylamino) propanoic acid (60.7 mg, 321 $\mu$mol, 1.0 equiv) in dichloromethane (1.00 mL) were added 3-(((ethylimino)methylene)amino)-N,N-dimethylpropan-1-amine hydrochloride (92.3 mg, 481 $\mu$mol, 1.5 equiv), 1H-benzo[d][1,2,3]triazol-1-ol (52.0 mg, 385 $\mu$mol, 1.2 equiv) and triethylamine (64.9 mg, 642 $\mu$mol, 2.0 equiv). The reaction mixture was stirred at 50 °C for 2 hours under nitrogen.

After completion of the reaction, the mixture was quenched with water (3.00 mL) and extracted with dichloromethane (3.00 mL × 3). The combined organic layers were washed with brine (2.50 mL × 2), dried over anhydrous sodium sulfate, filtered and concentrated under reduced pressure to give a residue. The residue was purified by prep-TLC (commercial hexanes/ethyl acetate = 0/1) to afford tert-butyl N-[(2R)-1-[(3R)-3-[3-(4-methylpyridine-2-amido)phenyl]methoxypyrrolidin-1-yl]-1-oxopropan-2-yl]carbamate (120 mg, 75.8% yield) as colorless gum; LCMS-Method 1 $[M+1]^+$ = 483.3.

Step F: PCSK9-compound-9: A solution of t tert-butylN-[(2R)-1-[(3R)-3-[3-(4-methylpyridine-2-amido)phenyl]methoxypyrrolidin-1-yl]-1-oxopropan-2-yl]carbamate (120 mg, 248 $\mu$mol, 1.0 equiv) in HCl/dioxane (4 M, 2.4 mL) was stirred at 25 °C for 2 hours under nitrogen. After completion of the reaction, the mixture was concentrated under reduced pressure to afford 30A (71.99 mg, 68.8% yield) as yellow solid; $^1$H NMR (400 MHz, Dimethyl sulfoxide-d6) $\delta$ = 10.77 (d, $J$ = 4.8 Hz, 1H), 8.62 (d, $J$ = 2.8 Hz, 1H), 8.35 (d, $J$ = 14.0 Hz, 3H), 8.17 (d, $J$ = 7.2 Hz, 1H), 7.92 (s, 1H), 7.80 (d, $J$ = 5.6 Hz, 1H), 7.59 (s, 1H), 7.34 (t, $J$ = 7.6 Hz, 1H), 7.09 (d, $J$ = 6.8 Hz, 1H), 4.59 - 4.47 (m, 2H), 4.31 - 4.16 (m, 1H), 4.13 - 3.95 (m, 1H), 3.69 - 3.46 (m, 3H), 3.44 - 3.32 (m, 1H), 2.48 (s, 3H), 2.20 - 1.88 (m, 2H), 1.34 (dd, $J$ = 6.4, 12.8 Hz, 3H); LCMS-Method 4 $[M+1]^+$ = 383.2; HPLC- Method 6 Rt = 1.085 min.

**Analytical method by SFC:**

Column: Chiralcel OJ-3 50 × 4.6 mm I.D., 3 $\mu$m;

Mobile phase: Phase A for $CO_2$, and Phase B for IPA (0.05% DEA);

Gradient elution: IPA (0.05%DEA) in CO2 from 5% to 40%;

Flow rate: 4mL/min; Detector: PDA;

Colum Temp: 35 °C; Back Pressure: 100 Bar;

Retention time: 1.218 min.

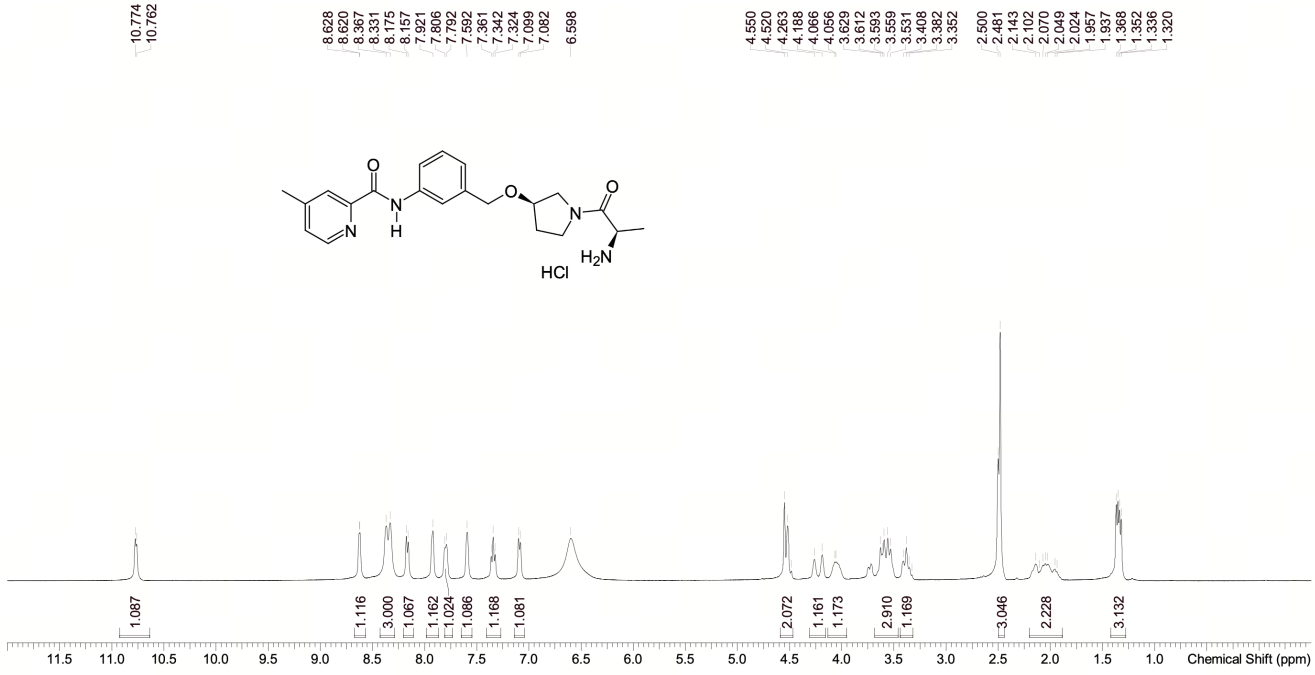

**NMR Spectrum of PCSK9-compound-9**

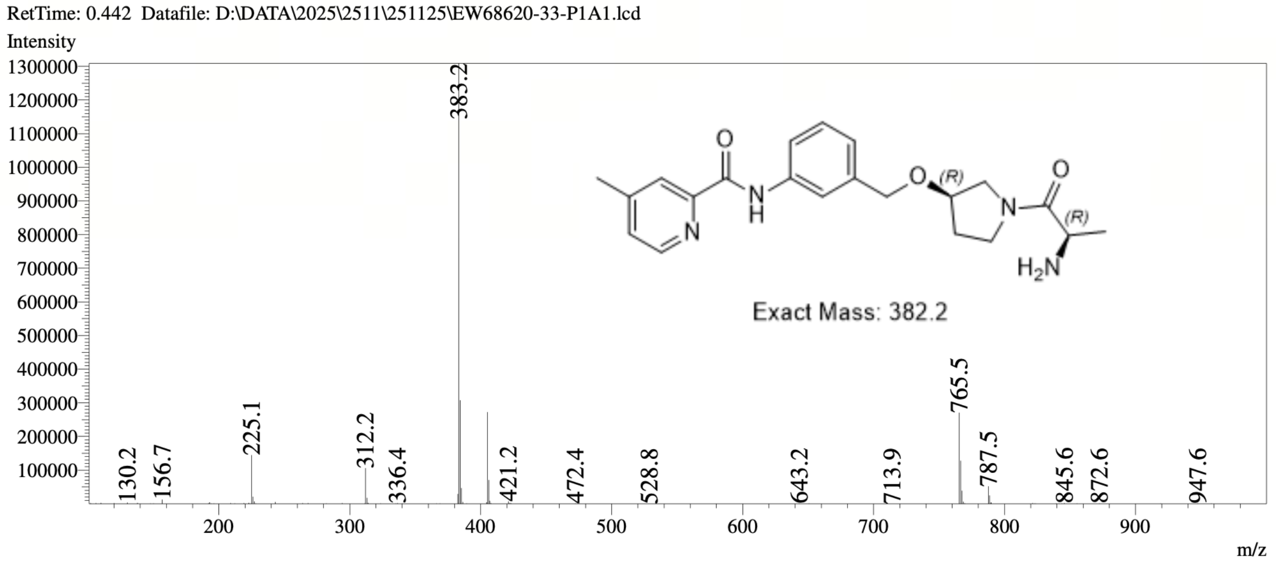

**MS Spectrum of PCSK9-compound-9**

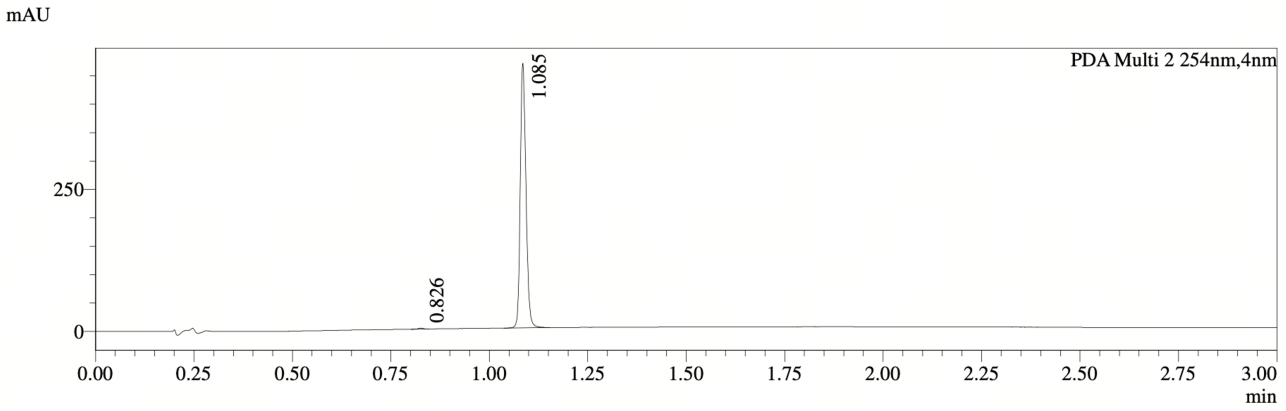

PDA Ch2 254nm

| Peak# | Ret. Time | Width | Height | Height% | Area | Area% |
|---|---|---|---|---|---|---|
| 1 | 0.826 | 0.027 | 1610 | 0.345 | 1561 | 0.335 |
| 2 | 1.085 | 0.028 | 465529 | 99.655 | 464837 | 99.665 |

**HPLC Chromatogram of PCSK9-compound-9**

### Supplementary Figures

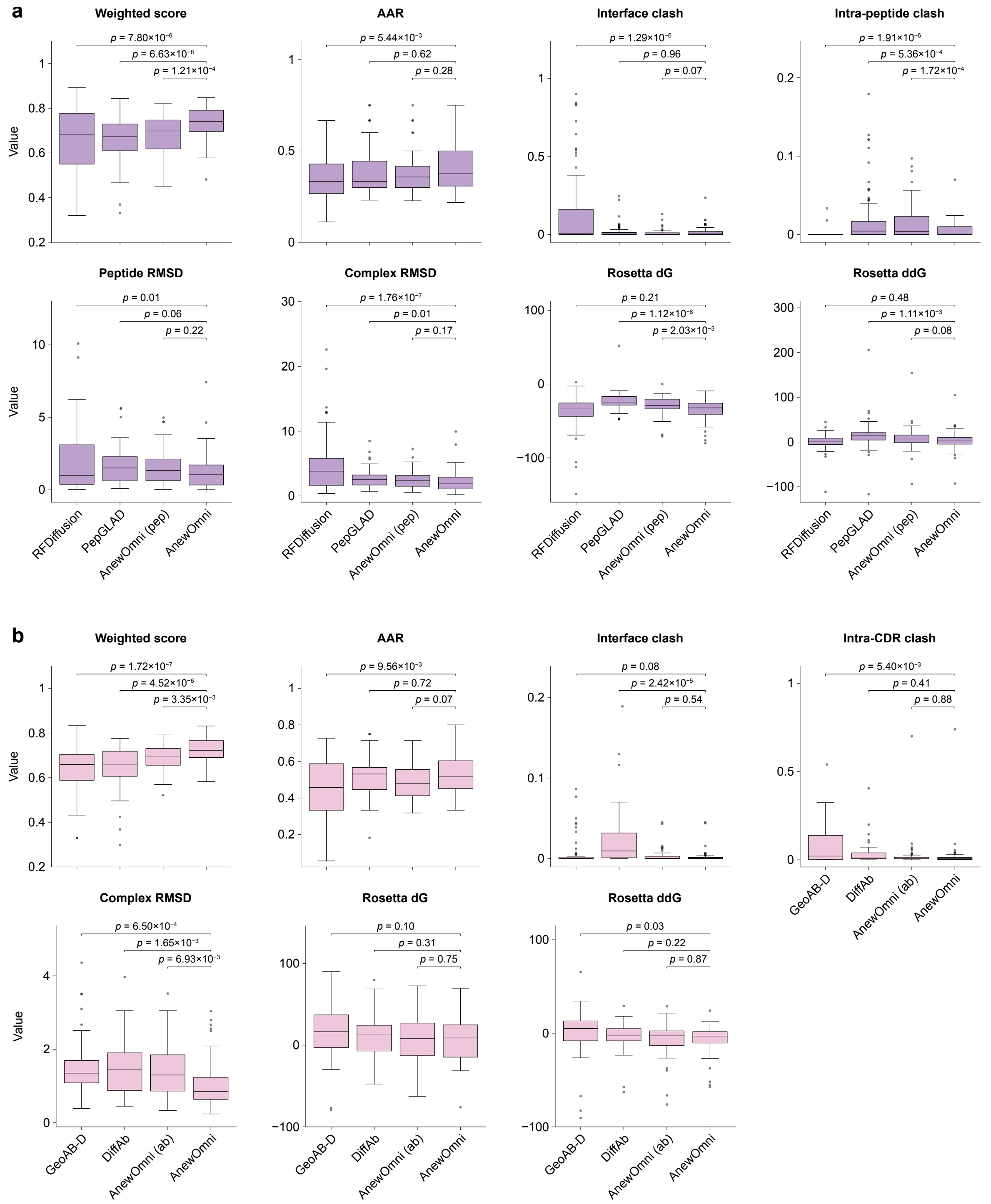

**Supplementary Figure 1.** ***In silico* benchmark for peptide and antibody. a-h**, Overall weighted score, amino acid recovery (AAR), interface clash ratio, intra-peptide clash ratio, root mean square deviation (RMSD) of $C_\alpha$ to the native peptides aligned by the peptide, RMSD of $C_\alpha$ to the native peptide aligned by the target, Rosetta $\Delta$G and Rosetta $\Delta\Delta$G of generated peptides (N=93). Ours (pep-only) denotes the ablation version of our model trained only with peptide data.

**i-o**, Overall weighted score, amino acid recovery (AAR), interface clash ratio, intra-peptide clash ratio, RMSD of $C_\alpha$ to the native CDRs aligned by the target, Rosetta $\Delta$G and Rosetta $\Delta\Delta$G of generated CDRs (N=60).Ours (ab-only) denotes the ablation version of our model trained only with antibody data.

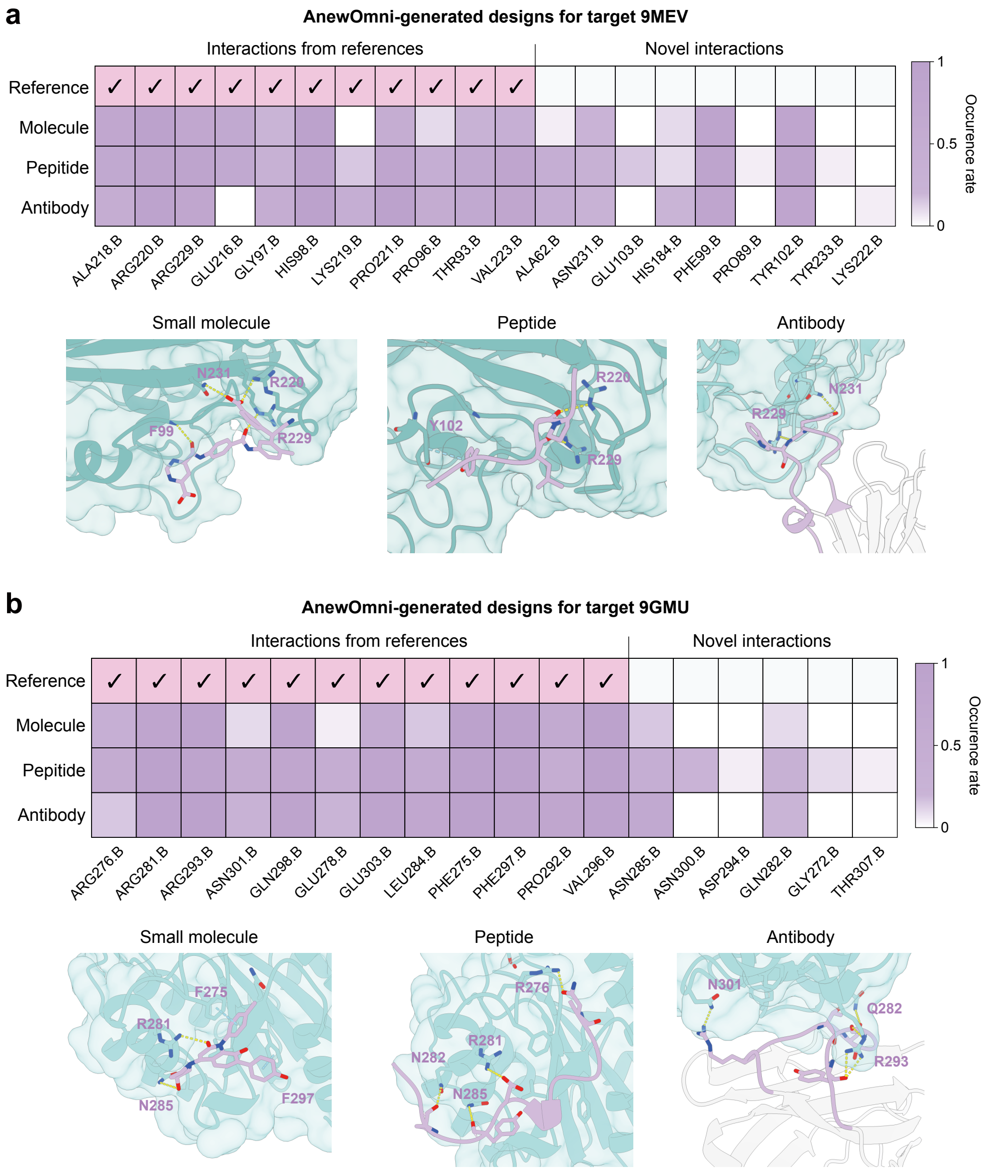

**Supplementary Figure 2. Visualization of small molecule, peptide, and antibody generated for the same binding site**. The targets' PDB code: **a**, 9MEV, **b**, 9GMU. The heatmaps indicate the interaction profile across modalities.

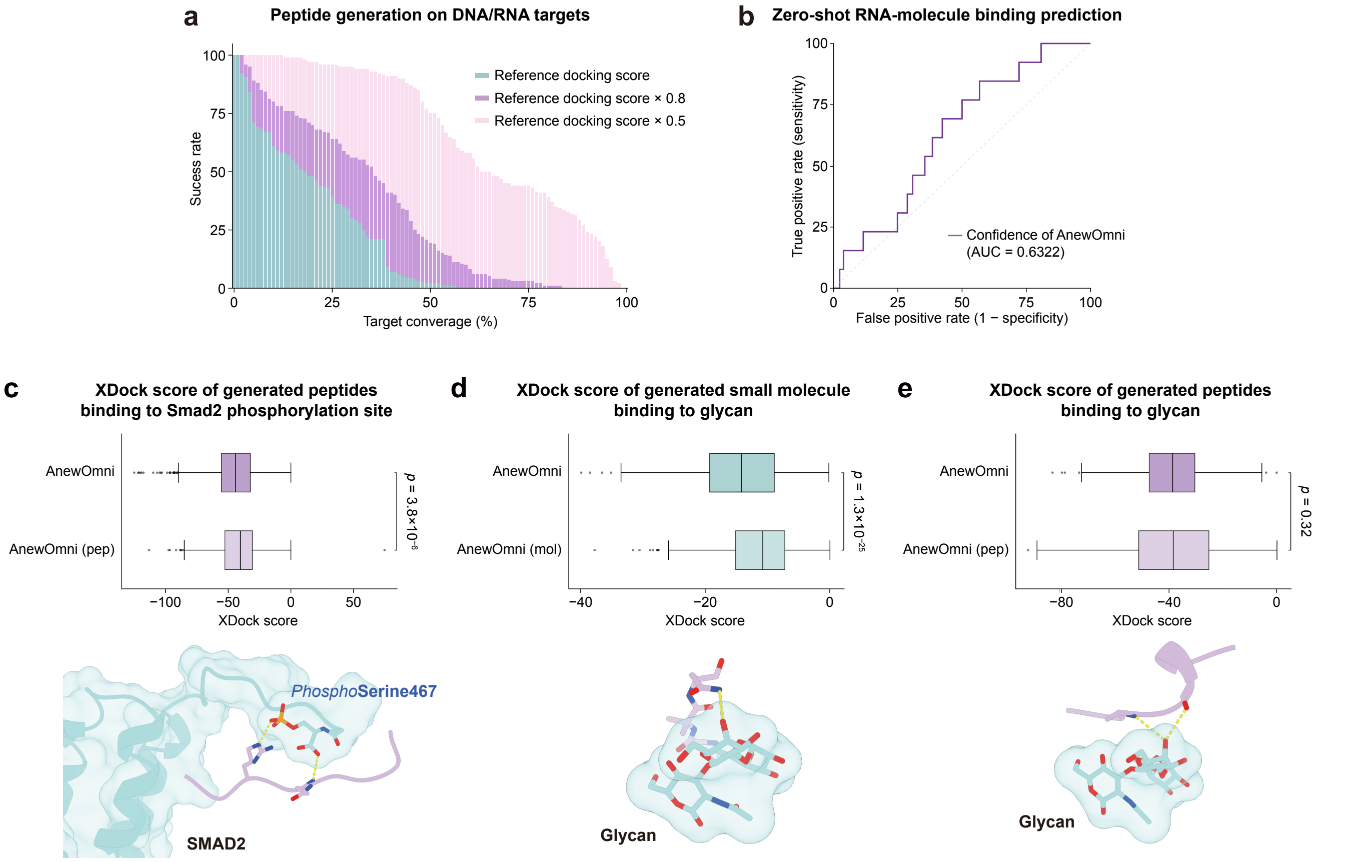

**Supplementary Figure 3.** ***In silico*** **analysis of AnewOmni on non-canonical targets.** **a**, Success rates of peptide generation on 139 DNA/RNA targets, with binding energies evaluated using XDock[41]. Multiple success thresholds are reported, defined as achieving 100%, 80%, or 50% of the native binder's binding energy. **b**, Zero-shot prediction of RNA-binding small molecules via model confidence on a newly reported experimental assay with binary binding labels[42]. **c**, Distribution of XDock scores for peptides generated against a phosphorylation site, along with a representative structural visualization. **d**, Distribution of XDock scores for small molecules generated against a glycan target, along with a representative structural visualization. **e**, Distribution of XDock scores for peptide generated against a glycan target, along with a representative structural visualization.

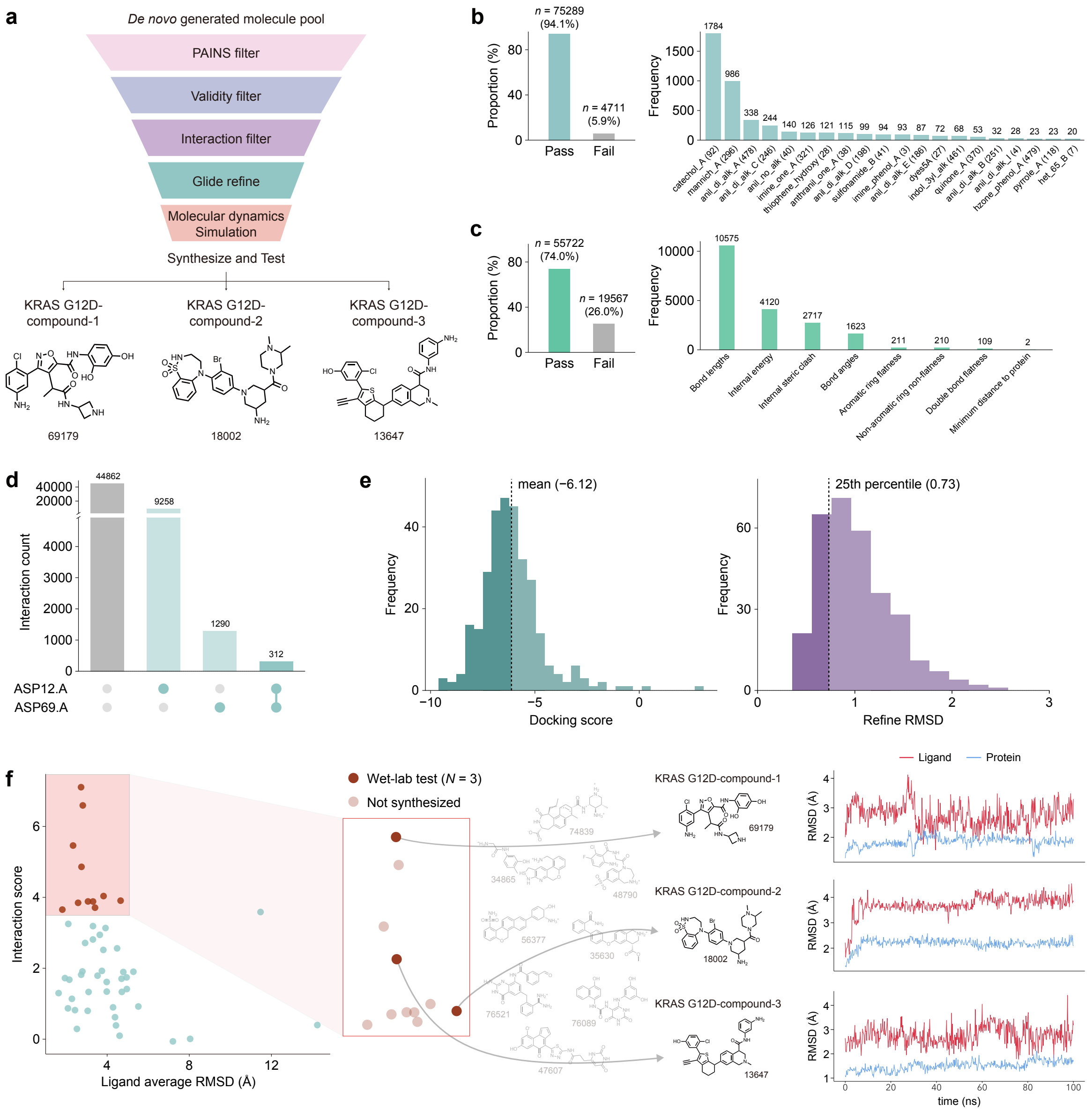

**Supplementary Figure 4. Pipeline of molecule design on KRAS G12D. a**, Overall filter workflow toward *de novo* generated molecule pool. Three compounds passed all criteria and were selected for wet-lab testing. **b**, PAINS filter analysis. Left: Proportion of molecules passing and failing the Pan-Assay Interference Compounds (PAINS) filter. Right: Frequency distribution of the top 20 detected "PAINS substructures". **c**, Validity and geometric filtration summary. Left: Ratio of molecules passing the validity check using PoseBusters. Right: Breakdown of specific failure reasons. **d**, Interaction filter analysis focusing on key residues ASP12 and ASP69. In the UpSetplot, the bars represent the count of molecules for each interaction combination. Solid dots in the matrix denote satisfied interactions at the specific residue. **e**, Statistical distributions of the Glide docking filter. The docking process was performed in 'refine only' mode. Left: distribution of docking scores. Right: the RMSD obtained after structure refinement, the 25th percentile utilized as the selection threshold. **f**, Candidate selection based on Molecular Dynamics (MD) simulations. The candidates with good interaction score and low ligand RMSD were selected to quotation. Finally, three compounds were synthesized and tested. the RMSD trajectories of ligand (sky blue)

and receptor (red) are shown.

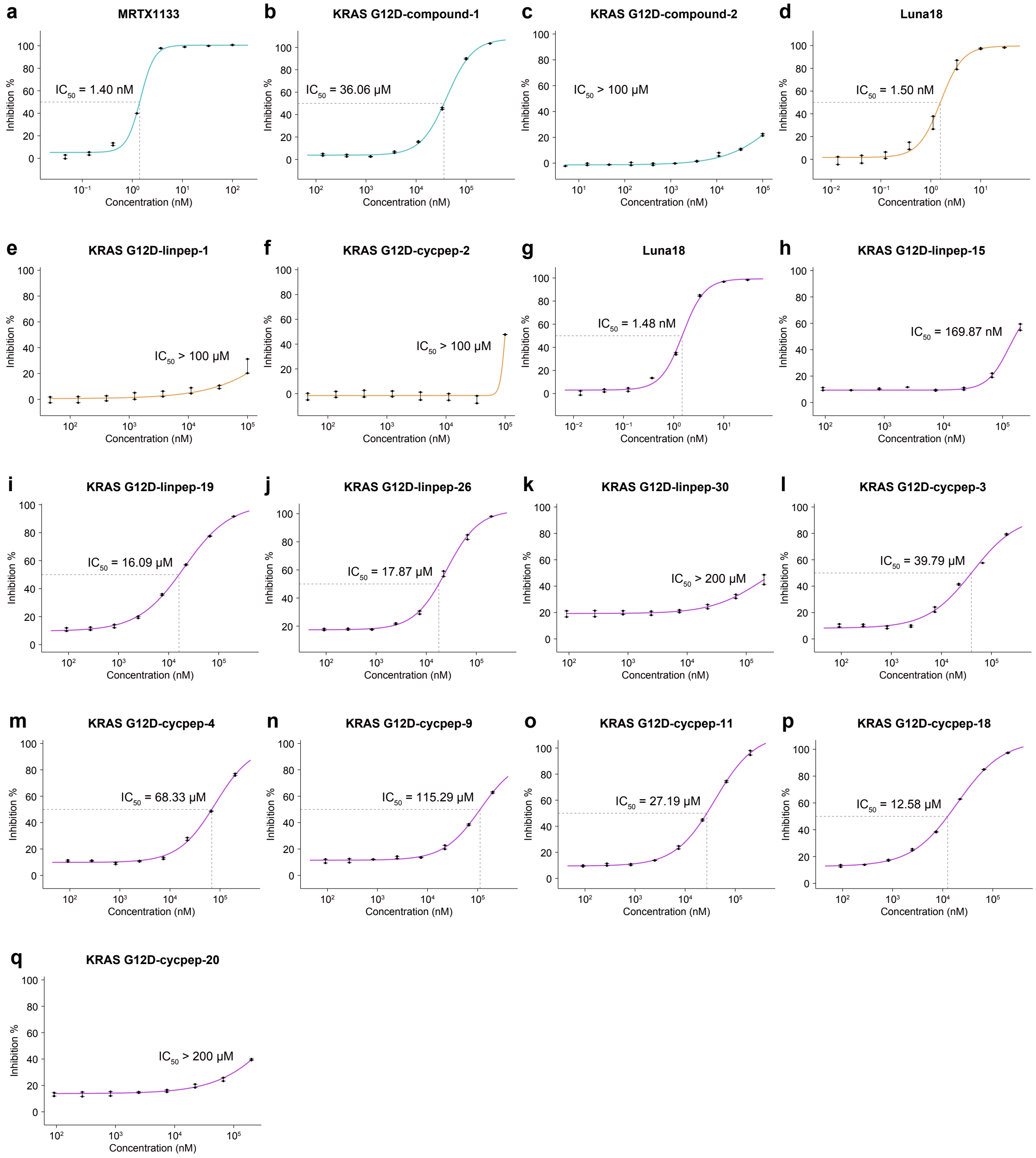

**Supplementary Figure 5. HTRF results for small molecules and peptides on KRAS G12D. a-c**, HTRF results for small molecules on KRAS G12D from two independent experiments, with MRTX1133[70] (synthesized by WuXi AppTec) as the positive control. **d-f**, HTRF results for KRAS G12D-linpep-1 and -2 from two independent experiments conducted along with the dual-point screening (Supplementary Table 1), with LUNA18[71] (from MedChemExpress, Cat. No. HY-156002) as the positive control. **g-q**, HTRF results for the other tested peptides from two independent experiments, with LUNA18[71] (from MedChemExpress, Cat. No. HY-156002) as the positive control.

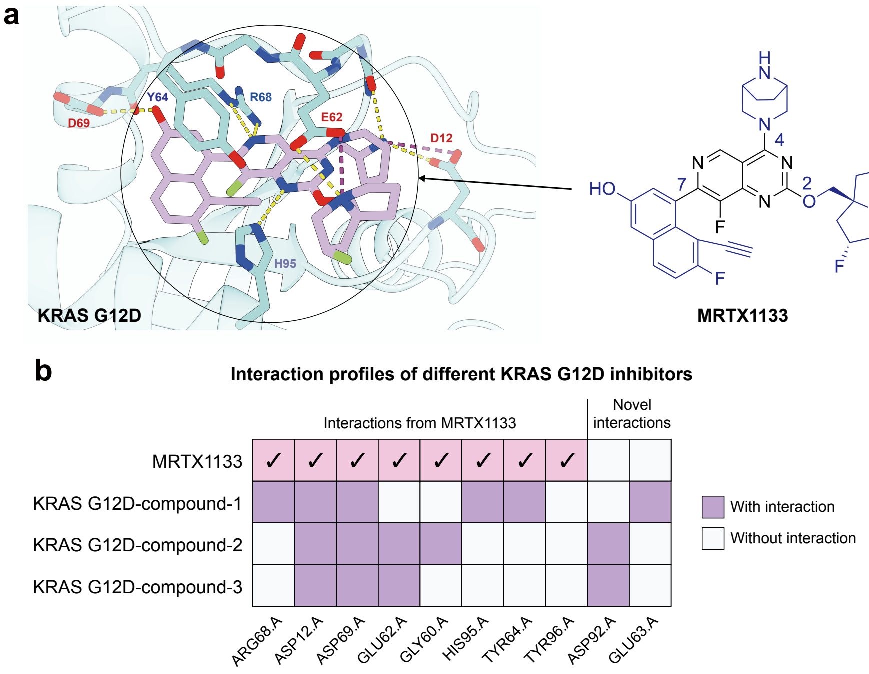

**Supplementary Figure 6. Interaction profile for KRAS G12D inhibitors.** **a**, Interaction profile of MRTX1133[70]. **b**, Comparison of interaction profiles of different KRAS G12D inhibitors.

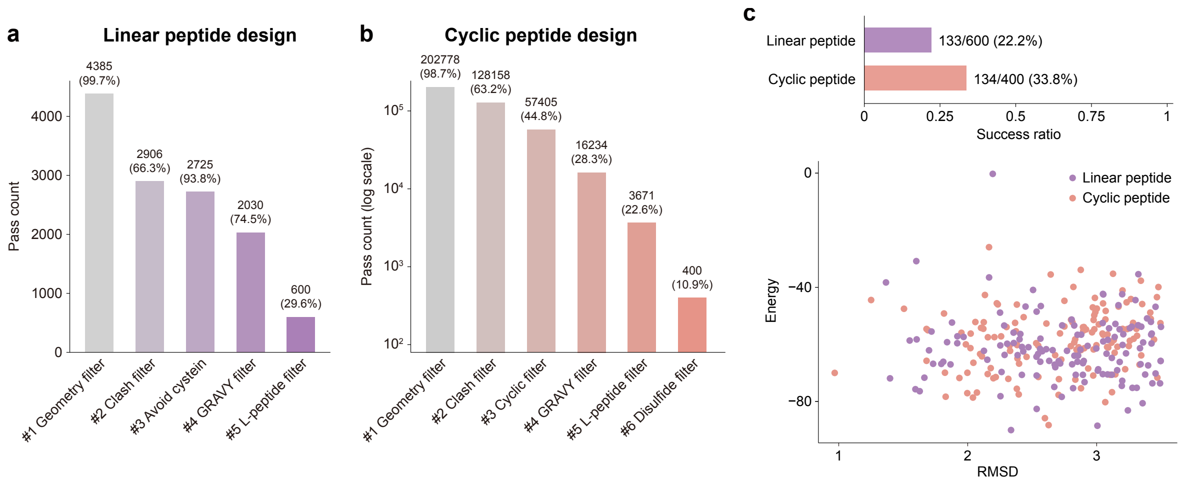

**Supplementary Figure 7. Pipeline of peptide design on KRAS G12D.** **a**, Number of generations passing each filter during linear peptide design targeting KRAS G12D. Generation was terminated once 600 candidates satisfied all filters. **b**, Number of generations passing each filter during cyclic peptide design targeting KRAS G12D. Generation was terminated once 400 candidates satisfied all filters. **c**, Top: Proportion of linear and cyclic peptides exhibiting stable molecular dynamics trajectories (RMSD $< 3.5$ Å). Bottom: Distributions of trajectory RMSD and binding energies given by MM/PBSA[72] are shown for the stable candidates.

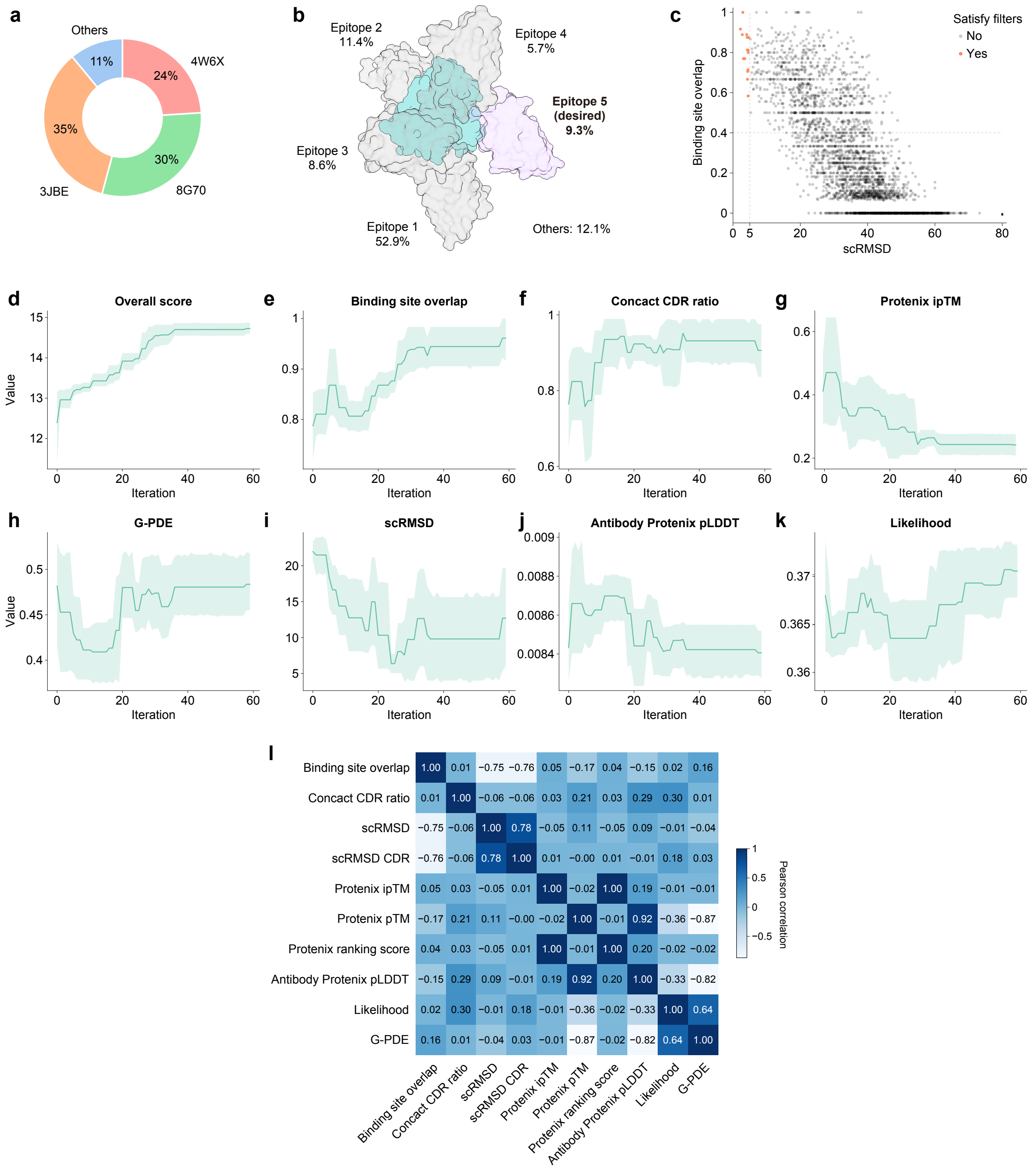

**Supplementary Figure 8. Metrics for antibody pipeline on KRAS G12D during iteration. a**, Distribution of the PDB IDs of the enriched frameworks among all generated candidates. **b**, Representative epitopes and their tendencies in Protenix predictions (N=148), estimated from the initial predictions together with the first 100 candidates during iteration, which approximate random antibodies and therefore reflect the intrinsic bias of Protenix toward KRAS G12D epitopes. **c**, Self-consistent root mean square deviation (scRMSD) versus binding-site overlap for all candidates. Thresholds of scRMSD below 5.0Å and binding-site overlap above 40% are indicated as filtering criteria. **d–k**, Metric trajectories of the top five candidates ranked by the overall weighted score (Supplementary Note 6.1). Iterative ranking progressively enriches structurally plausible conformations for CDR design, specifically those targeting the desired epitope and forming interactions mostly through CDR residues. Structure prediction confidence metrics may remain low because the targeted epitope is intrinsically disfavored in predictions, and the conformational plausibility is prioritized over confidences. **i**, Pearson correlation coefficients between different metrics.

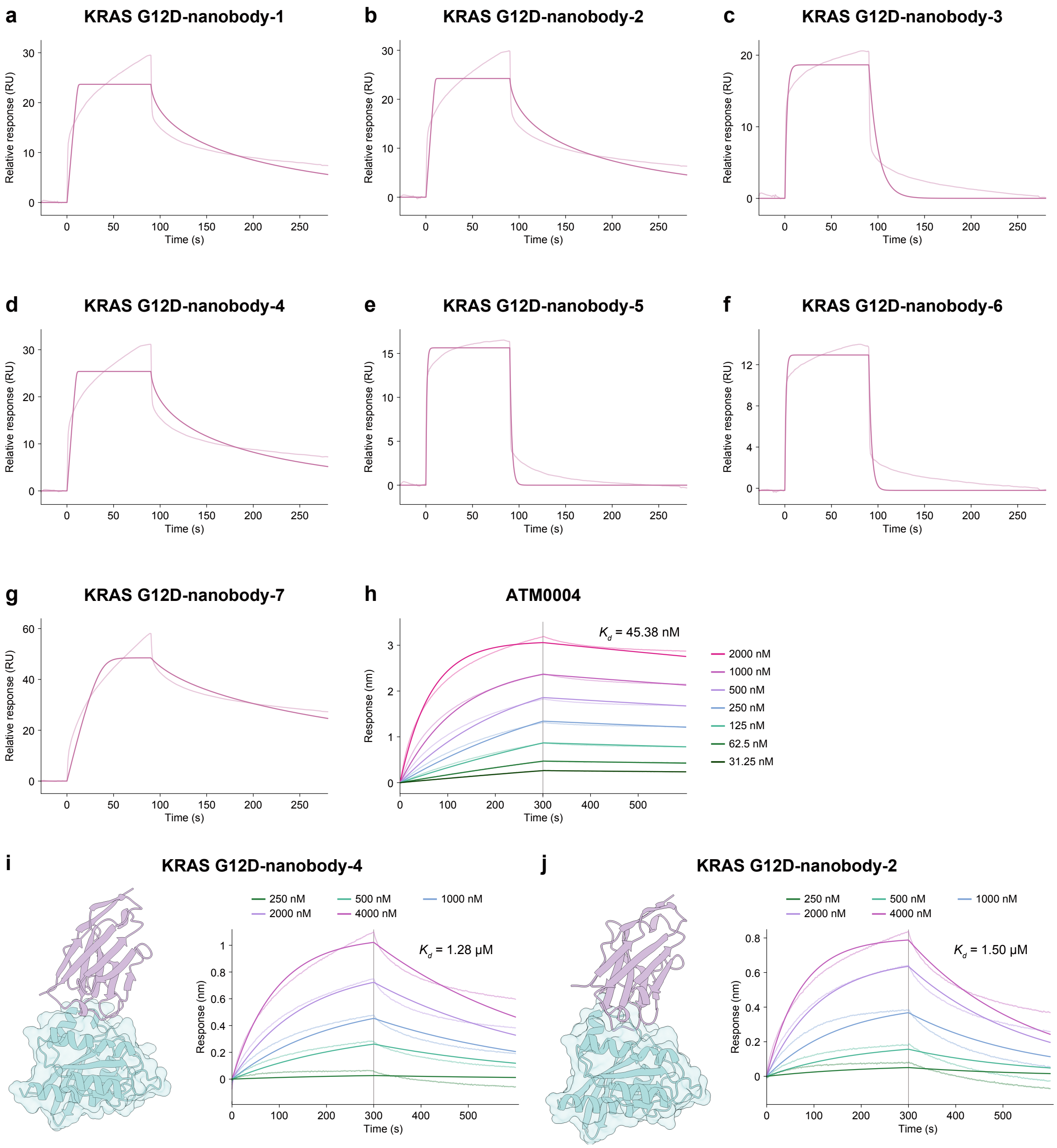

**Supplementary Figure 9. Initial screening and the positive control for nanobodies in complex with KRAS G12D.** Nanobodies were first screened in a single concentration using surface plasma resonance (SPR). Candidates with relative response (RU) above 20 were further validated in multiple concentrations using Bio-layer interferometry (BLI) to derive the exact dissociation constant ($K_d$). **a-g**, RU for the nanobodies in a single concentration of 150 nM using

SPR. **h**, BLI sensorgram of the positive control (from aviva, product number: ATM00004) for nanobodies binding to KRAS G12D. **i-j**, Structural visualization and BLI sensogram of KRAS G12D-nanobody-4 and -2.

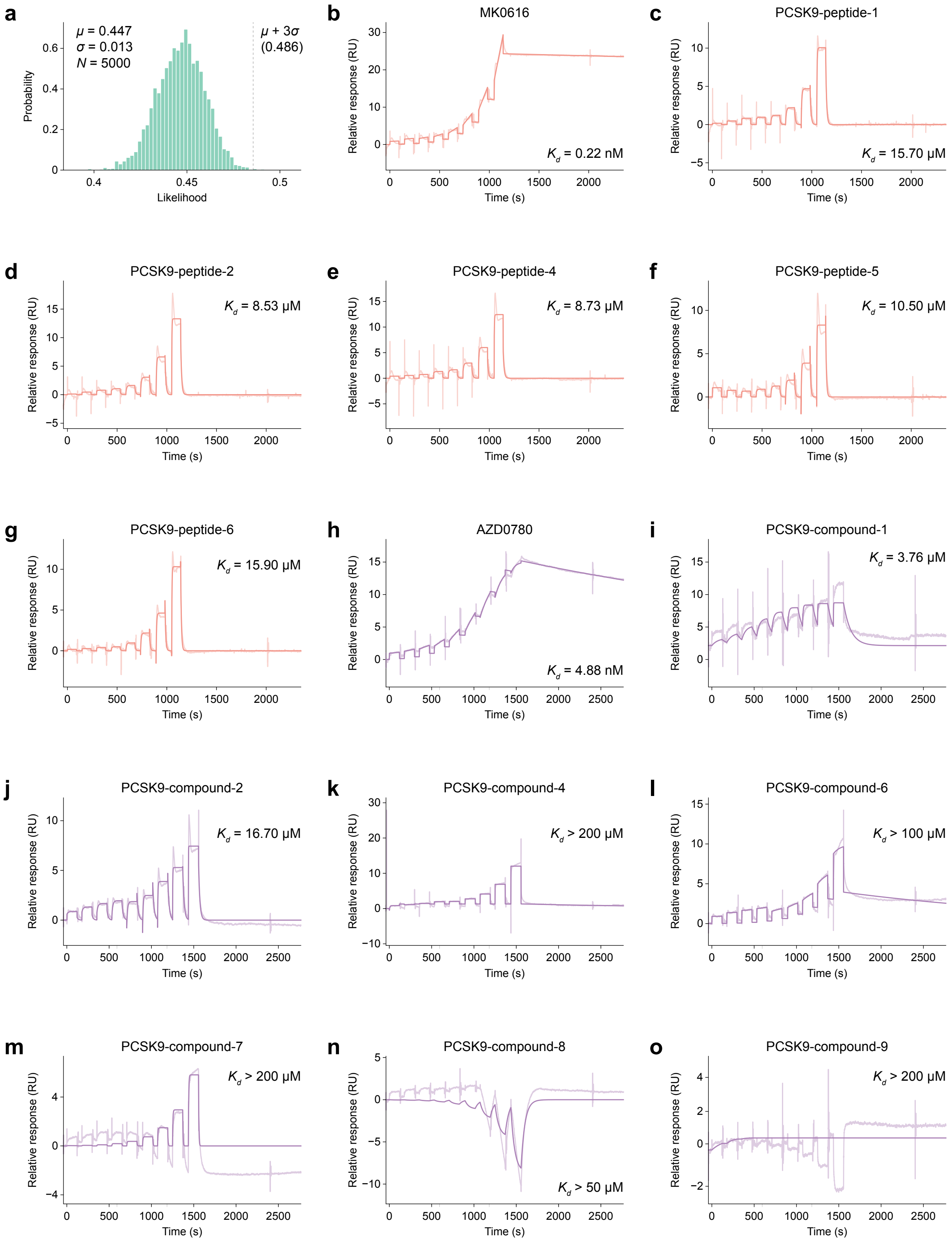

**Supplementary Figure 10.** **Results for PCSK9 inhibitor design.** **a**, Distribution of generative likelihood from all generated peptides on the orthosteric binding site of PCSK9. **b-h**, Binding affinities of tested peptides determined by SPR, with MK0616 (from MedChemExpress, Cat#HY-P4153) as the positive control. **i-p**, Binding affinities of tested small molecules determined by SPR, with AZD0780 (from TOPSCIENCE, TSID: T64362, CAS: 2455427-91-3) as the positive control.

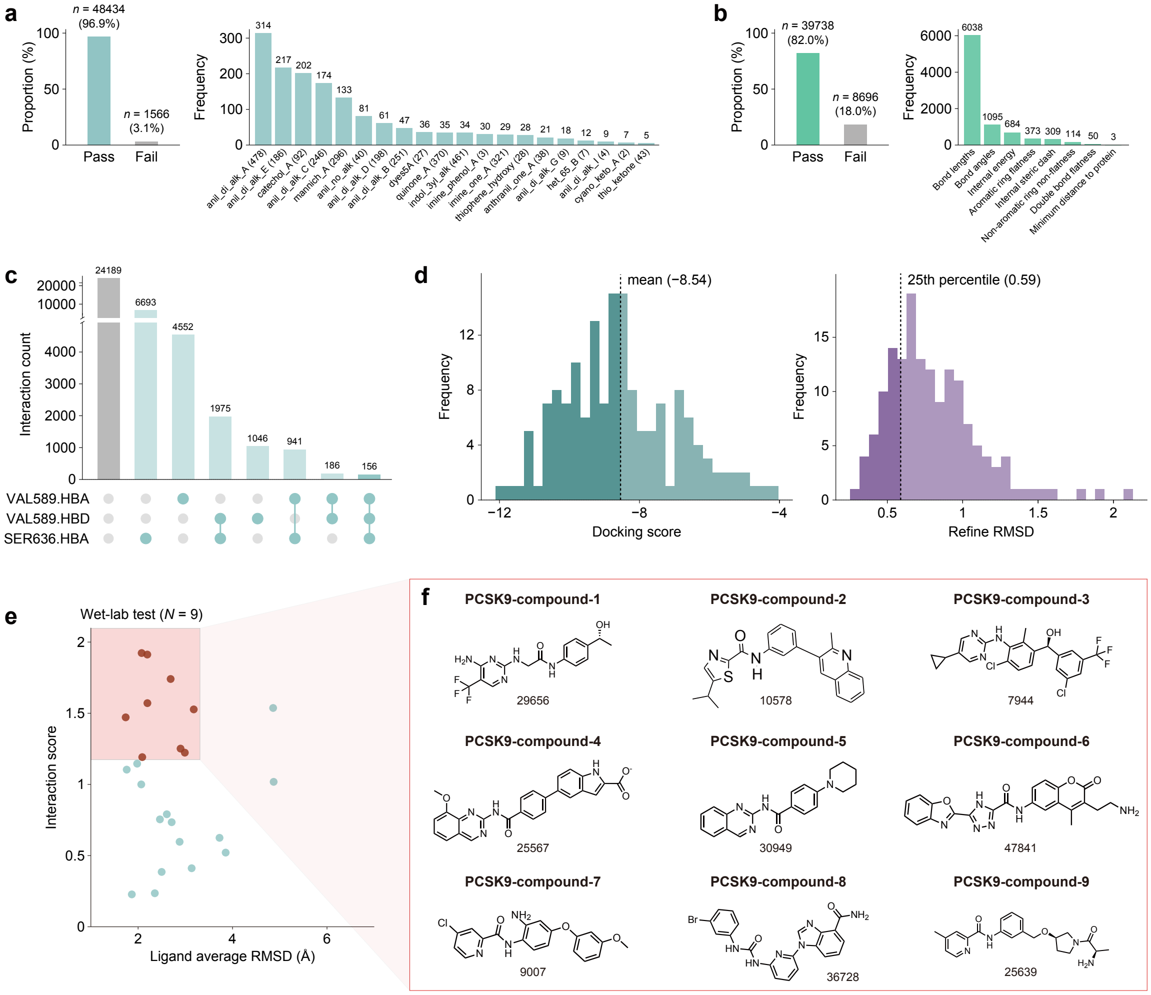

**Supplementary Figure 11.** **Pipeline of molecule design on PCSK9.** **a**, PAINS filter summary. Left: Proportion of molecules passing and failing the PAINS filter. Right: Frequency distribution of the top 20 detected PAINS substructure. **b**, Validity and geometric filtration summary. Left: Ratio of molecules passing the validity check using PoseBusters. Right: Breakdown of specific failure reasons. **c**, Interaction filter result summary. The interaction analysis focusing on key residues VAL589 and SER636. Specifically, both hydrogen bond donor and hydrogen bond acceptor were considered separately via known binding mode. In the UpSetplot, the bars represent the count of molecules for each interaction combination. Solid dots in the matrix denote satisfied interactions at the specific residue. **d**, Statistical distributions of the Glide docking filter. The docking process was performed in 'refine only' mode. Left: distribution of docking scores. Right: the RMSD obtained after structure refinement, the 25th percentile utilized as the selection threshold. **e**, Candidate selection based on Molecular Dynamics (MD) simulations. The candidates with good interaction score and low ligand RMSD were selected as final candidates, 8 compounds were synthesized and tested. **f**, 2D diagrams of all synthesized molecules derived from the pipeline.

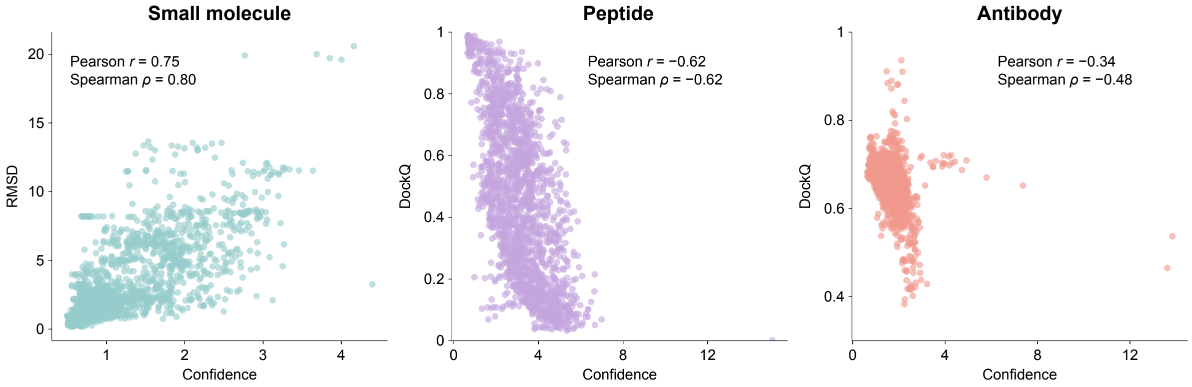

**Supplementary Figure 12. Correlation between confidence and docking metrics**. Evaluation of Pearson and Spearman correlation between model-derived confidence score and key evaluation metrics on the docking tasks, including RMSD for **(a)** the molecule testset (N=100), DockQ for **(b)** the peptide testset (N=93) and **(c)** the antibody testset (N=60), respectively.

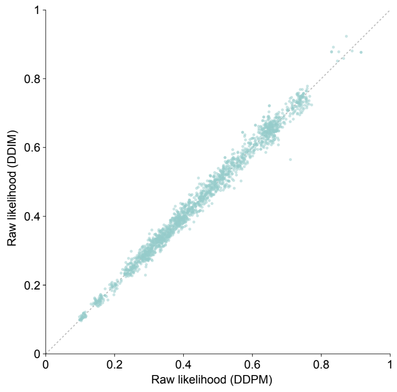

**Supplementary Figure 13. Correlation of likelihood between DDPM and DDIM samplers**. Scatter plot of raw likelihoods between DDPM and DDIM sampling strategies under identical noise conditions (N=1900).

#### Supplementary Tables

**Supplementary Table 1.** Dual-point HTRF screening of KRAS G12D peptides. The table shows the percentage inhibition of various cyclic (cycpep) and linear (linpep) peptides tested at concentrations of 100 $\mu$M and 10 $\mu$M. Data are presented as individual values from two independent experiments (Replicate 1 and 2) and their mean ($n = 2$).

| ID | Inhibition at 100 $\mu$M (%) | | | Inhibition at 10 $\mu$M (%) | | |
|---|---|---|---|---|---|---|
| | Replicate 1 | Replicate 2 | Mean | Replicate 1 | Replicate 2 | Mean |
| KRAS-cycpep-3 | 71.1 | 71.4 | 71.2 | 23.2 | 24.4 | 23.8 |
| KRAS-cycpep-5 | 89.2 | 88.0 | 88.6 | 39.0 | 39.3 | 39.1 |
| KRAS-cycpep-6 | 11.6 | 13.1 | 12.4 | 0.3 | 0.3 | 0.3 |
| KRAS-cycpep-7 | 6.6 | 6.6 | 6.6 | 1.5 | 2.7 | 2.1 |
| KRAS-cycpep-8 | 14.0 | 13.4 | 13.7 | 2.1 | 1.5 | 1.8 |
| KRAS-cycpep-10 | 16.4 | 16.4 | 16.4 | 0.9 | 0.9 | 0.9 |
| KRAS-cycpep-13 | 15.5 | 14.6 | 15.0 | -2.9 | -1.8 | -2.3 |
| KRAS-cycpep-14 | 15.2 | 16.4 | 15.8 | 2.1 | 3.6 | 2.9 |
| KRAS-cycpep-15 | 22.0 | 22.3 | 22.2 | 8.9 | 10.4 | 9.7 |
| KRAS-cycpep-16 | 25.3 | 25.6 | 25.4 | 4.5 | 4.8 | 4.6 |
| KRAS-cycpep-17 | 67.5 | 66.9 | 67.2 | 18.2 | 18.8 | 18.5 |
| KRAS-cycpep-19 | 23.2 | 21.7 | 22.5 | 6.0 | 5.1 | 5.5 |
| KRAS-linpep-2 | 9.2 | 9.2 | 9.2 | 9.2 | 9.5 | 9.4 |
| KRAS-linpep-3 | 72.5 | 75.2 | 73.9 | 7.8 | 7.8 | 7.8 |
| KRAS-linpep-4 | 20.2 | 22.6 | 21.4 | 0.9 | 1.5 | 1.2 |
| KRAS-linpep-5 | 82.7 | 82.9 | 82.8 | 41.3 | 41.6 | 41.5 |
| KRAS-linpep-7 | 14.3 | 14.0 | 14.1 | 8.3 | 10.1 | 9.2 |
| KRAS-linpep-8 | 17.3 | 17.0 | 17.1 | 8.9 | 8.9 | 8.9 |
| KRAS-linpep-9 | 20.8 | 21.1 | 21.0 | 6.6 | 8.3 | 7.5 |
| KRAS-linpep-10 | 12.2 | 13.4 | 12.8 | 9.5 | 11.0 | 10.3 |
| KRAS-linpep-11 | 9.2 | 10.4 | 9.8 | 5.7 | 8.1 | 6.9 |
| KRAS-linpep-12 | 95.7 | 95.7 | 95.7 | 71.4 | 72.8 | 72.1 |
| KRAS-linpep-13 | 14.3 | 15.8 | 15.0 | 6.0 | 13.4 | 9.7 |
| KRAS-linpep-14 | 8.6 | 11.9 | 10.3 | 0.9 | 4.8 | 2.9 |
| KRAS-linpep-16 | 31.2 | 30.3 | 30.8 | 11.6 | 11.9 | 11.8 |
| KRAS-linpep-17 | 76.7 | 77.6 | 77.2 | 13.7 | 15.8 | 14.7 |
| KRAS-linpep-18 | 19.9 | 22.0 | 21.0 | 9.5 | 12.2 | 10.9 |
| KRAS-linpep-19 | 77.9 | 78.8 | 78.3 | 33.3 | 36.6 | 34.9 |
| KRAS-linpep-20 | 23.2 | 25.0 | 24.1 | 8.9 | 10.4 | 9.7 |
| KRAS-linpep-21 | 13.4 | 14.9 | 14.1 | 10.7 | 13.7 | 12.2 |
| KRAS-linpep-22 | 24.4 | 28.3 | 26.3 | 8.6 | 12.5 | 10.6 |
| KRAS-linpep-23 | 15.8 | 20.5 | 18.2 | 2.7 | 8.1 | 5.4 |
| KRAS-linpep-24 | 44.3 | 52.9 | 48.6 | 29.5 | 31.5 | 30.5 |
| KRAS-linpep-25 | 29.5 | 30.9 | 30.2 | 18.5 | 20.5 | 19.5 |
| KRAS-linpep-27 | 17.0 | 20.8 | 18.9 | 15.2 | 18.5 | 16.8 |
| KRAS-linpep-28 | 17.9 | 20.8 | 19.3 | 13.4 | 19.0 | 16.2 |
| KRAS-linpep-29 | 41.6 | 46.7 | 44.2 | 22.6 | 29.2 | 25.9 |
| KRAS-cycpep-11 | 98.4 | 98.7 | 98.6 | 46.4 | 49.7 | 48.0 |
| KRAS-cycpep-12 | -15.4 | -7.1 | -11.3 | -23.5 | -12.8 | -18.1 |
| KRAS-linpep-6 | 24.4 | 23.2 | 23.8 | -16.6 | -2.9 | -9.8 |
| KRAS-linpep-15 | 85.6 | 82.7 | 84.1 | 15.2 | 12.5 | 13.8 |

**Supplementary Table 2.** Library of frameworks of antibodies and nanobodies. CDR regions are marked with -CDRn- (n=1,2,3). For antibodies, the heavy chain and the light chain are marked with "H:" and "L:", respectively.

| PDB | Type | Sequence |
|---|---|---|
| 8FK5 | antibody | H: ERLVESGGGVVQPGSSLRLSCAAS-CDR1-GMHWVRQAPGQGLEWVAFI-CDR2-KYHADSVWGRLSISRDNSKDTLYLQMNSLRVEDTATYFCVR-CDR3-WGKGTTVTVSS<br>L: SALTQPASVSGSPGQSITISC-CDR1-WYQQHPGKAPKVVIY-CDR2-GVSNRFSGSKSGNTASLTISGLQAEDEGDYYC-CDR3-FGTGTKLTVL |
| 7N65 | antibody | H: DIRIAESGGGLVQPGESLRLACEII-CDR1-WTTWVRQAPGKGLEWVADI-CDR2-KKYGPSVTGRFTISRDNGKNLVFLQMNSLRVEDTATYYCAR-CDR3-WGPGTLVTVSS<br>L: SVLTQPPSVSGAPGQRVVISC-CDR1-WYQQSPGKVPRIIIY-CDR2-GVPARFSGSKSGTSASLAITGLQAEDEADYYC-CDR3-FGGGTKV |
| 7UKM | antibody | H: QVQLVESGGGVVQPGRSLRLSCEAS-CDR1-DMHWARQAPGKGLEWVAVI-CDR2-KFYADSVKGRFTISRDNSKNTLYLQMNSLRAEDTAVYYCAR-CDR3-WGQGTTVTVSS<br>L: DNVMTQTPFSLSVTPGQPASISC-CDR1-WYLQKPGQSPQLLIY-CDR2-GVPERFSGSGSGTDFTLKISRVEAEDVGVYYC-CDR3-FGQGTKLEIK |
| 4IOF | antibody | H: EVQLVESGGGLVQPGGSLRLSCAAS-CDR1-SIHWVRQAPGKGLEWVASI-CDR2-TSYADSVKGRFTISADTSKNTAYLQMNSLRAEDTAVYYCAR-CDR3-WGQGTLVTVSS<br>L: TQSPSSLSASVGDRVTITC-CDR1-WYQQKPGKAPKLLIY-CDR2-GVPSRFSGSRSGTDFTLTISSLQPEDFATYYC-CDR3-FGQGTKVEIK |
| 7LLK | antibody | H: EMRLEESGGALVKPGGSLRLSCVAS-CDR1-SLMWVRQAPGKGLQWVSSI-CDR2-ISYAGSVRGRFDVFRDRTQTSLLLNMNKLKTEDTGVYFCAR-CDR3-WGRGTLVTV<br>L: ATQPPSVSVSKGETARITC-CDR1-WYRQRPGQSPVLVMF-CDR2-RIPERFSASDSGDTATLTIAGAQDVDEAEYYC-CDR3-FGGGTKLTVL |
| 7T86 | antibody | H: EVQLVETGGGLVKPGGSLRLSCSAS-CDR1-YMTWIRQAPGKGPEWVSHI-CDR2-IYYADSVRGRFTISRDNAKSSLYLQMDSLQADDTAVYYCAR-CDR3-WGPGTLVTVSS<br>L: EIVLTQSPATLSLSPGERATLSC-CDR1-WYQQKPGQAPRLLIY-CDR2-GIPDRFSGSGSGTDFTLTISRLEPEDFAVYYC-CDR3-FGGGTKLEIK |
| 8FSJ | antibody | H: EVQLLEQSGPEVKKPGDSLRISCKMS-CDR1-WIGWVRQKPGQGLEWMGII-CDR2-TRYGPSFQGQVTISIDKSTSTAYLQWNNVKASDTGIYYCAR-CDR3-WGQGTLVIVSS<br>L: LTLTQSPGTLSLSPGERATLSC-CDR1-WYQQKPGQAPRLLIY-CDR2-GIPDRFSGSGSGTGFTLIISRLEPEDFAVYYC-CDR3-FGQGTRLEIK |
| 6XDG | antibody | H: QVQLVESGGGVVQPGRSLRLSCAAS-CDR1-AMYWVRQAPGKGLEWVAVI-CDR2-KYYADSVKGRFTISRDNSKNTLYLQMNSLRTEDTAVYYCAS-CDR3-WGQGTLVTVSS<br>L: QSALTQPASVSGSPGQSITISC-CDR1-WYQQHPGKAPKLMIY-CDR2-GVSNRFSGSKSGNTASLTISGLQSEDEADYYC-CDR3-FGGGTKLTVL |
| 6X05 | nanobody | QLQLVESGGGSVQAGGSLTLSCTAS-CDR1-GIGWFRQRRGEQREEIAYI-CDR2-PNLGDSVKDRFTISRDKSNGTVYLQMNSLKPEDTAVYYCHG-CDR3-WGQGTQVTVSS |
| 9G5L | nanobody | QVQLQESGGGLVQPGGSLRLSCASA-CDR1-AIGWFRQAPGKEREVVSCI-CDR2-TYYADSVKGRFTISSDNAKNTVYLQMNSLKPEDTAVYYCAA-CDR3-WGQGTQVTVSS |
| 9G1Y | nanobody | QVQLQESGGGLVQPGGSLRLSCAAS-CDR1-AIGWFRQAPGKEREGVSCI-CDR2-TYYADSVKGRFTISSDNAKNTVYLQMNSLKPEDTAVYYCAA-CDR3-WGQGTQVTVSS |
| 7WD2 | nanobody | QVQLQESGGGLVQPGGSLRLTCAPS-CDR1-AIGWFRQAPGKEREGVSCI-CDR2-TYYADSVKGRFTISRDNAKNTVYLQMNSLKPEDTAVYYCAA-CDR3-WGQGTQVTVSS |
| 3JBE | nanobody | QVQLQESGGGSVQAGGSLRLSCAAS-CDR1-CMAWFRQVLGEGREGVAFI-CDR2-MRYADTVKGRFTVSQDKDKNTVYLQMNSLKPEDTAIYYCAA-CDR3-WGQGTQVTVSS |
| 7PAF | nanobody | QVQLVESGGGSVQAGGSLRLSCAAS-CDR1-YLGWFRQAPGKEREGVAAL-CDR2-TYYADSVKGRFTVSLDNAKNTVYLQMNSLKPEDTALYYCAA-CDR3-WGQGTQVTVSA |
| 9G4G | nanobody | QVQLQESGGGLVQPGGSLRLSCVVS-CDR1-AIGWFRQAPGKEREGVSCI-CDR2-TNYADSVKGRFTISRDNAKNTVYLQMNSLKPEDTAVYYCAA-CDR3-WGQGTQVTVSS |

| PDB | Type | Sequence |
|---|---|---|
| 7UIB | nanobody | QLQLVETGGGLVKPGGSLRLSCVVS-CDR1-RMAWVRQAPGKELEWVSSI-CDR2-TYYEDSVKGRFTISTDNAKNTLYLQMSSLKPEDTAVYYCAA-CDR3-WGQGTQVTVSS |
| 9G48 | nanobody | QVQLQESGGGLVQPGGSLRLSCAAS-CDR1-AIGWFRQAPGKEREGVSCI-CDR2-TYYADSVKGRFTISRDNAKNTVYLEMNSLKPEDTAVYYCAA-CDR3-WGQGTQVTVSS |
| 7FAT | nanobody | QVQLQESGGGSVQAGGSLRLSCAAS-CDR1-CLGWFRQAPGKEREGVAAI-CDR2-TSYADSVKGRFTISRDNAKNTLYLQMNSLKPEDTAMYYCAA-CDR3-WGQGTQVTVSS |
| 3G9A | nanobody | DVQLQESGGGSVQAGGSLRLSCAAS-CDR1-SMAWFRQAPGKECELVSNI-CDR2-TTYAGSVKGRFTISRDDAKNTVYLQMVNLKSEDTARYYCAA-CDR3-WGKGTQVTVSS |
| 8Q95 | nanobody | QVQLVESGGDLVQSGGSLKLACAVS-CDR1-SIGWFRQAPGKEREAVSYS-CDR2-TYYVASVKGRFTISRDNAKNTAYLQMNNLKPEDTGIYYCAA-CDR3-WGQGTQVTVSS |
| 7ZW1 | nanobody | QVQLQESGGGLVQAGGSLRLSCAAS-CDR1-TVGWFCQAPGKEREFVAAV-CDR2-TWYADSVKGRFTISRDNAKNTVYLQMNSLKQEDTAVYYCAA-CDR3-WGQGTQVTVSS |
| 5VXK | nanobody | QLQLVESGGGLVQPGGSLRLSCAAS-CDR1-PIAWFRQAPGKEREGVSCI-CDR2-QSYSDSVKGRFTISRDTANNRVHLQMNNLKPEDTAVYYCAA-CDR3-WGQGTQVTVSS |
| 9G13 | nanobody | QVQLQASGGGFVQPGGSLRLSCAAS-CDR1-IMGWFRQAPGKEREFVSAI-CDR2-TYYADSVKGRFTISRDNSKNTVYLQMNSLRAEDTATYYCAP-CDR3-WGQGTQVTVSS |
| 8IEE | nanobody | QVQLQESGGGSVQTGGSLRLSCAGS-CDR1-SMGWYRQAPGRERELIGTI-CDR2-TYYSDAVKGRFTISLDNAKNTVYLQMNNLKPEDTAMYICNT-CDR3-WGQGTQVTVSS |
| 4N1H | nanobody | QVQLQESGGGLVQAGASLKLSCAAS-CDR1-AMGWFRQAPGKEREFVAAI-CDR2-TKYADSVKGRFAISRDNDKNTVWLRMNSLKPEDTAVYYCAA-CDR3-WGQGTQVTVSS |
| 7S2S | nanobody | QVQLQESGGGSVQAGGSLRLSCAAS-CDR1-CMGWFRQAPGKEREGVAGI-CDR2-TGYGDSVKGRFTISKDNAKNTLYLQMNSLKPEDTAMYYCAA-CDR3-WGQGTQVTVSS |
| 8ELO | nanobody | EVQLQESGGGLVQPGGSLRLSCAAS-CDR1-AMGWYRQAPGKEREWVCAI-CDR2-TYYADSVKGRFTCSRDNSKNTLYLQMNSLKPEDTAVYYCAR-CDR3-WGQGTQVTVSS |
| 8CYJ | nanobody | QVQLVESGGGLVQPGGSLRLSCAAS-CDR1-AMSWVRQAPGKGREWVSGI-CDR2-TYYTESVKGRFTISRDNAKNMLYLQMNSLKPEDTALYYCAK-CDR3-WGQGTQVTVSS |
| 8PIZ | nanobody | QLQLVESGGGLVQPGGSLRLSCASS-CDR1-AIVWFRQAPGKEREGVSCI-CDR2-TIYADSVKGRFTISRDNAKNTVYLQMNSLKPEDTAVYYCAA-CDR3-WGKGTLATVSS |
| 7NKT | nanobody | QVQLVESGGGSVQPGGSLRLSCLGS-CDR1-AIGWFRQAPGKEREGVSCI-CDR2-TIYADSVKGRFTISRDYGKNTVYLQMNSLKPEDTAMYYCAA-CDR3-WGQGAPVTVSS |
| 9G2A | nanobody | QVQLQESGGGLVQPGGSLRLSCAAS-CDR1-AIGWFRQAPGKEREGVSCI-CDR2-TYYADSVKGRFTISSDNAKNTVYLQMNSLKSEDTAVYYCAA-CDR3-WGQGTQVTVSS |
| 8G70 | nanobody | QVQLQESGGGLVQPGGSLRLSCVAS-CDR1-SMGWYRQAPGKQRELVAQI-CDR2-THYADSVKGRFTISEHRGKNAVYLEMHSLKPEDTAVYYCHL-CDR3-WGQGTQVTVSS |
| 8Q94 | nanobody | QVQLVETGGDLVQSGGSLRLACVLS-CDR1-SIGWFRQAPGKEREGISYS-CDR2-TYYVDSVKGRFTVSRDNAKNTAYLQMNSLKPEDSGIYYCAA-CDR3-WGQGTQVTVSS |
| 7ME7 | nanobody | VQLVESGGGLVQAGGSLRLSCAVS-CDR1-GMAWFRQAPGKERDFVATI-CDR2-TLYADSVKGRFTISRDNAKNTVYLQMNSLKIEDTAVYYCAV-CDR3-WGQGTQVTVS |
| 6XW6 | nanobody | QVQLQESGGGLVQAGDSLRVSCAAS-CDR1-PMGWFRQAPGKEREFVAAI-CDR2-TYYLDSVKGRFTTSRDNAKNTVYLQLNNLKPEDTAIYYCAA-CDR3-WGQGTQVTVSS |
| 4Y7M | nanobody | QVQLVESGGGLVQAGGSLRLSCAAS-CDR1-AIGWFRQAPGKEREGVSCI-CDR2-TYYPDSVKGRFTASSDKAKNMVYLQMNSLKPEDTAVYYCAA-CDR3-WGKGTQVTVSS |
| 6QX4 | nanobody | QVQLVESGGGLVQPGGSLRLSCAAS-CDR1-PMSWVRQAPGKGLEWVSDI-CDR2-TYYADSVKGRFTISRDNAKNTLYLQMNSLKPEDTAVYYCAT-CDR3-WGQGTQVTVSS |
| 4W6X | nanobody | QVQLQESGGGSVQAGGSLRLSCTAS-CDR1-CMGWFRQAPGKEREGVACI-CDR2-SYYADSVKGRFTISQDNAKDTVFLRMNSLKPEDTAIYYCAL-CDR3-WGQGTQVTVSS |
| 8OWT | nanobody | QVQLVESGGGLVQAGGSLRLSCAAS-CDR1-AMGWFRQAPGEERECVASI-CDR2-TWYADSVKGRFTISRDNPQNTVYLQMNNLKSGDTAVYYCAA-CDR3-WGQGIQVTVSS |
| 7ZML | nanobody | QVQLQESGGGLVQAGGSLRLSCAAS-CDR1-RMGWYRQAPGKQRELVAAI-CDR2-TNYAYSVKGRFTISRDNAKNTVYLQMNSLKPEDTAIYYCEA-CDR3-WGQGTQVTVSA |
| 4W6Y | nanobody | QVQLQESGGGSVQAGGSLRLSCAAS-CDR1-CMAWFRQVPGKEREGVASI-CDR2-TYYADSVKGRFTISRDNAKNTVSLQMNSLKPEDTATYYCAA-CDR3-WGQGTQVTVSS |

| PDB | Type | Sequence |
|---|---|---|
| 7NS6 | nanobody | QVQLVESGGGLVQPGGSLRLSCAAS-CDR1-AIGWFRQAPGKEREGVSFI-CDR2-TYYVDSVKGRFTISRDNAKNTVYLQMNSLTPEDTAIYYCAV-CDR3-WGQGTQVTVSS |
| 6OYH | nanobody | DVQLQESGGGLVQTGGSLTLSCATS-CDR1-AMAWFRQAPGKEREFVAGV-CDR2-TAYADAVKGRFTISRDNAANTVYLQMTSLKPEDTAVYFCAA-CDR3-WGQGTQVTVSS |
| 8DT8 | nanobody | EVQLQESGGGLVQPGGSLRLSCAAS-CDR1-ALGWYRQAPGKEREWVCAI-CDR2-TYYADSVKGRFTCSRDNSKNTLYLQMNSLKPEDTAVYYCAR-CDR3-WGQGTQVTVSS |
| 7P14 | nanobody | QVQLVESGGGSVQAGGSLRLSCAAS-CDR1-YLGWFRQAPGKEREGVAAL-CDR2-TYYADSVKGRFTVSLDNAKNTVYLQMNSLKPEDTALYYCAA-CDR3-WGQGTQVTVS |
| 7UST | nanobody | QVQLQESGGGLVQPGGSLRLSCAAS-CDR1-AIGWFRQAPGKEREGVSCI-CDR2-TYYADSVKGRFTISRDNAKNTVYLQMNSLKPEDTAVYYCAR-CDR3-WGKGTQVTVSS |
| 5E1H | nanobody | VQLAETGGGLVQAGGSLRLSCAAS-CDR1-AMAWFRQAPGKEREFVAGI-CDR2-TNYADSVKGRFTVSRDNAKNTVYLQMNSLKPEDTAVYYCAG-CDR3-WGQGTQVTVSS |
| 3EAK | nanobody | QVQLVESGGGLVQPGGSLRLSCAAS-CDR1-SLGWFRQAPGQGLEAVAAI-CDR2-TYYADSVKGRFTISRDNSKNTLYLQMNSLRAEDTAVYYCAA-CDR3-WGQGTLVTVSS |

**Supplementary Table 3.** CCD codes and sources of excluded ligands.

| Type | CCD codes | Source |
|---|---|---|
| Crystal aids | SO4, GOL, EDO, PO4, ACT, PEG, DMS, TRS, PGE, PG4, FMT, EPE, MPD, MES, CD, IOD | AlphaFold3 paper |
| Ions | 118, 119, 1AL, 1CU, 2FK, 2HP, 2OF, 3CO, 3MT, 3NI, 3OF, 4MO, 4PU, 4TI, 543, 6MO, AG, AL, ALF, AM, ATH, AU, AU3, AUC, BA, BEF, BF4, BO4, BR, BS3, BSY, CA, CAC, CD, CD1, CD3, CD5, CE, CF, CHT, CO, CO5, CON, CR, CS, CSB, CU, CU1, CU2, CU3, CUA, CUZ, CYN, DME, DMI, DSC, DTI, DY, E4N, EDR, EMC, ER3, EU, EU3, F, FE, FE2, FPO, GA, GD3, GEP, HAI, HG, HGC, HO3, IN, IR, IR3, IRI, IUM, K, KO4, LA, LCO, LCP, LI, LU, MAC, MG, MH2, MH3, MMC, MN, MN3, MN5, MN6, MO, MO1, MO2, MO3, MO4, MO5, MO6, MOO, MOS, MOW, MW1, MW2, MW3, NA2, NA5, NA6, NAO, NAW, NET, NI, NI1, NI2, NI3, NO2, NRU, O4M, OAA, OC1, OC2, OC3, OC4, OC5, OC6, OC7, OC8, OCL, OCM, OCN, OCO, OF1, OF2, OF3, OH, OS, OS4, OXL, PB, PBM, PD, PER, PI, PO3, PR, PT, PT4, PTN, RB, RH3, RHD, RU, SB, SE4, SEK, SM, SMO, SO3, T1A, TB, TBA, TCN, TEA, TH, THE, TL, TMA, TRA, V, VN3, VO4, W, WO5, Y1, YB, YB2, YH, YT3, ZCM, ZN, ZN2, ZN3, ZNO, ZO3, ZR | AlphaFold3 paper |
| AF3 excluded ligands | 144, 15P, 1PE, 2F2, 2JC, 3HR, 3SY, 7N5, 7PE, 9JE, AAE, ABA, ACE, ACN, ACT, ACY, AZI, BAM, BCN, BCT, BDN, BEN, BME, BO3, BTB, BTC, BU1, C8E, CAD, CAQ, CBM, CCN, CIT, CL, CLR, CM, CMO, CO3, CPT, CXS, D10, DEP, DIO, DMS, DN, DOD, DOX, EDO, EEE, EGL, EOH, EOX, EPE, ETF, FCY, FJO, FLC, FMT, FW5, GOL, GSH, GTT, GYF, HED, IHP, IHS, IMD, IOD, IPA, IPH, LDA, MB3, MEG, MES, MLA, MLI, MOH, MPD, MRD, MSE, MYR, N, NA, NH2, NH4, NHE, NO3, O4B, OHE, OLA, OLC, OMB, OME, OXA, P6G, PE3, PE4, PEG, PEO, PEP, PG0, PG4, PGE, PGR, PLM, PO4, POL, POP, PVO, SAR, SCN, SEO, SEP, SIN, SO4, SPD, SPM, SR, STE, STO, STU, TAR, TBU, TME, TPO, TRS, UNK, UNL, UNX, UPL, URE | AlphaFold3 paper |

| Type | CCD codes | Source |
|---|---|---|
| Glycans | 045, 05L, 07E, 07Y, 08U, 09X, 0BD, 0H0, 0HX, 0LP, 0MK, 0NZ, 0UB, 0V4, 0WK, 0XY, 0YT, 10M, 12E, 145, 147, 149, 14T, 15L, 16F, 16G, 16O, 17T, 18D, 18O, 1CF, 1FT, 1GL, 1GN, 1LL, 1S3, 1S4, 1SD, 1X4, 20S, 20X, 22O, 22S, 23V, 24S, 25E, 26O, 27C, 289, 291, 293, 2DG, 2DR, 2F8, 2FG, 2FL, 2GL, 2GS, 2H5, 2HA, 2M4, 2M5, 2M8, 2OS, 2WP, 2WS, 32O, 34V, 38J, 3BU, 3DO, 3DY, 3FM, 3GR, 3HD, 3J3, 3J4, 3LJ, 3LR, 3MG, 3MK, 3R3, 3S6, 3SA, 3YW, 40J, 42D, 445, 44S, 46D, 46Z, 475, 48Z, 491, 49A, 49S, 49T, 49V, 4AM, 4CQ, 4GC, 4GL, 4GP, 4JA, 4N2, 4NN, 4QY, 4R1, 4RS, 4SG, 4UZ, 4V5, 50A, 51N, 56N, 57S, 5GF, 5GO, 5II, 5KQ, 5KS, 5KT, 5KV, 5L3, 5LS, 5LT, 5MM, 5N6, 5QP, 5SP, 5TH, 5TJ, 5TK, 5TM, 61J, 62I, 64K, 66O, 6BG, 6C2, 6DM, 6GB, 6GP, 6GR, 6K3, 6KH, 6KL, 6KS, 6KU, 6KW, 6LA, 6LS, 6LW, 6MJ, 6MN, 6PZ, 6S2, 6UD, 6YR, 6ZC, 73E, 79J, 7CV, 7D1, 7GP, 7JZ, 7K2, 7K3, 7NU, 83Y, 89Y, 8B7, 8B9, 8EX, 8GA, 8GG, 8GP, 8I4, 8LR, 8OQ, 8PK, 8S0, 8YV 95Z, 96O, 98U, 9AM, 9C1, 9CD, 9GP, 9KJ, 9MR, 9OK, 9PG, 9QG, 9S7, 9SG, 9SJ, 9SM, 9SP, 9T1, 9T7, 9VP, 9WJ, 9WN, 9WZ, 9YW, A0K, A1Q, A2G, A5C, A6P, AAL, ABD, ABE, ABF, ABL, AC1, ACR, ACX, ADA, AF1, AFD, AFO, AFP, AGL, AH2, AH8, AHG, AHM, AHR, AIG, ALL, ALX, AMG, AMN, AMU, AMV, ANA, AOG, AQA, ARA, ARB, ARI, ARW, ASC, ASG, ASO, AXP, AXR, AY9, AZC, B0D, B16, B1H, B1N, B2G, B4G, B6D, B7G, B8D, B9D, BBK, BBV, BCD, BDF, BDG, BDP, BDR, BEM, BFN, BG6, BG8, BGC, BGL, BGN, BGP, BGS, BHG, BM3, BM7, BMA, BMX, BND, BNG, BNX, BO1, BOG, BQY, BS7, BTG, BTU, BW3, BWG, BXF, BXP, BXX, BXY, BZD, C3B, C3G, C3X, C4B, C4W, C5X, CBF, CBI, CBK, CDR, CE5, CE6, CE8, CEG, CEZ, CGF, CJB, CKB, CKP, CNP, CR1, CR6, CRA, CT3, CTO, CTR, CTT, D1M, D5E, D6G, DAF, DAG, DAN, DDA, DDL, DEG, DEL, DFR, DFX, DG0, DGO, DGS, DGU, DJB, DJE, DK4, DKX, DKZ, DL6, DLD, DLF, DLG, DNO, DO8, DOM, DPC, DQR, DR2, DR3, DR5, DRI, DSR, DT6, DVC, DYM, E3M, E5G, EAG, EBG, EBQ, EEN, EEQ, EGA, EMP, EMZ, EPG, EQP, EQV, ERE, ERI, ETT, EUS, F1P, F1X, F55, F58, F6P | Matched by unbranched hydrocarbon length |

| Type | CCD codes | Source |
|---|---|---|
| | F8X, FBP, FCA, FCB, FCT, FDP, FDQ, FFC, FFX, FIF, FK9, FKD, FMF, FMO, FNG, FNY, FRU, FSA, FSI, FSM, FSW, FUB, FUC, FUD, FUF, FUL, FUY, FVQ, FX1, FYJ, G0S, G16, G1P, G20, G28, G2F, G3F, G3I, G4D, G4S, G6D, G6P, G6S, G7P, G8Z, GAA, GAC, GAD, GAF, GAL, GAT, GBH, GC1, GC4, GC9, GCB, GCD, GCN, GCO, GCS, GCT, GCU, GCV, GCW, GDA, GDL, GE1, GE3, GFP, GIV, GL0, GL1, GL2, GL4, GL5, GL6, GL7, GL9, GLA, GLC, GLD, GLF, GLG, GLO, GLP, GLS, GLT, GM0, GMB, GMH, GMT, GMZ, GN1, GN4, GNS, GNX, GP0, GP1, GP4, GPH, GPK, GPM, GPO, GPQ, GPU, GPV, GPW, GQ1, GRF, GRX, GS1, GS9, GTK, GTM, GTR, GU0, GU1, GU2, GU3, GU4, GU5, GU6, GU8, GU9, GUF, GUL, GUP, GUZ, GXL, GXV, GYE, GYG, GYP, GYU, GYV, GZL, H1M, H1S, H2P, H3S, H53, H6Q, H6Z, HBZ, HD4, HNV, HNW, HSG, HSH, HSJ, HSQ, HSX, HSY, HTG, HTM, HVC, IAB, IDC, IDF, IDG, IDR, IDS, IDU, IDX, IDY, IEM, IN1, IPT, ISD, ISL, ISX, IXD, J5B, JFZ, JHM, JLT, JRV, JSV, JV4, JVA, JVS, JZR, K5B, K99, KBA, KBG, KD5, KDA, KDB, KDD, KDE, KDF, KDM, KDN, KDO, KDR, KFN, KG1, KGM, KHP, KME, KO1, KO2, KOT, KTU, L0W, L1L, L6S, L6T, LAG, LAH, LAI, LAK, LAO, LAT, LB2, LBS, LBT, LCN, LDY, LEC, LER, LFC, LFR | |

| Type | CCD codes | Source |
|---|---|---|
| | LGC, LGU, LKA, LKS, LM2, LMO, LNV, LOG, LOX, LRH, LTG, LVO, LVZ, LXB, LXC, LXZ, LZ0, M1F, M1P, M2F, M3M, M3N, M55, M6D, M6P, M7B, M7P, M8C, MA1, MA2, MA3, MA8, MAB, MAF, MAG, MAL, MAN, MAT, MAV, MAW, MBE, MBF, MBG, MCU, MDA, MDP, MFB, MFU, MG5, MGC, MGL, MGS, MJJ, MLB, MLR, MMA, MN0, MNA, MQG, MQT, MRH, MRP, MSX, MTT, MUB, MUR, MVP, MXY, MXZ, MYG, N1L, N3U, N9S, NA1, NAA, NAG, NBG, NBX, NBY, NDG, NFG, NG1, NG6, NGA, NGC, NGE, NGK, NGR, NGS, NGY, NGZ, NHF, NLC, NM6, NM9, NNG, NPF, NSQ, NT1, NTF, NTO, NTP, NXD, NYT, OAK, OI7, OPM, OSU, OTG, OTN, OTU, OX2, P53, P6P, P8E, PA1, PAV, PDX, PH5, PKM, PNA, PNG, PNJ, PNW, PPC, PRP, PSG, PSV, PTQ, PUF, PZU, QDK, QIF, QKH, QPS, QV4, R1P, R1X, R2B, R2G, RAE, RAF, RAM, RAO, RB5, RBL, RCD, RER, RF5, RG1, RGG, RHA, RHC, RI2, RIB, RIP, RM4, RP3, RP5, RP6, RR7, RRJ, RRY, RST, RTG, RTV, RUG, RUU, RV7, RVG, RVM, RWI, RY7, RZM, S7P, S81, SA0, SCG, SCR, SDY, SEJ, SF6, SF9, SFU, SG4, SG5, SG6, SG7, SGA, SGC, SGD, SGN, SHB, SHD, SHG, SIA, SID, SIO, SIZ, SLB, SLM, SLT, SMD, SN5, SNG, SOE, SOG, SOL, SOR, SR1, SSG, SSH, STW, STZ, SUC, SUP, SUS, SWE, SZZ, T68, T6D, T6P, T6T, TA6 TAG, TCB, TDG, TEU, TF0, TFU, TGA, TGK, TGR, TGY, TH1, TM5, TM6, TMR, TMX, TNX, TOA, TOC, TQY, TRE, TRV, TS8, TT7, TTV, TU4, TUG, TUJ, TUP, TUR, TVD, TVG, TVM, TVS, TVV, TVY, TW7, TWA, TWD, TWJ, TWY, TXB, TYV, U1Y, U2A, U2D, U63, U8V, U97, U9A, U9D, U9G, U9J, U9M, UAP, UBH, UBO, UDC, UEA, V3M, V3P, V71, VG1, VJ1, VJ4, VKN, VTB, W9T, WIA, WOO, WUN, WZ1, WZ2, X0X, X1P, X1X, X2F, X2Y, X34, X6X, X6Y, XDX, XGP, XIL, XKJ, XLF, XLS, XMM, XS2, XXM, XXR, XXX, XYF, XYL, XYP, XYS, XYT, XYZ, YDR, YIO, YJM, YKR, YO5, YX0, YX1, YYB, YYH, YYJ, YYK, YYM, YYQ, YZ0, Z0F, Z15, Z16, Z2D, Z2T, Z3K, Z3L, Z3Q, Z3U, Z4K, Z4R, Z4S, Z4U, Z4V, Z4W, Z4Y, Z57, Z5J, Z5L, Z61, Z6H, Z6J, Z6W, Z8H, Z8T, Z9D, Z9E, Z9H, Z9K, Z9L, Z9M, Z9W, ZB0, ZB1, ZB2, ZB3, ZCD, ZCZ, ZD0, ZDC, ZDO, ZEE, ZEL, ZGE, ZMR | |

| Type | CCD codes | Source |
|---|---|---|
| Non-biological | NUC, ZN, CA, MG, III, MN, FE, CU, SF4, FE2, CO, FES, GOL, NA, CL, K, CU1, GOL, XE, NO2, EDO, NI, BR, CD, O, CS, NO, TL, HG, UNL, KR, SR, RB, F, AG, AR, U, AU, MO, SE, GD, YB, VX, SM, LI, RE, N, W, OS, HO, PI, EDO, PG4, OGA, SO4, HEZ, FEO, CL, DMS, ACT, MPD, GOL, NH2, CUA, SIW, PGW, IOD, BR, 3NI, ZRW, 78M, UNX, MES, CCN, PO4 | RosettaFold paper |
| Excluded by Schrodinger | 1PE, 2HT, 2PE, 7PE, ACT, ACY, AKG, BCT, BMA, BME, BOG, BU3, BUD, CAC, CIT, CME, CO3, DMS, DTT, DTV, EDO, EGL, EPE, FES, FMT, FS3, FS4, GBL, GOL, GSH, HEC, HED, HEM, IMD, IOD, IPA, MAN, MES, MG8, MLI, MO6, MPD, MYR, NAG, NCO, NH3, NO3, OCT, OGA, OPG, P2U, PG4, PGE, PGO, PHO, PLP, PO4, POP, PSE, PSU, PTL, SEO, SGM, SO4, SPD, SPM, SRT, SUC, SUL, TAM, TAR, TFA, TLA, TPP, TRS | Maestro |
| Manually removed | ATP, ADP, AMP, GDP, UMP, PPV, PMP, DCP, TMP, ADE, GNP, TTP, DTP, NAP, NDP, CMP, BMP, DMP, SAM | - |

**Supplementary Table 4.** Dynamic batching configurations for training of each module, where $C$ denotes the predefined capacity threshold of each batch. The length of data is either measured by the number of atoms or the number of blocks depending on the operating space of the modules.

| Dataset | All-Atom VAE | | Latent Diffusion | | Confidence Module | |
|---|---|---|---|---|---|---|
| | $C$ | Length Type | $C$ | Length Type | $C$ | Length Type |
| BioLip2 | 12,000,000 | atom | 400,000 | block | 8,000,000 | atom |
| CrossDocked | 12,000,000 | atom | 800,000 | block | - | - |
| PDBbind v2020 | 12,000,000 | atom | 800,000 | block | 12,000,000 | atom |
| SIU | 12,000,000 | atom | 800,000 | block | - | - |
| ProtFrag | 12,000,000 | atom | 1,800,000 | block | - | - |
| PepBench | 12,000,000 | atom | 1,800,000 | block | 12,000,000 | atom |
| SAbDab | 12,000,000 | atom | 900,000 | block | 12,000,000 | atom |

**Supplementary Table 5.** Hyperparameters for AnewOmni.

| Name | Value | Description |
|---|---|---|
| | | Variational AutoEncoder |
| embed_size | 512 | Dimension of the atom and block type embeddings. |
| hidden_size | 512 | Dimension of hidden states. |
| latent_size | 8 | Dimension of latent states. |
| edge_size | 64 | Dimension of edge type embeddings. |
| n_rbf | 64 | Number of RBF kernels for embedding the spatial distances. |
| cutoff | 10.0 | Cutoff distance for RBF kernels. |
| n_layers | 6 | Number of layers in the encoder and the decoder. |
| n_head | 8 | Number of heads for multi-head attention. |
| $\lambda_1$ | 0.8 | The weight of KL divergence on the sequence. |
| $\lambda_2$ | 1.0 | The weight of KL divergence on the structure. |
| $\lambda_{\text{dist}}$ | 0.5 | The weight of local distance loss. |
| mask_ratio | 0.05 | The ratio of residues on the binding site for reconstruction. |
| N | 10 | Number of iterations for reconstruction. |
| | | Latent Diffusion Model |
| hidden_size | 512 | Dimension of hidden states. |
| T | 100 | Number of total steps for diffusion. |
| n_rbf | 64 | Number of RBF kernels for embedding the spatial distances. |
| cutoff | 3.0 | Cutoff distance for RBF kernels in the normalized latent space. |
| n_layers | 6 | Number of layers in the denoising network. |
| n_head | 8 | Number of heads for multi-head attention. |
| | | Supervised Confidence Model |
| hidden_size | 512 | Dimension of hidden states. |
| n_rbf | 64 | Number of RBF kernels for embedding the spatial distances. |
| cutoff | 10.0 | Cutoff distance for RBF kernels. |
| n_layers | 6 | Number of layers in the denoising network. |
| n_head | 8 | Number of heads for multi-head attention. |

**Supplementary Table 6.** IUPAC Names and SMILES for All Synthesized Small Molecules.

| Compound Name | IUPAC Name | SMILES |
|---|---|---|
| KRAS G12D-compound-1 | 3-(5-amino-2-chlorophenyl)-4-(1-(azetidin-3-ylamino)-1-oxopropan-2-yl)-N-(2,4-dihydroxyphenyl)isoxazole-5-carboxamide | O=C(C1=C(C(C)C(NC2CNC2)=O)C(C3=CC(N)=CC=C3Cl)=NO1)NC4=CC=C(O)C=C4O |
| KRAS G12D-compound-2 | cis-(5-amino-1-(3-bromo-4-(1,1-dioxido-3,4-dihydrobenzo[f][1,2,5]thiadiazepin-5(2H)-yl)phenyl)piperidin-3-yl)((R)-3,4-dimethylpiperazin-1-yl)methanone | O=C([C@H]1CN(C2=CC=C(N(C3=CC=CC=C34)CCNS4(=O)=O)C(Br)=C2)C[C@@H](N)C1)N5C[C@@H](C)N(C)CC5 |
| KRAS G12D-compound-3 | N-(3-aminophenyl)-7-(2-(2-chloro-5-hydroxyphenyl)-3-ethynyl-4,5,6,7-tetrahydrobenzo[b]thiophen-7-yl)-2-methyl-1,2,3,4-tetrahydroisoquinoline-4-carboxamide | O=C(C1CN(C)CC2=C1C=CC(C3CCCC4=C3SC(C5=CC(O)=CC=C5Cl)=C4C#C)=C2)NC6=CC=CC(N)=C6 |
| PCSK9-compound-1 | (R)-2-((4-amino-5-(trifluoromethyl)pyrimidin-2-yl)amino)-N-(4-(1-hydroxyethyl)phenyl)acetamide | O=C(NC1=CC=C([C@H](O)C)C=C1)CNC2=NC=C(C(F)(F)F)C(N)=N2 |
| PCSK9-compound-2 | 5-isopropyl-N-(3-(2-methylquinolin-3-yl)phenyl)thiazole-2-carboxamide | O=C(C1=NC=C(C(C)C)S1)NC2=CC=CC(C3=CC4=CC=CC=C4N=C3C)=C2 |
| PCSK9-compound-3 | (4-chloro-3-((5-cyclopropylpyrimidin-2-yl)amino)-2-methylphenyl)(3-chloro-5-(trifluoromethyl)phenyl)methanol | OC(C1=CC=C(Cl)C(NC2=NC=C(C3CC3)C=N2)=C1C)C4=CC(C(F)(F)F)=CC(Cl)=C4 |
| PCSK9-compound-4 | 5-(4-((8-methoxyquinazolin-2-yl)carbamoyl)phenyl)-1H-indole-2-carboxylicacid | O=C(C(N1)=CC2=C1C=CC(C3=CC=C(C(NC4=NC=C5C=CC=C(OC)C5=N4)=O)C=C3)=C2)O |
| PCSK9-compound-5 | 4-(piperidin-1-yl)-N-(quinazolin-2-yl)benzamide | O=C(NC1=NC=C2C=CC=CC2=N1)C3=CC=C(N4CCCCC4)C=C3 |
| PCSK9-compound-6 | N-(3-(2-aminoethyl)-4-methyl-2-oxo-2H-chromen-6-yl)-5-(benzo[d]oxazol-2-yl)-4H-1,2,4-triazole-3-carboxamide | O=C(C1=NN=C(C2=NC3=CC=CC=C3O2)N1)NC4=CC(C(C)=C(CCN)C5=O)=C(O5)C=C4 |
| PCSK9-compound-7 | N-[2-amino-4-(3-methoxyphenoxy)phenyl]-4-chloropyridine-2-carboxamide | O=C(C1=NC=CC(Cl)=C1)NC2=CC=C(OC3=CC=CC(OC)=C3)C=C2N |
| PCSK9-compound-8 | 1-(6-(3-(3-bromophenyl)ureido)pyridin-2-yl)-1H-benzo[d]imidazole-4-carboxamide | O=C(C1=C2C(N(C3=NC(NC(NC4=CC=CC(Br)=C4)=O)=CC=C3)C=N2)=CC=C1)N |
| PCSK9-compound-9 | N-[3-([(3R)-1-[(2R)-2-aminopropanoyl]pyrrolidin-3-yl]oxymethyl)phenyl]-4-methylpyridine-2-carboxamide | O=C(C1=NC=CC(C)=C1)NC2=CC=CC(CO[C@H]3CN(C([C@H](N)C)=O)CC3)=C2 |

**Supplementary Table 7.** Sequences of tested cyclic and linear peptides for KRAS G12D.

| ID | Sequence | ID | Sequence |
|---|---|---|---|
| KRAS G12D-cycpep-1 | cyclo[CYPWMRWRRC] | KRAS G12D-linpep-6 | QEHRHTIVYQWQ |
| KRAS G12D-cycpep-2 | cyclo[CWHYYSEYKC] | KRAS G12D-linpep-7 | YYYLRERFV |
| KRAS G12D-cycpep-3 | cyclo[CERWYVEHC] | KRAS G12D-linpep-8 | EPFLYERHIR |
| KRAS G12D-cycpep-4 | cyclo[CYWKYVDRRC] | KRAS G12D-linpep-9 | RWTWEFRH |
| KRAS G12D-cycpep-5 | cyclo[CTWFFKRQC] | KRAS G12D-linpep-10 | HYQRWTLWGRE |
| KRAS G12D-cycpep-6 | cyclo[CNRWWRDVSLC] | KRAS G12D-linpep-11 | TWRRWDRG |
| KRAS G12D-cycpep-7 | cyclo[CAWHFRYHKC] | KRAS G12D-linpep-12 | YIYFAVRRRVE |
| KRAS G12D-cycpep-8 | cyclo[CYFGRYDRYIC] | KRAS G12D-linpep-13 | FRWKGLYRL |
| KRAS G12D-cycpep-9 | cyclo[CVNERWLRVYRC] | KRAS G12D-linpep-14 | RVQQRPRW |
| KRAS G12D-cycpep-10 | cyclo[CSIYRIRYDQHC] | KRAS G12D-linpep-15 | KQFMFIKY |
| KRAS G12D-cycpep-11 | cyclo[CFRWKELKLC] | KRAS G12D-linpep-16 | EFRQRFPQY |
| KRAS G12D-cycpep-12 | cyclo[CQFRQQHRHC] | KRAS G12D-linpep-17 | PHFLYLYRP |
| KRAS G12D-cycpep-13 | cyclo[CHETWSKWFVC] | KRAS G12D-linpep-18 | YYLFRERH |
| KRAS G12D-cycpep-14 | cyclo[CHAVRWWRC] | KRAS G12D-linpep-19 | YMEWLWEK |
| KRAS G12D-cycpep-15 | cyclo[CPLWYKRKC] | KRAS G12D-linpep-20 | KFYSTFPRY |
| KRAS G12D-cycpep-16 | cyclo[CEHFIHREIC] | KRAS G12D-linpep-21 | RFNRRRYKVW |
| KRAS G12D-cycpep-17 | cyclo[CSRSFFWRC] | KRAS G12D-linpep-22 | ERLWATFRR |
| KRAS G12D-cycpep-18 | cyclo[CYLERTVRWEC] | KRAS G12D-linpep-23 | THEWWFLE |
| KRAS G12D-cycpep-19 | cyclo[CWTPWDDRIC] | KRAS G12D-linpep-24 | HWFWAFQQT |
| KRAS G12D-cycpep-20 | cyclo[CVYIEQYSQRC] | KRAS G12D-linpep-25 | VRYDYLWRPFI |
| KRAS G12D-linpep-1 | KQQFRFFRYEL | KRAS G12D-linpep-26 | YWKTYFFNYT |
| KRAS G12D-linpep-2 | RAWWRWRKNT | KRAS G12D-linpep-27 | QYPRWVWD |
| KRAS G12D-linpep-3 | YYRYFELLGR | KRAS G12D-linpep-28 | YDEYIWKSRQ |
| KRAS G12D-linpep-4 | YSHWWYHIT | KRAS G12D-linpep-29 | FWQNRRWKFLA |
| KRAS G12D-linpep-5 | EYRISFEYREQ | KRAS G12D-linpep-30 | WYEHDRLDYIAR |

**Supplementary Table 8.** Sequences of experimentally tested peptides for PCSK9.

| ID | Sequence |
|---|---|
| PCSK9-peptide-1 | WHRRFQYKRQ |
| PCSK9-peptide-2 | DNRWWTKQWRH |
| PCSK9-peptide-3 | WWWVKWRMFE |
| PCSK9-peptide-4 | RQRYTWRRYY |
| PCSK9-peptide-5 | FLHDWWTWRI |
| PCSK9-peptide-6 | HKYWEWIFEKY |
| PCSK9-peptide-7 | RLEYVRESWHYD |

**Supplementary Table 9.** Sequences, selection strategies, and PDB IDs of frameworks of experimentally tested nanobodies for KRAS G12D.

| ID | Sequence | Strategy | Framework |
|---|---|---|---|
| KRAS G12D-nanobody-1 | QVQLQESGGGSVQAGGSLRLSCAASGYQYSLL CMAWFRQVLGEGREGVAFITTYNGAMRYAD TVKGRFTVSQDKDKNTVYLQMNSLKPEDT AIYYCAAREQGGSEGYEDTSDMYDWGQGT QVTVSS | scRMSD | 3JBE |
| KRAS G12D-nanobody-2 | QVQLQESGGGSVQAGGSLRLSCAASGYQYSLL CMAWFRQVLGEGREGVAFITTYNGAMRYAD TVKGRFTVSQDKDKNTVYLQMNSLKPEDT AIYYCAADQAGGLSPPPTTKVYARWGQGT QVTVSS | scRMSD | 3JBE |
| KRAS G12D-nanobody-3 | QVQLQESGGGSVQAGGSLRLSCAASGYQYSLL CMAWFRQVLGEGREGVAFITTYNGAMRYAD TVKGRFTVSQDKDKNTVYLQMNSLKPEDT AIYYCAARSDYPGYGVASPLEMEYWGQGT QVTVSS | scRMSD | 3JBE |
| KRAS G12D-nanobody-4 | QVQLQESGGGSVQAGGSLRLSCAASGYQYSLL CMAWFRQVLGEGREGVAFITTYNGAMRYAD TVKGRFTVSQDKDKNTVYLQMNSLKPEDT AIYYCAARQCLYPPVSSSNEAMDDWGQGT QVTVSS | scRMSD | 3JBE |
| KRAS G12D-nanobody-5 | QVQLQESGGGSVQAGGSLRLSCTASGYTYRKY CMGWFRQAPGKEREGVACINSGGGTSYYAD SVKGRFTISQDNAKDTVFLRMNSLKPEDTA IYYCALMARYSTSWMEWSHS-DTWGQGTQVT VSS | likelihood | 4W6X |
| KRAS G12D-nanobody-6 | QVQLQESGGGSVQAGGSLRLSCTASGYTYRKY CMGWFRQAPGKEREGVACINSGGGTSYYAD SVKGRFTISQDNAKDTVFLRMNSLKPEDTA IYYCALIAQPWYGTEPASHAHSWGQGTQVT VSS | likelihood | 4W6X |
| KRAS G12D-nanobody-7 | QVQLQESGGGSVQAGGSLRLSCTASGYTYRKY CMGWFRQAPGKEREGVACINSGGGTSYYAD SVKGRFTISQDNAKDTVFLRMNSLKPEDTA IYYCALEYRIFGSYSVSPSSDYWGQGTQVT VSS | likelihood | 4W6X |

**Supplementary Table 10.** Overall rankings for *de novo* small molecule design.

| Model | substruct. 0.2 | Chem. 0.2 | Interact. 0.4 | Geom. 0.2 | Weighted Score | Rank |
|---|---|---|---|---|---|---|
| LIGAN | 1.13 | 1.40 | **4.27** | 1.25 | 8.05 | 6 |
| 3DSBDD | 1.13 | 1.60 | 2.23 | 0.70 | 5.67 | 9 |
| GraphBP | 0.17 | 1.50 | 0.37 | 0.10 | 2.13 | 14 |
| Pocket2Mol | 0.73 | 1.25 | 2.83 | 0.70 | 5.52 | 10 |
| TargetDiff | 1.77 | 1.50 | 3.50 | 1.70 | 8.47 | 5 |
| DiffSBDD | 0.77 | 1.75 | 1.20 | 0.95 | 4.67 | 12 |
| DiffBP | 0.27 | 1.10 | 2.10 | 1.35 | 4.82 | 11 |
| FLAG | 0.70 | 1.40 | 1.40 | 0.60 | 4.10 | 13 |
| D3FG | 1.47 | **2.25** | 1.80 | 0.70 | 6.22 | 8 |
| DecompDiff | 1.90 | 1.80 | 2.50 | 1.80 | 8.00 | 7 |
| MolCRAFT | 1.93 | 1.55 | 3.93 | **2.20** | 9.62 | 2 |
| VoxBind | 1.53 | 2.00 | 3.83 | 2.00 | 9.37 | 3 |
| Ours (mol) | 2.23 | 2.15 | 2.70 | 1.95 | 9.03 | 4 |
| Ours (all) | **2.27** | **2.25** | 3.47 | **2.20** | **10.38** | **1** |

**Supplementary Table 11.** Substructure analysis for *de novo* small molecule design.

| Model | Atom Type | | Ring Type | | Functional Group | | Rank |
|---|---|---|---|---|---|---|---|
| | JSD | MAE | JSD | MAE | JSD | MAE | |
| LIGAN | 0.1167 | 0.8680 | 0.3163 | 0.2701 | 0.2468 | 0.0378 | 8.33 |
| 3DSBDD | 0.0860 | 0.8444 | 0.3188 | 0.2457 | 0.2682 | 0.0494 | 8.33 |
| GraphBP | 0.1642 | 1.2266 | 0.5061 | 0.4382 | 0.6259 | 0.0705 | 13.17 |
| Pocket2Mol | 0.0916 | 1.0497 | 0.3550 | 0.3545 | 0.2961 | 0.0622 | 10.33 |
| TargetDiff | 0.0533 | 0.2399 | 0.2345 | 0.1559 | 0.2876 | 0.0441 | 5.17 |
| DiffSBDD | 0.0529 | 0.6316 | 0.3853 | 0.3437 | 0.5520 | 0.0710 | 10.17 |
| DiffBP | 0.2591 | 1.5491 | 0.4531 | 0.4068 | 0.5346 | 0.0670 | 12.67 |
| FLAG | 0.1032 | 1.7665 | 0.2432 | 0.3370 | 0.3634 | 0.0666 | 10.50 |
| D3FG | 0.0644 | 0.8154 | 0.1869 | 0.2204 | 0.2511 | 0.0516 | 6.67 |
| DecompDiff | 0.0431 | 0.3197 | 0.2431 | 0.2006 | 0.1916 | 0.0318 | 4.50 |
| MolCRAFT | 0.0490 | 0.3208 | 0.2469 | **0.0264** | 0.1196 | 0.0477 | 4.33 |
| VoxBind | 0.0942 | 0.3564 | 0.2401 | 0.0301 | **0.1053** | 0.0761 | 6.33 |
| Ours (mol) | 0.0352 | 0.2796 | 0.1252 | 0.0976 | 0.1562 | 0.0173 | 2.83 |
| Ours (all) | **0.0280** | **0.2108** | **0.1073** | 0.0726 | 0.1356 | **0.0161** | **1.67** |

**Supplementary Table 12.** Results of chemical properties for *de novo* small molecule design. LogP between -0.4 and 5.6 are all reasonable[28], without preference to higher or lower values.

| Model | QED | LogP | SA | LPSK | Rank |
|---|---|---|---|---|---|
| LIGAN | 0.46 | 0.56 | 0.66 | 4.39 | 7.00 |
| 3DSBDD | 0.48 | 0.47 | 0.63 | 4.72 | 6.00 |
| GraphBP | 0.44 | 3.29 | 0.64 | 4.73 | 6.50 |
| Pocket2Mol | 0.39 | 2.39 | 0.65 | 4.58 | 7.75 |
| TargetDiff | 0.49 | 1.13 | 0.60 | 4.57 | 6.50 |
| DiffSBDD | 0.49 | -0.15 | 0.34 | 4.89 | 5.25 |
| DiffBP | 0.47 | 5.27 | 0.59 | 4.47 | 8.50 |
| FLAG | 0.41 | 0.29 | 0.58 | **4.93** | 7.00 |
| D3FG | 0.49 | 1.56 | 0.66 | 4.84 | **2.75** |
| DecompDiff | 0.49 | 1.22 | 0.66 | 4.40 | 5.00 |
| MolCRAFT | 0.48 | 0.87 | 0.66 | 4.39 | 6.25 |
| VoxBind | 0.54 | 2.22 | 0.65 | 4.70 | 4.00 |
| Ours (mol) | 0.53 | 1.36 | 0.68 | 4.69 | 3.25 |
| Ours (all) | **0.55** | 1.55 | **0.70** | 4.68 | **2.75** |

**Supplementary Table 13.** Geometry analysis for *de novo* small molecule design.

| Model | Static Geometry | | Clash | | Rank |
|---|---|---|---|---|---|
| | $JSD_{BL}$ | $JSD_{BA}$ | $Ratio_{cca}$ | $Ratio_{cm}$ | |
| LIGAN | 0.4645 | 0.5673 | 0.0096 | 0.0718 | 7.75 |
| 3DSBDD | 0.5024 | 0.3904 | 0.2482 | 0.8683 | 10.50 |
| GraphBP | 0.5182 | 0.5645 | 0.8634 | 0.9974 | 13.50 |
| Pocket2Mol | 0.5433 | 0.4922 | 0.0576 | 0.4499 | 10.50 |
| TargetDiff | 0.2659 | 0.3769 | 0.0483 | 0.4920 | 5.50 |
| DiffSBDD | 0.3501 | 0.4588 | 0.1083 | 0.6578 | 9.25 |
| DiffBP | 0.3453 | 0.4621 | 0.0449 | 0.4077 | 7.25 |
| FLAG | 0.4215 | 0.4304 | 0.6777 | 0.9769 | 11.00 |
| D3FG | 0.3727 | 0.4700 | 0.2115 | 0.8571 | 10.50 |
| DecompDiff | 0.2576 | 0.3473 | 0.0462 | 0.5248 | 5.00 |
| MolCRAFT | **0.2250** | **0.2683** | 0.0264 | 0.2691 | **3.00** |
| VoxBind | 0.2701 | 0.3771 | 0.0103 | 0.1890 | 4.00 |
| Ours (mol) | 0.3362 | 0.4166 | 0.0042 | 0.0715 | 4.25 |
| Ours (all) | 0.3223 | 0.3848 | **0.0040** | **0.0708** | **3.00** |

**Supplementary Table 14.** Results of interaction analysis for *de novo* small molecule design.

| Model | Vina Score | | Vina Min | | Vina Dock | | | | PLIP Interaction | | | | Rank |
|---|---|---|---|---|---|---|---|---|---|---|---|---|---|
| | E | IMP (%) | E | IMP (%) | E | IMP (%) | MPBG (%) | LBE | $JSD_{OA}$ | $MAE_{OA}$ | $JSD_{PP}$ | $MAE_{PP}$ | |
| LIGAN | **-6.47** | **62.13** | **-7.14** | **70.18** | **-7.70** | **72.71** | 4.22 | 0.3897 | 0.0346 | 0.0905 | 0.1451 | **0.3416** | **3.33** |
| 3DSBDD | - | 3.99 | -3.75 | 17.98 | -6.45 | 31.46 | 9.18 | 0.3839 | 0.0392 | 0.0934 | 0.1733 | 0.4231 | 8.42 |
| GraphBP | - | 0.00 | - | 1.67 | -4.57 | 10.86 | -30.03 | 0.3200 | 0.0462 | 0.1625 | 0.2101 | 0.4835 | 13.08 |
| Pocket2Mol | -5.23 | 31.06 | -6.03 | 38.04 | -7.05 | 48.07 | -0.17 | **0.4115** | 0.0319 | 0.2455 | 0.1535 | 0.4152 | 6.92 |
| TargetDiff | -5.71 | 38.21 | -6.43 | 47.09 | -7.41 | 51.99 | 5.38 | 0.3537 | 0.0198 | 0.0600 | 0.1757 | 0.4687 | 5.25 |
| DiffSBDD | - | 12.67 | -2.15 | 22.24 | -5.53 | 29.76 | -23.51 | 0.2920 | 0.0333 | 0.1461 | 0.1777 | 0.5265 | 11.00 |
| DiffBP | - | 8.60 | - | 19.68 | -7.34 | 49.24 | 6.23 | 0.3481 | 0.0249 | 0.1430 | **0.1256** | 0.5639 | 8.75 |
| FLAG | - | 0.04 | - | 3.44 | -3.65 | 11.78 | -47.64 | 0.3319 | 0.0170 | 0.0277 | 0.2762 | 0.3976 | 10.50 |
| D3FG | - | 3.70 | -2.59 | 11.13 | -6.78 | 28.9 | -8.85 | 0.4009 | 0.0638 | **0.0135** | 0.1850 | 0.4641 | 9.50 |
| DecompDiff | -5.18 | 19.66 | -6.04 | 34.84 | -7.10 | 48.31 | -1.59 | 0.3460 | 0.0215 | 0.0769 | 0.1848 | 0.4369 | 7.75 |
| MolCRAFT | -6.15 | 54.25 | -6.99 | 56.43 | -7.79 | 56.22 | 8.38 | 0.3638 | 0.0214 | 0.0780 | 0.1868 | 0.4574 | 4.17 |
| VoxBind | -6.16 | 41.80 | -6.82 | 50.02 | -7.68 | 52.91 | **9.89** | 0.3588 | 0.0257 | 0.0533 | 0.1850 | 0.4606 | 4.42 |
| Ours (mol) | -5.50 | 26.37 | -5.96 | 38.21 | -7.19 | 49.60 | 2.15 | 0.3521 | 0.0175 | 0.0447 | 0.2372 | 0.4706 | 7.25 |
| Ours (all) | -5.72 | 30.40 | -6.08 | 39.23 | -7.25 | 51.59 | 7.50 | 0.3473 | **0.0135** | 0.0173 | 0.2012 | 0.4277 | 5.33 |

**Supplementary Table 15.** Non canonical targets used for testing AnewOmni.

| **Type** | **PDB ID** |
|---|---|
| DNA/RNA | 1QD3, 1PBR, 2G5K, 2Z74, 1AJU, 3GOG, 2KTZ, 407D, 2FD0, 3C44, 3DVV, 1DB6, 3OWZ, 2Z75, 1NBK, 1HNW, 3Q50, 1U8D, 1R4E, 1Q8N, 3GAO, 2YDH, 2BEE, 3Q3Z, 1J7T, 4ERJ, 2O3W, 1KOC, 1P96, 408D, 2O3X, 1KOD, 3DS7, 2ET8, 1ARJ, 2FCY, 1NTA, 1O9M, 2AU4, 3NPQ, 1YKV, 4KQY, 1YRJ, 2FCZ, 3E5C, 3GX5, 2JUK, 3S4P, 3SUX, 1BYJ, 2ET3, 1UUI, 1EI2, 2G9C, 1QV8, 3GX7, 3FU2, 1CVY, 1F27, 2ET4, 1Y26, 2KU0, 2B57, 1MWL, 1Y27, 1XPF, 3NPN, 2O3V, 2XO0, 1UUD, 3GER, 2BE0, 1LC4, 4JF2, 2JWQ, 3GOT, 2OGN, 2OE8, 1F1T, 3GX2, 2XO1, 3GES, 1UTS, 1QV4, 2FCX, 2ESJ, 1AM0, 3GX3, 1FYP, 3LA5, 1NZM, 2LOA, 1NEM, 2L94, 1NYI, 3F2Q, 2D55, 2GIS, 2TOB, 3SD3, 316D, 2XNZ, 1CVX, 4AOB, 2OE5, 3MUR, 2GDI, 3FO4, 1TOB, 4FE5, 2F4S, 3SLM, 3G4M, 1FJG, 1O15, 2MB3, 2F4T, 1I9V, 1LVJ, 1FMN, 3MXH, 2KX8, 3D2X, 2ESI, 3FO6, 1XBP, 3MUM, 2F4U, 1NTB, 1ZZ5, 2G5Q, 2KGP, 1AKX, 4LVV, 3DIL, 2PWT, 2XNW, 1EHT, 2YGH |
| Post-translational | 1U7V |
| Glycan | 6HA0 |

3. Chengxin Zhang, Xi Zhang, Lydia Freddolino, and Yang Zhang. Biolip2: an updated structure database for biologically relevant ligand–protein interactions. *Nucleic Acids Research*, 52(D1):D404–D412, 2024.

4. Renxiao Wang, Xueliang Fang, Yipin Lu, and Shaomeng Wang. The pdbbind database: Collection of binding affinities for protein- ligand complexes with known three-dimensional structures. *Journal of medicinal chemistry*, 47(12):2977–2980, 2004.

5. Xingang Peng, Shitong Luo, Jiaqi Guan, Qi Xie, Jian Peng, and Jianzhu Ma. Pocket2mol: Efficient molecular sampling based on 3d protein pockets. In *International conference on machine learning*, pages 17644–17655. PMLR, 2022.

6. Yanwen Huang, Bowen Gao, Yinjun Jia, Hongbo Ma, Wei-Ying Ma, Ya-Qin Zhang, and Yanyan Lan. Siu: A million-scale structural small molecule-protein interaction dataset for unbiased bioactivity prediction. *arXiv preprint arXiv:2406.08961*, 2024.

7. Xiangzhe Kong, Yinjun Jia, Wenbing Huang, and Yang Liu. Full-atom peptide design with geometric latent diffusion. *Advances in Neural Information Processing Systems*, 37:74808–74839, 2024.

8. Xiangzhe Kong, Rui Jiao, Haowei Lin, Ruihan Guo, Wenbing Huang, Wei-Ying Ma, Zihua Wang, Yang Liu, and Jianzhu Ma. Peptide design through binding interface mimicry with pepmimic. *Nature biomedical engineering*, pages 1–16, 2025.

9. Xiangzhe Kong, Wenbing Huang, and Yang Liu. Conditional antibody design as 3d equivariant graph translation. In *The Eleventh International Conference on Learning Representations*, 2023.

10. Xiangzhe Kong, Zishen Zhang, Ziting Zhang, Rui Jiao, Jianzhu Ma, Wenbing Huang, Kai Liu, and Yang Liu. Unimomo: Unified generative modeling of 3d molecules for de novo binder design. In *Forty-second International Conference on Machine Learning*, 2025.

11. James Dunbar, Konrad Krawczyk, Jinwoo Leem, Terry Baker, Angelika Fuchs, Guy Georges, Jiye Shi, and Charlotte M Deane. Sabdab: the structural antibody database. *Nucleic acids research*, 42(D1):D1140–D1146, 2014.

12. Jared Adolf-Bryfogle, Oleks Kalyuzhniy, Michael Kubitz, Brian D Weitzner, Xiaozhen Hu, Yumiko Adachi, William R Schief, and Roland L Dunbrack Jr. Rosettaantibodydesign (rabd): A general framework for computational antibody design. *PLoS computational biology*, 14(4):e1006112, 2018.

**13.** Pascal Vincent, Hugo Larochelle, Isabelle Lajoie, Yoshua Bengio, Pierre-Antoine Manzagol, and Léon Bottou. Stacked denoising autoencoders: Learning useful representations in a deep network with a local denoising criterion. *Journal of machine learning research*, 11(12), 2010.

**14.** Diederik P Kingma and Max Welling. Auto-encoding variational bayes. *arXiv preprint arXiv:1312.6114*, 2013.

**15.** Yaron Lipman, Ricky TQ Chen, Heli Ben-Hamu, Maximilian Nickel, and Matthew Le. Flow matching for generative modeling. In *The Eleventh International Conference on Learning Representations*, 2023.

**16.** Dan Hendrycks and Kevin Gimpel. Gaussian error linear units (gelus). *arXiv preprint arXiv:1606.08415*, 2016.

**17.** Rui Jiao, Xiangzhe Kong, Ziyang Yu, Wenbing Huang, and Yang Liu. Equivariant pretrained transformer for unified geometric learning on multi-domain 3d molecules. In *ICLR 2024 Workshop on Generative and Experimental Perspectives for Biomolecular Design*, 2024.

**18.** Shitong Luo, Yufeng Su, Xingang Peng, Sheng Wang, Jian Peng, and Jianzhu Ma. Antigen-specific antibody design and optimization with diffusion-based generative models for protein structures. *Advances in Neural Information Processing Systems*, 35:9754–9767, 2022.

**19.** Alexander Quinn Nichol and Prafulla Dhariwal. Improved denoising diffusion probabilistic models. In *International Conference on Machine Learning*, pages 8162–8171. PMLR, 2021.

**20.** Jonathan Ho, Ajay Jain, and Pieter Abbeel. Denoising diffusion probabilistic models. *Advances in neural information processing systems*, 33:6840–6851, 2020.

**21.** Jonathan Ho and Tim Salimans. Classifier-free diffusion guidance. *arXiv preprint arXiv:2207.12598*, 2022.

**22.** Xiangzhe Kong, Yinjun Jia, Wenbing Huang, and Yang Liu. Full-atom peptide design with geometric latent diffusion. *Advances in Neural Information Processing Systems*, 37:74808–74839, 2024.

**23.** Srinivas Ramachandran, Pradeep Kota, Feng Ding, and Nikolay V Dokholyan. Automated minimization of steric clashes in protein structures. *Proteins: Structure, Function, and Bioinformatics*, 79(1):261–270, 2011.

**24.** Noel M O'Boyle, Michael Banck, Craig A James, Chris Morley, Tim Vandermeersch, and Geoffrey R Hutchison. Open babel: An open chemical toolbox. *Journal of cheminformatics*, 3:1–14, 2011.

**25.** Yang Song, Jascha Sohl-Dickstein, Diederik P Kingma, Abhishek Kumar, Stefano Ermon, and Ben Poole. Score-based generative modeling through stochastic differential equations. In *International Conference on Learning Representations*, 2021.

**26.** John Skilling. *The Eigenvalues of Mega-dimensional Matrices*, pages 455–466. Springer Netherlands, Dordrecht, 1989.

**27.** M.F. Hutchinson. A stochastic estimator of the trace of the influence matrix for laplacian smoothing splines. *Communications in Statistics - Simulation and Computation*, 19(2):433–450, 1990.

**28.** Haitao Lin, Guojiang Zhao, Odin Zhang, Yufei Huang, Lirong Wu, Zicheng Liu, Siyuan Li, Cheng Tan, Zhifeng Gao, and Stan Z Li. Cbgbench: Fill in the blank of protein-molecule complex binding graph. *arXiv preprint arXiv:2406.10840*, 2024.

**29.** Tomer Tsaban, Julia K Varga, Orly Avraham, Ziv Ben-Aharon, Alisa Khramushin, and Ora Schueler-Furman. Harnessing protein folding neural networks for peptide–protein docking. *Nature communications*, 13(1):176, 2022.

**30.** Xiangzhe Kong, Wenbing Huang, and Yang Liu. Conditional antibody design as 3d equivariant graph translation. In *The Eleventh International Conference on Learning Representations*, 2023.

**31.** Jiahan Li, Chaoran Cheng, Zuofan Wu, Ruihan Guo, Shitong Luo, Zhizhou Ren, Jian Peng, and Jianzhu Ma. Full-atom peptide design based on multi-modal flow matching. In *Forty-first International Conference on Machine Learning*, 2024.

**32.** Saul B Needleman and Christian D Wunsch. A general method applicable to the search for similarities in the amino acid sequence of two proteins. *Journal of molecular biology*, 48(3):443–453, 1970.

**33.** Steven Henikoff and Jorja G Henikoff. Amino acid substitution matrices from protein blocks. *Proceedings of the National Academy of Sciences*, 89(22):10915–10919, 1992.

**34.** Rebecca F Alford, Andrew Leaver-Fay, Jeliazko R Jeliazkov, Matthew J O'Meara, Frank P DiMaio, Hahnbeom Park, Maxim V Shapovalov, P Douglas Renfrew, Vikram K Mulligan, Kalli Kappel, et al. The rosetta all-atom energy function for macromolecular modeling and design. *Journal of chemical theory and computation*, 13(6):3031–3048, 2017.

**35.** Fei Ye, Zaixiang Zheng, Dongyu Xue, Yuning Shen, Lihao Wang, Yiming Ma, Yan Wang, Xinyou Wang, Xiangxin Zhou, and Quanquan Gu. Proteinbench: A holistic evaluation of protein foundation models. *arXiv preprint arXiv:2409.06744*, 2024.

**49.** Todd J Dolinsky, Paul Czodrowski, Hui Li, Jens E Nielsen, Jan H Jensen, Gerhard Klebe, and Nathan A Baker. Pdb2pqr: expanding and upgrading automated preparation of biomolecular structures for molecular simulations. *Nucleic acids research*, 35(suppl_2):W522–W525, 2007.

**69.** Yuxuan Zhang, Chengyue Gong, Hanyu Zhang, Wenzhi Ma, Zhenyu Liu, Xinshi Chen, Jiaqi Guan, Lan Wang, Yanping Yang, Yu Xia, and Wenzhi Xiao. Protenix-v1: Toward high-accuracy open-source biomolecular structure prediction. *bioRxiv*, 2026.

**70.** Xiaolun Wang, Shelley Allen, James F Blake, Vickie Bowcut, David M Briere, Andrew Calinisan, Joshua R Dahlke, Jay B Fell, John P Fischer, Robin J Gunn, et al. Identification of mrtx1133, a noncovalent, potent, and selective krasg12d inhibitor. *Journal of medicinal chemistry*, 65(4):3123–3133, 2021.

**71.** Mikimasa Tanada, Minoru Tamiya, Atsushi Matsuo, Aya Chiyoda, Koji Takano, Toshiya Ito, Machiko Irie, Tomoya Kotake, Ryuuichi Takeyama, Hatsuo Kawada, et al. Development of orally bioavailable peptides targeting an intracellular protein: from a hit to a clinical kras inhibitor. *Journal of the American Chemical Society*, 145(30):16610–16620, 2023.

**72.** Samuel Genheden and Ulf Ryde. The mm/pbsa and mm/gbsa methods to estimate ligand-binding affinities. *Expert opinion on drug discovery*, 10(5):449–461, 2015.
